# *In vitro* selection of cyclized, glycosylated peptide antigens that tightly bind HIV high mannose patch antibodies

**DOI:** 10.1101/2025.03.24.645033

**Authors:** Jennifer K. Bailey, Satoru Horiya, Mahesh Neralkar, Viktor Horvath, Kosuke Nakamoto, J. Sebastian Temme, Raphael J. Turra, Isaac J. Krauss

## Abstract

*In vitro* selection is typically limited to discovery of peptides, proteins and nucleic acids. Given the importance of carbohydrate-protein interactions in diverse areas of biology including cell adhesion/recognition, immunoregulation and host-pathogen interactions, directed-evolution-based methods for discovery of potent glycoligands are greatly needed. We have previously reported a method for *in vitro* selection of glycopeptides that combines mRNA display, alkynyl amino acid incorporation, and CuAAC “click” glycosylation. Herein, we describe extensions of this method that incorporate chemical cyclization, removal of N-terminal glycosylation sites and next-generation sequencing; as an approach to HIV immunogen design, we have then used this method to develop mimics of the High Mannose Patch (HMP), which is the region on HIV envelope protein gp120 most commonly targeted by HIV broadly neutralizing antibodies (bnAbs). We prepared libraries of 10^12-14^ glycopeptides about 50 amino acids in length, containing variable numbers of high mannose (Man_9_GlcNAc_2_) glycans and cyclization at varied sites. We performed selections to obtain binders of HIV bnAbs PGT128, PGT122, and gl-PGT121, a germline precursor of PGT122, and prepared numerous glycopeptide hits by chemical synthesis. Selected glycopeptides in some cases bound very tightly to their target HIV bnAb, e.g., with a *K*_D_ as low as 0.5 nM for PGT128. These glycopeptides are of interest as immunogens and tools for HIV vaccine design.

## Main

The HIV pandemic has resulted in 40 million deaths and continues to infect about 1.5 million people per year.^1^ Although antiretroviral drugs can suppress viral load to undetectable levels, usually indefinitely, they do not eliminate the reservoir of latently infected cells, so must be taken for the rest of a patient’s life to prevent viral rebound, at significant cost.^2^ By comparison, an effective vaccine, once developed, will be a more efficient and cost-effective tool to combat the HIV epidemic.^3^ However, after 40 years of research, all HIV vaccines tested in humans have demonstrated little or no efficacy.^1^

The HIV vaccine field has taken a great deal of inspiration in recent decades from the discovery of HIV broadly neutralizing antibodies (bnAbs),^4-6^ which neutralize genetically diverse HIV strains, in exceptional cases up to 99%. A large fraction of infected individuals eventually produce bnAbs (20-50% depending on definitions of breadth), but only after months or years of chronic infection.^7-9^ Although this is typically insufficient to control infection, some monoclonal bnAbs have been shown to provide protection in animal models when administered prior to SHIV exposure.^10-14^ Thus, elicitation of bnAbs is one of the major goals of HIV vaccine design. These design efforts are supported by extensive structural studies of how bnAbs bind to the HIV envelope (Env) proteins gp120 and gp41, which have revealed which conserved structural features bnAbs bind to.^5, 6, 15^ The most common class of bnAbs, exemplified by PGT128,^16-18^ PGT122,^19, 20^ and 2G12,^21-23^ binds to a densely glycosylated region of gp120 known as the High Mannose Patch (HMP), which is centered around the N332 glycan and comprises several other glycans that are mainly Man_8-9_GlcNAc_2_ glycoforms.^24-26^ As an approach to vaccine design, there has been significant interest in the engineering of Env variants that better display the HMP or engage HMP bnAb germline precursors,^27-33^ and of synthetic glycopeptides that mimic the HMP and might elicit HMP-binding antibodies.^34-44^

We have sought to mimic HMP epitopes without using either the entire Env protein, which contains a plethora of “distracting” epitopes that easily elicit off-target antibodies;^45-47^ or short glycopeptides directly derived from the Env sequence, which may not sufficiently recapitulate the HMP structure without the conformational support of the remaining Env protein. Aiming to design glycostructures that better mimic the glycan presentation in the HMP, we have pioneered various techniques for the directed evolution of DNA,^48, 49^ RNA,^50^ F-RNA,^51^ or peptide-supported^52, 53^ glycan clusters. The latter method combines mRNA display,^54^ non-canonical amino acids,^55, 56^ and CuAAC^57, 58^ chemical attachment of glycans. We have used this technique for the *de novo* design of glycopeptide HIV antigens that present glycans in a manner tightly recognized by bnAb 2G12 with low nM to pM *K*_D_ values.^52^ We later showed that some of these glycopeptides could elicit gp120-binding antibodies without being based on the gp120 peptide sequence.^41, 42^ Compared to 2G12, which binds to only the glycans and has modest breadth (∼30%) and potency (EC_50_ ∼2.4 µg/mL), other HMP-binding antibodies such as PGT128 and PGT122 bind to a combination of glycan and polypeptide elements and exhibit much greater breadth (72 and 65%, respectively) and potency (EC_50_ 0.02 and 0.05 µg/mL).^59^ In the present work, we report multiple selections of glycopeptides that mimic PGT128- and mature or germline PGT122 epitopes using a modified glycopeptide mRNA display method that adds library N-terminal processing, peptide cyclization, and high-throughput sequencing to our previously described workflow.

## Results and discussion

### Selection of glycopeptide binders for PGT128

PGT128 is a broad and very potent bnAb, neutralizing 72% of a diverse HIV pseudovirus panel with a median IC_50_ of 0.02 µg/mL.^16^ It exhibits significant binding to high-mannose glycans on glycan arrays,^16, 26, 59^ and structural studies^16-18^ show that it makes contacts to varying extents with several high mannose glycans including N332 but also some peptide residues including the ^323^IGDIR^327^ sequence, a highly conserved gp120 motif that binds the CCR5 co-receptor (Fig. 1).^60^ We thus sought to display multiple high mannose glycans and the IGDIR motif within peptide scaffolds of a size substantial enough to resemble miniproteins.^61^ Thus, we designed a 49-residue peptide library in which IGDIR appeared early, late, or in the middle of the random region (the Less Biased or “L” library). In a parallel library (the Heavily Biased or “H” library), we extended this constant region to IGDIRXAXC*M* (X = random, *M* = a methionine analog, homopropargylglycine (HPG) for CuAAC glycosylation). This provided a fixed cysteine and glycosylation site shortly after the IGDIR sequence, mimicking the presence of the ^331^CN(glycan)^332^ in gp120. Remaining amino acids were random, encoded by NNS codons (N = A/T/G/C, S = G/C) to achieve frequencies of additional cysteine or glycosylation site codons of 1/32. To provide additional opportunities for peptides to adopt stable folds, we opted to cyclize the libraries by crosslinking cysteines with *m-*dibromoxylene.^62, 63^ We wished to avoid potentially exposed or conformationally floppy presentations of a glycan at the N-terminus of peptides resulting from glycan attachment at the N-terminal HPG encoded by the *Start*/*Met* codon; thus, we designed the library with a constant alanine at the second position, enabling enzymatic cleavage of the N-terminal HPG with peptide deformylase (PDF) and methionine aminopeptidase (MAP), which is capable of cleaving N-terminal methionine analogs (Supplementary Fig. 2).^64, 65^

**Fig. 1:**
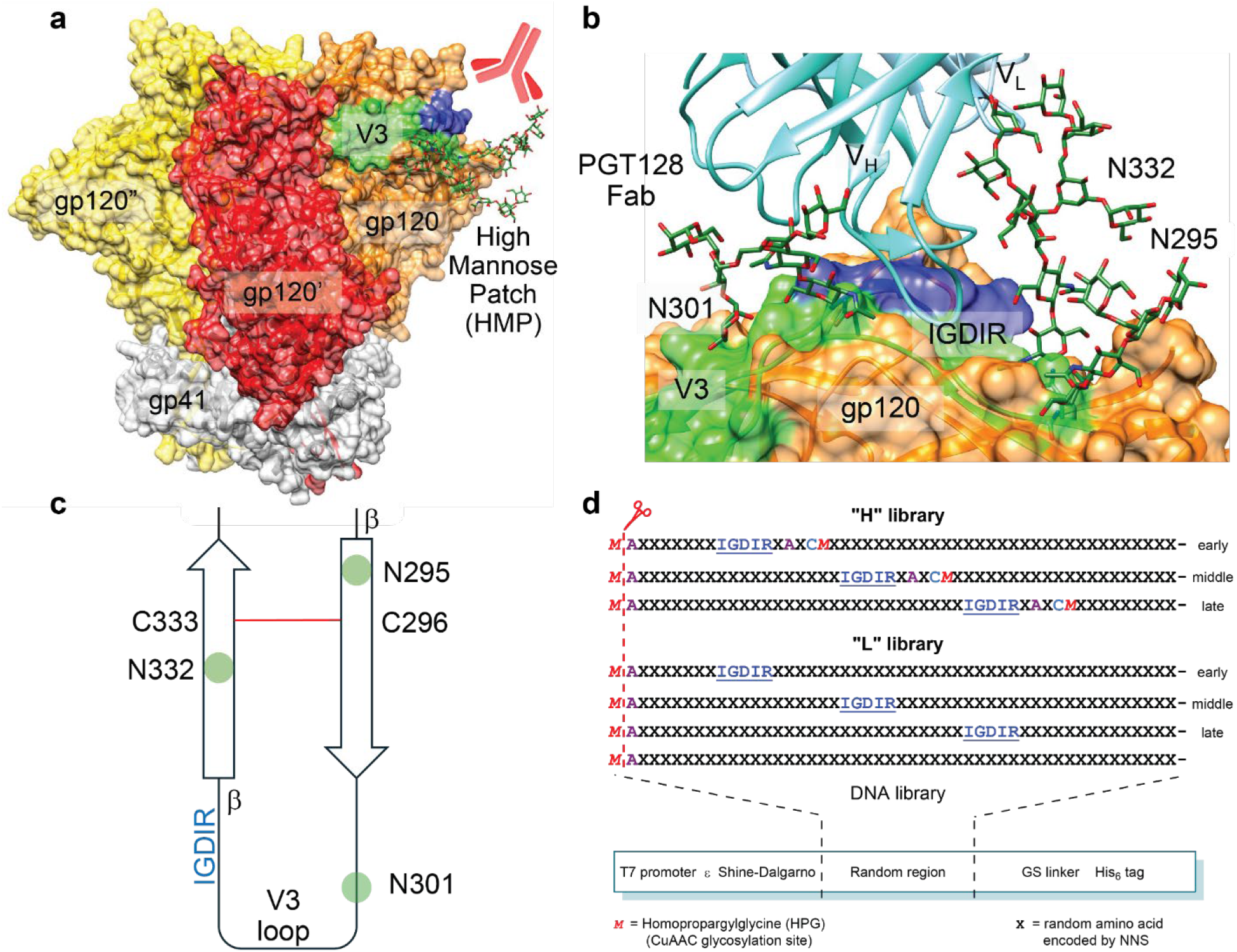
Epitope design for high mannose patch directed HIV bnAbs. a) Crystal structure of BG505 SOSIP.664, a native-like trimer (PDB ID: 5ACO). Rightmost protomer V3 loop is shown in green, HMP glycans at N332, N301, and N295 are displayed in forest green and IGDIR region in blue. b) PGT128 Fab makes contact with N332 and N301 glycans and IGDIR region of the V3 loop. c) Schematic representation of V3 loop PGT128 epitope region. d) Library design for selection with PGT128. All sequences followed by GSGSLGHHHHHHR. N-terminal *M* is cleaved co-translationally by PDF/MAP.

**Fig. 2:**
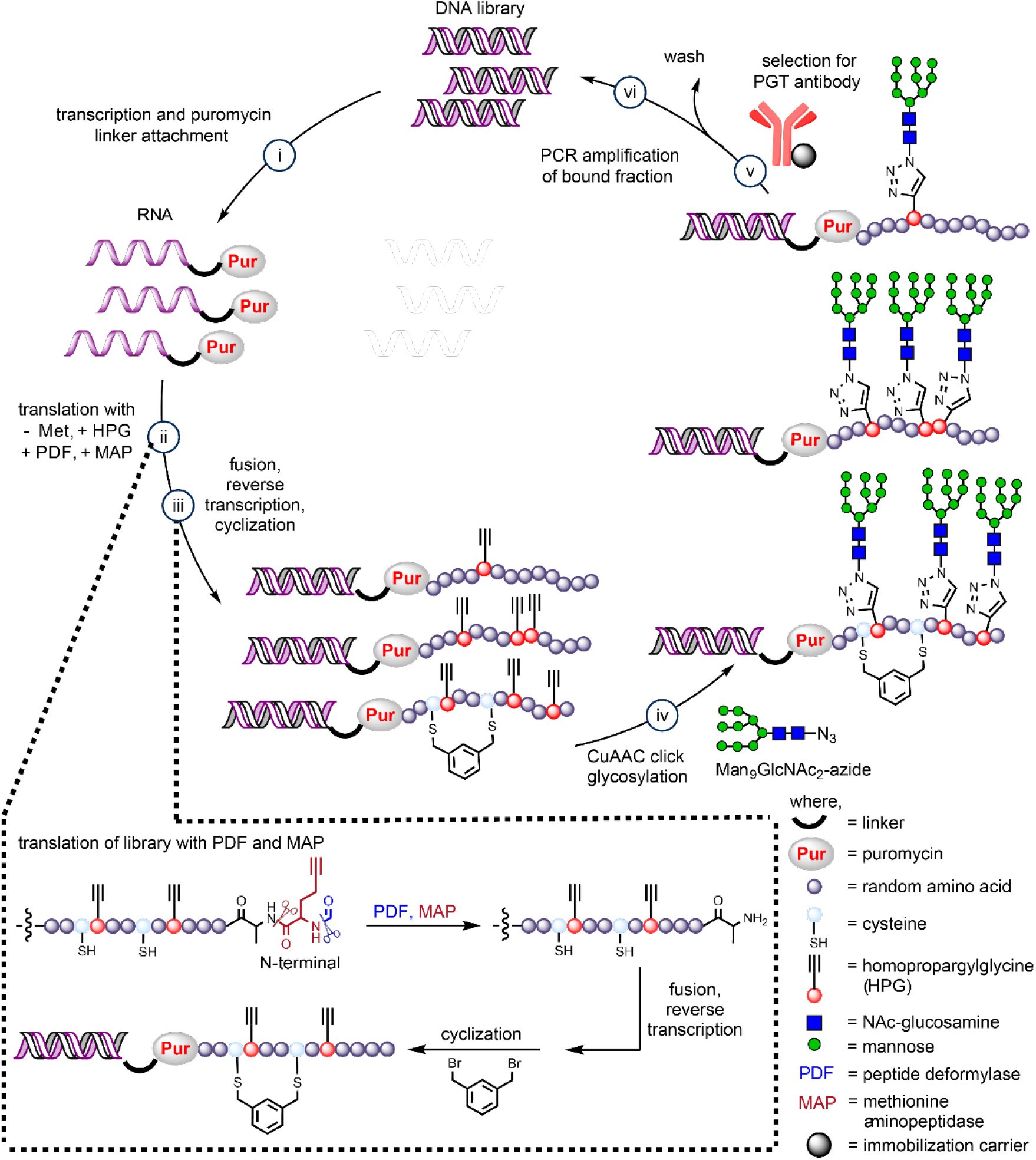
Scheme for *in vitro* mRNA display glycopeptide directed evolution.

Overall, the two libraries thus had theoretical multivalency distributions with 2-3 glycans in 59% of H library and 35% of L library peptides (see Supplementary Fig. 4).

DNA encoding the peptide libraries described above was prepared by assembly of several synthetic fragments and included T7 promoter, epsilon enhancer, Shine-Dalgarno sequence, a flexible linker, and a His_6_ tag for purification (Figs. 1d and 2). Following transcription, libraries were crosslinked to a puromycin oligo and translated in the presence of PDF/MAP to generate library fusions. Fusions were then captured on oligo(dT) beads and cyclized on-resin with *m-*dibromoxylene. Following elution from oligo(dT) beads, fusions were reverse transcribed and further purified on Ni-NTA resin to remove any free RNA. Following CuAAC attachment of synthetic Man_9_GlcNAc_2_ glycan azide,^66^ 7.3 and 6.8 pmol of libraries H and L were obtained, corresponding to 4.4 × 10^12^ and 4.1 × 10^12^ sequences, respectively.

For selection, libraries were each incubated with various concentrations of PGT128 IgG and the bound complexes retrieved on Protein G or Protein A beads (Fig. 3a). Antibody concentrations were decreased from 200 nM to 10 nM over the first five rounds of selection. In the next five rounds, selection pressure was increased not only by further decreasing antibody concentration to 2.5 nM, but also by increasing selection temperature from room temperature to 37 °C. At every round except the first, a negative selection was performed to discard Protein G or Protein A bead binders. Additional negative selections were performed in rounds 6-8 to remove the portion of the library that bound to PGT128 prior to glycosylation. A final two rounds of selection were performed with 50 nM biotinylated PGT128 Fab instead of IgG.

**Fig. 3:**
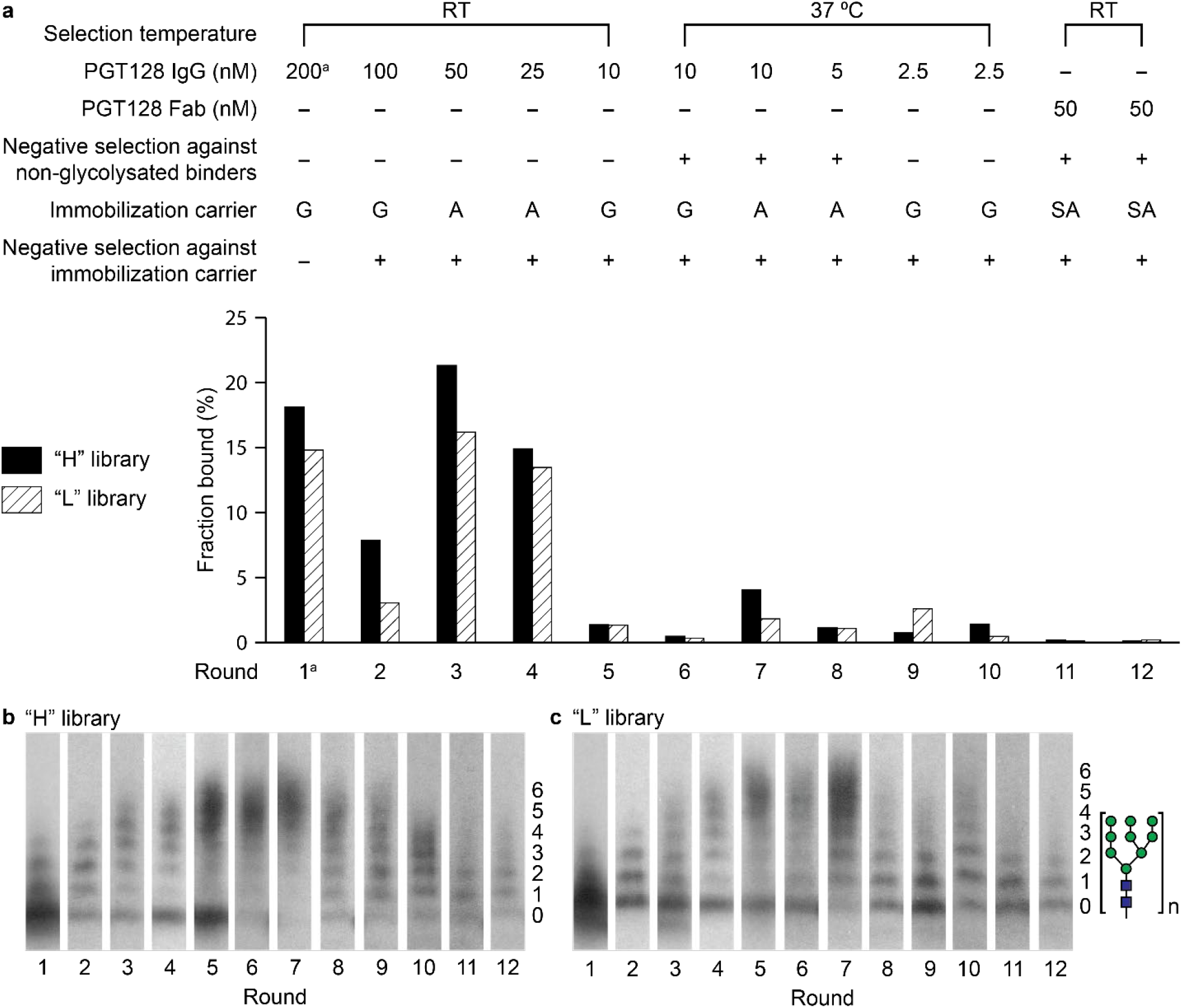
Summary of PGT128 selection progress. (a) PGT128 selection conditions and fractions bound for each round. The immobilization carrier used was Protein G (G), Protein A (A), or Streptavidin (SA); (b) Glycosylation from round to round for the “H” library and (c) “L” library, as visualized by SDS-PAGE and fluorography of library fusions after P1 nuclease digestion. (^a^ Round 1 selection was conducted with targets of parallel selections, see Supplementary Section II.B)

**Fig. 4:**
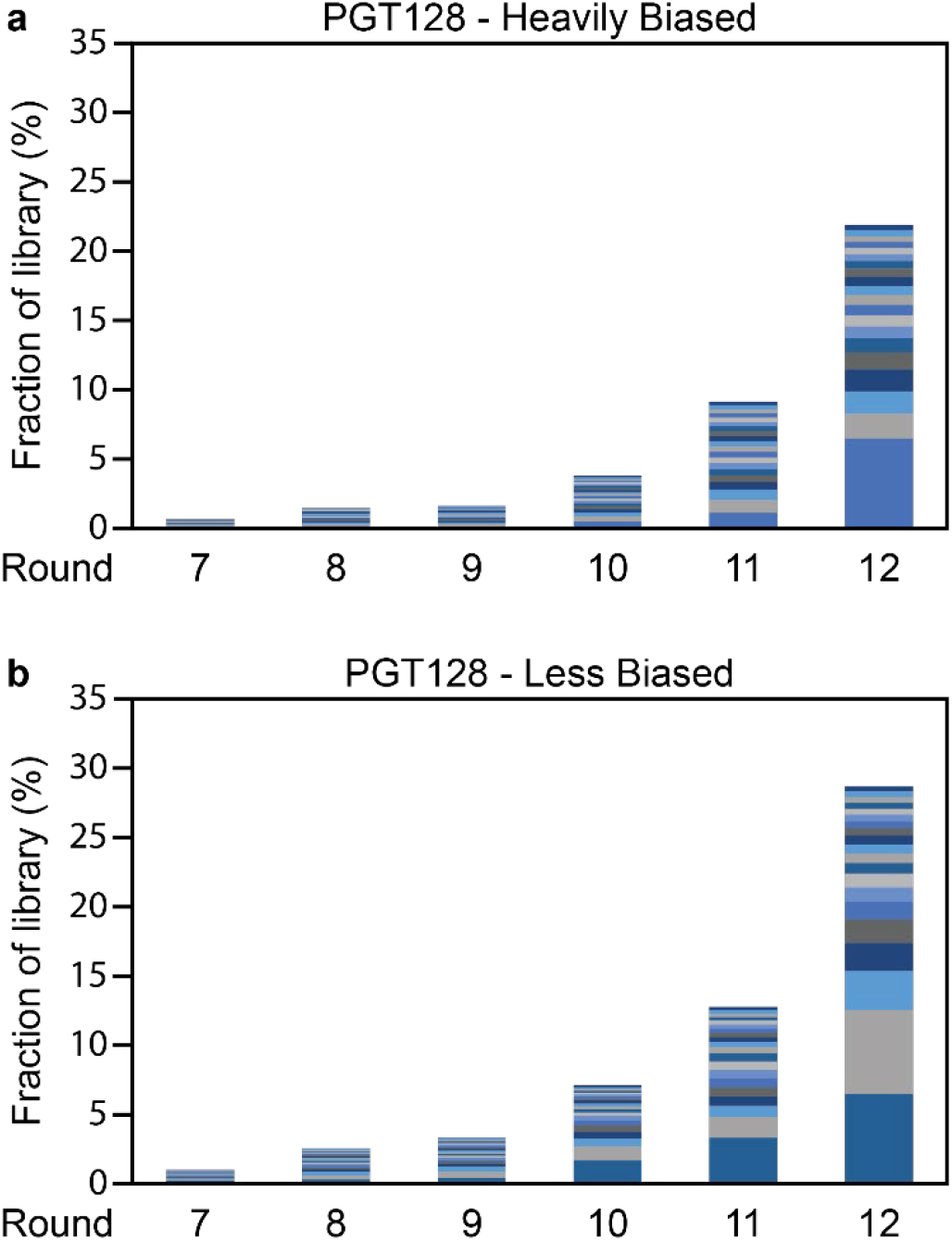
Enrichment of the top 20 most abundant sequence clusters in PGT128 selection. Each bar represents a sequence cluster (see Supplementary Tables 6 and 7), with the height relative to its portion of the library.

In the first five rounds of selection, conducted at room temperature, both H and L libraries became enriched (Fig. 3b,c) in highly multivalent glycopeptides, i.e., > 4 Man_9_GlcNAc_2_ sugars per peptide, which is undesirable because it exceeds the number of glycans present in the PGT128 epitope. Subsequent selection at 37 °C resulted in a loss of highly multivalent sequences and enrichment for glycopeptides of low multivalency. The multivalency decrease observed by gel was also confirmed in library sequencing data (Supplementary Fig. 7). These effects of selection temperature on multivalency parallel those we previously observed during selection of Man_9_-decorated libraries with 2G12.^49, 52^

Throughout the selections there was no increase in the fraction of library captured on the beads; instead, the fraction bound decreased to very low levels in more stringent rounds of selection. We were interested in whether tight binding sequences could nevertheless be identified in these libraries by inspection of sequencing data. DNA recovered from round 7-12 libraries was submitted for Amplicon-EZ next-generation sequencing (NGS) (GENEWIZ). To facilitate analysis of library diversity, sequence clusters were generated with CD-HIT (v4.8.1)^67^ with a threshold chosen such that peptide sequences with up to six mutations were grouped into one cluster. As shown in Fig. 4, the top 20 sequence clusters made up less than one percent of the round 7 libraries but had increased to over 20% of the round 12 libraries.

Although no individual sequences yet dominated the library at round 12, the most enriched clusters comprised up to 6% of each library.

To select sequences for synthesis and validation of binding to PGT128, clusters were ranked in terms of a scoring function that included points for high enrichment, two cysteines for cyclization, and the presence of 2-4 glycosylation sites (Supplementary Table 9). Based on this scoring function, clusters H1, H22, and L1 (6.47%, 0.46%, and 6.05% of their respective round 12 libraries) were chosen for synthesis. Peptides were prepared via microwave-heated Fmoc solid-phase peptide synthesis (SPPS) using DIC/Oxyma activation and 20% piperidine deprotections buffered with 0.1 M Oxyma (Fig. 5).^68^ For the purpose of binding analysis, sequences were prepared with a C-terminal biotinyllysine and a flexible GS linker. For cyclization, peptides were generally treated with TCEP followed by *m-*dibromoxylene in (NH_4_)HCO_3_ and acetonitrile. The resulting peptides (cyclic or acyclic, if applicable) were CuAAC glycosylated with Man_9_GlcNAc_2_ azide, and the final purity and identity of HPLC-purified glycopeptides were verified by LC/MS (Supplementary Section III).

**Fig. 5:**
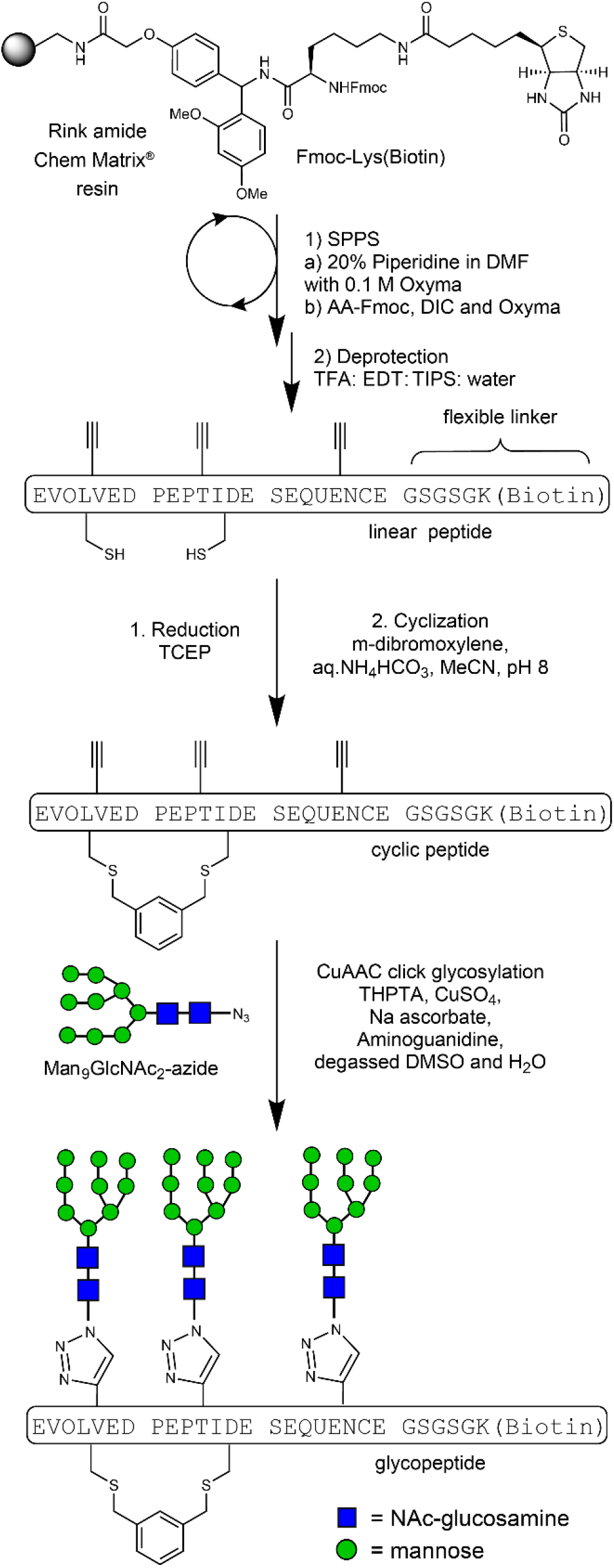
Generic scheme for the synthesis of selected glycopeptides.

Biolayer interferometry was used to assess the binding of the synthetic glycopeptides to PGT128 (Fig. 6). We were pleased to see that H1 and H22 glycopeptides bound to PGT128 with *K*_D_ values of 0.52 nM and 16.9 nM, respectively (peptides **1** and **2**). Given the very tight PGT128 recognition of glycopeptide H1, we synthesized various mutants to investigate the sequence dependence of this interaction. Omission of the glycan at the first of the three glycosylation sites (peptide **3**) resulted in ∼2-fold loss of recognition, whereas loss of the glycan at the second and third glycosylation sites resulted in a ∼40-fold decrease in binding (peptide **4**). No binding was observed at all to a deglycosylated mutant of H1 (Supplementary Fig. 33). H1 also contains the gp120 ^323^IGDIR^327^ sequence that was included in the H library design, and we sought to test whether this motif was important for PGT128 recognition as is the case for gp120. In gp120, mutation of ^323^IGDIR^327^ can significantly decrease binding and neutralization by HMP antibodies, with the G, D, and R positions being especially important.^59, 60^ In the context of H1, mutation of this motif to IAAIA resulted in a 26-fold decrease in PGT128 binding (peptide **5**). We also wondered whether an arbitrary sequence containing three Man_9_GlcNAc_2_ glycans and the same amino acids would bind similarly. Thus, we prepared a random scrambled version of the H1 sequence containing three glycans and cyclization (peptide **6**). It bound to PGT128 with a *K*_D_ value of 251 nM, ∼500-fold weaker than the parent sequence, demonstrating that glycan presentation created by H1 is clearly more antigenic than a random arrangement. Finally, to assess the importance of peptide cyclization on the presentation of epitope elements, we prepared an acyclic H1 variant (peptide **7**) in which cysteines were alkylated with iodoacetamide rather than the xylyl linker. 7 was recognized by PGT128 with a *K*_D_ value of 90 nM, ∼180-fold more weakly than the cyclized variant. However, another of the top library hits, L1 (peptide **8**), was acyclic but recognized by PGT128 with a *K*_D_ value of 2.37 nM, showing that tight binding can evolve without cyclization. Collectively, these data illustrate that directed evolution can furnish arrangements of high-mannose glycans with high antigenicity that is very dependent on peptide sequence/context.

**Fig. 6:**
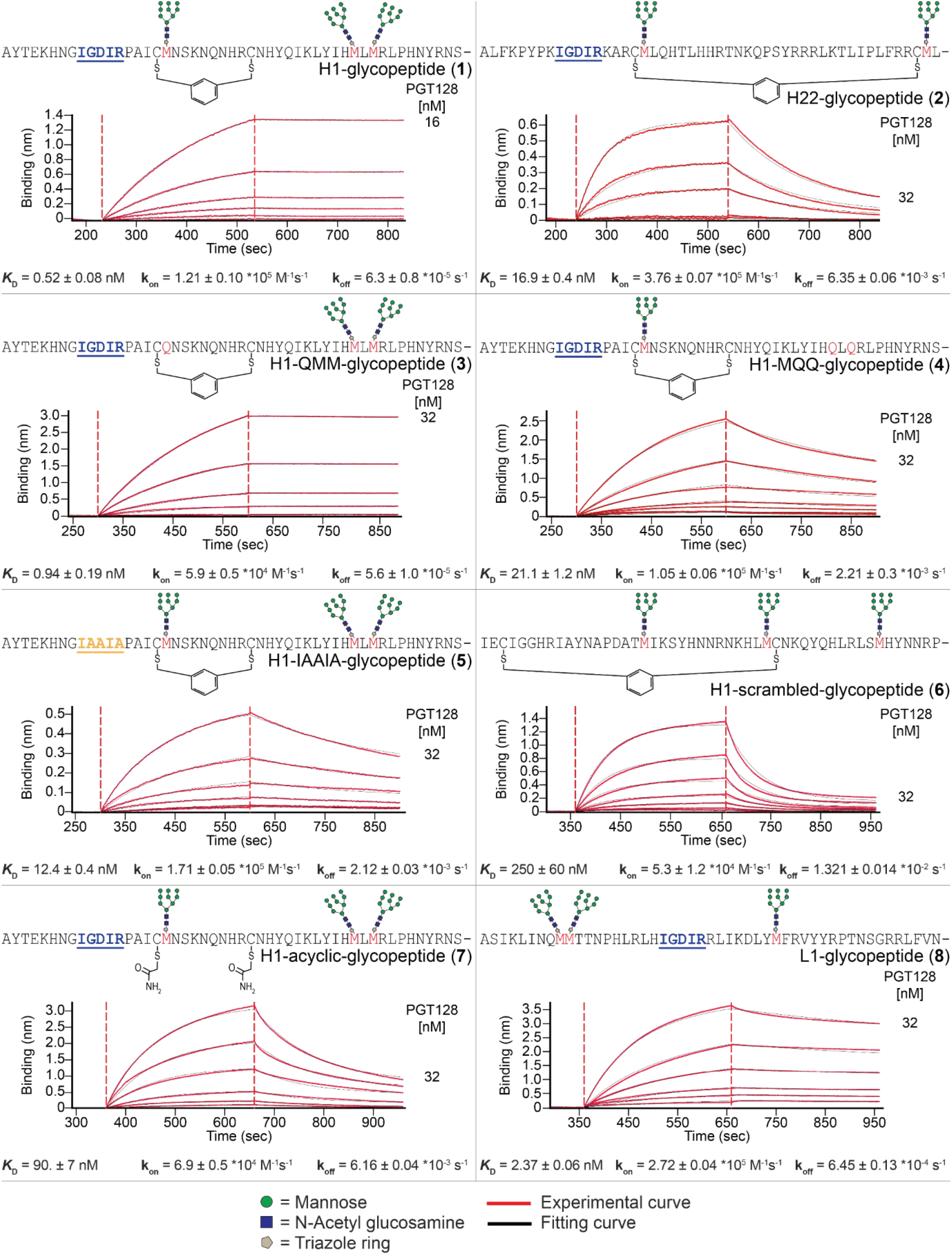
Biolayer interferometry (BLI) measurement of glycopeptides immobilized via biotin on a streptavidin biosensor binding to PGT128. All sensorgrams represent a series of two-fold dilutions of antibody with highest concentration tested labeled. All sequences followed by GSGSGK(Biotin). M denotes homopropargylglycine, modified by CuAAC as in Fig. 5.

**Fig. 7:**
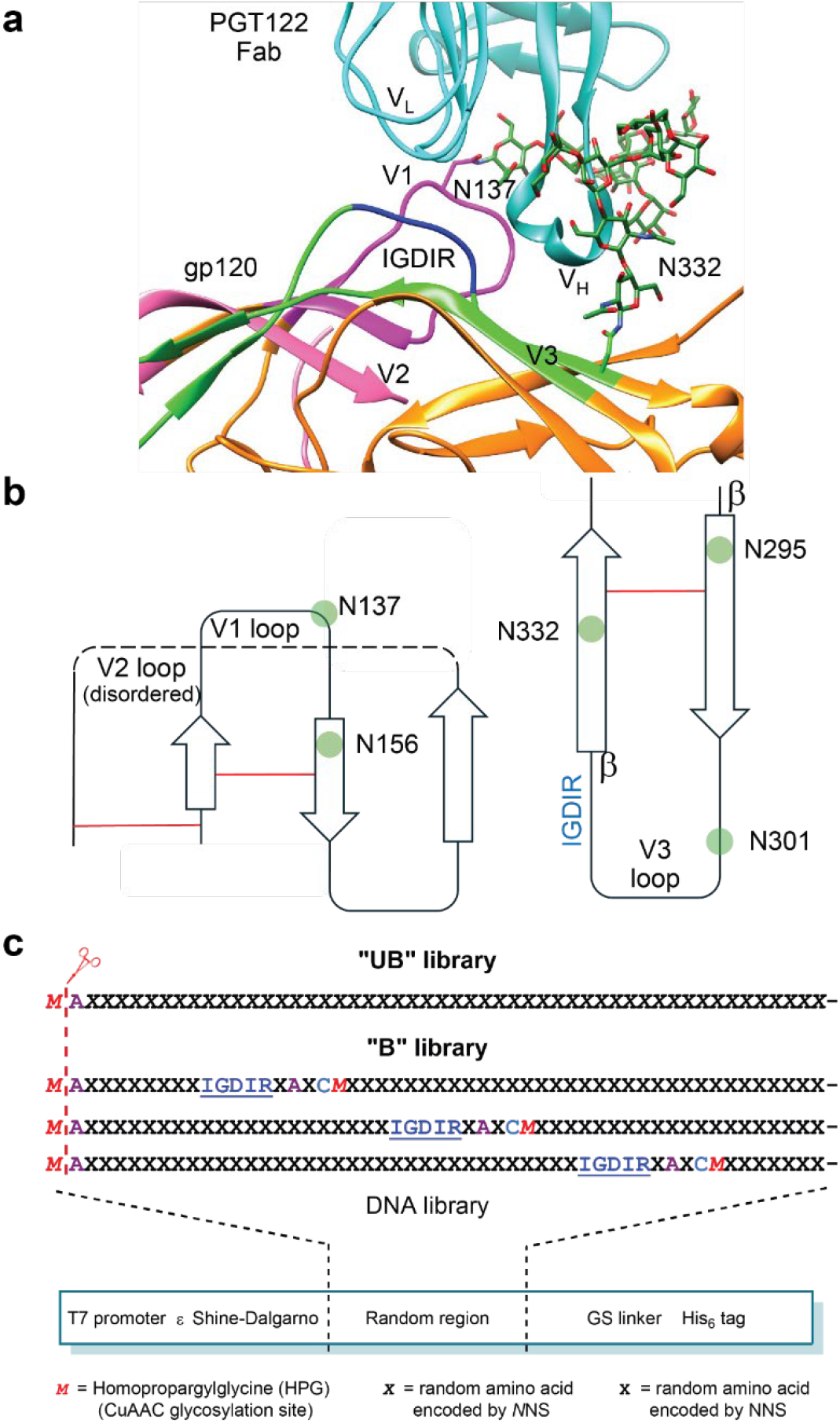
Epitope design for PGT122 HIV bnAbs. a) Crystal structure of BG505 SOSIP.664, a native-like trimer (PDB ID: 4NCO). V3 loop is shown in green, super site glycans at N332 and N137 are shown in forest green and IGDIR region in blue, b) Schematic representation of V1/V2/V3 loop PGT122 epitope region; PGT122 Fab makes contact with N332 glycan, the IGDIR region of V3 loop and the N137 glycan of V1/V2 loop c) library design for selection with PGT122 and gl-PGT121. All sequences followed by GSGSLGHHHHHHR. N-terminal *M* is cleaved co-translationally by PDF/MAP.

### Selection of glycopeptide binders for PGT122 and its germline precursor

PGT122 neutralizes 65% of HIV pseudoviruses in a representative panel with a median IC_50_ of 0.05 µg/mL and binds to the HMP in a manner similar to PGT128 despite originating from a different HIV positive donor.^25, 59^ The germline precursor of PGT122 (gl-PGT121) has also been the subject of studies to engineer Env protein immunogens that might initiate a PGT122-like antibody response by activating germline B cells derived from the same V/D/J genes as those that led to PGT122.^28-30^ PGT122 exhibits little binding to oligomannose glycans alone on arrays^26, 69^ although neutralization is dependent on the glycan at N332, which is exclusively high mannose.^70^ PGT122 also neutralizes viruses produced in the presence of kifunensine,^69^ which bear exclusively high-mannose glycans, and crystal structure data show that it can accommodate high mannose glycans (Fig. 7a,b).^20^ Similar to PGT128, PGT122 also recognizes the IGDIR motif.^59, 60^ Therefore, we designed libraries similar to those described for PGT128 selections to use in selections for binders of PGT122 and gl-PGT121. As shown in Fig. 7c, we created a Biased or “B” library of 51 amino acids in length containing the sequence IGDIRXAXC*M* at early, middle, or late positions with *M* again representing the HPG for glycan attachment. An Unbiased or “UB” library was also created with all random amino acids other than the N- and C-terminus using *N*NS codons (*N* = 40% A, 20/20/20% C/G/T). With a Met frequency of 1/32 among NNS codons and 1/20 among *N*NS codons, 85% of the B library and 68% of the UB library were calculated to contain 1-3 glycosylation sites (Supplementary Fig. 34). These libraries were prepared in a manner similar to that described for PGT128 selections (Supplementary Section IV.A), and the B and UB libraries were obtained with respective yields of 50 and 209 pmol (diversity ∼10^13^-10^14^ sequences).

Since the Round 1 libraries are the largest and hardest to prepare, this round was conducted with both planned targets PGT122 and gl-PGT121 combined in one pot. Selections began with 200 nM PGT122 and 200 nM gl-PGT121, and the library was then split for separate selections with the two antibodies in subsequent rounds. For PGT122 selection, stringency was increased as shown in Fig. 8a. As with the PGT128 selection, we monitored library glycosylation by SDS-PAGE of digested fusions (Fig. 8b,c); however, unlike the PGT128 selection, barely any increase of library glycosylation was observed in early selection rounds, and we observed a decrease starting in round 6, perhaps related to the lack of PGT122 affinity for high mannose.^26, 69^ An increase of non-glycosylated species was particularly obvious for the B library. We performed NGS starting in round 8 and found that most non-glycosylated sequences were frameshifted products (Supplementary Fig. 37). In round 12, we performed a selection to enrich peptides that contained at least one glycosylation site. This enrichment was accomplished by CuAAC attachment of biotin rather than glycans followed by capture of biotinylated peptides on streptavidin beads. This procedure successfully reduced species that lacked internal HPG sites (Fig. 8b,c, rounds 11 vs 13 and Supplementary Fig. 36a,b). It is noteworthy that, since translation starts with HPG in every peptide, this enrichment for internal HPGs was only possible due to our introduction of the N-terminal HPG cleavage step. Following this procedure, we then continued three more rounds of selection through round 15.

**Fig. 8:**
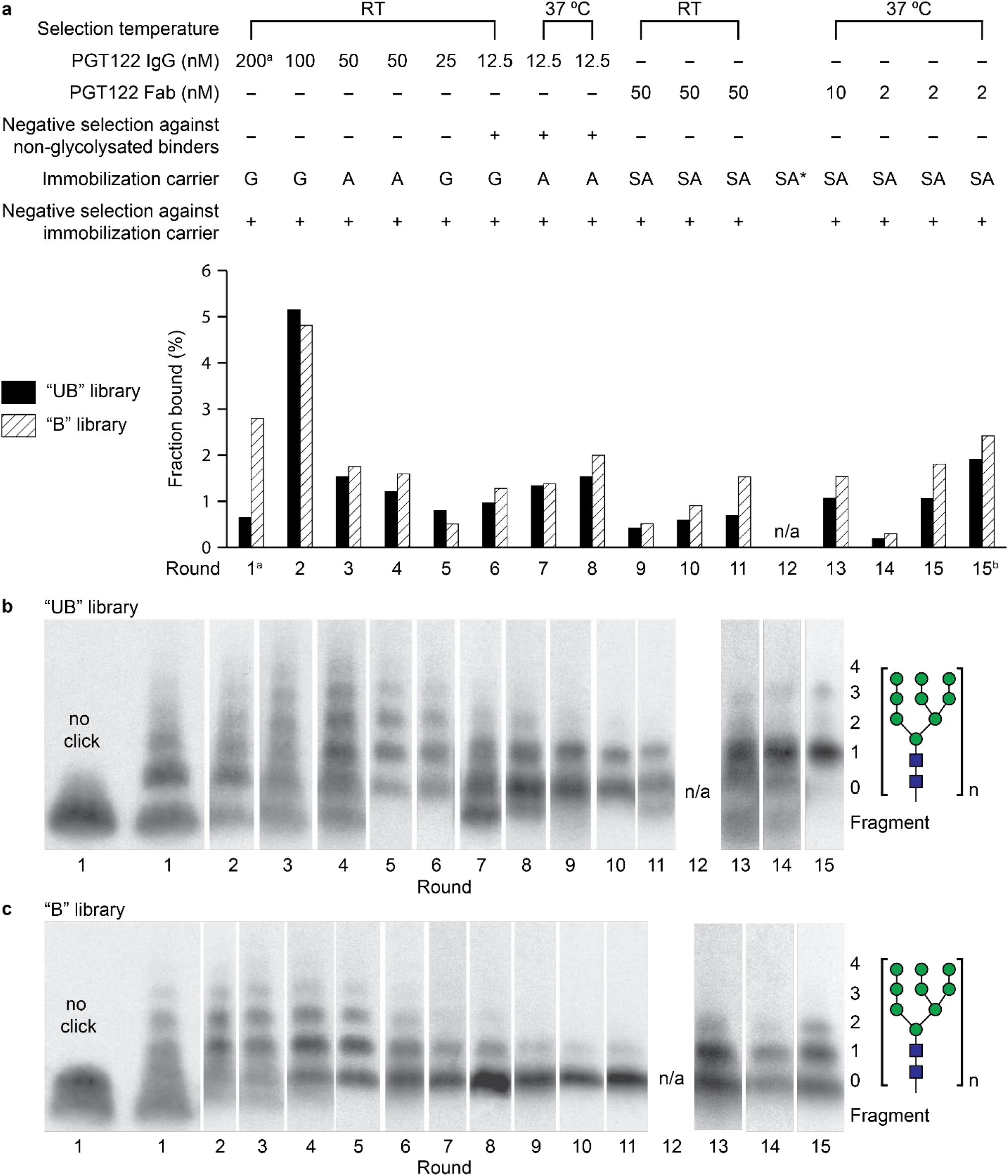
Summary of PGT122 selection progress. (a) PGT122 selection conditions and fractions bound for each round. The immobilization carrier used was Protein G (G), Protein A (A), or Streptavidin (SA); (b) Glycosylation from round to round for the “UB” library and (c) “B” library, as visualized by SDS-PAGE and fluorography of library fusions after P1 nuclease digestion. (^a^ Round 1 selection was conducted with both PGT122 and gl-PGT121 in one pot, ^b^ Round 15 selection with non-glycosylated library)

Compared with the PGT128 selection, sequences in the PGT122 selection converged to a greater extent in earlier rounds; therefore, sequences obtained through high throughput sequencing were grouped into larger clusters for analysis (threshold set to up to 12 sequence differences permitted per cluster). Supplementary Fig. 38 shows the most enriched clusters (by percentage), from rounds 8 to 15, with some clusters representing more than 10% of each library. We performed preliminary binding assays for those sequences and others using translated peptides with or without attached glycans. No sequences required glycans for binding, but some bound more strongly when glycosylated (Supplementary Table 22). Based on this screening and high final frequency in the round 15 library, we selected 2U7 (cluster frequency of 51%) for BLI binding assays with and without glycans (peptides 9 and 10). Glycosylated 2U7 bound to PGT122 with a *K*_D_ value of 111 nM (Figure 9). Though weaker than the H1-PGT128 interaction, this *K*_D_ value is similar to the reported affinities of PGT122 for HIV Env (100-174 nM).^69, 71^ Without glycosylation, 2U7 was recognized by PGT122 with a *K*_D_ of 215 nM. Lack of strong binding dependence on a high-mannose glycan is consistent with glycan array data indicating a preference of antibodies in the PGT122 family for other glycan types (e.g., sialylated biantennary complex)^26^ and with the observed decrease in library glycosylation throughout selection.

**Fig. 9:**
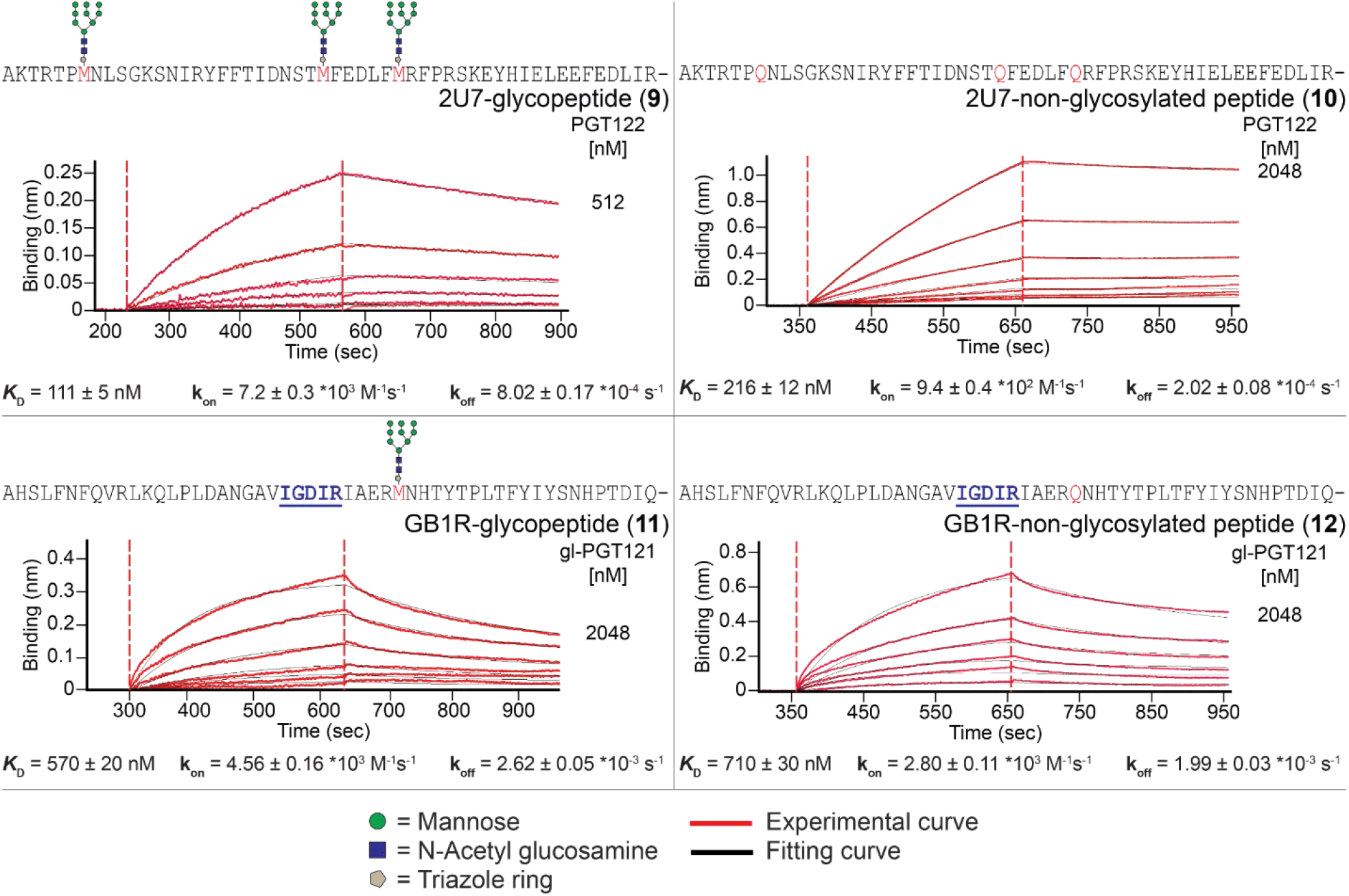
Biolayer interferometry (BLI) measurement of glycopeptides immobilized via biotin on a streptavidin biosensor binding to PGT122 and gl-PGT121. All sensorgrams represent a series of two-fold dilutions of antibodies with highest concentration tested labeled. All sequences followed by GSGSGK(Biotin). M denotes homopropargylglycine, modified by CuAAC as in Fig. 5.

Since the majority of the clusters in the PGT122 selection were already prominent among NGS data at rounds 8 and 9, we did not continue the selection for gl-PGT121 after round 9 and synthesized some sequences within the top 2 clusters of each library test for binding analysis by BLI. Among the sequences tested, only GB1R bound to gl-PGT121 in a detectable manner (574 nM *K*_D_, see Figure 9, peptide **11**), and with very little glycan dependence (708 nM *K*_D_ without glycosylation, peptide **12**). Although this affinity is modest, it is comparable to that of an HIV Env variant that was engineered to bind gl-PGT121, and WT HIV Env typically exhibits little or no affinity for this germline antibody.^28^ However, GB1R binding was specific for gl-PGT121, with no binding observed to mature PGT122 (Supplementary Fig. 45).

## Conclusions

Herein, we have demonstrated a modified glycopeptide mRNA display system including cyclization and N-terminal processing for the directed evolution of glycopeptide binders to three different antibodies in two HIV bnAb families. Selections demonstrated substantial enrichment by NGS data, and we used biolayer interferometry of chemically synthesized hits to validate highly enriched glycopeptide sequences for binding. In the case of PGT128, the best hit exhibited very strong (∼single digit nM) affinity that is to our knowledge, the strongest PGT128 binding to any glycopeptides.^35, 39, 40^ We also showed that tight binding for this sequence was contingent on the specific presentation of glycans achieved by full glycosylation, cyclization, and an unmodified peptide backbone with binding motif present. We also discovered binders of PGT122 and its germline precursor with antibody affinities comparable to those of the HIV Env,^69, 71^ albeit weaker than our PGT128 binders. This may result from the comparatively low affinity of PGT122-family antibodies for the high-mannose glycans used in our libraries.^26^ Overall, the HIV epitope mimic glycopeptides described here are attractive as candidate immunogens to elicit bnAbs to HIV, likely when employed in a multimeric format such as carrier protein conjugates or nanoparticles. As they lack HIV Env elements other than HMP epitope features, they may also be useful as cell sorting baits to isolate further HMP-directed bnAbs from the B cell repertoire of immunized or infected humans or rhesus macaques. With these potential applications in mind, the work described here may lead to useful improvements in HIV vaccine development.

## Acknowledgements

This work was supported by NIH grants R01-AI113737, R01-AI090745, and R21-AI140030. David Nemazee, Linghang Peng, and Jenny Tran are acknowledged for providing antibodies, and Dennis Burton and Devin Sok are acknowledged for supplying expression plasmids. Dan Polyak is acknowledged for preparation of Fmoc-homopropargylglycine, and Richard Redman is acknowledged for preliminary work on peptide synthesis.

## Author Contributions

† J.K.B. and S.H. are co-first authors. I.J.K., S.H., and J.K.B. conceived the project and designed the selection experiments. S.H. and J.K.B executed the selection experiments and preliminary binding studies with ribosomally-prepared hits. M.N. conducted the chemical synthesis and binding measurements of most glycopeptides. V. H. processed and clustered NGS data. K. N. conducted chemical synthesis and binding measurements of some glycopeptides. J.S.T. synthesized the glycan used for library production. All authors participated in data analysis and J.K.B., S.H., M.N., V.H., R.J.T and I.J.K. prepared the manuscript.

