## Supporting Information for "*In vitro* selection of cyclized, glycosylated peptide antigens that tightly bind HIV high mannose patch antibodies"

### Supplementary Information

### I. Preselection Experiments

#### A. Materials

All synthetic DNA oligos were purchased from Integrated DNA Technologies unless otherwise stated. Oligos with a length of 50 bases or more were purified by denaturing urea-PAGE before use. Stock solutions of individual dNTPs and NTPS were each obtained as a set from Thermo Scientific. Restriction endonucleases were purchased from New England Biolabs.

All reagents and buffers, generally Molecular Biology grade, were purchased from Fisher Scientific, Ambion, or Sigma. All water used was either purified by a Milli-Q Ultrapure water purification system or purchased RNA grade water (Fisherbrand, BP561-1). Homemade buffers were sterilized by passage through 0.22 µm filters (Millipore). $Man_9(GlcNAc)_2$-$N_3$, $Man_9$(cyclohexyl)-$N_3$ and Tris(3-hydroxypropyltriazolylmethyl)amine ligand (THPTA) used in the click reaction were chemically synthesized in-house.[1]

All PCR amplification was performed on an Eppendorf Mastercycler Gradient thermal cycler. Quantitative absorbance measurements were performed on a NanoDrop 2000c (Thermo Scientific) or NanoDrop One$^C$ (Thermo Scientific).

Precast SDS-PAGE gels (4-20% gradient) were purchased from Bio-Rad. All other SDS-PAGE gels were homemade using a Mini-PROTEAN® Tetra Handcast System (Bio-Rad). For $^{35}$S-cysteine-containing samples, gels were fixed, dried on chromatography paper, exposed to phosphorimaging screen, and visualized using a Storm or Typhoon phosphorimager. For $^{3}$H-histidine-containing samples, gels were fixed, treated with NAMP100 fluorographic reagent (GE Healthcare), dried on chromatography paper, and exposed to film.

Samples for liquid scintillation counting were prepared in 20 mL scintillation vials (Fisherbrand, 03-337-23) with 2-15 mL of Econo-Safe Biodegradable Counting Cocktail (Atlantic Nuclear, 11175). Scintillation counting was performed on a Beckman LS6000TA scintillation counter.

A Voyager DE Pro instrument was used for all MALDI-TOF-MS. Except where otherwise noted, the MALDI matrix used was α-cyano-4-hydroxycinnamic acid (CHCA; Sigma). CHCA was prepared as a 10 mg/mL solution in 0.1% (v/v) TFA/MeCN and stored at -20 °C in between uses for up to a month.

#### B. Cyclization of Free Peptide

DNA for the preselection sequence 12A from our previous 2G12 selection[2] was PCR amplified from its encoding plasmid as a template. The plasmid was diluted 50-fold into a 100 µL PCR reaction with final 0.2 mM of each dNTP, 0.5 µM of each forward primer (Library-FP; see **Table S3** for preselection primer sequences) and reverse primer (V/L-RP), 0.2 ng/µL plasmid template, and 0.02 U/µL Phusion™ Hot Start II Polymerase (Thermo Scientific) in 1X Phusion HF Buffer. The following PCR protocol was used: 98 °C for 30 seconds, followed by 25 cycles of 98 °C for 5 seconds, 62 °C for 10 seconds, and 72 °C for 10 seconds, followed by 72 °C for 5 minutes. The product DNA was purified by extraction with phenol: chloroform: isoamyl alcohol (25:24:1, v/v/v; VWR) and then chloroform followed by ethanol precipitation. The resulting DNA was transcribed using T7 polymerase as described in **Section II.A.2**, and the RNA was purified by 5% denaturing urea-PAGE with UV shadowing and passive elution. Passive elution was performed by soaking gel pieces in 0.4 M KOAc (pH 5.5) overnight at room temperature with tumbling, and RNA were collected by isopropanol precipitation followed by 70% (v/v) ethanol rinse. 12A RNA was translated to peptide in the presence of $^{35}$S-cysteine and HPG and in the absence of methionine as previously described.[3] Crude peptides were split and purified with Ni-NTA agarose with or without cyclization while captured on resin.

The protocol for Ni-NTA agarose purification of peptides without cyclization is as follows: To the crude translation (25 µL) was added four volumes (100 µL) of bind buffer (50 mM Tris-HCl, pH 7.8, 300 mM NaCl, 5 mM BME) and one volume (25 µL) of Ni-NTA Agarose (Qiagen). The mixture was tumbled at room temperature for 1 hour. Next, the suspension was transferred to a 0.22 µm Ultrafree®-MC Centrifugal Filter (MilliporeSigma UFC30GV00), and the tube was rinsed with four volumes (100 µL) of bind buffer to transfer remaining resin to the same filter. The filter was spun down at 14,000 ×g for 30 seconds to remove the flow-through. To wash the resin, eight volumes (200 µL) of bind buffer was added, and the filter was spun down as before. This was repeated for a total of three washes with bind buffer. Next, the resin was washed in the same way, but with wash buffer (50 mM Tris-HCl, pH 7.8, 5 mM BME) for a total of two times. To elute the peptide, one volume (25 µL) of 0.2% (v/v) trifluoroacetic acid (TFA) was added to the resin and allowed to sit for 2 minutes before spinning down into a new 1.5 mL centrifuge tube. The elution step was repeated twice more with collection each time into a new 1.5 mL centrifuge tube.

The protocol used for cyclization while captured on resin is as follows: To 25 µL of crude translated peptide was added 100 µL of bind buffer without BME (50 mM Tris-HCl, pH 7.8, 300 mM NaCl) and 25 µL of Ni-NTA Agarose (Qiagen). The mixture was tumbled at room temperature for 1 hour. Next, the mixture was spun down at 21,000 ×g and the supernatant was carefully removed from the loosely-pelleted agarose. The resin was washed six times by resuspending with 200 µL of bind buffer with TCEP (50 mM Tris-HCl, pH 7.8, 300 mM NaCl, 0.2 mM TCEP), spinning down, and removing the supernatant. The resin was then washed twice with 200 µL of wash buffer with TCEP (50 mM Tris-HCl, pH 8.0, 0.2 mM TCEP). To the resin was then added 100 µL of free peptide cyclization buffer (40 mM Tris-HCl, pH 8.0, 0.2 mM TCEP, 5 mM m-dibromoxylene, 20% (v/v) MeCN), and the mixture was covered in foil and tumbled at room temperature for 30 minutes. The reaction was spun down, and the supernatant was removed. Next, the resin was washed twice with 200 µL of bind buffer (50 mM Tris-HCl, pH 7.8, 300 mM NaCl, 5 mM BME). The resin was resuspended in 100 µL of wash buffer (50 mM Tris-HCl, pH 7.8, 5 mM BME) and transferred to a 0.22 µm Ultrafree-MC Centrifugal Filter (MilliporeSigma). The tube was rinsed with 100 µL of wash buffer, and the rinse was added to the filter. The filter was then spun down at 10,000 ×g for 30 seconds. The resin in the filter was washed with 200 µL of wash buffer. To elute the peptide, 25 µL 0.2% (v/v) TFA was added to the resin and allowed to sit at room temperature for 2 minutes before spinning down and collecting in a new tube. The elution step was carried out twice more, with each collected separately. The linear and cyclized peptides were analyzed by MALDI-TOF-MS (**Figure S1**).

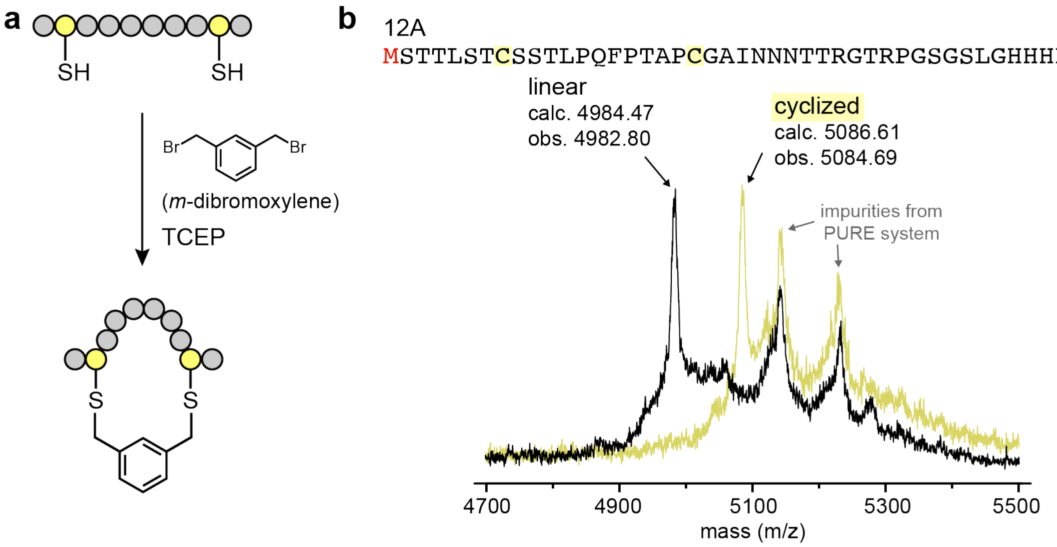

**Figure S1**. Peptide cyclization with m-dibromoxylene. (a) Generic depiction of bis-alkylation using m-dibromoxylene; (b) The amino acid sequence of a test peptide (12A) and MALDI-TOF-MS spectra of cyclized and non-cyclized 12A peptides. M in red denotes HPG.

#### C. MAP and PDF cleavage tests on model peptides

Peptide deformylase (PDF) and methionine aminopeptidase (MAP) were prepared as reported by the Suga group.[4]

_1. Preparation of RNA for test MAP/PDF cleavage_

In order to analyze N-terminal formyl HPG digestion by PDF and MAP using MALDI-TOF-MS, easily detectable short peptides were designed (**Table S1**). The dsDNA templates for T7 transcription of those sequences were prepared by bidirectional polymerase extension of annealed oligos. The oligos used are listed in **Table S1**, as well as the peptide sequences encoded by combinations thereof.

**Table S1**. Oligos for polymerase extension and encoded peptide sequences.

| Name | Sequence (5′ to 3′) |
|---|---|
| MH6C-1 | TAATACGACTCACTATAGGGTTAACTTTAGTAAGGAGGACAGCTAAATGCACCAC CATCACCATCACTGCAAA |
| MAH6C-1 | TAATACGACTCACTATAGGGTTAACTTTAGTAAGGAGGACAGCTAAATGGCGCAC CACCATCACCATCACTGCAAA |
| MH6C-2AAdeltaM | CTAGCTACCTATAGCCGGTGGCATTTATGGGTCAGACGCGCCGGGTCGATGTAGG CTTTGCAGTGATGGTGATGGTG |
| MH6C-AAΔM[a] | MHHHHHHCKAYIDPARLTHKCHRL |
| MAH6C-AAΔM[b] | MAHHHHHHCKAYIDPARLTHKCHRL |

Peptide sequences are encoded by the following oligo combinations: a MH6C-1 and MH6C-2AAdeltaM; b MAH6C-1 and MH6C-2AAdeltaM. The annealing sites are underlined. M in red denotes HPG.

Polymerase extension reactions (100 μL) were assembled by combining 10 μL of 10X Apex Buffer II (Genesee Scientific), 3 μL of 50 mM $MgCl_2$, 2 μL of 10 mM each dNTP mix, 1 μL of 100 μM oligo-1 (MH6C-1 or MAH6C-1), 1 μL of 100 μM oligo-2 (MH6C-2AAdeltaM), and 0.5 μL of Apex Hot Start Taq (Genesee Scientific). Using a thermal cycler, the reactions were heated at 98 °C for 10 minutes, 55 °C for 15 minutes, and 72 °C for 45 minutes. The crude reaction mixtures were diluted with water to 300 μL, extracted with phenol/chloroform, and precipitated with ethanol followed by a 70% (v/v) ethanol wash. The pellets were dissolved in 25 μL of water. The DNAs were analyzed on a 2% agarose gel, compared to controls with known concentrations to approximate the yield for assembling T7 transcription. DNA was transcribed, and the crude transcripts were purified by 8% denaturing urea-PAGE for use in translation.

*2. PDF/MAP Cleavage Tests*

RNA Sequences prepared above were translated in the presence of $^{35}$S-cysteine with one of the following conditions: translations were conducted either as previously described[3] with 1) neither PDF nor MAP, 2) with 6 μM PDF only, or 3) with 6 μM PDF and 15 μM MAP with 100 μM $CoCl_2$ cofactor. Peptides were purified by Ni-NTA agarose purification without cyclization and analyzed by MALDI-TOF-MS. As expected, we observed mass reductions for MAH6C-AAΔM corresponding to cleavage of the N-terminal formyl group with PDF only, and loss of HPG as well when MAP was also included (**Figure S2**). For MH6C-AAΔM, the histidine immediately following HPG strongly inhibited MAP cleavage, and only formyl cleavage was observed, consistent with a previous report[6] (data not shown).

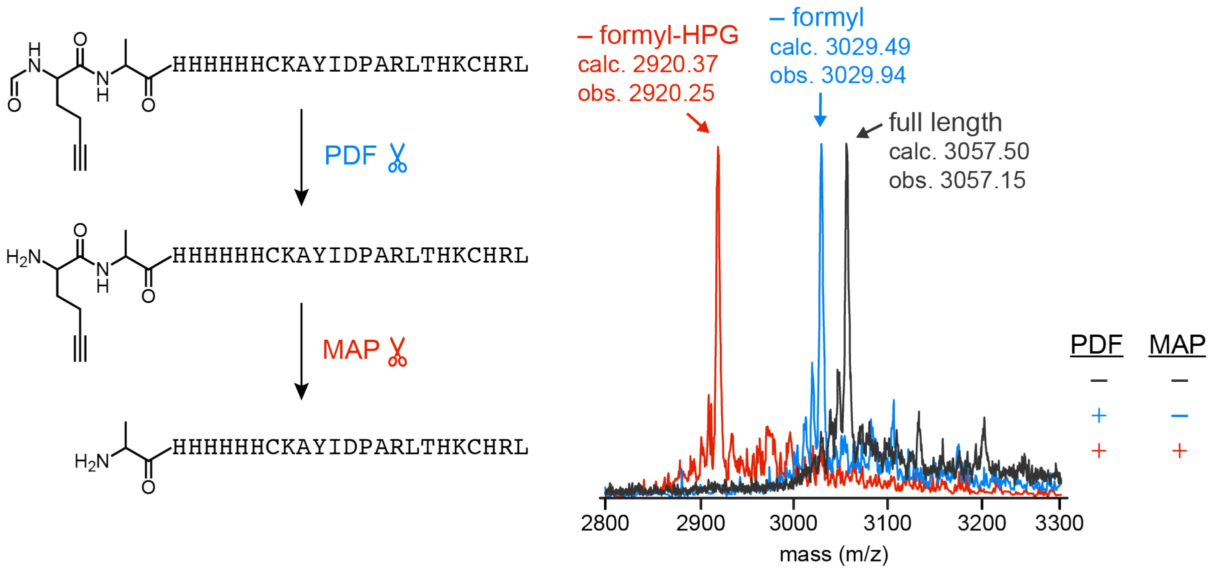

**Figure S2**. MALDI-TOF-MS analysis of N-terminal formyl HPG processing with PDF/MAP.

To observe the reduction in CuAAC glycosylation sites that results from cleavage of the N-terminal HPG, two 2G12-binding glycopeptides, 10V1 and 10F2, which contain 3 and 5 HPG residues respectively, but differ in their second amino acid so that only 10V1 should be cleaved by MAP.[2] These were prepared from their encoding plasmids as was done for 12A described in **Section I.B**. Both sequences were translated in the presence of $^{35}$S-cysteine either with both PDF and MAP or with neither, purified by Ni-NTA agarose purification without cyclization, and click glycosylated with $Man_9$(cyclohexyl)-$N_3$[2] as described in **Section II.A.8**. Samples were then run on a 4-20% SDS-PAGE (Bio-Rad) gel to check for number of glycosylation events. 10V1 has an alanine after the N-terminal HPG; as expected, it thus underwent N-terminal HPG cleavage, with a gel exhibiting 3 glycosylation events after translation without PDF/MAP but only 2 glycosylation events with PDF/MAP (**Figure S3A, left**). In contrast, 10F2, which has a histidine following the N-terminal HPG, exhibited negligible reduction in glycosylation in the same experiment (**Figure S3A, right**). To verify that use of PDF/MAP in translation is compatible with generation of glycopeptide library, the 10V1 glycosylation experiment was repeated with its RNA-peptide fusion: following the procedure in **Section II.A.2-9**, 10V1 RNA was photo-crosslinked with a puromycin-containing oligo, translated with or without MAP/PDF, reverse transcribed, cyclized, click glycosylated with $Man_9$(cyclohexyl)-$N_3$, and treated with Nuclease P1. As shown in **Figure S3B**, fusion integrity is not affected by the presence/absence of MAP/PDF, and the expected reduction in glycosylation was observed for MAP/PDF-treated sample (**Figure S3C**).

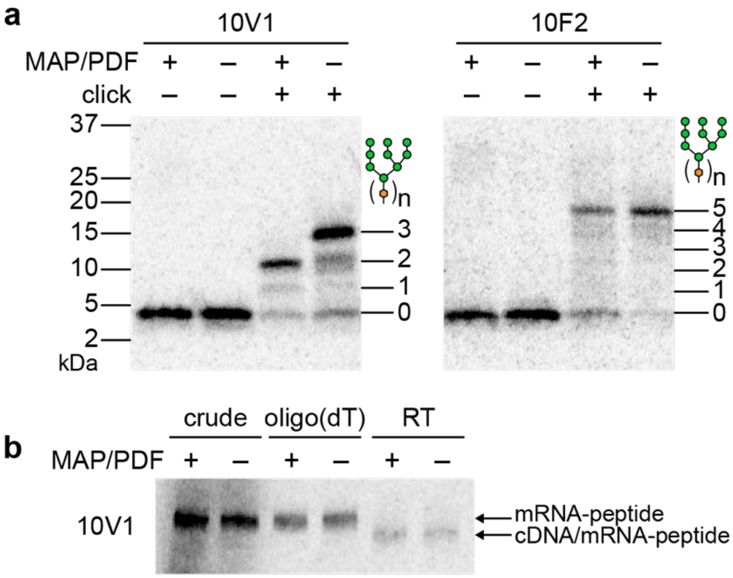

**Figure S3**. PDF/MAP processing analysis by SDS-PAGE. (a) Glycosylation of peptides 10V1 and 10F2 prepared by translation as free peptides with or without PDF/MAP treatment; (b) 10V1 fusions prepared with or without PDF/MAP in translation: mRNA-peptides, cDNA/mRNA-peptides, and (c) glycosylated peptides prepared as cDNA/mRNA fusions followed by nuclease digestion. All samples were radiolabeled with $^{35}$S-cysteine. Peptides (a) were run on a 4-20% SDS-PAGE gel (Bio-Rad). Fusions (b) were run on a homemade 7.5% SDS-PAGE gel. Glycosylated fusions (c) were digested with Nuclease-P1 for analysis on a 4-20% SDS-PAGE (Bio-Rad). Gels were visualized by phosphorimaging. 10V1 sequence: **M**ATKTNCKREKT**M**DNHVTI**M**RSIPWYTYRWLPNGSGSGLGHHHH-HHRL ; 10F2 sequence: **M**HPYNTSRTSA**MM**AALK**M**QVTD**M**YALALFHRILGSGSGLGHHHHHHRL (**M** = HPG).

### II. PGT128 Selection

#### A. Library Generation for Round 1 PGT128 Selection

*1. Preparation of Library DNA*

To create each library pool with fixed elements in various locations, separate DNA libraries were purchased and combined. The desired DNA lengths were 249 bases, longer than the limitations quoted by our commercial suppliers. To generate the full-length DNAs, we purchased the libraries as 197-mer oligos containing the entire randomized region and part of the 5' and 3' constant regions for extension by PCR to the desired 249 base length. The antisense strands of the DNA libraries to be extended and used in this study were purchased from Integrated DNA Technologies (Ultramer® DNA Oligos, 4 nmol; summarized in **Table S2**; theoretical multivalency distribution given in **Figure S4**). The lyophilized powders were dissolved in 100 µL of water. To 40 µL of library in water was added 60 µL of 8 M urea. The libraries were heated at 70 °C for 5 minutes before loading onto a pre-run 5% denaturing urea-PAGE gel. The library bands were visualized with a 254 nm handheld UV lamp and cut out with fresh razors for electroelution with Elutrap Electroelution System (Whatman) with 0.5X TBE running buffer. The eluents were precipitated with isopropanol followed by a 70% (v/v) ethanol wash. The library pellets were dissolved in 26 µL of water and quantified using NanoDrop.

The purified DNA templates were extended by PCR amplification. In a representative example, PCR (1.2 mL) was performed for each library with 15 nM of purified template (18 pmol), 0.2 mM of each dNTP, 1 µM of the forward extending primer (JB-HL-fwd-X), 1 µM of the reverse extending primer (JB-H-rev-X for the Heavily Biased library; JB-L-rev-X for the Less Biased, see **Table S3** for PGT128 selection primer sequences), and 0.025 U/µL Taq DNA Polymerase (New England Biolabs) in 1X Standard Taq Buffer. The following PCR protocol was used: 5 cycles of 95 °C for 30 seconds, 57 °C for 15 seconds, and 68 °C for 15 seconds, followed by a final extension at 68 °C for 5 minutes. Minimal cycling was used in order to keep the copy number of each sequence low. The crude PCR products for the Heavily Biased pool were combined into a 15 mL conical centrifuge tube (likewise for the Less Biased pool). The DNA solutions were extracted with phenol/chloroform, followed by 3 extractions with n-butanol to concentrate the DNA solution. The resulting DNA solution (~2-3 mL) was aliquoted into 2 mL microcentrifuge tubes and precipitated with isopropanol. The pellets were rinsed with 70% (v/v) ethanol and dissolved in 150 µL of water. The extended DNA library pools (summarized in **Table S4**) were analyzed on a 2% agarose gel. The yields were estimated to be 188-235 pmol based on comparisons to control samples with known concentrations.

**Table S2**. Individual libraries for selection with PGT antibodies. The Heavily Biased library comprises three individual libraries with the conserved IGDIRXAXCM motif early (E), middle (M), or late (L) in the sequence. The Less Biased library comprises four individual libraries with the conserved IGDIR motif early (E), middle (M), or late (L) in the sequence, or not at all (0). Codon degeneracy was exploited in the C-terminal constant region to design orthogonal primers for the two library pools. The peptide sequence pools derived from each individual library are shown, lacking the MAP-cleaved N-terminal HPG. The antisense DNA sequences of each library, as ordered from IDT, are also shown.

| Library | Name | Sequence 5′ to 3′ [a] |
| --- | --- | --- |
| Heavily Biased | H-E-2 | AXXXXXXXIGDIRXAXCMXXXXXXXXXXXXXXXXXXXXXXXXXXXXXXX |
| | | GTGGTGATGACCCAGAGAACCGGAGCCSNNSNNSNNSNNSNNSNNSNNSNNSNN<br>SNNSNNSNNSNNSNNSNNSNNSNNSNNSNNSNNSNNSNNSNNSNNSNNSNNSNN<br>SNNSNNSNNSNNCATGCASNNCGCSNNACGAATATCGCCAATSNNSNNSNNSNN<br>SNNSNNSNNCGCCATTTAGCTGTCCTCCTTACTAA |
| | H-M-2 | AXXXXXXXXXXXXXXXXXXIGDIRXAXCMXXXXXXXXXXXXXXXXXXXX |
| | | GTGGTGATGACCCAGAGAACCGGAGCCSNNSNNSNNSNNSNNSNNSNNSNNSNN<br>SNNSNNSNNSNNSNNSNNSNNSNNSNNSNNSNNCATGCASNNCGCSNNACGAAT<br>ATCGCCAATSNNSNNSNNSNNSNNSNNSNNSNNSNNSNNSNNSNNSNNSNNSNN<br>SNNSNNSNNCGCCATTTAGCTGTCCTCCTTACTAA |
| | H-L-2 | AXXXXXXXXXXXXXXXXXXXXXXXXXXXXXIGDIRXAXCMXXXXXXXXX |
| | | GTGGTGATGACCCAGAGAACCGGAGCCSNNSNNSNNSNNSNNSNNSNNSNNSNN<br>CATGCASNNCGCSNNACGAATATCGCCAATSNNSNNSNNSNNSNNSNNSNNSNN<br>SNNSNNSNNSNNSNNSNNSNNSNNSNNSNNSNNSNNSNNSNNSNNSNNSNNSNN<br>SNNSNNSNNCGCCATTTAGCTGTCCTCCTTACTAA |
| Less Biased | L-E-3 | AXXXXXXXIGDIRXXXXXXXXXXXXXXXXXXXXXXXXXXXXXXXXXXXX |
| | | GTGATGGTGGCCTAAGCTACCGGAGCCSNNSNNSNNSNNSNNSNNSNNSNNSNN<br>SNNSNNSNNSNNSNNSNNSNNSNNSNNSNNSNNSNNSNNSNNSNNSNNSNNSNN<br>SNNSNNSNNSNNSNNSNNSNNSNNSNNACGAATATCGCCAATSNNSNNSNNSNN<br>SNNSNNSNNCGCCATTTAGCTGTCCTCCTTACTAA |
| | L-M-3 | AXXXXXXXXXXXXXXXXXXIGDIRXXXXXXXXXXXXXXXXXXXXXXXXX |
| | | GTGATGGTGGCCTAAGCTACCGGAGCCSNNSNNSNNSNNSNNSNNSNNSNNSNN<br>SNNSNNSNNSNNSNNSNNSNNSNNSNNSNNSNNSNNSNNSNNSNNSNNACGAAT<br>ATCGCCAATSNNSNNSNNSNNSNNSNNSNNSNNSNNSNNSNNSNNSNNSNNSNN<br>SNNSNNSNNCGCCATTTAGCTGTCCTCCTTACTAA |
| | L-L-3 | AXXXXXXXXXXXXXXXXXXXXXXXXXXXXXIGDIRXXXXXXXXXXXXXX |
| | | GTGATGGTGGCCTAAGCTACCGGAGCCSNNSNNSNNSNNSNNSNNSNNSNNSNN<br>SNNSNNSNNSNNSNNACGAATATCGCCAATSNNSNNSNNSNNSNNSNNSNNSNN<br>SNNSNNSNNSNNSNNSNNSNNSNNSNNSNNSNNSNNSNNSNNSNNSNNSNNSNN<br>SNNSNNSNNCGCCATTTAGCTGTCCTCCTTACTAA |
| | L-0-3 | AXXXXXXXXXXXXXXXXXXXXXXXXXXXXXXXXXXXXXXXXXXXXXXXX |
| | | GTGATGGTGGCCTAAGCTACCGGAGCCSNNSNNSNNSNNSNNSNNSNNSNNSNN<br>SNNSNNSNNSNNSNNSNNSNNSNNSNNSNNSNNSNNSNNSNNSNNSNNSNNSNN<br>SNNSNNSNNSNNSNNSNNSNNSNNSNNSNNSNNSNNSNNSNNSNNSNNSNNSNN<br>SNNSNNSNNCGCCATTTAGCTGTCCTCCTTACTAA |

a Each peptide sequence is followed by −GSGSLGHHHHHHRL.

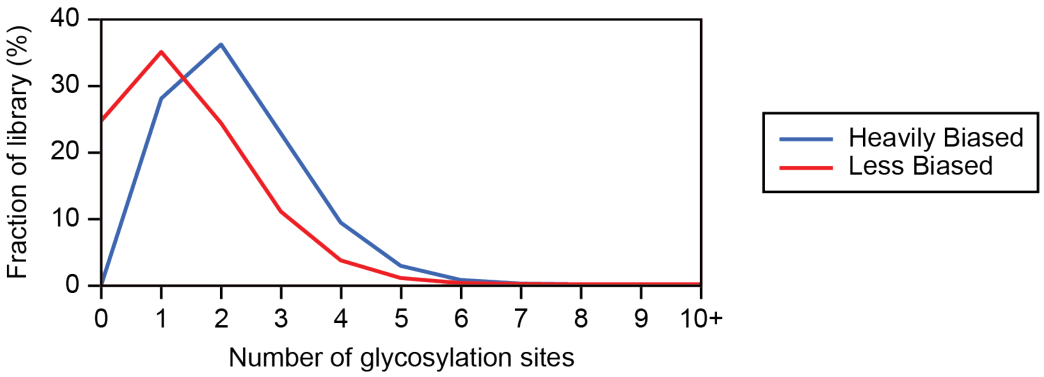

**Figure S4**. Theoretical multivalency distributions for Heavily Biased and Less Biased libraries of PGT128 selection.

**Table S3**. List of primers used in preselection experiments and PGT128 selection.

| Name | Sequence 5′ to 3′ [a] |
|---|---|
| JB-HL-fwd-X | TAATACGACTCACTATAGGGTTAACTTTAGTAAGGAGGACAGCTAA |
| JB-H-rev-X | CTAGCTACCTATAGCCGGTGGTGATGGTGGTGATGACCCAGAGAAC |
| JB-L-rev-X | CTAGCTACCTATAGCCGGTGGTGATGGTGATGGTGGCCTAAGCTAC |
| Library-FP1 | TAATACGACTCACTATAGGGTTAACTTTAGTAAGGAGG |
| F/H-RP | CTAGCTACCTATAGCCGGTGGTGATGGTGGTGATGACCCAGAG |
| V/L-RP | CTAGCTACCTATAGCCGGTGGTGATGGTGATGGTGGCCTAAGC |
| H-RT-RP | TTTTTTTTTTTTTTTGTGATGGTGGTGATGACCCAGAG |
| L-RT-RP | TTTTTTTTTTTTTTTGTGATGGTGATGGTGGCCTAAGC |

**Table S4**. Extended library DNA sequences. The full sense DNA sequences after PCR with the extending primers are shown, with the added constant regions underlined.

| Library | Name | Sequence 5′ to 3′ |
|---|---|---|
| Heavily Biased | H-E-2 full | <u>TAATACGACTCACTATAGGGTTAACT</u>TTAGTAAGGAGGACAGCTAAATGGCG<br>NNSNNSNNSNNSNNSNNSNNSATTGGCGATATTCGTNNSGCGNNSTGCATGN<br>NSNNSNNSNNSNNSNNSNNSNNSNNSNNSNNSNNSNNSNNSNNSNNSNNSNN<br>SNNSNNSNNSNNSNNSNNSNNSNNSNNSNNSNNSNNSNNSGGCTCCGGTTCT<br>CTGGGTCATCACCAC<u>CATCACCACCGGCTATAGGTAGCTAG</u> |
| | H-M-2 full | <u>TAATACGACTCACTATAGGGTTAACT</u>TTAGTAAGGAGGACAGCTAAATGGCG<br>NNSNNSNNSNNSNNSNNSNNSNNSNNSNNSNNSNNSNNSNNSNNSNNSNNSN<br>NSATTGGCGATATTCGTNNSGCGNNSTGCATGNNSNNSNNSNNSNNSNNSNN<br>SNNSNNSNNSNNSNNSNNSNNSNNSNNSNNSNNSNNSNNSGGCTCCGGTTCT<br>CTGGGTCATCACCAC<u>CATCACCACCGGCTATAGGTAGCTAG</u> |
| | H-L-2 full | <u>TAATACGACTCACTATAGGGTTAACT</u>TTAGTAAGGAGGACAGCTAAATGGCG<br>NNSNNSNNSNNSNNSNNSNNSNNSNNSNNSNNSNNSNNSNNSNNSNNSNNSN<br>NSNNSNNSNNSNNSNNSNNSNNSNNSNNSNNSNNSATTGGCGATATTCGTNN<br>SGCGNNSTGCATGNNSNNSNNSNNSNNSNNSNNSNNSNNSGGCTCCGGTTCT<br>CTGGGTCATCACCAC<u>CATCACCACCGGCTATAGGTAGCTAG</u> |
| Less Biased | L-E-3 full | <u>TAATACGACTCACTATAGGGTTAACT</u>TTAGTAAGGAGGACAGCTAAATGGCG<br>NNSNNSNNSNNSNNSNNSNNSATTGGCGATATTCGTNNSNNSNNSNNSNNSN<br>NSNNSNNSNNSNNSNNSNNSNNSNNSNNSNNSNNSNNSNNSNNSNNSNNSNN<br>SNNSNNSNNSNNSNNSNNSNNSNNSNNSNNSNNSNNSNNSGGCTCCGGTAGC<br>TTAGGCCACCATCAC<u>CATCACCACCGGCTATAGGTAGCTAG</u> |
| | L-M-3 full | <u>TAATACGACTCACTATAGGGTTAACT</u>TTAGTAAGGAGGACAGCTAAATGGCG<br>NNSNNSNNSNNSNNSNNSNNSNNSNNSNNSNNSNNSNNSNNSNNSNNSNNSN<br>NSATTGGCGATATTCGTNNSNNSNNSNNSNNSNNSNNSNNSNNSNNSNNSNN<br>SNNSNNSNNSNNSNNSNNSNNSNNSNNSNNSNNSNNSNNSGGCTCCGGTAGC<br>TTAGGCCACCATCAC<u>CATCACCACCGGCTATAGGTAGCTAG</u> |
| | L-L-3 full | <u>TAATACGACTCACTATAGGGTTAACT</u>TTAGTAAGGAGGACAGCTAAATGGCG<br>NNSNNSNNSNNSNNSNNSNNSNNSNNSNNSNNSNNSNNSNNSNNSNNSNNSN<br>NSNNSNNSNNSNNSNNSNNSNNSNNSNNSNNSNNSATTGGCGATATTCGTNN<br>SNNSNNSNNSNNSNNSNNSNNSNNSNNSNNSNNSNNSNNSGGCTCCGGTAGC<br>TTAGGCCACCATCAC<u>CATCACCACCGGCTATAGGTAGCTAG</u> |
| | L-0-3 full | <u>TAATACGACTCACTATAGGGTTAACT</u>TTAGTAAGGAGGACAGCTAAATGGCG<br>NNSNNSNNSNNSNNSNNSNNSNNSNNSNNSNNSNNSNNSNNSNNSNNSNNSN<br>NSNNSNNSNNSNNSNNSNNSNNSNNSNNSNNSNNSNNSNNSNNSNNSNNSNN<br>SNNSNNSNNSNNSNNSNNSNNSNNSNNSNNSNNSNNSNNSGGCTCCGGTAGC<br>TTAGGCCACCATCAC<u>CATCACCACCGGCTATAGGTAGCTAG</u> |

*2. Preparation of Puromycin-modified RNA*

Several batches of T7 transcription were carried out to produce sufficient RNA for subsequent steps. As a representative example, 75 pmol of library DNA was used for a T7 transcription reaction assembled on ice with 1X Transcription buffer (5X: 400 mM HEPES-KOH, pH 7.6, 125 mM $MgCl_2$, 200 mM DTT, 10 mM spermidine), 50 mg/mL PEG-8000, 4 mM of each NTP, 2.5 U/mL Inorganic Pyrophosphatase, 100 nM purified dsDNA, and 0.05 µg/µL T7 RNA Polymerase, with water added to 1 mL volume. The reaction was incubated at 37 °C overnight. DNA template was removed by incubation with Turbo DNase (Invitrogen) at 37 °C for 15 minutes, and the reaction was quenched with EDTA. The crude transcripts were precipitated with isopropanol followed by a 70% (v/v) ethanol rinse. The library RNA pellets were dissolved in 8 M urea and purified on a 5% denaturing urea-PAGE gel. Note: The glass plates of the gel were soaked in 1 N NaOH for 1 hour prior to use. The RNA bands were visualized with a 254 nm handheld UV lamp and cut out with fresh razors for electroelution with Elutrap Electroelution System (Whatman) with 0.5X TBE running buffer. The eluents were precipitated with isopropanol followed by a 70% (v/v) ethanol wash. The library pellets were dissolved in 140 µL of water and quantified by NanoDrop. In this representative experiment, yields were 2.5-3.7 nmol. The purified RNA was combined with other batches for puromycin modification.

Purified library RNA (4.8 nmol) was subjected to puromycin modification via photo-crosslinking with psoralen (**Figure S5**). Reactions (1600 µL) were assembled on ice with 3 µM Library RNA and 7.5 µM XL-PSO oligonucleotide [(C6 psoralen)-2′-OMe(UAGCCGGUG)$(dA)_{15}$(Spacer $9)_2$dA$(dC)_2$-Puromycin, purchased from Keck] in 1X XL Buffer (10X XL buffer: 200 mM HEPES-KOH, 1 M KCl, 10 mM spermidine, 10 mM EDTA, pH 7.5). The reaction mixtures were aliquoted into 8-strip 0.2 mL PCR tubes (50 µL each) and annealed on a thermal cycler (Bio-Rad S1000). Reactions were heated at 70 °C for 3 minutes, then slowly cooled to 25 °C with a ramp of 0.1 °C/s, incubated at 25 °C for 5 minutes, and then chilled at 4 °C. The annealed solutions were transferred to a Costar 96-well plate (100 µL per well), and a sample (5 µL) was set aside for gel analysis. The plate was placed on an ice water bath prepared in an empty pipette-tip box lid. In the 4 °C room, a handheld UV lamp was placed on top of the plate to irradiate the solutions at 365 nm for 20 minutes. The solutions from the wells were combined, and a sample (5 µL) was set aside for gel analysis. Next, the solutions were precipitated with isopropanol followed by a 70% (v/v) ethanol wash before 5% denaturing urea-PAGE purification. Bands were visualized by RNA Staining Solution (Abnova). The crosslinked RNA ("XL-RNA") bands were cut out and electroeluted with Elutrap Electroelution System (Whatman) with 0.5X TBE running buffer. The eluents were collected and precipitated with ethanol followed by a 70% (v/v) ethanol wash. The XL-RNA pellets were dissolved in 100 µL of water and quantified by NanoDrop. Yields ranged from 1050-1695 pmol. The purified XL-RNA was analyzed on a 5% denaturing urea-PAGE with comparison to the saved samples of non-XL RNA and crude XL-RNA.

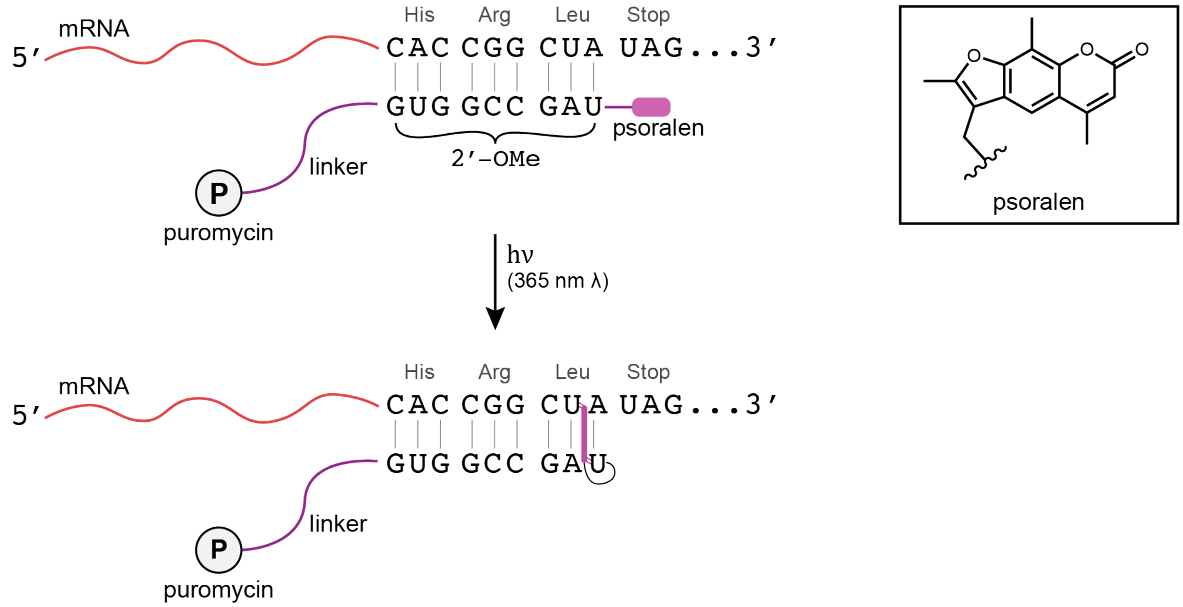

**Figure S5**. Modification of mRNA with puromycin-containing oligonucleotide.

*3. Translation for the Formation of Library mRNA-peptide Fusions*

Purified XL-RNA was subjected to mRNA-display translation with homemade PURE system with the inclusion of homopropargylglycine (HPG). As a representative example, 1500 pmol of XL-RNA was used for a translation reaction (1500 µL) assembled as previously described[3] with the addition of 15 µM MAP, 6 µM PDF, and 100 µM $CoCl_2$ for the cleavage of the N-terminal HPG and with 3.3 µM [2,5-$^3$H]-L-histidine (Moravek Biochemicals) for radiolabeling. The translation reactions were incubated at 37 °C for 30 minutes. Next, 450 µL (0.3 volumes) of a solution of 2.05 M KCl and 172 mM $Mg(OAc)_2$ were added. The mixtures were incubated at room temperature for 15 minutes and then stored at -20 °C overnight to allow for efficient fusion formation.

*4. Oligo(dT) Purification and Cyclization*

Using the 1500 µL translation plus 450 µL $Mg^{2+}/K^+$ added as a representative example, the following is the general protocol used for oligo(dT) purification and cyclization: A sample of the crude translation was set aside (48 µL) for analytical purposes. Six portions of 250 µL each of Oligo $d(T)_{25}$ Magnetic Beads (New England Biolabs) were placed into 2 mL microcentrifuge tubes. The beads were equilibrated twice with 1 mL of 1X oligo(dT) binding buffer with high EDTA (20 mM Tris-HCl, 1 M NaCl, 50 mM EDTA, pH 8.0, 0.2% (v/v) Triton X-100, 5 mM BME). The beads were resuspended in 317 µL of 2X oligo(dT) binding buffer with high EDTA and 317 µL of crude translation was added to each. The suspensions were tumbled at room temperature for 30 minutes. The supernatants were removed, and each of the tubes with beads were washed four times with 1268 µL of oligo(dT) binding buffer with TCEP (20 mM Tris-HCl, 1 M NaCl, 10 mM EDTA, pH 8.0, 0.2% (v/v) Triton X-10, 0.5 mM TCEP). The beads were then washed three times with 1268 µL of oligo(dT) wash buffer with TCEP (20 mM Tris-HCl, 300 mM NaCl, 10 mM EDTA, pH 8.0, 0.1% (v/v) Tween-20, 0.5 mM TCEP). Next, 1268 µL of cyclization buffer (20 mM Tris-HCl, pH 8.0, 660 mM NaCl, 0.2% (v/v) Triton X-100, 0.5 mM TCEP, 3.3 mM m-dibromoxylene, 33% (v/v) MeCN) was added, and the suspension was tumbled at room temperature for 30 minutes. The supernatant was removed, and 1268 µL of cyclization wash buffer with BME (20 mM Tris-HCl, pH 8.0, 660 mM NaCl, 0.2% (v/v) Triton X-100, 10 mM BME, 33% (v/v) MeCN) was added. The suspension was tumbled at room temperature for 10 minutes to quench the reaction. The supernatant was removed, and the beads were washed twice with 1268 µL portions of oligo(dT) wash buffer with BME/no EDTA (20 mM Tris-HCl, 300 mM NaCl, 0.1% (v/v) Tween-20, 10 mM BME). mRNA-peptide fusions were eluted from the beads with 317 µL of 0.1% (v/v) Tween-20. The elution step was repeated five times.

The yields of oligo(dT) purification and cyclization, as well as subsequent steps, were calculated using liquid scintillation counting (LSC) measurements as previously described.[3] As the random libraries had a varying number of histidine residues within each peptide sequence, the average number of histidine residues was estimated to complete the specific activity calculation. Each library had six fixed histidine residues as part of the affinity tag on the C-terminus. Next, the probability of a histidine residue appearing in the random region was calculated. As histidine is encoded by 1/32 NNS codons, with 40 random NNS codons in the Heavily Biased library, 1.25 random histidine residues would be present on average. The Less Biased library comprised three libraries with 43 random NNS codons and one library with 48 NNS codons. Assuming equal representation of the four pools within the Less Biased library, 1.38 histidine residues would be present in the random regions. Adding the six fixed histidine residues, the average number of histidine residues was 7.25 for the Heavily Biased library and 7.38 in the Less Biased library.

In the oligo(dT) purification and cyclization example described, the yields were 18.2 pmol and 12.9 pmol for the Heavily Biased and Less Biased libraries, respectively. The fusion formation (i.e. mRNA-peptides formed) is determined from the yield of oligo(dT) purification, with the theoretical yield as the pmol of input XL-RNA. In the representative case, 1500 pmol of XL-RNA were used in translation, corresponding to fusion formation rates of 1.2% and 0.9% for the Heavily Biased and Less Biased libraries, respectively.

*5. Reverse Transcription*

For each library, the first two eluent fractions from oligo(dT) purification and cyclization were combined, totaling 3.8 mL. The eluents were passed through an 0.22 µm Ultrafree-MC Centrifugal Filter (MilliporeSigma) into 10 separate 1.5 mL microcentrifuge tubes to remove residual magnetic beads. A portion of the filtered eluents (50 µL total) was saved for LSC and SDS-PAGE. Next, the mRNA-peptide fusions were precipitated with isopropanol in the presence of linear acrylamide carrier, followed by a 70% (v/v) ethanol wash. The pellets were allowed to dry at room temperature before dissolving in 27.5 µL of 0.18% (v/v) Triton X-100 such that the final concentration of Triton X-100 in the 50 µL reaction would be 0.1% (v/v). To the dissolved fusions were then added 2.5 µL of a mix of 10 mM each dNTP to a final concentration of 0.5 mM each and 6.25 µL of 80 µM reverse transcription primer to a final concentration of 10 µM (H-RT-RP for the for Heavily Biased library, L-RT-RP for the Less Biased library). The solutions were heated at 65 °C for 5 minutes on a metal heat block and then incubated on ice for >5 minutes. Next, 10 µL of 5X Reaction Buffer (ThermoScientific), 1.25 µL of 40 U/µL of RiboLock RNase Inhibitor (ThermoScientific), and 2.5 µL of 200 U/µL of RevertAid Reverse Transcriptase (ThermoScientific) were added, with final concentrations of 1X, 1 U/µL, and 10 U/µL, respectively. The reactions were incubated at 42 °C on a metal heat block for 30 minutes, chilled on ice, and then diluted to 200 µL with water. A total of 25 µL was saved for LSC and SDS-PAGE. Yields were 14.3 pmol and 10.1 pmol for the Heavily Biased and Less Biased libraries, respectively.

Note: For subsequent rounds of selection, the reverse transcription protocol was altered slightly from that described above. Instead of addition before heating at 65 °C, the dNTP mix was added after (with Reaction Buffer, RiboLock, and RevertAid).

*6. Ni-NTA Agarose Purification*

The cDNA/mRNA-peptide fusions were precipitated with ethanol in the presence of linear acrylamide carrier for Ni-NTA affinity purification. Each of the pellets in the 10 tubes was dissolved in 17.5 µL of denaturing bind buffer (100 mM $NaH_2PO_4$, 10 mM Tris-HCl, 6 M guanidinium hydrochloride, pH 8.0, with 0.2% (v/v) Triton X-100, 5mM BME). Next, 75 µL of HisPur™ Ni-NTA Resin (ThermoScientific) was equilibrated as follows: 600 µL of denaturing bind buffer was added to the resin, spun down at 21,000 ×g for 30 seconds, and the supernatant was removed. This wash was repeated once more. The resin was resuspended in 200 µL of denaturing bind buffer and divided into the 10 tubes with fusions (20 µL each) for a volume of 37.5 µL per tube (altogether: 375 µL). The suspensions were tumbled at room temperature for 1 hour. Next, each of the 10 suspensions were transferred into one 0.22 µm Ultrafree-MC Centrifugal Filter (MilliporeSigma), and the filter was spun down at 2,000 ×g for 30 seconds to collect the flow-through fraction. The tubes from which the resin suspesions had been taken were each rinsed with 15 µL of denaturing bind buffer each (total: 150 µL) and transferred to the filter unit. The filter was spun down at 2,000 ×g for 30 seconds. The tubes were rinsed with denaturing bind buffer once more. Next, the resins in the filter unit were washed with 150 µL of wash buffer (50 mM $NaH_2PO_4$, pH 8.0, 300 mM NaCl, 20 mM imidazole, 5 mM BME) and spun down at 2,000 ×g for 30 seconds. This wash was repeated twice more with the last centrifugation at 10,000 ×g instead of 2,000 ×g. The cDNA/mRNA-peptide fusions were eluted from the resins as follows: the filter was placed into a new 1.5 mL microcentrifuge tube, 75 µL of native elute buffer (50 mM $NaH_2PO_4$, 300 mM NaCl, 250 mM imidazole, pH 8.0, with 0.2% (v/v) Triton X-100, 5 mM BME) was added, and the filter was spun down at 10,000 ×g for 30 seconds. The elution step was repeated twice more with collection of each eluent in a separate 1.5 mL microcentrifuge tube. A portion of the flow-through, washes, and eluents, as well as the filter with resin, were measured by LSC. The combined eluents of the Heavily Biased and Less Biased libraries contained 4.6 pmol and 4.0 pmol fusions, respectively, while the flow-through fractions contained 3.1 pmol and 2.0 pmol fusions, respectively.

Ni-NTA agarose purification was repeated to retrieve more cDNA/mRNA-peptides from the flow-through fractions. To the flow-through fraction (375 μL) was added 75 μL of fresh Ni-NTA resin. The suspension was tumbled at room temperature for 1 hour before transferring to a new centrifugal filter unit. Wash and elution steps were carried out as before. A portion of the flow-through, washes, and eluents, as well as the filter with resin, were measured by LSC. In the first eluent, another 1.1 pmol and 0.8 pmol of fusions were recovered for the Heavily Biased and Less Biased libraries, respectively.

Note 1: Resubjection of the flow-through was only done for the first three rounds of each selection.
Note 2: The wash buffer used contained 20 mM imidazole, which may elute some fusions from the Ni NTA agarose prior to elution. It is not recommended to add imidazole to the wash buffer.

*7. Gel Filtration*
All three elution fractions from Ni-NTA agarose purification, as well as the first eluent from resubjection, were desalted by gel filtration to remove imidazole. Per manufacturer instructions, a NAP5 column (GE Healthcare) was equilibrated with 10 mL of gel filtration buffer (10 mM Tris-HCl, pH 7.5, 1 mM EDTA pH 8.0, with 0.2% (v/v) Triton X-100 and 5 mM BME). The eluents (300 μL total) were then loaded onto the column and collected in a 1.5 mL tube as the first flow-through fraction. Next, 200 μL of gel filtration buffer was added and collected in a new 1.5 mL tube as the second flow-through fraction. The cDNA/mRNA-peptide fusions were then eluted from the column with 700 μL of gel filtration buffer, followed by a second elution (300 μL of gel filtration buffer), collected separately. Yields were 6.2 pmol and 5.1 pmol for the Heavily Biased and Less Biased libraries, respectively.

*8. Click Reaction of Fusion Batches I*
As previously discussed, several batches of fusions were prepared. A summary of the first 5 batches of cDNA/mRNA-peptides prior to the click reaction (just after gel filtration) is summarized in **Table S5**.

**Table S5**. Summary of cDNA/mRNA-peptides quantities after gel filtration as estimated by LSC.

| Batch | Heavily Biased Library (pmol) | Less Biased Library (pmol) |
|---|---|---|
| 1 | 1.20 | 1.02 |
| 2 | 4.08 | 3.50 |
| 3 | 1.44 | 1.44 |
| 4 | 10.41 | 6.12 |
| 5 | 6.17 | 5.09 |
| Sum: | 23.32 | 17.17 |

As the specific activity values were similar, batches 1, 2, and 3 and batches 4 and 5 were combined for the click reaction. A smaller portion (80-90 μL) of the combined batches were set aside for analysis. Batches 1-3 were combined and isopropanol precipitated into three tubes followed by 70% (v/v) ethanol wash, while batches 4-5 were combined and isopropanol precipitated into four tubes followed by 70% (v/v) ethanol wash. Next, the pellets of separate tubes for batches 1-3 and batches 4-5 were dissolved, recombined into a single tube, and ethanol precipitated again for the click reaction.

The click reaction with $Man_9(GlcNAc)_2$-azide was carried out under an inert atmosphere as previously described,[3,7] except using a 10 μL total volume in a 1.5 mL microcentrifuge tube. Reagents were prepared in several 0.5 mL centrifuge tubes and placed into a two-neck pointed-bottom flask filled with argon (**Figure S6**). The peptide pellet (Tube A) was dissolved in 3 μL of water, and 1 μL of 1 M HEPES-KOH, pH 7.6 and 1 μL of 50 mM aminoguanidine were added. To Tube B was added 2.8 μL of water, 0.5 μL of 20 mM $CuSO_4$, and 0.5 μL of 20 mM of Tris(3-hydroxypropyltriazolylmethyl)amine ligand (THPTA).[8] Next, $Man_9(GlcNAc)_2$-azide was added to Tube B (1.2 µL of a 25 mM solution). To Tube C was added 4.9 µL of water, 0.5 µL of 10 mM THPTA, and 0.6 µL of 25 mM $Man_9(GlcNAc)_2$-azide. A few milligrams of sodium-L-ascorbate powder was added to Tube D. The caps of Tubes A-D were cut off and the tubes were added to the two-neck pointed-bottom flask. The flask was capped with a septum fitted with an outlet needle and purged with argon for 1 hour to degas the solutions (**Figure S6B**). At that time, the septum was briefly removed to transfer the contents of Tube B to Tube A. The sodium-L-ascorbate powder (Tube D) was dissolved in degassed water to make a 100 mM solution. Next, 1 µL of the 100 mM sodium-L-ascorbate solution was added to Tube A. The septum with outlet needle was replaced, and the reaction was purged with argon for 15 minutes. Next, the outlet needle was removed to keep the system under positive argon pressure. The reaction was allowed to proceed for another 75 minutes. Next, the contents of Tube C were added to Tube A, as well as 0.5 µL of the 100 mM sodium-L-ascorbate solution. The septum with outlet needle was replaced and the reaction was purged with argon for 15 minutes. The outlet needle was removed, and the reaction was allowed to proceed for another 75 minutes under positive argon pressure. The reaction was quenched by addition of 2.5 µL of 10 mM EDTA (pH 8.0) and diluted with water. A small portion was used to quantify the glycosylated peptide by LSC for SDS-PAGE analysis and/or subsequent experiments.

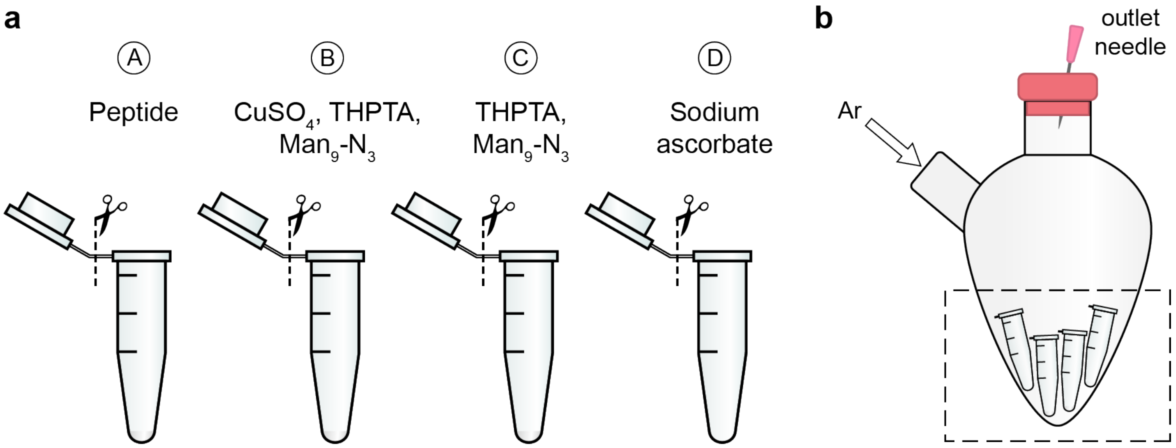

**Figure S6**. Click reaction set-up. (a) Separate tubes are prepared with click reagents, and the caps are removed; (b) The tubes are placed in a two-neck pointed-bottom flask with argon flow. An outlet needle is put in place to purge the system, then removed to keep the system under positive argon pressure.

Following the click reaction, each solution of glycosylated fusions was diluted to 150 µL. A portion of the solution was measured by LSC for quantification. Yields of batches 1-3 and batches 4-5 were 0.65 pmol and 3.25 pmol for the Heavily Biased library, respectively, and 0.85 pmol and 2.81 pmol for the Less Biased library, respectively. Loss of some fusions during this recombination process may have occurred due to adhesion to the plastic tubes.

*9. Nuclease-digestion of Fusions for SDS-PAGE*
The efficiency of the click reaction was analyzed by SDS-PAGE of nuclease-digested peptides. Saved samples of combined non-glycosylated fusions and glycosylated fusions for batches 1-3 and batches 4-5 were separately precipitated with ethanol in the presence of linear acrylamide. The pellets were dried and dissolved in 4 µL of 200 mM $NH_4OAc$, pH 5.3. Next, 1 µL of 1 U/µL Nuclease P1 from Penicillium citrinum (Sigma) was added and the samples were incubated at 37 °C for 1 hour. The reaction was quenched by addition of 2.5 µL of 1 M Tris-HCl, pH 7.8. Next, 7.5 µL of 2X Laemmli sample buffer with 5% BME was added to the samples for analysis on a 4-20% SDS-PAGE gel (Bio-Rad).

*10. Click Reaction and Analysis of Fusions Batches II*
To obtain sufficient yields of glycosylated library samples, three more batches (6-8) of fusions were prepared and subjected to the click reaction with $Man_9(GlcNAc)_2$-azide as described in **Section II.A.8**. Next, the old and new glycosylated fusion batches were resubjected to the click reaction to improve glycosylation efficiency. Each of the sample sets were separately analyzed by SDS-PAGE as well as combined.

**B. Round 1 Selection**

*1. Buffer Exchange and Quantification of Glycosylated Fusions Prior to Selection Round 1*
For each library, all glycosylated fusions from batches 1-8 were combined into a single 1.5 mL tube and the empty tubes rinsed with 50 µL of 0.2 % Triton X-100. The volume was reduced from ~1000 µL to 500 µL by speed-vac, and the fusions were then precipitated with isopropanol in the presence of linear acrylamide carrier followed by a 70% (v/v) ethanol wash. The pellets were dissolved in 153 µL 1X selection buffer (20 mM Tris-HCl, pH 7.5, 100 mM NaCl, 0.1% (v/v) Triton X-100), and yields based on LSC were calculated to be 7.3 and 6.8 pmol, equivalent to $4.4 \times 10^{12}$ and $4.1 \times 10^{12}$ sequences for Heavily Biased and Less Biased libraries, respectively. A portion of the library fusions were set aside (4.5 µL).

*2. Round 1 Selection*
To 148.5 µL of library fusions in 1X selection buffer were added PGT128, PGT130, and gl-PGT128 (the latter two being targets of parallel selections) to final concentrations of 200 nM each, with 1X selection buffer added to 200 µL. The mixture was tumbled at room temperature for 1 hour. While tumbling, Dynabeads™ Protein G (Invitrogen) were equilibrated with 1X selection buffer as follows: For each library, 150 µL of Protein G bead suspension was transferred to a 1.5 mL microcentrifuge tube. The tube was placed on a magnetic rack for 1 minute to separate the beads from solution, and the supernatant was removed. Next, 600 µL of 1X selection buffer was added to the beads. The mixture was resuspended by gently vortexing, spun down briefly, and placed on a magnetic rack for 1 minute. The supernatant was removed, and this wash was repeated two more times. The final supernatant was removed just prior to addition of the incubated library/antibody mixture. The mixture was allowed to tumble at room temperature for 30 minutes to capture antibodies and bound library fusions. Next, the suspension was briefly spun down, placed on a magnetic rack, and the supernatant was removed and saved. To wash the beads, 200 µL of 1X selection buffer was added, gently vortexed, spun down briefly, placed on a magnetic rack for 1 minute, and the wash was removed and saved. This wash step was repeated twice more for a total of three washes, each saved separately. The beads were resuspended in 200 µL of PCR mix A (1X Standard Taq Buffer, 0.1 mM each dNTP, 0.1% (v/v) Triton X-100) for PCR-based cDNA recovery from beads, as described in **Section II.B.4**.

*3. Determination of Fraction Bound*
The bead mixture in PCR mix A was resuspended by pipette, and 4 µL was transferred to a new 1.5 mL tube with 50 µL denaturing elute buffer (10 mM Tris-HCl, pH 7.5, 1 mM EDTA pH 8.0, 0.2% (w/v) SDS) to elute a small portion of fusions to measure the bound fraction. The suspension was heated at 95 °C for 5 minutes to denature antibodies, spun down briefly, placed on a magnetic rack for 1 minute, and the eluent was directly placed into a vial with liquid scintillation cocktail. The beads were washed with 50 µL of denaturing elute buffer, and the wash was placed into the same vial with liquid scintillation cocktail as the eluent. The beads were resuspended in 50 µL of denaturing elute buffer and placed into a separate vial with liquid scintillation cocktail for radioactivity measurements. The eluent with wash and the beads represented the "bound" fraction. Portions of the supernatant (4 µL) and washes (4 µL each) were measured by LSC as the "unbound" fraction. The tube in which the antibody and library fusions were incubated was also measured by LSC, but not included in calculations of fraction bound. The fraction bound was 18.1% for the Heavily Biased library and 14.8% for the Less Biased library.

*4. Recovery of Bound cDNA from Round 1 Selection*
Another 400 µL of PCR mix A was added to dilute the bead-bound fraction to about 600 µL. Approximately 150 µL of resuspended beads were used in pilot PCR experiments to determine the optimal PCR conditions for cDNA recovery used below. The remaining 450 µL of resuspended beads in PCR mix A were mixed with 2 parts PCR mix A and 1 part PCR mix B (1X Standard Taq Buffer, 0.1 mM each dNTP, 4 µM forward primer (Library FP1), 4 µM reverse primer (F/H-RP for Heavily Biased library; V/L-RP for Less Biased library), 0.1 U/µL Taq (New England Biolabs), 0.1% (v/v) Triton X-100) such that final concentrations were 1X Standard Taq Buffer, 0.1 mM each dNTP, 1 µM forward primer, 1 µM reverse primer, 0.025 U/µL Taq, and 0.1% (v/v) Triton X-100. For the large-scale PCR reactions, the total 1.8 mL volume was divided into 72 aliquots of 25 µL. Samples were briefly vortexed to resuspend the beads just prior to placing on the thermal cycler running the following program: 94 °C for 5 minutes, 24 (Heavily Biased library) or 28 (Less Biased library) cycles of 94 °C for 30 seconds, 73 °C for 30 seconds, and 74 °C for 30 seconds. The program was paused after the 5-minute initial denaturation step to briefly vortex the samples again before resuming the program. To maximize the total PCR product, some samples from pilot PCR experiments were subjected to further PCR to achieve the same number of cycles as in the large-scale.

PCR-amplified library DNA was isolated by removing the supernatant from magnetically isolated beads and then washing the beads once with 3-20 uL 0.1% (v/v) Triton X-100. Library DNA solutions were filtered by 0.22 µm Ultrafree-MC Centrifugal Filter (MilliporeSigma) to remove residual beads, extracted with phenol/chloroform and then chloroform, and precipitated with isopropanol followed by rinsing of pellets with 70% (v/v) ethanol. Yields were estimated by comparison to samples with known concentration on 6% native PAGE and determined to be approximately 38 pmol for each library.

For the next round of selection, a portion of the recovered DNA (12 pmol) was further amplified to obtain enough RNA for fusion preparation (including for other parallel selections). Several pilot experiments were carried out to optimize conditions for PCR. Ultimately, a 50-fold dilution of recovered DNA with 8 cycles provided enough material for subsequent experiments (75 to 94 pmol), though higher molecular weight bands were present in the samples, indicating the samples were cycled too much.

#### C. Subsequent Rounds of Selection

*1. General*
Fusions were prepared for the subsequent rounds of selection in a similar manner as the fusions for the first round. Fusion formation was much more efficient in the second round (4.3-6.0% in round 2 vs. 0.5-2.1% in round 1), thus improving overall yields and reducing the amount of translation volume required. Volumes of the subsequent transformations and purifications were decreased accordingly, except for the click reaction. The volume of the selection step was adjusted in each round to ensure excess of the antibody over the library fusions. Thus, as the concentration of the antibody decreased through selection, the volume of selection increased. For instance, the selection volume was 4.8 mL for round 9 with 2.5 nM PGT128 (12 pmol) with 4.9 or 5.6 pmol of library fusions. Following are procedures used for negative selections in subsequent rounds of selection.

*2. Negative Selection Against Immobilization Carrier*
Negative selections against the immobilization carrier were done just prior to the selection step as follows: Magnetic beads (Protein A/G or Streptavidin) were equilibrated with 1X selection buffer. Fusions in 1X selection buffer were added to the beads and tumbled at room temperature for 30 minutes. The supernatant was transferred to a new tube, and the beads were washed twice with 1X selection buffer with washes combined with the supernatant. The unbound fraction was directly used for selection with target.

*3. Negative Selection Against Non-Glycosylated Binders*
Fusions from gel filtration were concentrated by speedvac and precipitated with isopropanol in the presence of linear acrylamide carrier followed by a 70% (v/v) ethanol wash. The fusion pellets were dissolved in 1X selection buffer. Following negative selection against the immobilization carrier (described in **Section II.C.2**), the fusions were tumbled with 200 nM of antibody target for 1 hour at room temperature. The antibody-fusion complexes were captured on immobilization carrier for 30 minutes at room temperature. The supernatant was removed, and the beads were washed four times with 1x selection buffer. The combined supernatant and washes were filtered through a 0.22 µm Ultrafree-MC Centrifugal Filter (MilliporeSigma) and isopropanol precipitated followed by a 70% (v/v) ethanol wash for the click reaction.

### D. Next-generation Sequencing

*1. Preparation of DNA for NGS*

Recovered fraction bound DNA was minimally PCR amplified for sequencing analysis. DNA (generally 50-100 ng) was diluted around 100-fold into 200 µL PCR reactions with final 0.2 mM of each dNTP, 0.5 µM of each forward primer (Library-FP) and reverse primer (F/H-RP for Heavily Biased library; V/L-RP for Less Biased library), variable concentration of plasmid template, and 0.02 U/µL Phusion™ Hot Start II Polymerase (Thermo Scientific) in 1X Phusion HF Buffer. The following PCR protocol was used: 98 °C for 30 seconds, then 2-4 cycles of 98 °C for 5 seconds, 66 °C for 10 seconds, and 72 °C for 15 seconds. The crude PCR products were purified using a PCR clean-up kit (New England Biolabs) according to manufacturer instructions, eluting with 11 µL of 10 mM Tris-HCl, pH 8.0. The purified DNA was quantified by NanoDrop and assessed for purity by 8% native PAGE. DNA samples were submitted to GENEWIZ for Amplicon-EZ Next Generation Sequencing (NGS).

*2. Clustering and Analysis*

From the unique sequence identification and abundance analysis performed by GENEWIZ, we noticed that multivalency decreased in later rounds of selection (**Figure S7**). This is consistent with the library glycosylation patterns observed by SDS-PAGE (**Figure 3**). The average numbers of HPG residues found in the sequences shifted from 4.4 and 3.2 in round 7 to 2.9 and 2.1 in round 12 for the Heavily Biased and Less Biased libraries, respectively.

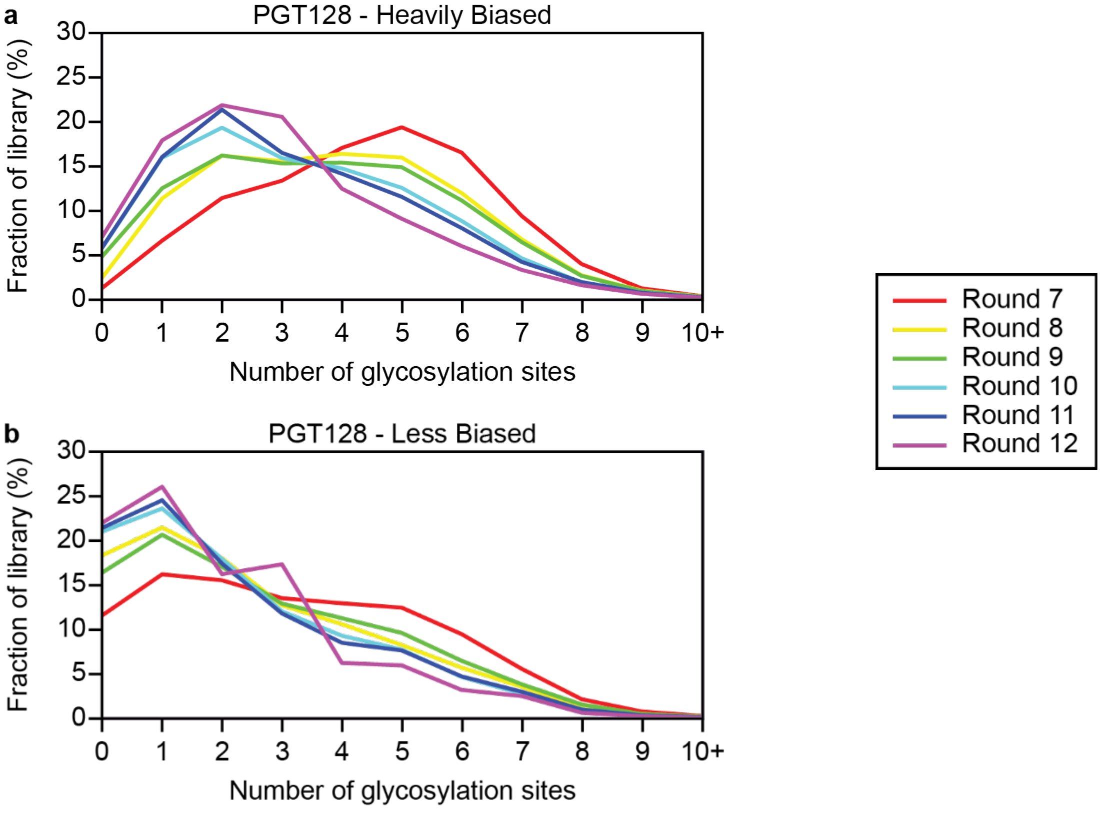

**Figure S7**. NGS analysis of multivalency through PGT128 selection for (a) the Heavily Biased library and (b) the Less Biased library.

Upon examination of sequences individually, we noticed many point mutations. These point mutations could be derived from sequencing read errors, PCR over several rounds of selection to regenerate the libraries, and/or PCR to generate samples for sequencing analysis. In order to account for these mutations, sequences were clustered into families of closely-related sequences. Starting from the raw Illumina paired-end reads generated by Amplicon-EZ and reported in FASTQ files, we used Paired-End reAd MergeR (PEAR v0.9.11)[9] to assemble the reads into 240 bp fragments using a PHRED quality score cutoff of 36 for trimming, p-value of 0.001, and minimum overlap of 200 bp. Typically, 90-95% of reads were successfully assembled into fragments. A collection of AWK and BASH scripts were used to orient the fragments into the forward direction, crop off the 5' and 3' constant regions, and align the fragments at their start codon. Nucleotide sequences were then translated into protein sequences using the standard codon table. Protein sequence libraries were clustered using Cluster Database at High Identity with Tolerance (CD-HIT v4.8.1)[10] with default settings and a minimum identity threshold of 0.875 (maximum 6 amino acid mismatch allowance). Finally, clustering results were analyzed using R (v3.6.3) and dplyr (v.1.0.7), and cluster sizes were normalized by total reads for each round.

For each cluster, a consensus sequence was generated using Biostrings (v2.54.0). Each consensus sequence was built position by position, based on which residue was most frequent at that position. In the event of a tie, the residue letter occurring earlier in the alphabet was chosen. As there are 20 amino acids, the consensus residue chosen would be present in at least 5% of the sequences. The consensus sequence could thus differ from the representative sequence chosen to create the cluster.

Cluster strength, or the sameness of the sequences within the cluster, was then evaluated based on the percent of sequences within the cluster having the consensus residue at a certain position. Each position was assigned a letter or number that represented a percentage range. The score range is as follows, where x is the consensus residue:

| | | | | | |
|---|---|---|---|---|---|
| 0: | | | x | $\leq$ | 85% |
| 1: | 85% | < | x | $\leq$ | 86% |
| 2: | 86% | < | x | $\leq$ | 87% |
| 3: | 87% | < | x | $\leq$ | 88% |
| 4: | 88% | < | x | $\leq$ | 89% |
| 5: | 89% | < | x | $\leq$ | 90% |
| 6: | 90% | < | x | $\leq$ | 91% |
| 7: | 91% | < | x | $\leq$ | 92% |
| 8: | 92% | < | x | $\leq$ | 93% |
| 9: | 93% | < | x | $\leq$ | 94% |
| a: | 94% | < | x | $\leq$ | 95% |
| b: | 95% | < | x | $\leq$ | 96% |
| c: | 96% | < | x | $\leq$ | 97% |
| d: | 97% | < | x | $\leq$ | 98% |
| e: | 98% | < | x | $\leq$ | 99% |
| f: | 99% | < | x | $\leq$ | 100% |

With this score guide, the most abundant cluster in the Heavily Biased library in round 12 was evaluated as follows, excluding N-terminal and C-terminal constant peptides:

| | **Cluster 1 – Heavily Biased, PGT128 round 12** |
|---|---|
| Representative | `YTEKHNGIGDIRPAICMNSKNQNYRCNHYQIKLYIHMLMRLSHNYRNS` |
| Consensus | `YTEKHNGIGDIRPAICMNSKNQNHRCNHYQIKLYIHMLMRLPHNYRNS` |
| Strength | `ebc7dacdddededdebcda6eb8eeebded5eeebeedbdbdcbdcd` |

For this cluster, the representative sequence and consensus sequence are not the same; the two differ in two locations (yellow highlights). Typically, the consensus sequence is also the most abundant sequence, as it is here.

The 20 most abundant clusters after round 12 are summarized in **Tables S6 and S7** with the consensus sequence and strength score. Starting from round 7, a clear increase in these clusters can be observed from round to round, though some clusters were not detected in earlier rounds (**Figure 4**).

**Table S6**. Top 20 most abundant sequence clusters in round 12 of PGT128 selection for the Heavily Biased library.

| PGT128 – Heavily Biased, Round 12 | | |
|---|---|---|
| ID | Consensus sequence and score | % of all sequences |
| 1 | YTEKHNGIGDIRPAICMNSKNQNHRCNHYQIKLYIHMLMRLPHNYRNS<br>ebc7dacdddededdebcda6eb8eeebded5eeebeedbdbdcbdcd | 6.47 |
| 11 | LTLRYLKIGDIRLANCMTVFPHFLSKKFFENGHRNLARPCTFRRNRHL<br>bcddededdeecdbddcdceceedebddcccdddecdddedcddccdde | 1.79 |
| 52 | YHKHRVMHHHEDKATSLTSNLVRLRLKTRIGDIRRALCMLSKFRYLIN<br>eebeedceddccbdddcdccdcddecb8bcdee8dce6cdddbceeec | 1.61 |
| 14 | LLHHLRMIGDIRPAHCMVSHQRRYVPISRKNVFFKRGFNSHPLRKILW<br>eeeceeededcddeedbddcdedaaecdceedddcee3ac5deecddd | 1.59 |
| 25 | RFRHSNNYYLTPFLTPLKTLISLQLRYRLIGDIRNASYMHKFSNRNRF<br>dddedddeeeeecdeeedddcadededeedeecedcd0acdddd6cdd | 1.24 |
| 5 | HQTHSYRIGDIRIAHLHGQPHAPVQGLPPVLRRRRELQVPLRARALLV<br>dddeddde ddddeceeeddddddceeeeedeeee00ddcddddceeb | 1.00 |
| 66 | IFNQGYRIKAWNDLKDIAIGDIRHALCMLVLARIKLQRRMVKYKHDHR<br>eebbefedee7ddd0cbdde0bee987cd9deec70ddcced7dd8dd | 0.88 |
| 9 | HQHHHPNYALMQRRLSIAIGDIRLAICMFAHLYHCYRKHLMANTIPMK<br>deddddddcecdbeddc9dddcdddedbcddaecdbedceddbbccdcc | 0.79 |
| 36 | FVTYQHMSQKNFRRYQILRNHFHPQNYRFIGDIRHALCMFIFKNLMRH<br>bdeeddcddddeeecbbeededecdcddcdeddedde9ccdcdedcdc | 0.76 |
| 103 | RLHHNIHSHPQKYLEHPLAHLAGHVLGRHHWRYSSGVVHGHRVRHLDQ<br>dc8eddedceedeefd0dedddddcdcaedddeddcdcbcdcddedddd | 0.68 |
| 375 | AMKIRSKIGDIRTAVCMFMHRHHHHHILDPYYLKMIVMYYSLKSRITL<br>cbcededcdbddd8d7ebddedddea9998daa97aaacaaa99a89a | 0.67 |
| 42 | FIKPCMMYLLPPTMLNLYIGDIRRAKCMEAMNNFHMNNKPLMATMPPH<br>bdeed0dde0eedddecde3ddbdddeefdeadddbbbdecbbb7ecd | 0.66 |
| 443 | KDILKLRIPFATLSGHRNIGDIRHAYCMSLKRPYIQVYSYLNHLKVRF<br>dceecdddeecce5bced6fddeded4accdecedcedceeeeedeed | 0.57 |
| 158 | TLHNIHDLNHYYRNLNTRIGDIRHATCMYFFMKLKLLKHNRFMDRAIY<br>becdeddddeeeedc0cdbecdecdd2ddbdbeedadd2cdcccecdd | 0.55 |
| 221 | PYRINQQMNFPWSSALFQIGDIRHARCMDSCRRFTNIMRYVYLKRRMN<br>edfee9b7ecdaebcbc7ed8cddcccacabedc6cb2ddbdddedcb | 0.47 |
| 22 | LFKPYPKIGDIRKARCMLQHTLHHRTNKQPSYRRRLKTLIPLFRRCML<br>ecdaeedeedbdefddacddcebdfddbeedddecebdeabbbedcbe | 0.46 |
| 247 | TNHLHRTIGDIRHAQCMIYLIYLVQNDQYKRNNRTFRLMLNPKLLKRF<br>eedeb6ddddddeddceaffdddfdebdcebeddeedddbebdceeddd | 0.45 |
| 61 | TNSYYHHNPLMRRTHVVMTLKPMNFWAKMIGDIRRAHCMTTINMLKRR<br>dccecfeaddccecdcd9de9dbadecaadcb9cbdededbabac8dd | 0.43 |
| 13 | ILLHVSTRSRYPHHHMAIIGDIRCASCMYPVLKWFYNFNRLKTYRKQF<br>eddcacecdeedeccaecbedbdeddd6cdde9dabdceeeb9cdacd | 0.41 |
| 74 | YRTHKLLHHHNDKWKSNIFPRIFVCHYYLIGDIRHARCMIPLEILRRY<br>ecdbd9eecdbdaa5ddadddbdeedde7dbcdecddd9cdebddece | 0.41 |

**Table S7**. Top 20 Most abundant sequence clusters in round 12 of PGT128 selection for the Less Biased library.

| PGT128- Less Biased, Round 12 | | |
|---|---|---|
| ID | Consensus sequence and score | % of all sequences |
| 8 | YSKHRFSFRHNNMLRDRKLIRKFSYHNHSIGDIRVANKFRYLHVFKFI<br>ecbddbddcbcbbedcddddddb9ededddddddddeddccedcdcdd5 | 6.48 |
| 1 | SIKLINQMMTTNPHLRLHIGDIRRLIKDLYMFRVYYRPTNSGRRLFVN<br>edecddcbacedeeeeeeeeeeeeeeeeeee0edededddddddedddd | 6.05 |
| 52 | HSHHHSPMIEFHSNGRLHIGDIRKFYADALMVLFFKMAFIDRIPFHDA<br>edededeeddbccd9edeeeeeeeeeeeeeedecbdddceddccccce | 2.82 |
| 67 | NIYFCSRRTNFHNSCYLMIGDIRGLSIYHHIMIHNKLHLLIMYNLLMM<br>dddbceddcccb6eddedeeddedec9dcccdccbddbdeccccceddd | 2.02 |
| 11 | IHLLPLRHNRRSHNRPSRLSWQKNDYFKSIGDIRATYWLRHNFLYRLS<br>eeeeeedddddedceedee8eeddbeddddeedeeddeeededdeeec | 1.71 |
| 61 | HVVVLHHSGFHGNRFSRLPKLLRNQHYQNIGDIRRLYNWIPYTKRYFQ<br>dcdbdcdbdcdcceddccedeeec0ee5ddddedbdb0cedededab | 1.29 |
| 2 | IHYHHPIIGDIRLKHNMINAHTKHVPQKLYLDIKFRRLFGLYILRMLN<br>eddedddeeddedcdaa0beddccbea0bcd8adeeeecddededced | 1.03 |
| 120 | IRKNFPMTFGHRPHLRVAHAQRAQHALLVLRRARRLLDQEVDAPGGRR<br>eecddd7beeddbdedcddecddedddeeecbeeeed9de9deeaee | 1.00 |
| 65 | ILYHYHNIGDIRRSQRHLNMQMRLYVSTLLHSSHTLRRASITHRIRKF<br>ebbcedbddddebccdbedccaeebd2bead4eddddcdddbeedd9d | 0.74 |
| 44 | TFSRYHTIGDIRHHTLKHHQSKGLQMRLIFLKRQFKAMGNCLRWKILF<br>deceeeeeeeededcbdededdbdcdcedbdcceddeeadcddecddde | 0.69 |
| 272 | TKDYRQKVRKIFSHHITKIGDIRLAEHQHFAKSRLLKGFVRARNRVRY<br>eebfdeeddcdec8eeddeeedcdedd4dededdeedd0edd0ceddc | 0.66 |
| 128 | HHYPNYHMRSHGDRLTLLRHLMSFLVDHKQILMFLLRMRKNHVSMMMT<br>bddb84ddebcdce9edeecedbdec5cddcdbcddebdcdd8adddb | 0.66 |
| 4 | KYTHIHSIGDIRNTYRNKHKHMALNKTNWALFQQHHRMLIRLFYRRLL<br>cd9dbcdebedddcdeaddbe9ddb4bcdc67dcdbecdceeeeedee | 0.51 |
| 15 | LNKHKHLRNHTRHHSVPTIGDIRKRIHNLLHYLAGFRFFNQMHSKMGV<br>fcdfeeeedf6cecbeadeeddebeddddddbdfebdeeddebbbbcc8 | 0.50 |
| 182 | IRNQTKKIGDIRGHHRTKPQYFEHPFVDLYKHYQYRVFHRGYLKLFRE<br>eddedddddedeeededecfdeddcdbede7ceeddeededdeeddee4 | 0.48 |
| 310 | YLHNHHNYSSNNKLHHLEIGDIRLIYQKYLRNPMFMTFLSRKHMNWQR<br>eeecdd0d0e6bcddddbedbddddd8bdcecbacab5cbdbdabdad | 0.43 |
| 1079 | NLTAMSRIGDIRKHHFGRPLYLTKHGAYPRYHTRYKHLLTYRHHFPLI<br>eedffffecefededdeeeddddcc8beede9ecbddfddebbdcfe | 0.42 |
| 98 | KHTHLRPMNFTQRLRKAHIGDIRLPRNISTSRIRTHIKFHLIRMHLRN<br>fedddce0d92dbdedfaddea6c0dddcecce9caedccfdbcbecd | 0.42 |
| 43 | FLLNHKRIGDIRKLPPLNLMATKTLTKERIRKIVNGFVQRLKGHSWWI<br>eeddddddddedeedfeddacfddddd7eddccdcecb5ded9deedb | 0.39 |
| 160 | IHHSYRGFTLRIPLTTNKIGDIRTAFPYPMLSHLFDRRRWKRGLHNWF<br>dfeefdededeaedcdceeeeedceeeddc48eec7dddd2efcdbea | 0.37 |

*3. Ranking Clusters by Scoring Function*

To help select sequences for synthesis and validation, clusters were scored based on the following function with variables defined in **Table S8**:

$$Score = 6 * \log(E_{7\to12}) + 2 * \log(E_{10\to12}) + 2 * M_{2-4} + 6 * H_{\leq6} - 8 * C_{odd} + 8 * C_2$$

**Table S8**. Scoring function variables.

| Symbol | Description |
|---|---|
| $E_{7\to12}$ | Fold increase in cluster frequency from round 7 to round 12. If the cluster was not present in round 7, then the frequency of the first instance the cluster was present in either round 8 or round 9 was used |
| $E_{10\to12}$ | Fold increase in cluster frequency from round 10 to round 12. If the cluster was not present in round 10, then the frequency from round 11 was used |
| $M_{2-4}$ | Boolean variable for whether the sequence contains between 2 and 4 (inclusive) AUG codons in the randomized region |
| $H_{\leq X}$ | Boolean variable for whether the sequence contains a number of histidine residues less than or equal to a set number ($X$) in the randomized region |
| $C_{odd}$ | Boolean variable for whether the sequence contains an odd number of cysteine residues in the randomized region |
| $C_2$ | Boolean variable for whether the sequence contains exactly 2 cysteine residues in the randomized region |

The top 20 scoring clusters are given in **Table S9** with their relative frequency each round plotted in **Figure S8**. We decided to synthesize H1, H22, and L1 as three of the four highest-scoring sequences. H31 was not selected because its two cysteine residues are very close together.

**Table S9**. Top 20 scoring clusters.*

| ID | Sequence | $E_{7\rightarrow12}$ | $E_{10\rightarrow12}$ | Score |
|---|---|---|---|---|
| H1 | YTEKHNGIGDIRPAICMNSKNQNYRCNHYQIKLYIHMLMRLSHNYRNS | 431.3 | 431.3 | 37.1 |
| H31 | FRYNMFDNLFRRSRHNDVRTRSLRKYSGAIGDIRSARCMNCYLKVRHK | 97.5 | 10.3 | 30.0 |
| H22 | LFKPYPRIGDIRKARCMLQHTLHHRTNKQPSYRRRLKTLIPHFRRCML | 115.0 | 5.8 | 29.9 |
| L1 | LIKLINQMMTTNPHLRLHIGDIRRLIKDLYMFRVYYRPTNSGGRLFVN | 756.4 | 137.5 | 29.5 |
| H409 | TIQHLRHYHGLGSYTNEMIGDIREALCMHSHTPFTWLWRMPVRDCLRN | 76.7 | 7.7 | 29.1 |
| H13 | ILLHVSTRNRYPHHHMATIGDIRCASCMYPVLKWFYNFNRLKTYRKQF | 102.5 | 2.8 | 29.0 |
| H455 | YNHYLTHVMIYVLTKAFVRRHKNLNEFFRIGGIRFAPCMMCRSFNQHN | 65.0 | 8.7 | 28.8 |
| H138 | FNHSGYHPSLHLRIHMRPIGDIRKAFCMSRMHGLNLVCFPLGLKRHTF | 67.5 | 4.2 | 28.2 |
| L6 | HDSVNRTLLCNAYMTSRRIGDIRCNDYQILIFNLMRFMSWKYMHMNNN | 103.7 | 10.4 | 28.1 |
| H42 | FIKPCVMYLLPPTMLNLYIGDIRRAKCMEAMNNFHMNNKPLMATMPPH | 94.3 | 12.5 | 28.0 |
| H622 | NNHANFEHSNNYHKLNFPRLVLTRIGLRLIGDIRNALCMNCMHKRLRR | 55.0 | 4.9 | 27.8 |
| L178 | TLIPMTTFRFLYHRYGDIGCYFCAMMSLPPRKSRHTQTHINKFNFAKR | 28.5 | 28.5 | 27.6 |
| H198 | VNNYLLYIGDIRCAICMYNLHMKKISTIFRHRFNKGYLKHFHRMYLLR | 60.0 | 2.8 | 27.6 |
| H11 | LTLRYLKIGDIRLANCMTVFPHFLSKKFFENGHRNLARPCTFRRNRHL | 99.4 | 3.7 | 27.1 |
| H62 | YLINPLSIGDIRHASCMNCPRAFMNRFVKSIHGKFFWHFFRNYHLKPN | 42.5 | 3.0 | 26.7 |
| H49 | FRNLFQRKHTRYLRSRMILAKIIRHQSLKIGDIRIAKCMLRCNTFRMQ | 30.9 | 5.7 | 26.4 |
| L67 | NIYFCSRRTNFHNSCYLMIGDIRGLSIYHHIMVHNKLHLLIMHNLLMM | 77.8 | 2.1 | 26.0 |
| L52 | HSHHHSPMIEFHSNGRLHIGDIRKFYADALMVLFFKMAFIDRIPFHDA | 941.3 | 941.3 | 25.8 |
| H152 | ILSHKFSIGDIRSACCMPKQGINYLRMISGMMHKSRPYRNQAYHLYHR | 30.0 | 2.7 | 25.7 |
| H350 | YHHSHFRIGDIRCATCMSLLKHSYPSSMLFNTNKKMRRLIFLRTIPFM | 30.0 | 1.9 | 25.4 |

* Calculated by scoring function as follows: $Score = 6 * \log(E_{7\rightarrow12}) + 2 * \log(E_{10\rightarrow12}) + 2 * M_{2-4} + 6 * H_{\leq6} - 8 * C_{odd} + 8 * C_2$

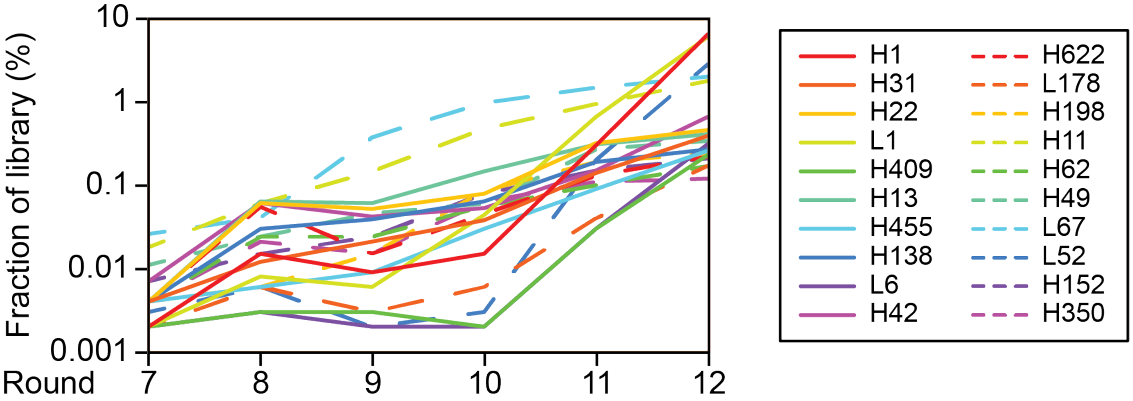

**Figure S8**. Enrichment of the top 20 scoring clusters from PGT128 selection.

### III. Synthesis of Individual Peptides and Glycopeptides from PGT128 Selection

#### A. Standard and Representative Procedures

*1. Materials*
For peptide synthesis, DMF (sequencing grade), TIPS (triisopropylsilane), EDT (ethane-1,2-dithiol), TCEP-HCl ((tris(2-carboxyethyl)phosphine hydrochloride)), *m*-dibromoxylene, iodoacetamide, DIC (N,N'-diisopropylcarbodiimide), pyridine, acetic anhydride, and phenol were purchased from Sigma-Aldrich, Acros, Alfa Aesar, TCI, or Fisher and used without further purification unless otherwise noted. All Fmoc-protected amino acids (except homopropargylglycine), piperidine, HATU, trifluoroacetic acid (TFA) (99.9%), and Oxyma were purchased from Chem-Impex International Inc. Fmoc-L-(homopropargyl)-Gly-OH[11] and $Man_9GlcNAc_2$-azide,[12] were prepared in-house. H-Rink amide Chemmatrix® resin was bought from Biotage and Sigma Aldrich. Milli-Q (MQ) ultrapure water was used for all the reactions in aqueous solutions. Pyridine and DIPEA were refluxed over $CaH_2$ and freshly distilled before use. Dichloromethane solvent was purified on a Pure Process Technologies solvent purification system. LC/LRMS analysis was performed on a Waters Acquity UPLC with reverse phase C4 column, and Waters Photodiode Array and Micromass ZQ4000 mass detectors. Peptides and glycopeptides were purified on Waters 2489 HPLC. Biolayer Interferometry (BLI) kinetics analysis was performed on a ForteBio BLItz instrument.

*2. Pre-Loading Resin with Fmoc-Lys(biotin)-OH for Solid-Phase Peptide Synthesis*

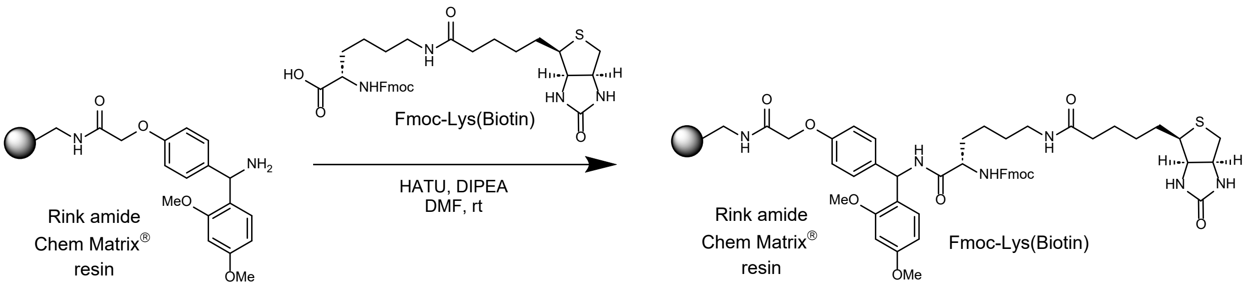

**Scheme S1**. Fmoc-Lys (biotin) loading on resin.

1 g (0.42 mmol/g) of H-Rink amide Chem-Matrix® resin was swelled in 30 mL dry dichloromethane in a 50 mL falcon tube for 30 minutes and then filtered. Fmoc-L-Lys(biotin)-OH (124 mg, 0.21 mmol) and HATU (175 mg, 0.46 mmol) in 10 mL of *N*, *N*-dimethylformamide was treated with DIPEA (183 µL, 1.05 mmol); this mixture was added to the resin immediately, and the suspension was agitated with shaking for 15 minutes at room temperature. The resin was filtered and washed twice with 10 mL of DMF and three times with 10 mL of DCM. The remaining amino groups were capped by acetylation with 10 mL of 60/40 v/v acetic anhydride/pyridine for 30 minutes. The resin was filtered and washed with DMF and DCM three times and kept under a high vacuum overnight.

*3. Resin Loading Determination*
To 6.5 mg of dry Fmoc-lysine(biotin)resin in a 4 mL glass vial was added 1 mL of 20% piperidine in DMF, and the suspension was allowed to react for 30 minutes. 100 µL of the reaction mixture was diluted with 900 µL DMF. A reference solution was prepared in the same way without the addition of resin. Absorbance for both solutions was recorded in a UV instrument at 301 nm wavelength (1 cm path length). The following formula for calculating the resin loading was used:
Fmoc loading(mmol/g) = [10*($Abs_{301(sample)}$ - $Abs_{301(reference)}$)*1mL] / ((7800 mL/mmol*cm)* 1 cm * g of resin sample)

*4. Microwave-Assisted Solid-Phase Peptide Synthesis*
Automated solid-phase peptide synthesis (SPPS) was performed on the Liberty Blue peptide synthesizer. The coupling and Fmoc deprotection were performed at 90 °C. Peptides were synthesized using DMF (ACS sequencing grade) as a solvent, Fmoc deprotection was done with piperidine in DMF (20/80, v/v) containing 0.1 M Oxyma, each coupling was performed using 0.2 M solution of Fmoc-amino acids (1.25 mL for each amino acid), 0.5 M Oxyma (0.5 mL) and 0.25 M DIC (1 mL) as coupling reagents. Double couplings were done for Arg and the last three amino acids in each sequence.

Amino acids used in SPPS (all L-): Fmoc-Ala-OH (A), Fmoc-Arg(Pbf)-OH (R), Fmoc-Asn(Trt)-OH (N), Fmoc-Asp(OMPE)-OH (D), Fmoc-Cys(Trt)-OH (C), Fmoc-Gln(Trt)-OH (Q), Fmoc-Gly-OH (G), Fmoc-His(Boc)-OH (H), Fmoc-Ile-OH (I), Fmoc-Leu-OH (L), Fmoc-Lys(Boc)-OH (K), Fmoc-Phe-OH (F), Fmoc-Pro-OH(P), Fmoc-Ser($^t$Bu)-OH (S), Fmoc-Thr($^t$Bu)-OH (T), Fmoc-Tyr-($^t$Bu)-OH (Y), Fmoc-Val-OH (V), Fmoc-HPG-OH (M).

*5. Resin Cleavage*

Resin after SPPS was transferred to a 50 ml falcon tube, 10 mL of freshly prepared cleavage cocktail TFA: EDT: water: TIPS (94:2.5:2.5:1) was added to the resin, and the tube was capped. After 3 hours at room temperature, the resin was filtered, and the cleavage cocktail was evaporated under a stream of nitrogen and triturated with cold ether to get a crude pellet, which was dissolved in 4 mL of 50:50 acetonitrile and water (containing 0.1% formic acid), filtered through a 0.22-micron filter and lyophilized overnight.

*6. Determination of Peptide/Glycopeptide Concentrations by UV NanoDrop*

$A_{280}$ Measurement: All measurements were recorded on a Thermo Scientific NanoDrop One instrument. For each sample or blank to be measured, 1 µL was added to the read position, and the absorbance at 280 nm was recorded. The molar extinction coefficient of each peptide/glycopeptide sequence was estimated by the number of aromatic side chains in amino acids.

The extinction coefficient at 280 nm was estimated as = number of Tryptophan (W) × 5500 + number of Tyrosine (Y) × 1490 (the small value for cysteine was not included in the calculation, as cysteines were alkylated in cyclization).

Peptide/glycopeptide concentrations were then calculated using Beer's law.

*7. Cyclization of Synthetic Peptides*

As a representative example, to the solution of H1-peptide (**S1**) (1.07 mg, 0.164 µmol) in 250 µL MQ water was added 70 µL of 3.4 mM TCEP-HCl solution in MQ water (1.5 equiv., 0.24 µmol). After 30 min, 320 µL of 200 mM aq. ammonium bicarbonate buffer, 240 µL of MeCN, and then 80 µL of 11 mM *m*-dibromoxylene (5.4 equiv., 0.88 µmol) solution in MeCN were added. The pH of the reaction was adjusted to 8 by the addition of 200 mM aq. ammonium bicarbonate buffer and checked with pH paper. Upon completion of the reaction, the reaction mixture was concentrated by lyophilization and redissolved in 200 µL of MQ water containing 0.1% FA for HPLC purification; this was purified by RP-HPLC on a Waters BEH-C4 column (10 X 250 mm, 300Å, 5 µm particle size) following a 95% A / 5% B to 65% A / 35% B gradient over 45 minutes with a flow rate of 4 mL/ min using 220 nm and 280 nm dual wavelength for UV-detection, where solvent A was water/ 0.1% formic acid and solvent B was acetonitrile/0.1% formic acid to afford 0.50 mg (0.075 µmol) of H1-cyclic peptide (**S2**) (46% yield) (quantified by UV-nanodrop). ESI-LRMS observed *m/z* of multiply charged ions 738.22 $[M+9H]^{9+}$, 830.46 $[M+8H]^{8+}$, 948.76 $[M+7H]^{7+}$, 1107.04 $[M+6H]^{6+}$, 1328.38 $[M+5H]^{5+}$ ; calculated average m/z for H1-cyclic peptide (**S2**) $C_{293}H_{445}N_{93}O_{79}S_3$ : 737.72 $[M+9H]^{9+}$, 829.81 $[M+8H]^{8+}$, 948.21 $[M+7H]^{7+}$, 1106.08 $[M+6H]^{6+}$, 1327.10 $[M+5H]^{5+}$.

*8. Click Glycosylation of Synthetic Peptides*

As a representative example, H1-cyclic peptide (**S2**) (0.19 mg, 0.028 µmol), $Man_9GlcNAc_2$-azide (2.6 uL, from 50 mM stock solution in water, 0.24 mg, 0.13 µmol), and aminoguanidine hemisulphate (14 µL from 100 mM stock solution in water, 1.4 µmol) were combined and lyophilized into 0.5 mL Eppendorf tube A. In the second 0.5 mL Eppendorf tube B, 1.8 µL (0.072 µmol) of a 40 mM solution of $CuSO_4$ and 0.9 µL (0.09 µmol) from a 100 mM solution of THPTA ligand were mixed and lyophilized. In the third 0.5 mL Eppendorf tube C, 5.6 µL of sodium ascorbate from a 250 mM stock solution (1.4 µmol) was lyophilized. Three tubes were placed in a two-neck pear flask, and a nitrogen atmosphere was set up by cycles of vacuum and nitrogen refill. Under nitrogen efflux, 14 µL of DMSO (degassed by freeze-pump-thaw) was added to tube A to dissolve sugar and peptide; 14 µL of water (degassed by freeze-pump-thaw) was added to tube B (producing a blue color). Next, the solution in tube B was transferred by pipette to tube C (blue color disappeared upon mixing), and the solution in tube C was immediately transferred to tube A and mixed by pipette (the final concentration for reaction was 1 mM in peptide). The reaction was checked by LCMS (typical reactions may take 2h to overnight to finish). After overnight incubation, the mixture was diluted with 200 µL water and injected into RP-HPLC (Waters BEH-C4, 5µm, 1.7 x 250 mm, 1 mL/ min, gradient of 2-30% (over 45 minutes) acetonitrile in water buffered with 0.1% formic acid to afford 65 µg (0.0051 µmol) of pure H1-glycopeptide (**1**) (18 %) yield after lyophilization (quantified by UV-nanodrop). ESI-LRMS observed *m/z* of multiply charged ions 1124.16 $[M+11H]^{11+}$, 1236.65 $[M+10H]^{10+}$, 1374.10 $[M+9H]^{9+}$, 1545.55 $[M+8H]^{8+}$, 1766.41 $[M+7H]^{7+}$; calculated average m/z for H1-glycopeptide (**1**) $C_{503}H_{796}N_{108}O_{244}S_3$ : 1124.32 $[M+11H]^{11+}$, 1236.66 $[M+10H]^{10+}$, 1373.95 $[M+9H]^{9+}$, 1545.57 $[M+8H]^{8+}$, 1766.22 $[M+7H]^{7+}$.

**B. Synthetic Scheme, Experimental Procedure, and LC-MS Chromatogram of Individual Peptides and Glycopeptides**

*1. H1-glycopeptide (**1**)*

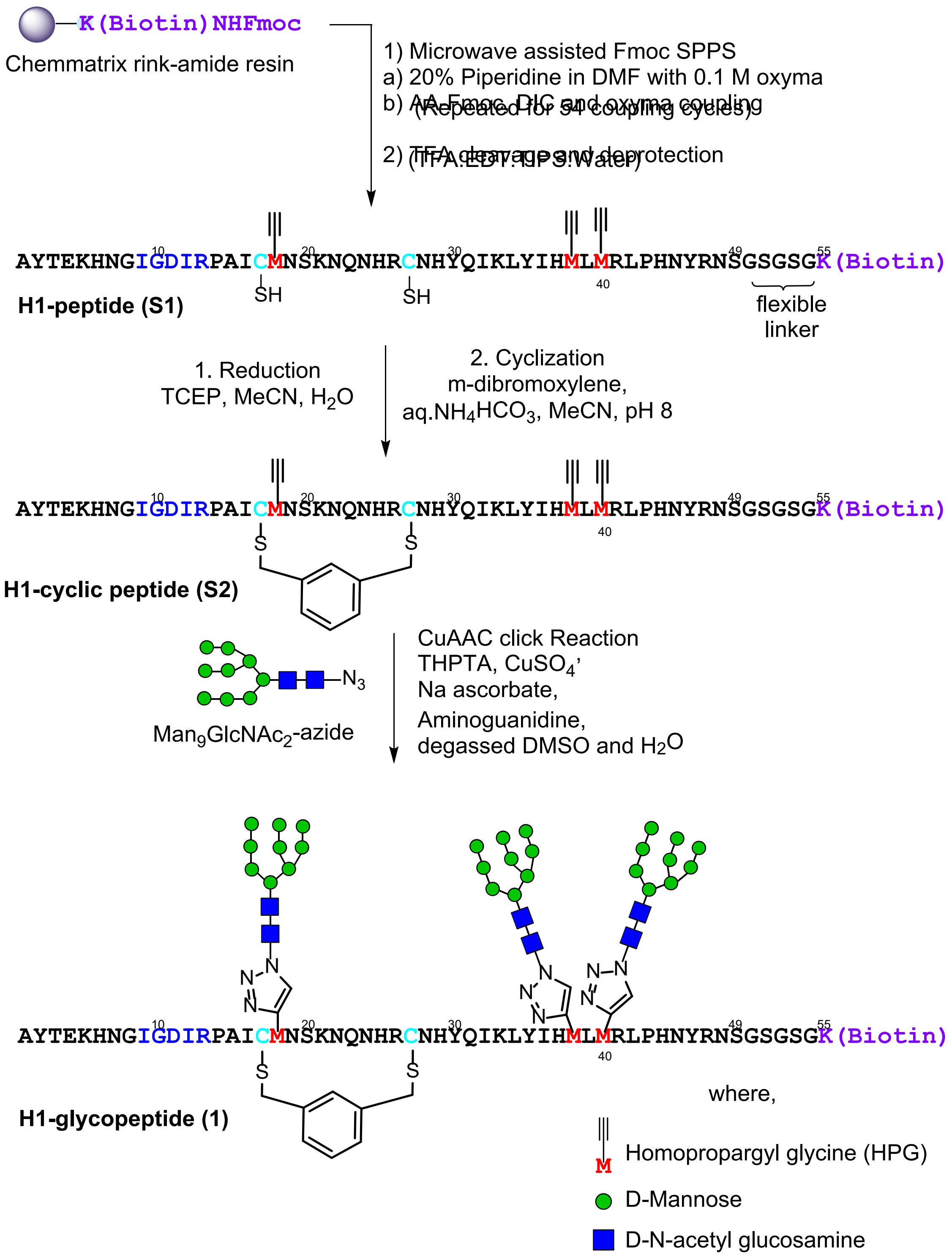

**Scheme S2.** Synthetic scheme to generate **H1-glycopeptide (1)**.

**H1-peptide (S1):** H1-peptide (**S1**) was prepared as in the representative procedure for solid phase peptide synthesis (Section III.A.4), starting with 220 mg of resin loaded at 0.11 mmol/g (24.2 μmol). Of the crude solid peptide, 36 mg was subjected to purification by RP-HPLC on the 10 x 250 mm C4 column (5-30% B over 45 min). 1.6 mg of pure peptide was obtained (4.4 % of the material that was purified). ESI-LRMS observed *m/z* of multiply charged ions 726.78 $[M+9H]^{9+}$, 817.32 $[M+8H]^{8+}$, 934.07 $[M+7H]^{7+}$, 1089.30 $[M+6H]^{6+}$, 1306.86 $[M+5H]^{5+}$; calculated average m/z for H1-peptide (**S1**) $C_{285}H_{439}N_{93}O_{79}S_3$ : 726.38 $[M+9H]^{9+}$, 817.05 $[M+8H]^{8+}$, 933.62 $[M+7H]^{7+}$, 1089.06 $[M+6H]^{6+}$, 1306.67 $[M+5H]^{5+}$.

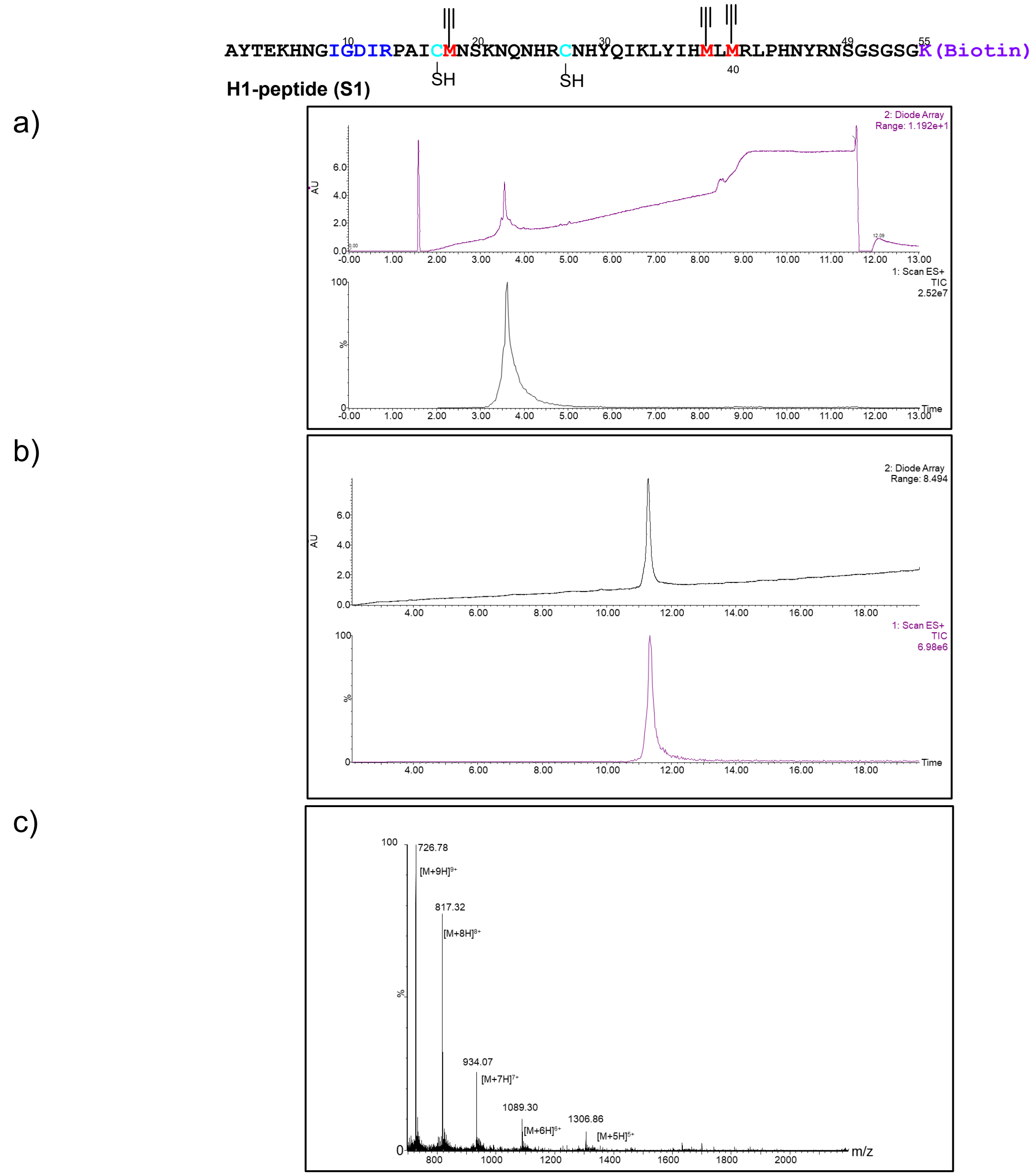

**Figure S9**. LC-MS of **H1-peptide (S1)**. a) UV chromatogram of crude H1-peptide, b) UV and ESI+ TIC chromatogram of purified H1-peptide c) ESI+ MS of purified H1-peptide

(LC-MS method for crude H1-peptide: 1% A to 5% B over 1 minute then 5%A to 45%B gradient over 8 minutes; LC-MS method for purified H1-peptide: 1% A to 5% B over 1 minute then 5%A to 30%B gradient over 18 minutes, solvent A was water/0.07% formic acid and solvent B was acetonitrile/0.07% formic acid, column: Acquity UPLC@Protein BEH C4, 2.1 mm * 150 mm, 1.7 μm, 300 Å; flow rate: 0.25 mL per minute.)

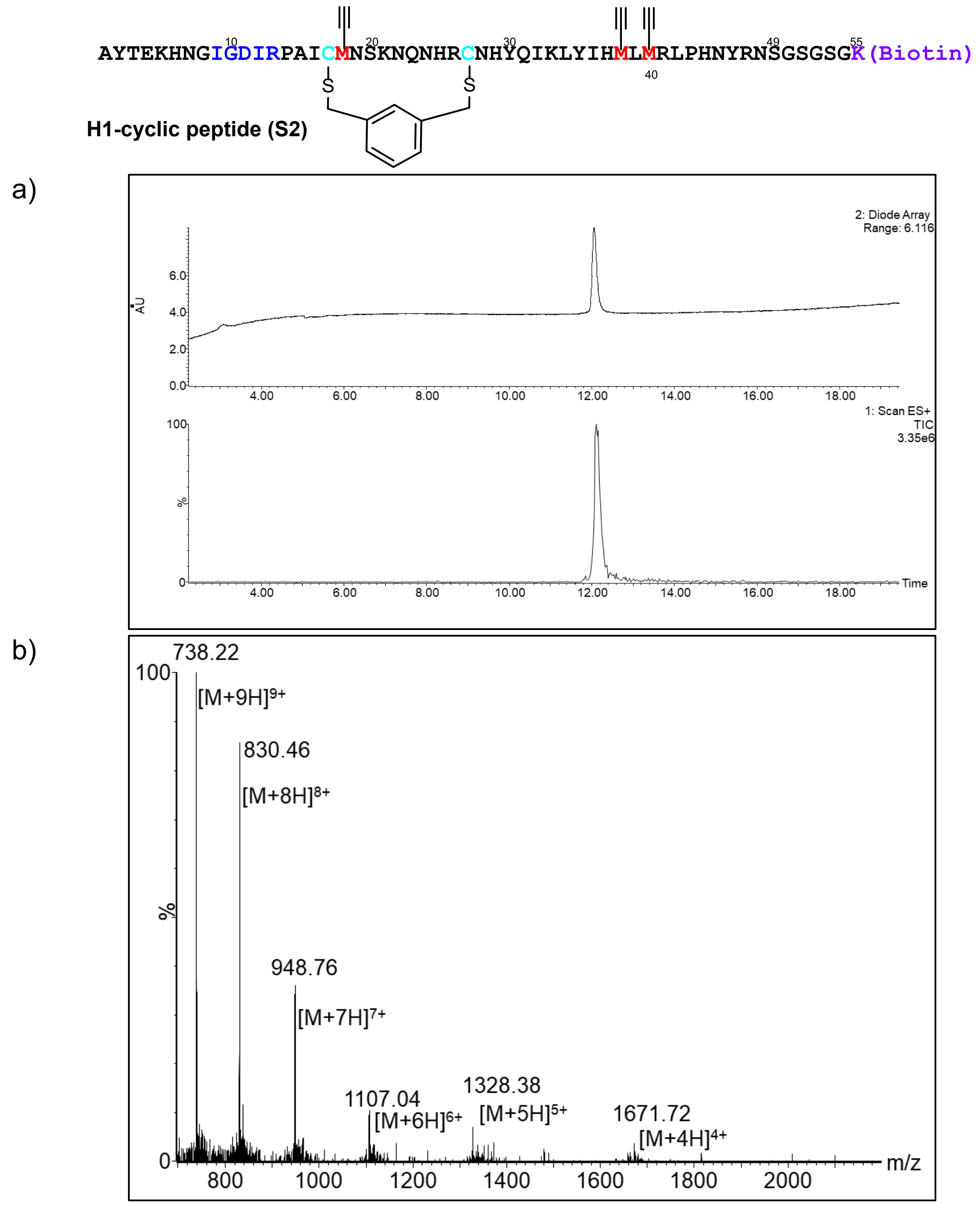

**Figure S10**. LC-MS of **H1-cyclic peptide (S2)**. a) UV and ESI+ TIC chromatogram of H1-peptide, b) ESI+ MS of H1-cyclic peptide.

(LC-MS method: 1% to 5% B over 1 minute then 5% A to 30% B gradient over 18 minutes, solvent A was water/0.07% formic acid and solvent B was acetonitrile/0.07% formic acid, column: Acquity UPLC@Protein BEH C4, 2.1 mm * 150 mm, 1.7 µm, 300 Å; flow rate: 0.25 mL per minute.)

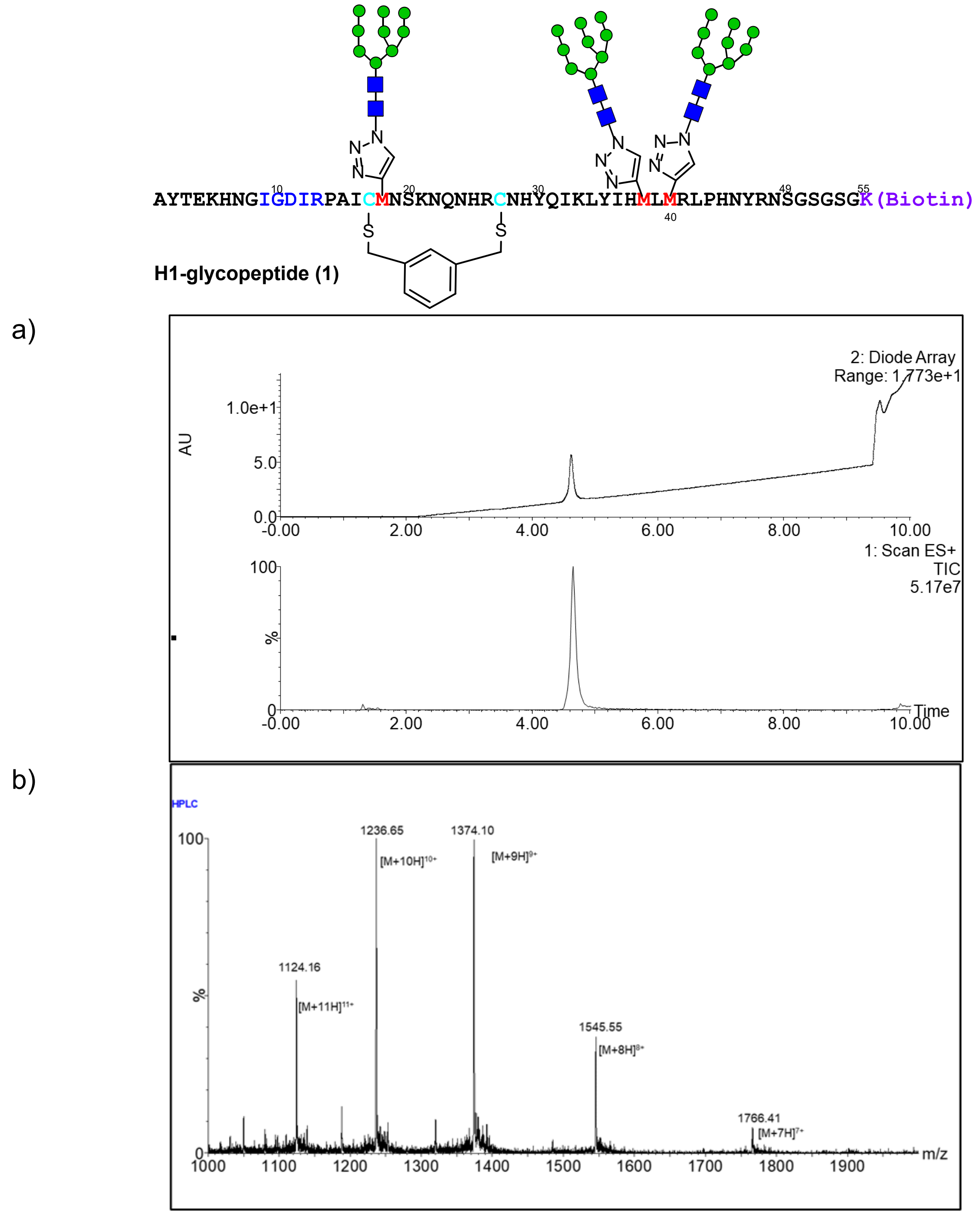

**Figure S11**. LC-MS of **H1-glycopeptide (1)**. a) UV and ESI+ TIC chromatogram of H1-peptide, b) ESI+ MS of H1-glycopeptide.

(LC-MS method: 1% A to 10% B over 1 minute then 10%A to 45%B gradient over 7 minutes, solvent A was water/0.07% formic acid and solvent B was acetonitrile/0.07% formic acid, column: Acquity UPLC@Protein BEH C4, 2.1 mm * 150 mm, 1.7 µm; flow rate: 0.25 mL per minute)

*2. H22-glycopeptide (2)*

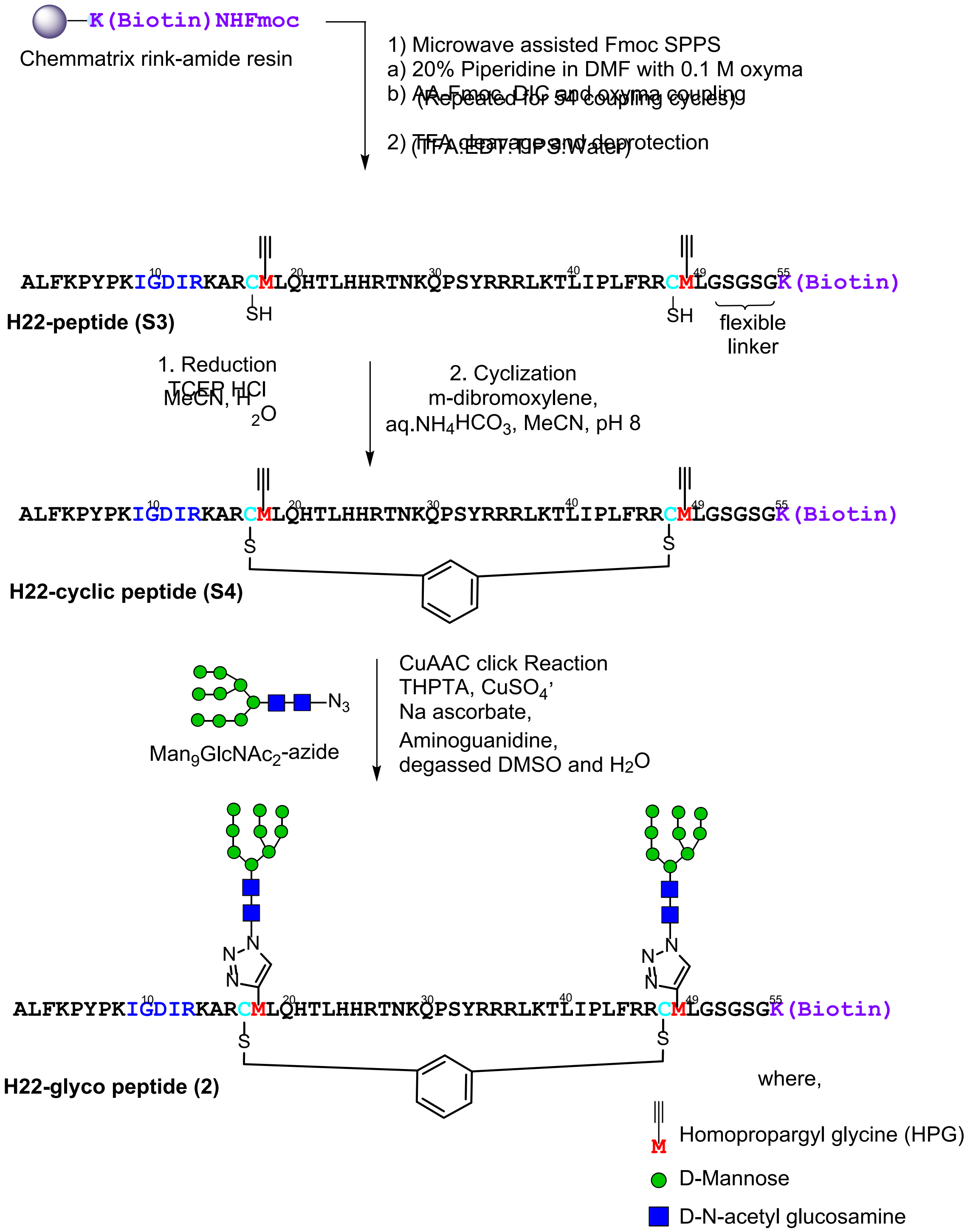

**Scheme S3.** Synthetic scheme to generate **H22-glycopeptide (2).**

**H22-peptide (S3):** H22-peptide (**S3**) was prepared as in the representative procedure for solid phase peptide synthesis (Section III.A.4), starting with 133 mg of resin loaded at 0.3 mmol/g (40 µmol). Of the crude solid peptide, 40 mg was subjected to purification by RP-HPLC on the 10 x 250 mm C4 column (5-30% B over 45 min). 1.07 mg of pure peptide was obtained (2.6 % of the material that was purified). ESI+ MS observed *m/z* of multiply charged ions 1111.41 $[M+6H]^{6+}$, 1333.07 $[M+5H]^{5+}$, 1665.89 $[M+4H]^{4+}$; calculated average m/z for H22-peptide (**S3**) $C_{299}H_{489}N_{97}O_{70}S_3$ : 1110.82 $[M+6H]^{6+}$, 1332.79 $[M+5H]^{5+}$, 1665.73 $[M+4H]^{4+}$.

**H22-cyclic peptide (S4):** The representative procedure for cyclization (Section III.A.7) was followed, using H22-peptide (**S3**) (1.06 mg, 0.16µmol). The crude was purified on the 10 x 250 mm C4 column (5-35 % B over 45 minutes), to afford 0.22 mg (0.03 µmol) of H22-cyclic peptide (**S4**) (20% yield) (quantified by UV-nanodrop). ESI-LRMS observed *m/z* of multiply charged ions 1127.38 $[M+6H]^{6+}$, 1352.99 $[M+5H]^{5+}$, 1690.85 $[M+4H]^{4+}$; calculated average m/z for H22-cyclic peptide (**S4**) $C_{307}H_{495}N_{97}O_{70}S_3$ : 1127.85 $[M+6H]^{6+}$, 1353.21 $[M+5H]^{5+}$, 1691.26 $[M+4H]^{4+}$.

**H22-glycopeptide (2):** The representative procedure for the click glycosylation of synthetic peptides (Section III.A.8) was conducted with H22-cyclic peptide (0.27 mg, 0.04 µmol) and $Man_9GlcNAc_2$-azide (1.6 µL from 50mM solution in water, 0.15 mg, 0.08 µmol). After HPLC purification on the 4.6 x 250 mm C4 column (2-30% B over 45 minutes), 18 µg (0.0017 µmol) of pure H22-glycopeptide (**2**) (4 %) yield was obtained (quantified by UV-nanodrop). ESI-LRMS observed *m/z* of multiply charged ions 1176.37 $[M+9H]^{9+}$, 1323.38 $[M+8H]^{8+}$, 1512.20 $[M+7H]^{7+}$, 1764.33 $[M+6H]^{6+}$; calculated average m/z for H22-glycopeptide (**2**) $C_{447}H_{729}N_{107}O_{180}S_3$ : 1176.38 $[M+9H]^{9+}$, 1323.31 $[M+8H]^{8+}$, 1512.21 $[M+7H]^{7+}$, 1764.07 $[M+6H]^{6+}$.

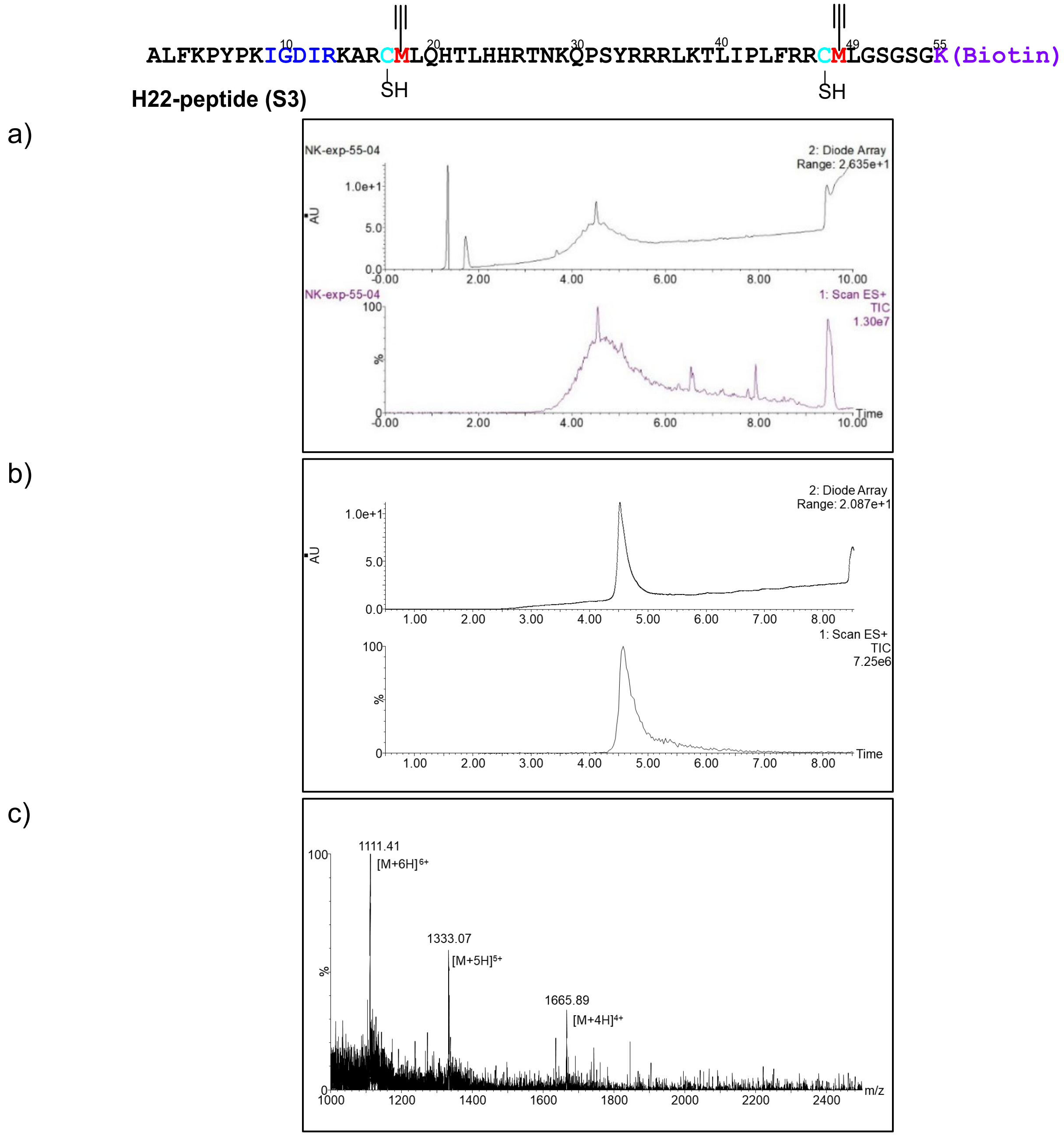

**Figure S12**. LC-MS of **H22 peptide (S3)**. a) UV and ESI+ TIC chromatogram of H22-peptide (crude), b) UV and ESI+ TIC chromatogram of H22-peptide, c) ESI+ MS of H22-peptide

(LC-MS method for crude H22-peptide: 1% A to 5% B over 1 minute then 5%A to 45%B gradient over 8 minutes; LC-MS method for purified H22 peptide: 1% A to 10% B over 1 minute then 10%A to 45%B gradient over 8 minutes, solvent A was water/0.07% formic acid and solvent B was acetonitrile/0.07% formic acid, column: Acquity UPLC@Protein BEH C4, 2.1 mm * 150 mm, 1.7 µm; flow rate: 0.25 mL per minute)

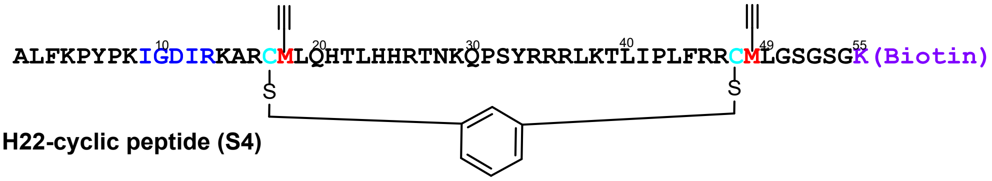

**H22-cyclic peptide (S4)**

a)

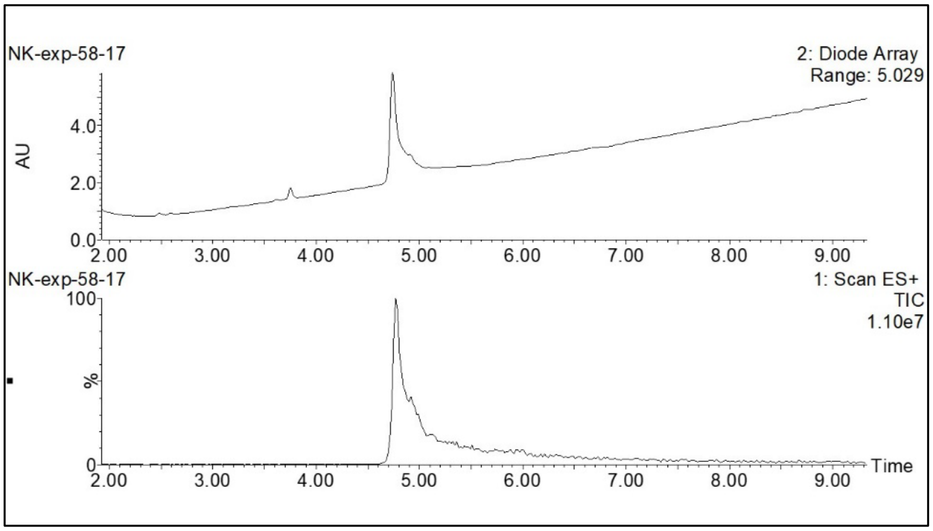

b)

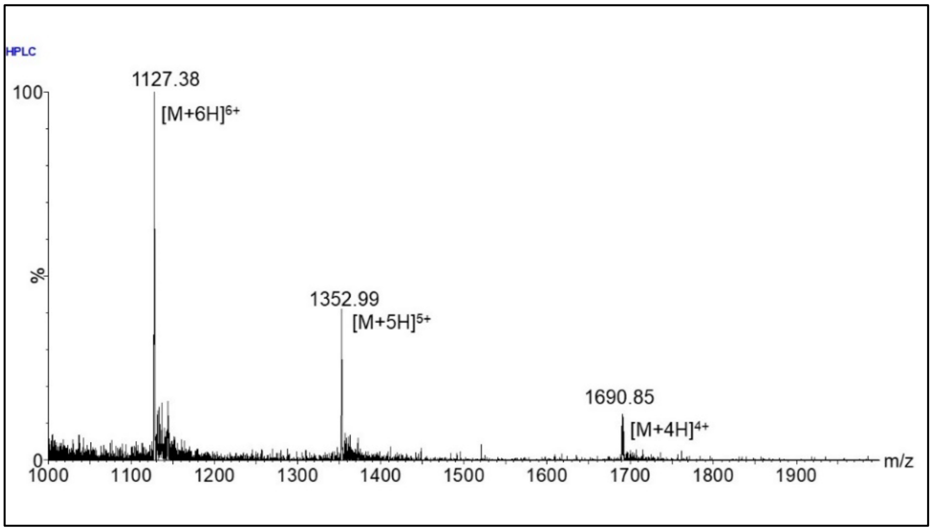

**Figure S13**. LC-MS of **H22 cyclic-peptide (S4)**. a) UV and ESI+ TIC chromatogram of H22-cyclic peptide, b) ESI+ MS of H22-cyclic peptide.
(LC-MS method: 1% A to 10% B over 1 minute then 10%A to 45%B gradient over 9 minutes, solvent A was water/0.07% formic acid, and solvent B was acetonitrile/0.07% formic acid, column: Acquity UPLC@Protein BEH C4, 2.1 mm * 150 mm, 1.7 µm; flow rate: 0.25 mL per minute.)

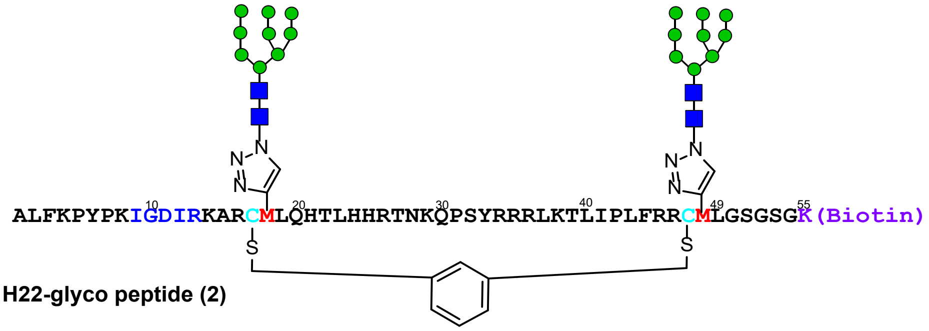

a)

b)

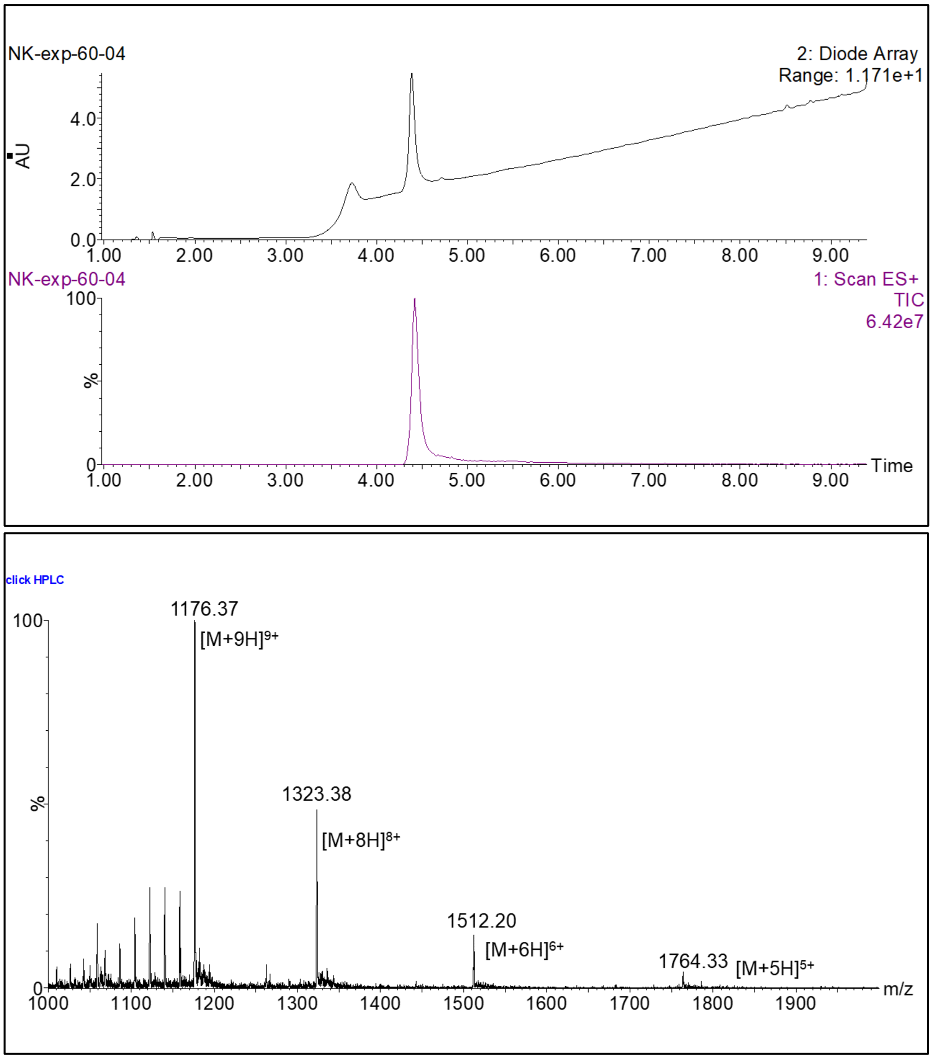

**Figure S14**. LC-MS of **H22 glycopeptide (2)**. a) UV and ESI+ TIC chromatogram of H22-glycopeptide, b) ESI+ MS of H22-glycopeptide.
(LC-MS method: 1% A to 10% B over 1 minute then 10%A to 45%B gradient over 9 minutes, solvent A was water/0.07% formic acid, and solvent B was acetonitrile/0.07% formic acid, column: Acquity UPLC@Protein BEH C4, 2.1 mm * 150 mm, 1.7 µm; flow rate: 0.25 mL per minute.)

3. *H1-QMM-glycopeptide (**3**)*

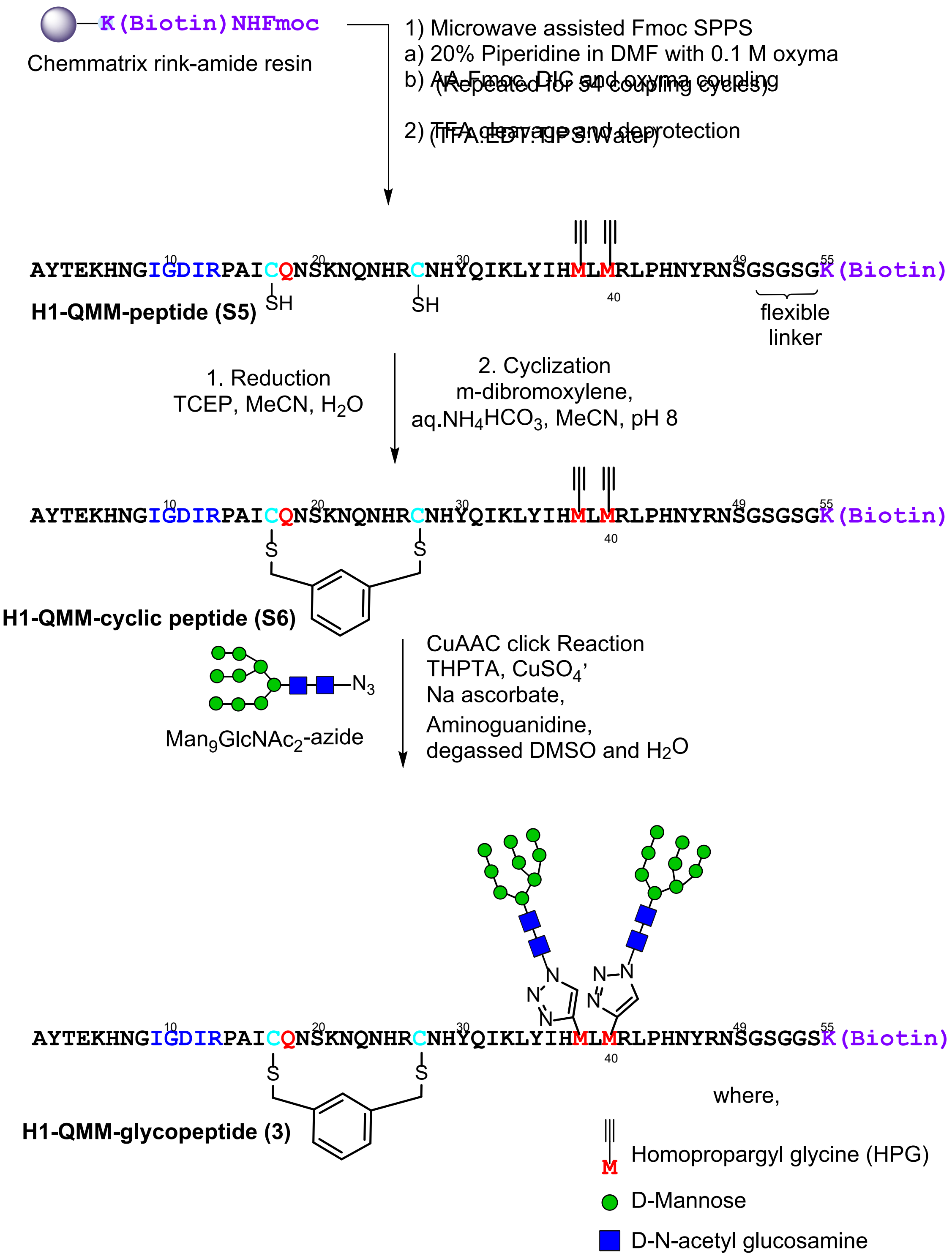

**Scheme S4.** Synthetic scheme to generate **H1-QMM-glycopeptide (3)**.

**H1-QMM-peptide (S5):** HQMM-peptide (**S5**) was prepared as in the representative procedure for solid phase peptide synthesis (Section III.A.4), starting with 180 mg of resin loaded at 0.17 mmol/g (30.6 µmol). Of the crude solid peptide, 60 mg was subjected to purification by RP-HPLC on the 10 x 250 mm C4 column (5-30% B over 45 min). 2.7 mg of pure peptide was obtained (4.5 % of the material that was purified). ESI-LRMS observed *m/z* of multiply charged ions 728.52 $[M+9H]^{9+}$, 819.33 $[M+8H]^{8+}$, 936.23 $[M+7H]^{7+}$, 1092.00 $[M+6H]^{6+}$, 1310.32 $[M+5H]^{5+}$; calculated average m/z values for H1-QMM peptide (**S5**) $C_{284}H_{440}N_{94}O_{80}S_3$ : 728.49 $[M+9H]^{9+}$, 819.42 $[M+8H]^{8+}$, 936.34 $[M+7H]^{7+}$, 1092.23 $[M+6H]^{6+}$, 1310.47 $[M+5H]^{5+}$.

**H1-QMM-cyclic peptide (S6):** The representative procedure for cyclization (Section III.A.7) was followed, using H1-QMM peptide **(S5)** (1.1 mg, 0.18 µmol). The crude was purified on the 10 x 250 mm C4 column (5-35 % B over 45 minutes), to afford 0.69 mg (0.10 µmol) of H1-MQQ-cyclic peptide (**S6**) (57% yield) (quantified by UV-nanodrop). ESI-LRMS observed *m/z* of multiply charged ions 739.96 $[M+9H]^{9+}$, 832.21 $[M+8H]^{8+}$, 950.98 $[M+7H]^{7+}$, 1109.23 $[M+6H]^{6+}$, 1330.87 $[M+5H]^{5+}$ ; calculated average m/z values for H1-QMM-cyclic peptide (**S6**) $C_{292}H_{446}N_{94}O_{80}S_3$ : 739.84 $[M+9H]^{9+}$, 832.19 $[M+8H]^{8+}$, 950.93 $[M+7H]^{7+}$, 1109.25 $[M+6H]^{6+}$, 1330.90 $[M+5H]^{5+}$.

**H1-QMM-glycopeptide (3):** The representative procedure for the click glycosylation of synthetic peptides (Section III.A.8) was conducted with H1-QMM-cyclic peptide **(S6)** (0.66 mg, 0.10 µmol) and $Man_9GlcNAc_2$-azide (8 µl from 50 mM solution in water, 0.76 mg, 0.40 µmol). After HPLC purification on the 4.6 x 250 mm C4 column (2-30% B over 45 minutes), 31 µg (0.0031 µmol) of pure H1-QMM-glycopeptide (**3**) (3 %) yield was obtained (quantified by UV-nanodrop). ESI-LRMS observed *m/z* of multiply charged ions 952.41 $[M+11H]^{11+}$, 1047.59 $[M+10H]^{10+}$, 1163.84 $[M+9H]^{9+}$, 1309.35 $[M+8H]^{8+}$, 1496.33 $[M+7H]^{7+}$; 1745.26 $[M+6H]^{6+}$; calculated average m/z values for H1-MQQ-glycopeptide **(3)** $C_{432}H_{680}N_{104}O_{190}S_3$ : 952.54 $[M+11H]^{11+}$, 1047.69 $[M+10H]^{10+}$, 1163.99 $[M+9H]^{9+}$, 1309.36 $[M+8H]^{8+}$, 1496.27 $[M+7H]^{7+}$; 1745.48 $[M+6H]^{6+}$.

a)

b)

c)

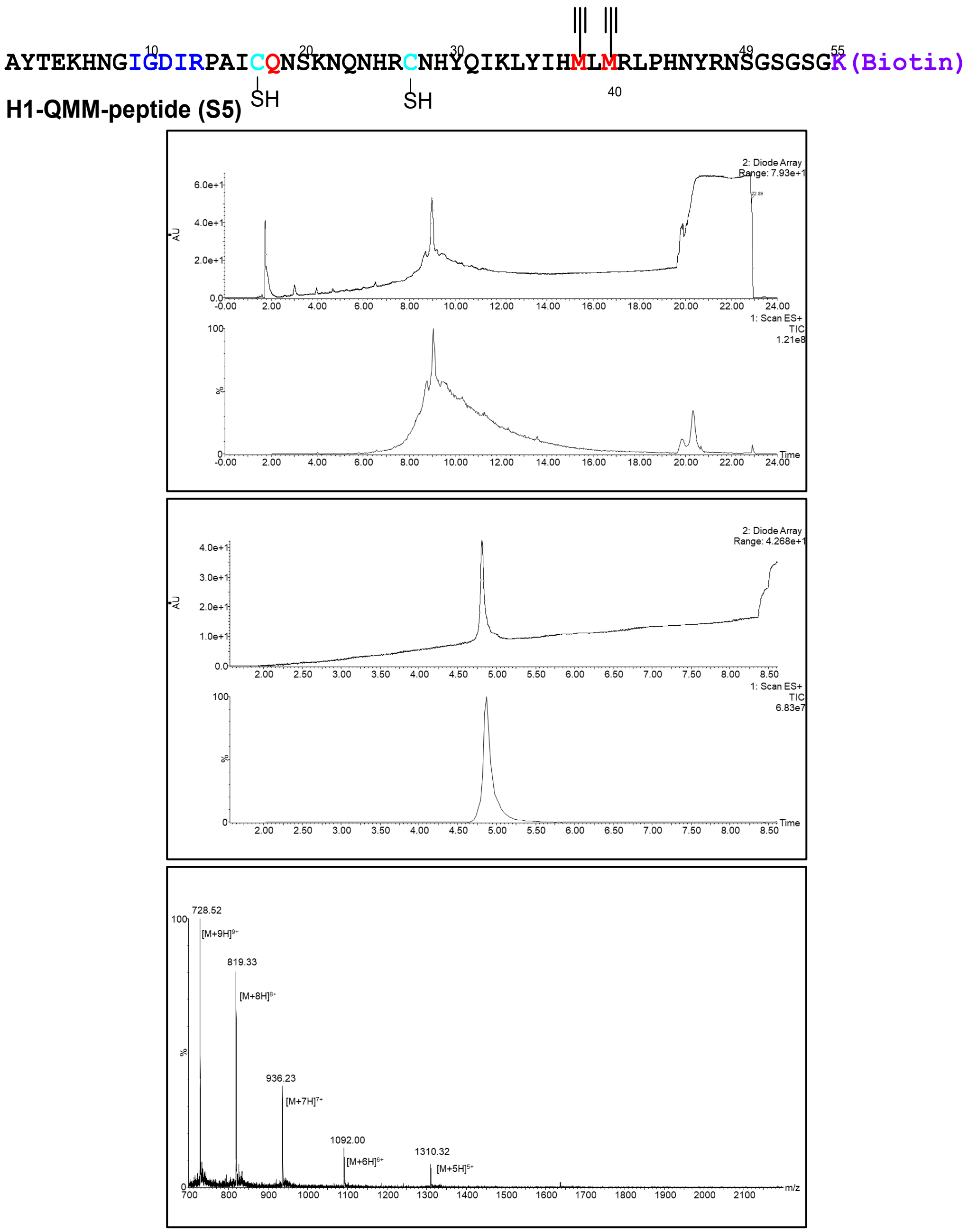

**Figure S15**. LC-MS of **H1-QMM-peptide (S5)**. a) UV- TIC chromatogram of H1-QMM-peptide (crude), b) UV and ESI+ TIC chromatogram of H-QMM-peptide (purified), c) ESI+ MS of H1-QMM-peptide.

(LC-MS method for crude H1-QMM-peptide: 1% A to 5% B over 1 minute then 5% A to 45% B gradient over 18 minutes, LC-MS method for purified H1-QMM-peptide: 1% A to 5% B over 1 minute then 5% A to 35% B gradient over 7 minutes, solvent A was water/0.07% formic acid and solvent B was acetonitrile/0.07% formic acid, column: Acquity UPLC@Protein BEH C4, 2.1 mm * 150 mm, 1.7 µm; flow rate: 0.25 mL per minute.)

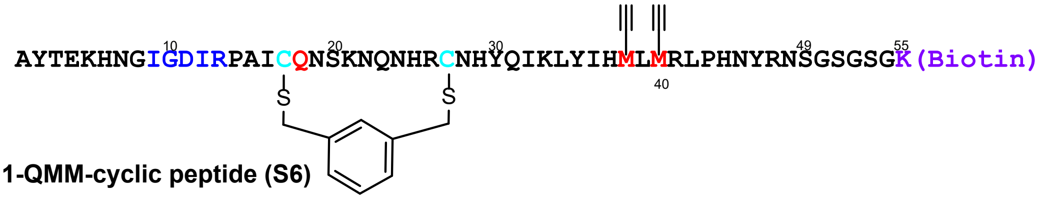

**H1-QMM-cyclic peptide (S6)**

a)

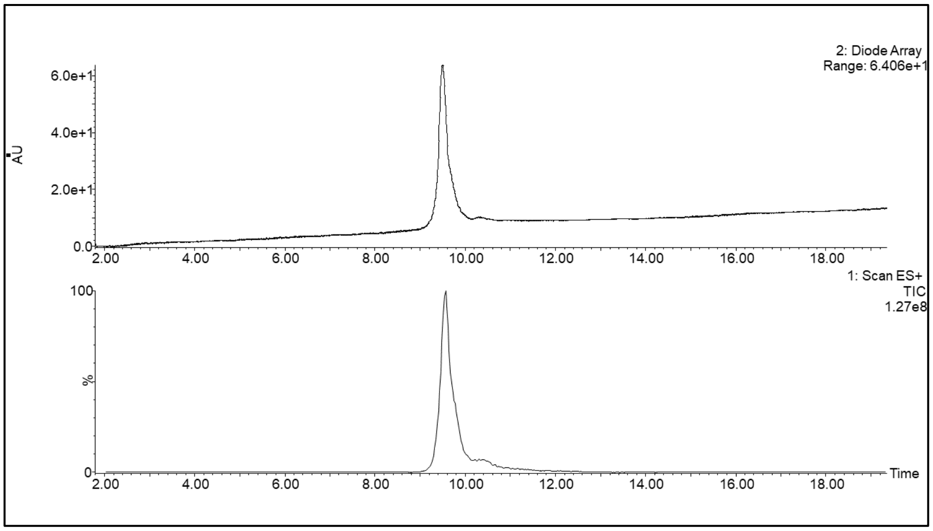

b)

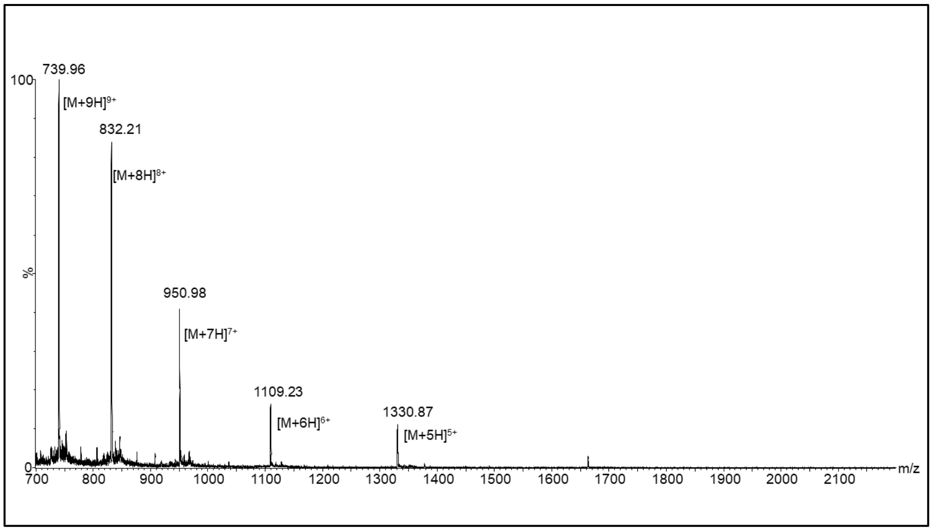

**Figure S16**. LC-MS of **H1-QMM-cyclic-peptide (S6)**. a) UV and ESI+ TIC chromatogram of H1-QMM-peptide, b) ESI+ MS of H1-QMM-cyclic-peptide.

(LC-MS method: 1% A to 5% B over 1 minute then 5% A to 35% B gradient over 18 minutes, solvent A was water/0.07% formic acid, and solvent B was acetonitrile/0.07% formic acid, column: Acquity UPLC@Protein BEH C4, 2.1 mm * 150 mm, 1.7 µm; flow rate: 0.25 mL per minute.)

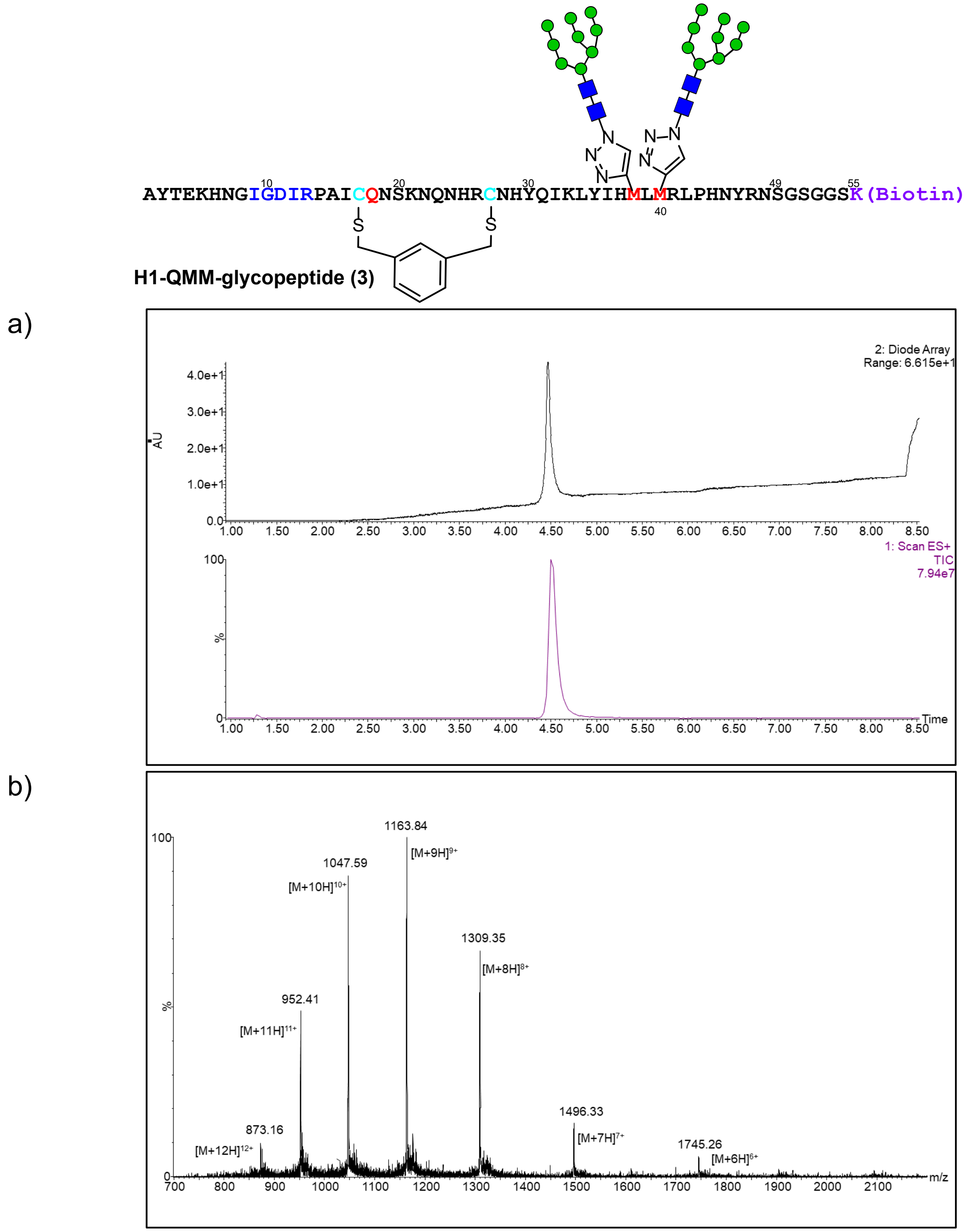

**Figure S17**. LC-MS of **H1-QMM-glycopeptide (3)**. a) UV and ESI+ TIC chromatogram of H1-QMM-peptide, b) ESI+ MS of H1-QMM-glycopeptide.

(LC-MS method: 1% A to 10% B over 1 minute then 10%A to 45%B gradient over 7 minutes, solvent A was water/0.07% formic acid and solvent B was acetonitrile/0.07% formic acid, column: Acquity UPLC@Protein BEH C4, 2.1 mm * 150 mm, 1.7 µm; flow rate: 0.25 mL per minute)

*4. H1-MQQ-glycopeptide (4)*

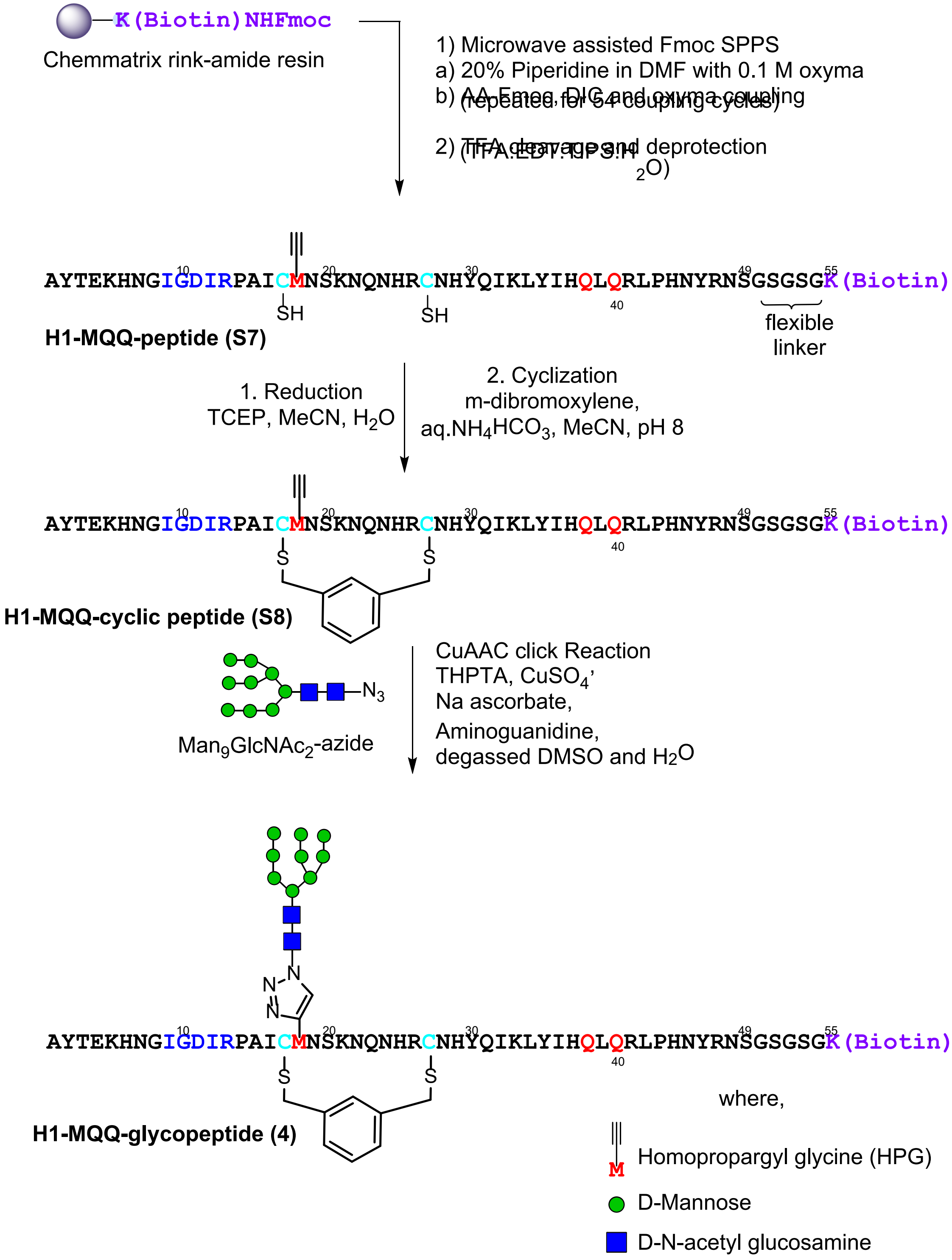

**Scheme S5.** Synthetic scheme to generate **H1-MQQ-glycopeptide (4)**.

**H1-MQQ-peptide (S7):** H1-MQQ-peptide **(S7)** was prepared as in the representative procedure for solid phase peptide synthesis (Section III.A.4), starting with 180mg of resin loaded at 0.17 mmol/g (30.6 µmol). Of the crude solid peptide, 60 mg was subjected to purification by RP-HPLC on the 10 x 250 mm C4 column (5-30% B over 45 min). 1.9 mg of pure peptide was obtained (3.1 % of the material that was purified). ESI-LRMS observed *m/z* of multiply charged ions 730.60 $[M+9H]^{9+}$, 821.74 $[M+8H]^{8+}$, 938.96 $[M+7H]^{7+}$, 1095.32 $[M+6H]^{6+}$, 1314.16 $[M+5H]^{5+}$; calculated average m/z values for H1-MQQ-peptide **(S7)** $C_{283}H_{441}N_{95}O_{81}S_3$ : 730.60 $[M+9H]^{9+}$, 821.80 $[M+8H]^{8+}$, 939.05 $[M+7H]^{7+}$, 1095.39 $[M+6H]^{6+}$, 1314.27 $[M+5H]^{5+}$.

**H1-MQQ-cyclic peptide (S8):** The representative procedure for cyclization (Section III.A.7) was followed, using H1-MQQ peptide **(S7)** (1.3 mg, 0.20 µmol). The crude was purified on the 10 x 250 mm C4 column (5-35 % B over 45 minutes), to afford 0.72 mg (0.10 µmol) of H1-MQQ-cyclic peptide (**S8**) (46% yield) (quantified by UV-nanodrop). ESI-LRMS observed *m/z* of multiply charged ions 741.91 $[M+9H]^{9+}$, 834.55 $[M+8H]^{8+}$, 953.52 $[M+7H]^{7+}$, 1112.28 $[M+6H]^{6+}$, 1334.77 $[M+5H]^{5+}$, 1667.90 $[M+4H]^{4+}$; calculated average m/z values for H1-MQQ-cyclic peptide **(S8)** $C_{291}H_{447}N_{95}O_{81}S_3$ : 741.95 $[M+9H]^{9+}$, 834.56 $[M+8H]^{8+}$, 953.64 $[M+7H]^{7+}$, 1112.42 $[M+6H]^{6+}$, 1334.70 $[M+5H]^{5+}$, 1668.12 $[M+4H]^{4+}$.

**H1-MQQ-glycopeptide (4):** The representative procedure for the click glycosylation of synthetic peptides (Section III.A.8) was conducted with H1-MQQ-cyclic peptide **(S8)** (0.71 mg, 0.11 µmol) and $Man_9GlcNAc_2$-azide (4.4 µl from 50 mM stock solution in water, 0.42 mg, 0.22 µmol). After HPLC purification on the 4.6 x 250 mm C4 column (2-30% B over 45 minutes), 81 µg (0.0094 µmol) of pure H1-MQQ-glycopeptide (**4**) (8 %) yield was obtained (quantified by UV-nanodrop). ESI-LRMS observed *m/z* of multiply charged ions 858.80 $[M+10H]^{10+}$, 954.10 $[M+9H]^{9+}$, 1073.21 $[M+8H]^{8+}$, 1226.45 $[M+7H]^{7+}$ 1430.41 $[M+6H]^{6+}$, 1716.14 $[M+5H]^{5+}$, calculated average m/z ratios for H1-MQQ-glycopeptide **(4)** $C_{361}H_{564}N_{100}O_{136}S_3$ : 858.72 $[M+10H]^{10+}$, 954.02 $[M+9H]^{9+}$, 1073.15 $[M+8H]^{8+}$, 1226.31 $[M+7H]^{7+}$ 1430.53 $[M+6H]^{6+}$, 1716.43 $[M+5H]^{5+}$.

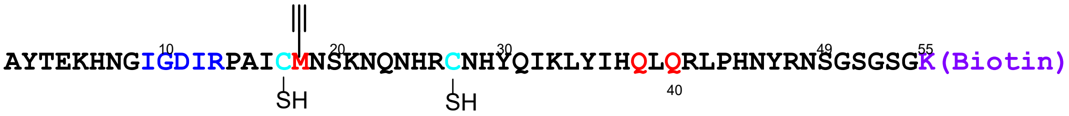

**H1-MQQ-peptide (S7)**

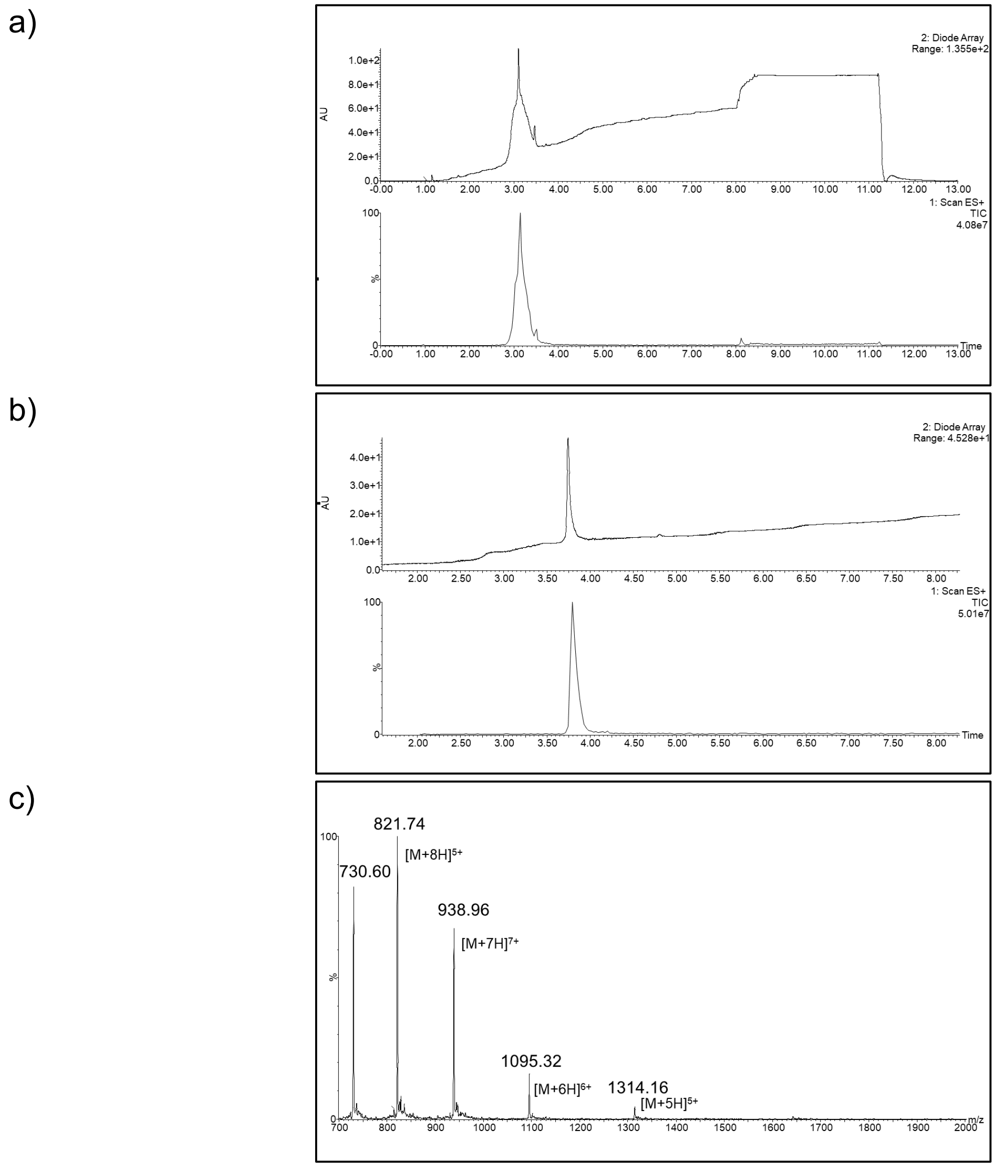

**Figure S18**. LC-MS of **H1-MQQ-peptide (S7)**. a) UV and ESI+ TIC chromatogram of H-MQQ-peptide (crude), b) UV and ESI+ TIC chromatogram of H-MQQ-peptide (purified), c) ESI+ MS of H1-MQQ-peptide (crude).

(LC-MS method for crude H1-MQQ-peptide: 1% A to 10% B over 1 minute then 10%A to 45%B gradient over 7 minutes; LC-MS method for H1-MQQ peptide: 1% A to 10% B over 1 minute then 10%A to 45%B gradient over 7 minutes, solvent A was water/0.07% formic acid and solvent B was acetonitrile/0.07% formic acid, column: Acquity UPLC@Protein BEH C4, 2.1 mm * 150 mm, 1.7 µm; flow rate: 0.25 mL per minute.)

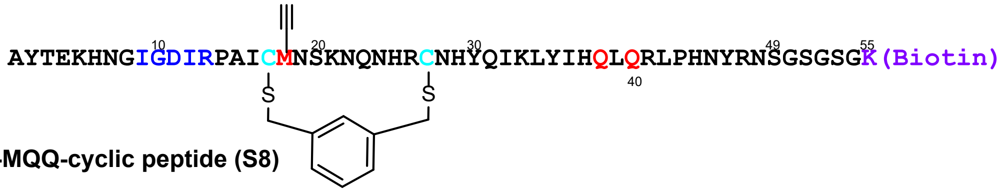

H1-MQQ-cyclic peptide (S8)

a)

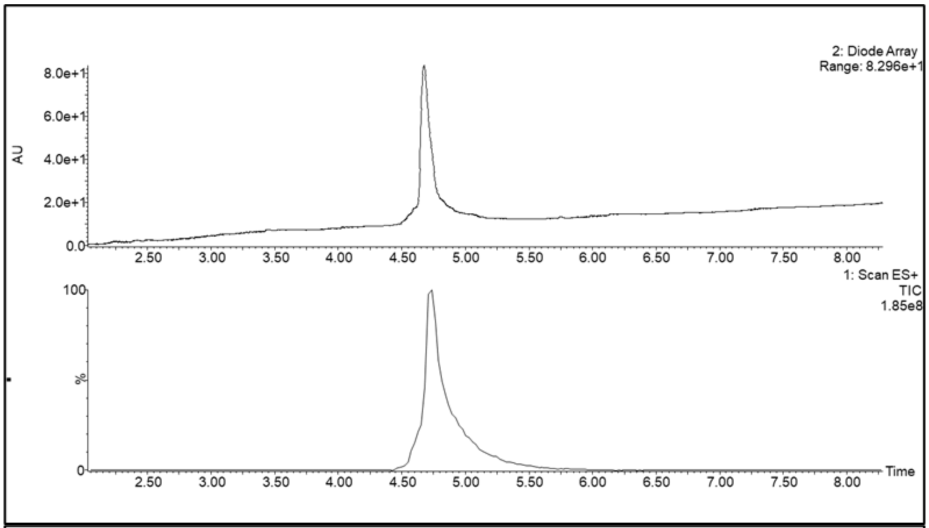

b)

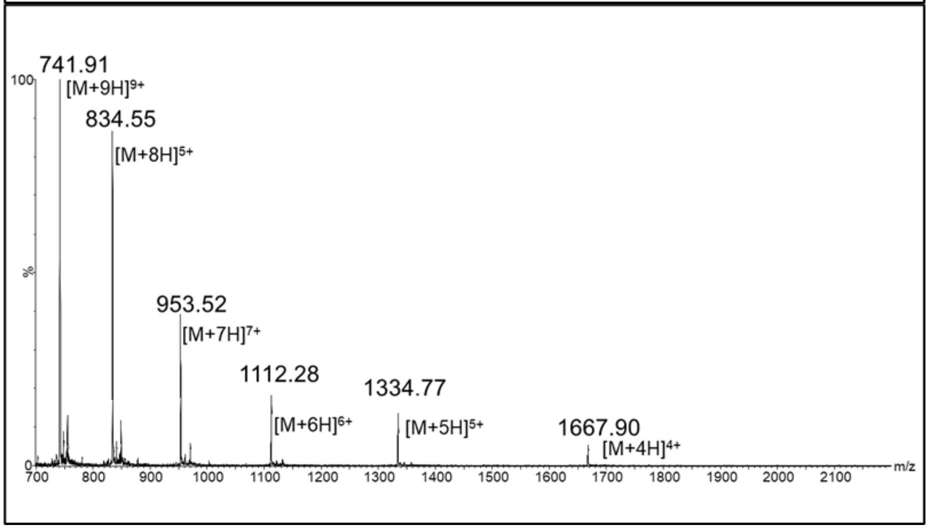

**Figure S19**. LC-MS of **H1-MQQ-cyclic-peptide (S8)**. a) UV and ESI+ TIC chromatogram of H1-MQQ-peptide, b) ESI+ MS of H1-MQQ-cyclic-peptide.

(LC-MS method: 1% A to 10% B over 1 minute then 10%A to 45%B gradient over 7 minutes, solvent A was water/0.07% formic acid and solvent B was acetonitrile/0.07% formic acid, column: Waters Acquity UPLC @Protein BEH C4, 2.1 mm * 150 mm, 1.7 µm; flow rate: 0.25 mL per minute.)

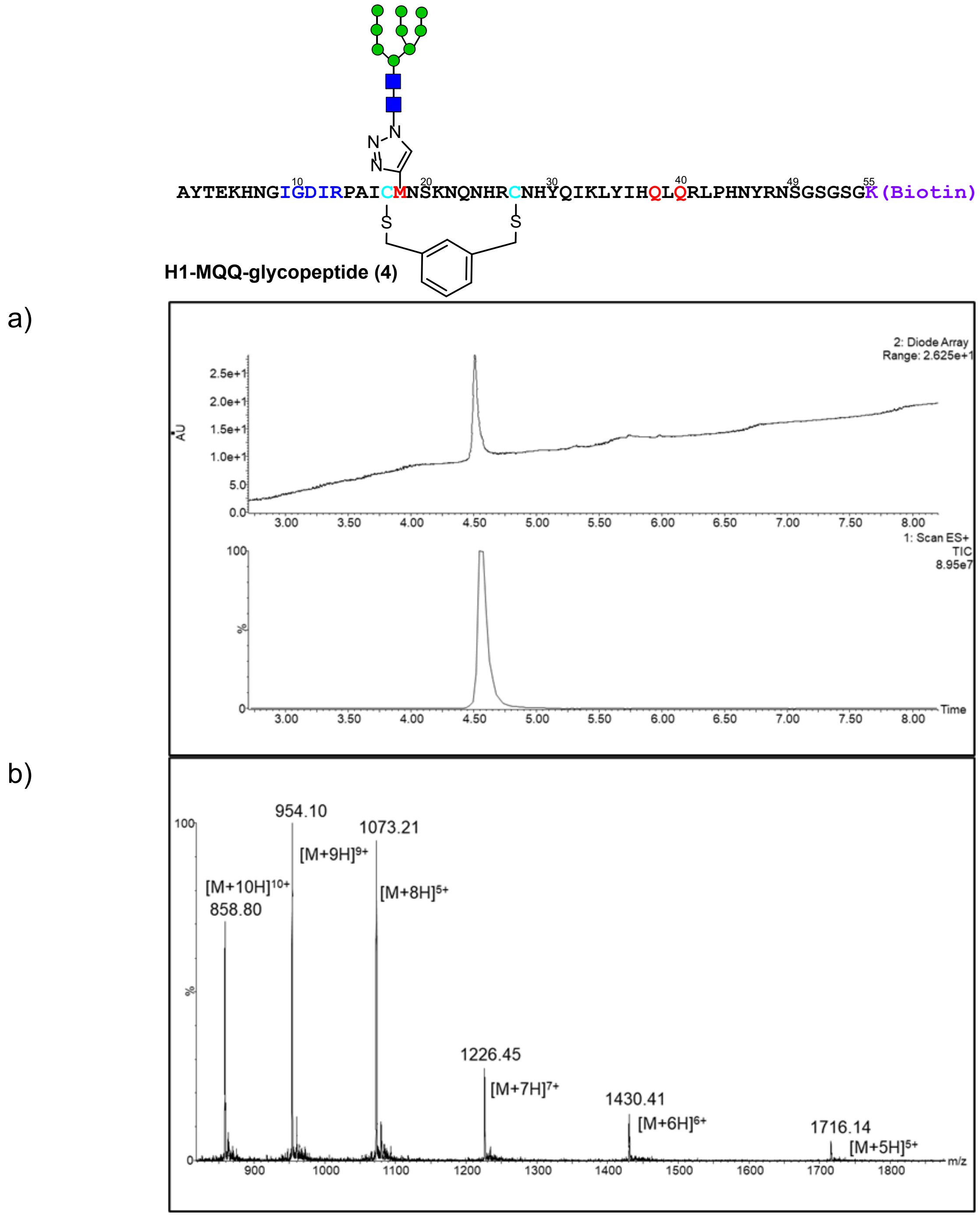

**Figure S20**. LC-MS of **H1-MQQ-glycopeptide (4)**. a) UV and ESI+ TIC chromatogram of H1-MQQ-peptide, b) ESI+ MS of H1-MQQ-glycopeptide.

(LC-MS method: 1% A to 10% B over 1 minute then 10%A to 45%B gradient over 7 minutes, solvent A was water/0.07% formic acid, and solvent B was acetonitrile/0.07% formic acid, column: Acquity UPLC@Protein BEH C4, 2.1 mm * 150 mm, 1.7 µm; flow rate: 0.25 mL per minute)

5. *H1-IAAIA-glycopeptide (5)*

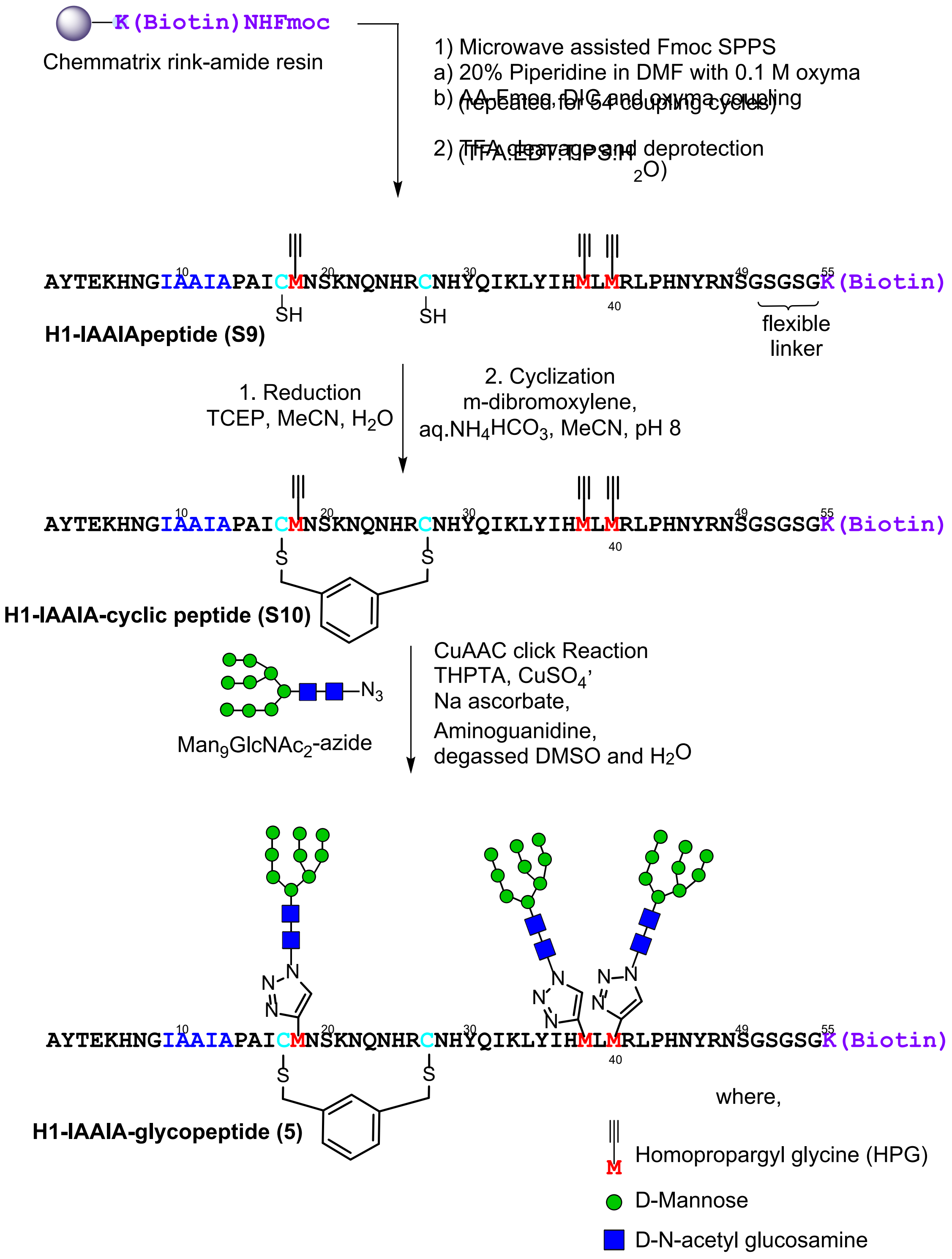

**Scheme S6.** Synthetic scheme to generate **H1-IAAIA-glycopeptide (5)**.

**H1-IAAIA-peptide (S9):** H1-IAAIA-peptide **(S9)** was prepared as in the representative procedure for solid phase peptide synthesis (Section III.A.4), starting with 180mg of resin loaded at 0.17 mmol/g (30.6 µmol). Of the crude solid peptide, 80 mg was subjected to purification by RP-HPLC on the 10 x 250 mm C4 column (5-30% B over 45 min). 2.1 mg of pure peptide was obtained (2.6 % of the material that was purified). ESI-LRMS observed *m/z* of multiply charged ions 1069.51 $[M+6H]^{6+}$, 1283.56 $[M+5H]^{5+}$, 1604.25 $[M+4H]^{4+}$, 2138.48 $[M+3H]^{3+}$; calculated average m/z values for H1-IAAIA-peptide **(S9)** $C_{282}H_{434}N_{90}O_{77}S_3$ : 1069.88 $[M+6H]^{6+}$, 1283.65 $[M+5H]^{5+}$, 1604.31 $[M+4H]^{4+}$, 2138.75 $[M+3H]^{3+}$.

**H1-IAAIA-cyclic peptide (S10):** The representative procedure for cyclization (Section III.A.7) was followed, using H1-IAAIA peptide **(S9)** (2.1 mg, 0.34 µmol). The crude was purified on the 10 x 250 mm C4 column (5-35 % B over 45 minutes), to afford 0.82 mg (0.12 µmol) of H1-IAAIA-cyclic peptide **(S10)** (37% yield) (quantified by UV-nanodrop). ESI-LRMS observed *m/z* of multiply charged ions 1086.69 $[M+6H]^{6+}$, 1304.00 $[M+5H]^{5+}$, 1629.82 $[M+4H]^{4+}$, 2172.51 $[M+3H]^{+3}$; calculated average m/z values for H1-IAAIA-cyclic peptide **(S10)** $C_{290}H_{440}N_{90}O_{77}S_3$ : 1086.90 $[M+6H]^{6+}$, 1304.08 $[M+5H]^{5+}$, 1629.85 $[M+4H]^{4+}$, 2172.79 $[M+3H]^{+3}$.

**H1-IAAIA-glycopeptide (5):** The representative procedure for the click glycosylation of synthetic peptides (Section III.A.8) was conducted with H1-IAAIA-cyclic peptide **(S10)** (0.71 mg, 0.11 µmol) and $Man_9GlcNAc_2$-azide (10 µL from 50 mM stock solution in water, 0.94 mg, 0.49 µmol). After HPLC purification on the 4.6 x 250 mm C4 column (2-45% B over 45 minutes), 55 µg (0.0044 µmol) of pure H1-IAAIA-glycopeptide (**5**) (4 %) yield was obtained (quantified by UV-nanodrop). ESI-LRMS observed *m/z* of multiply charged ions 1224.85 $[M+10H]^{10+}$, 1361.09 $[M+9H]^{9+}$, 1531.30 $[M+8H]^{8+}$, 1749.64 $[M+7H]^{7+}$, 2040.81 $[M+6H]^{6+}$ ; calculated average m/z values for H1-IAAIA-glycopeptide **(5)** $C_{500}H_{791}N_{105}O_{242}S_3$ : 1225.15 $[M+10H]^{10+}$, 1361.16 $[M+9H]^{9+}$, 1531.18 $[M+8H]^{8+}$, 1749.78 $[M+7H]^{7+}$, 2041.24 $[M+6H]^{6+}$.

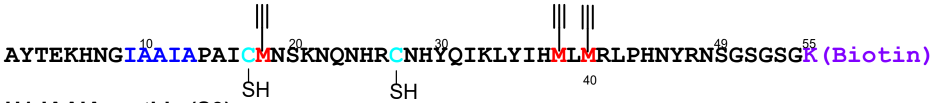

**H1-IAAIApeptide (S9)**

a)

b)

c)

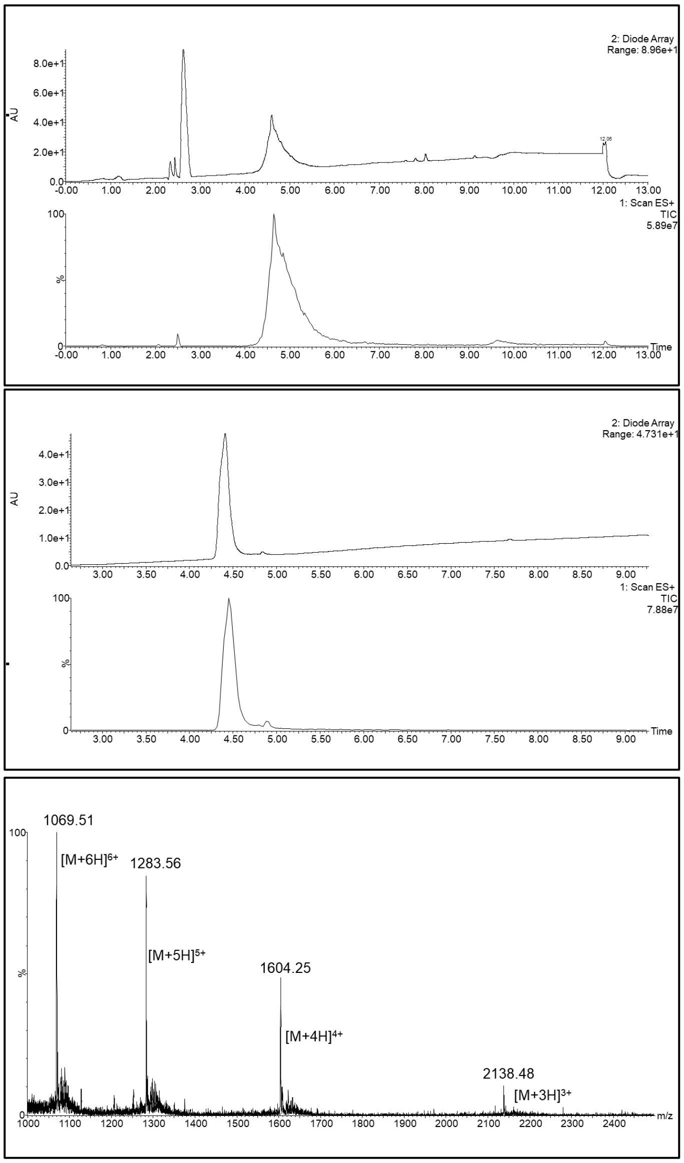

**Figure S21**. LC-MS of **H1-IAAIA-peptide (S9)**. a) UV and ESI+ TIC chromatogram of H1-IAAIA-peptide (crude), b) UV and ESI+ TIC chromatogram of H1-IAAIA-peptide (purified), c) ESI+ MS of H1-IAAIA-peptide.

(LC-MS method for crude HIAAIA-peptide: 1% A to 5% B over 1 minute then 10%A to 45%B gradient over 8 minutes LC-MS method for pure H1-IAAIA peptide: 1% A to 10% B over 1 minute then 10%A to 45%B gradient over 8 minutes, solvent A was water/0.07% formic acid and solvent B was acetonitrile/0.07% formic acid, column: Acquity UPLC@Protein BEH C4, 2.1 mm * 150 mm, 1.7 μm; flow rate: 0.3 mL per minute)

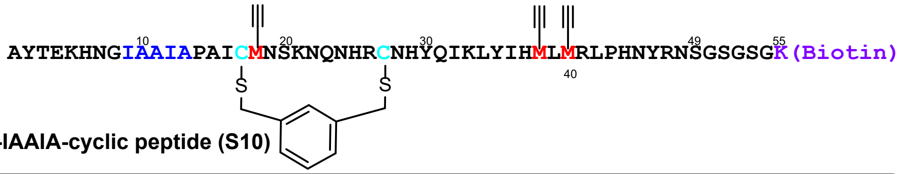

H1-IAAIA-cyclic peptide (S10)

a)

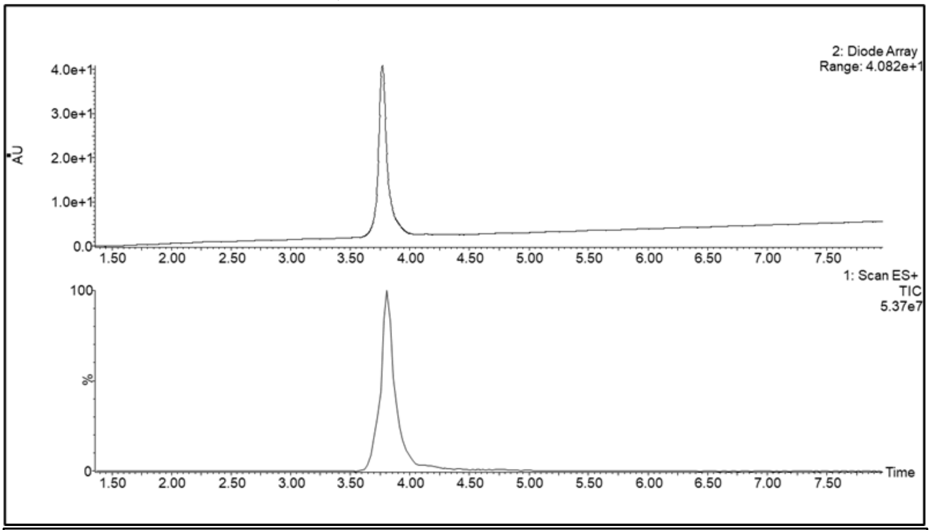

b)

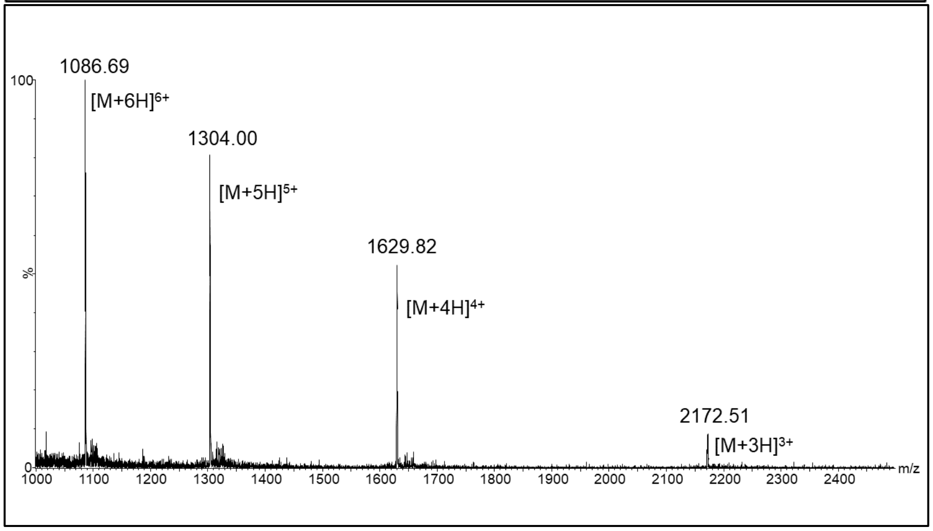

**Figure S22**. LC-MS of **H1-IAAIA-cyclic-peptide (S10)**. a) UV and ESI+ TIC chromatogram of H1-IAAIA-peptide, b) ESI+ MS of H1-IAAIA-cyclic-peptide.

(LC-MS method: 1% A to 10% B over 1 minute then 10%A to 45%B gradient over 7 minutes, solvent A was water/0.07% formic acid, and solvent B was acetonitrile/0.07% formic acid, column: Acquity UPLC@Protein BEH C4, 2.1 mm * 150 mm, 1.7 µm; flow rate: 0.25 mL per minute.)

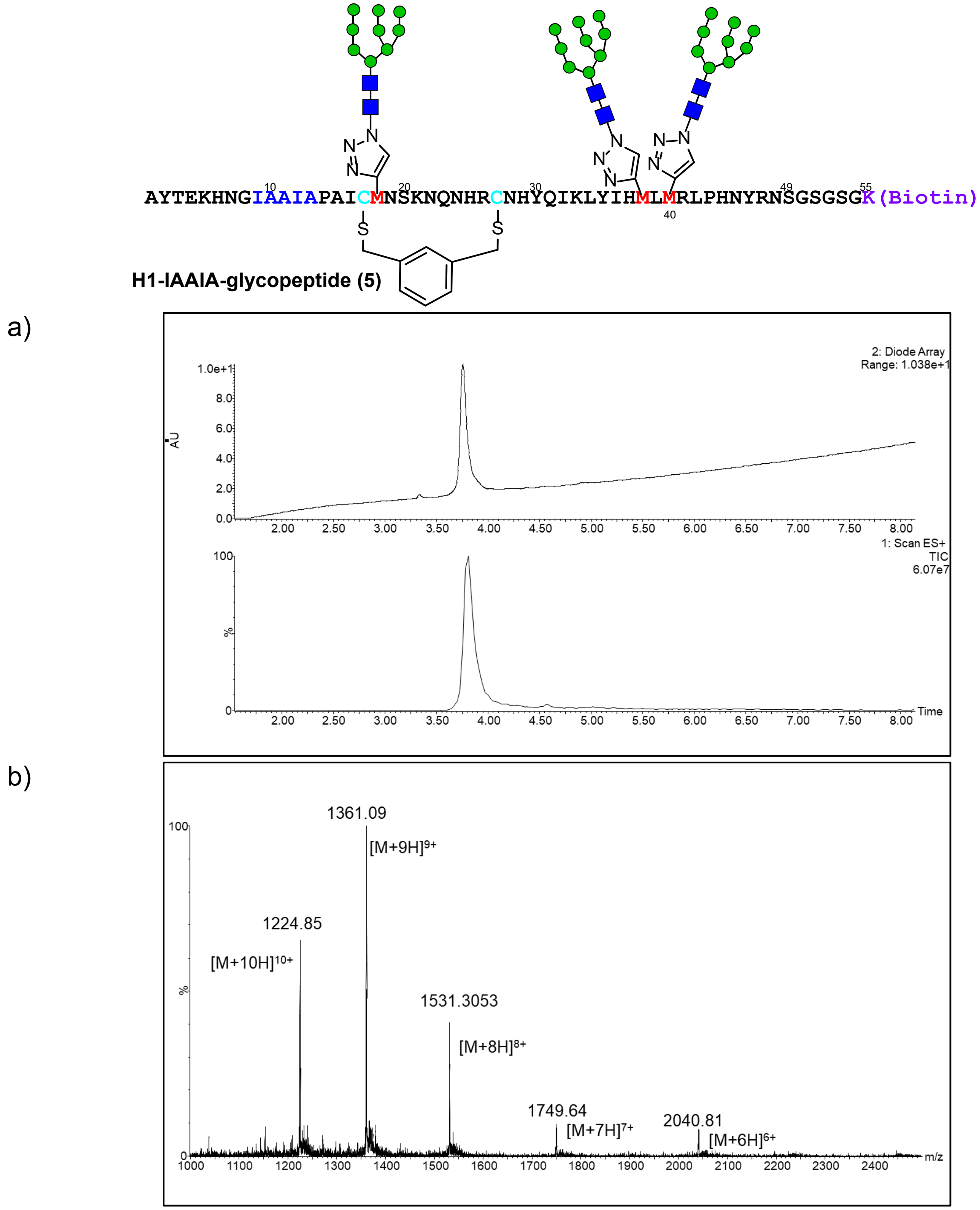

**Figure S23**. LC-MS of **H1-IAAIA-glycopeptide (5)**. a) UV and ESI+ TIC chromatogram of H1-peptide, b) ESI+ MS of H1-IAAIA-glycopeptide.

(LC-MS method: 1% A to 10% B over 1 minute then 10%A to 45%B gradient over 7 minutes, solvent A was water/0.07% formic acid, and solvent B was acetonitrile/0.07% formic acid, column: Acquity UPLC@Protein BEH C4, 2.1 mm * 150 mm, 1.7 µm; flow rate: 0.25 mL per minute)

*6. H1-QQQ-non-glycosylated Peptide (**S12**)*

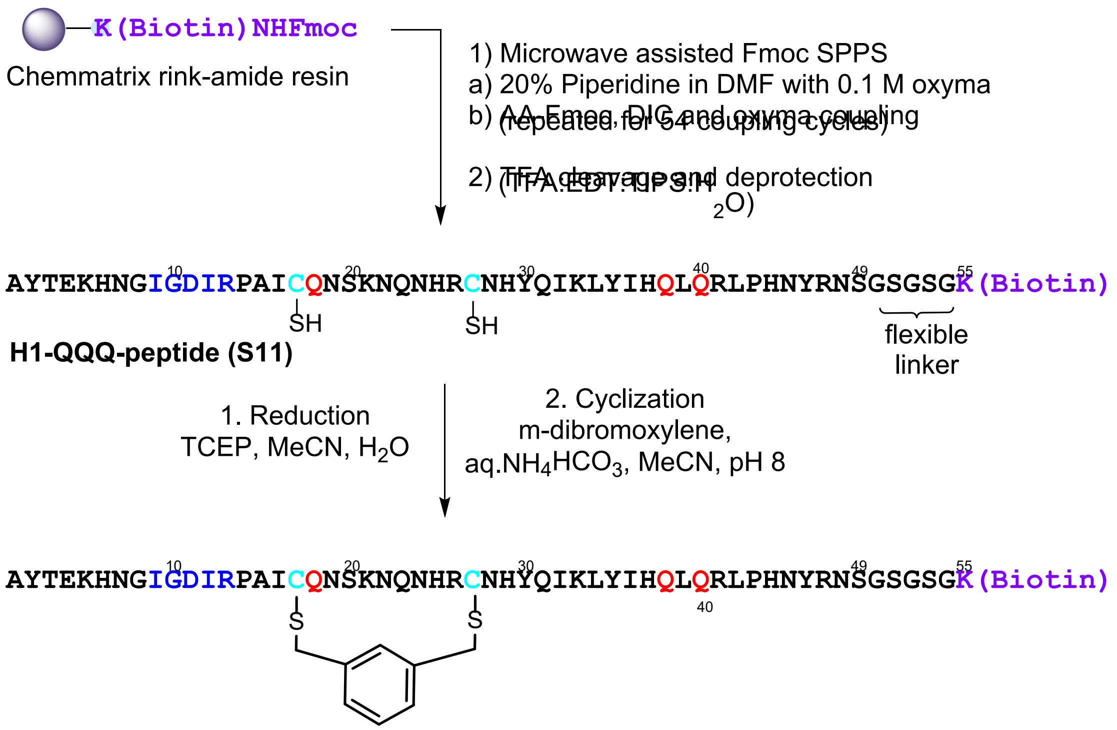

**H1-QQQ-cyclic non-glycosylated peptide (S12)**

**Scheme S7.** Synthetic scheme to generate **H1-QQQ-cyclic-(non-glycosylated) (S12).**

**H1-QQQ-peptide (S11):** H1-QQQ-peptide **(S11)** was prepared as in the representative procedure for solid phase peptide synthesis (Section III.A.4), starting with 180 mg of resin loaded at 0.17 mmol/g (30.6 µmol). Of the crude solid peptide, 40 mg was subjected to purification by RP-HPLC on the 10 x 250 mm C4 column (5-30% B over 45 min). 1.1 mg of pure peptide was obtained (2.6 % of the material that was purified). ESI-LRMS observed *m/z* of multiply charged ions 1098.44 $[M+6H]^{6+}$, 1317.81 $[M+5H]^{5+}$, 1646.91 $[M+4H]^{4+}$; calculated average m/z values for H1-QQQ-peptide (**S11**) $C_{282}H_{442}N_{96}O_{82}S_3$ : 1098.56 $[M+6H]^{6+}$, 1318.07 $[M+5H]^{5+}$, 1647.34 $[M+4H]^{4+}$.

**H1-QQQ-cyclic non-glycosylated peptide (S12):** The representative procedure for cyclization (Section III.A.7) was followed, using H1-QQQ-peptide (1.1 mg, 0.16µmol). The crude was purified on the 10 x 250 mm C4 column (5-35 % B over 45 minutes), to afford 0.49 mg (0.073 µmol) of H1-QQQ cyclic peptide (46% yield) (quantified by UV-nanodrop). ESI-LRMS observed *m/z* of multiply charged ions 744.33 $[M+9H]^{9+}$, 837.22 $[M+8H]^{8+}$, 956.63 $[M+7H]^{7+}$, 1115.69 $[M+6H]^{6+}$, 1138.52 $[M+5H]^{5+}$, 1673.09 $[M+4H]^{4+}$; calculated average m/z values for H1-QQQ-cyclic peptide (**S12**) $C_{290}H_{448}N_{96}O_{82}S_3$ : 744.06 $[M+9H]^{9+}$, 836.94 $[M+8H]^{8+}$, 956.36 $[M+7H]^{7+}$, 1115.58 $[M+6H]^{6+}$, 1138.50 $[M+5H]^{5+}$, 1672.87 $[M+4H]^{4+}$.

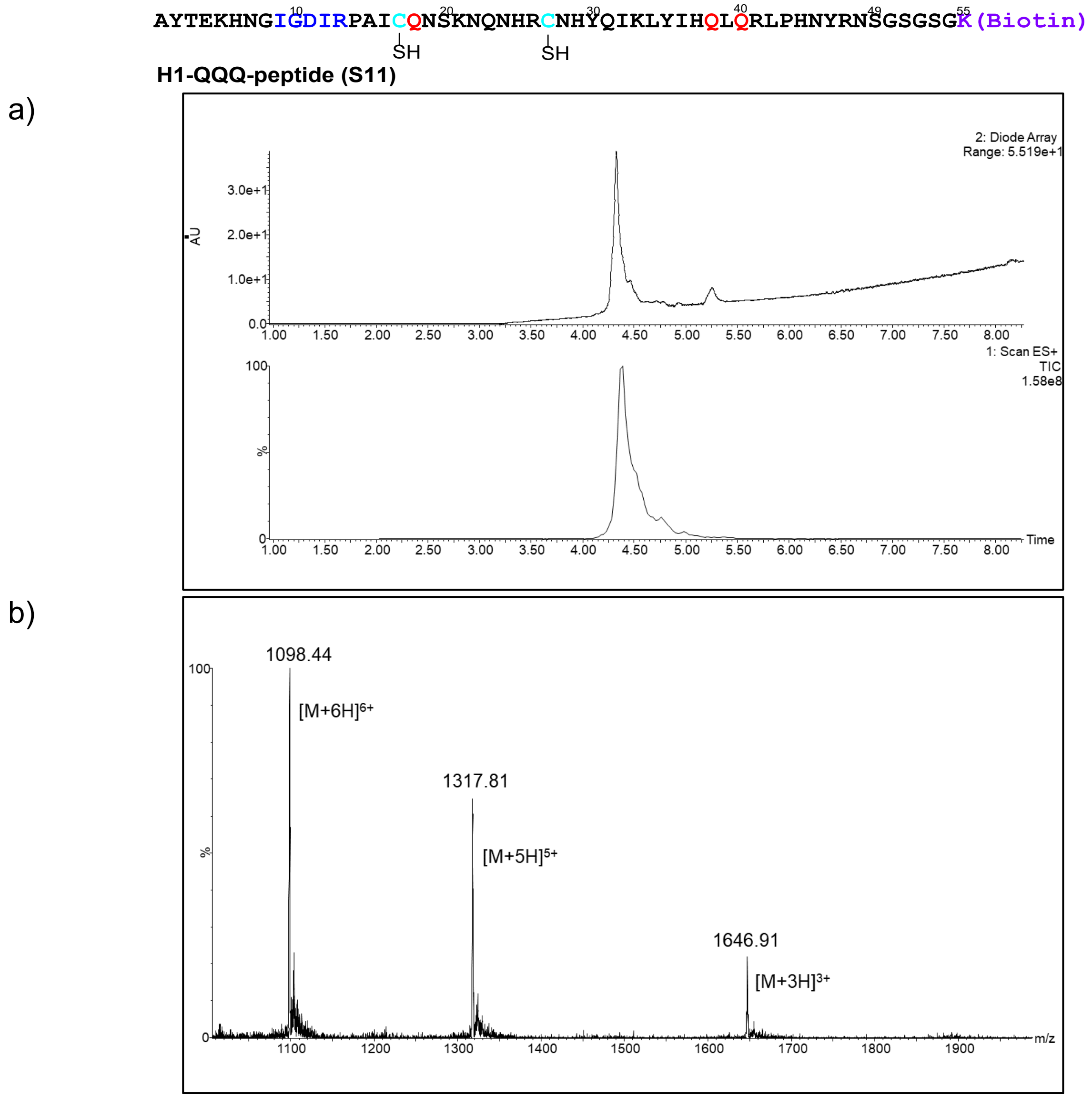

**Figure S24**. LC-MS of **H1-QQQ-peptide (S11)**. a) UV and ESI+ TIC chromatogram of H1-QQQ- peptide, b) ESI+ MS of H1-QQQ- peptide.

(LC-MS method: 1% A to 10% B over 1 minute then 10%A to 45%B gradient over 7 minutes, solvent A was water/0.07% formic acid and solvent B was acetonitrile/0.07% formic acid, column: Acquity UPLC@Protein BEH C4, 2.1 mm * 150 mm, 1.7 µm; flow rate: 0.25 mL per minute.)

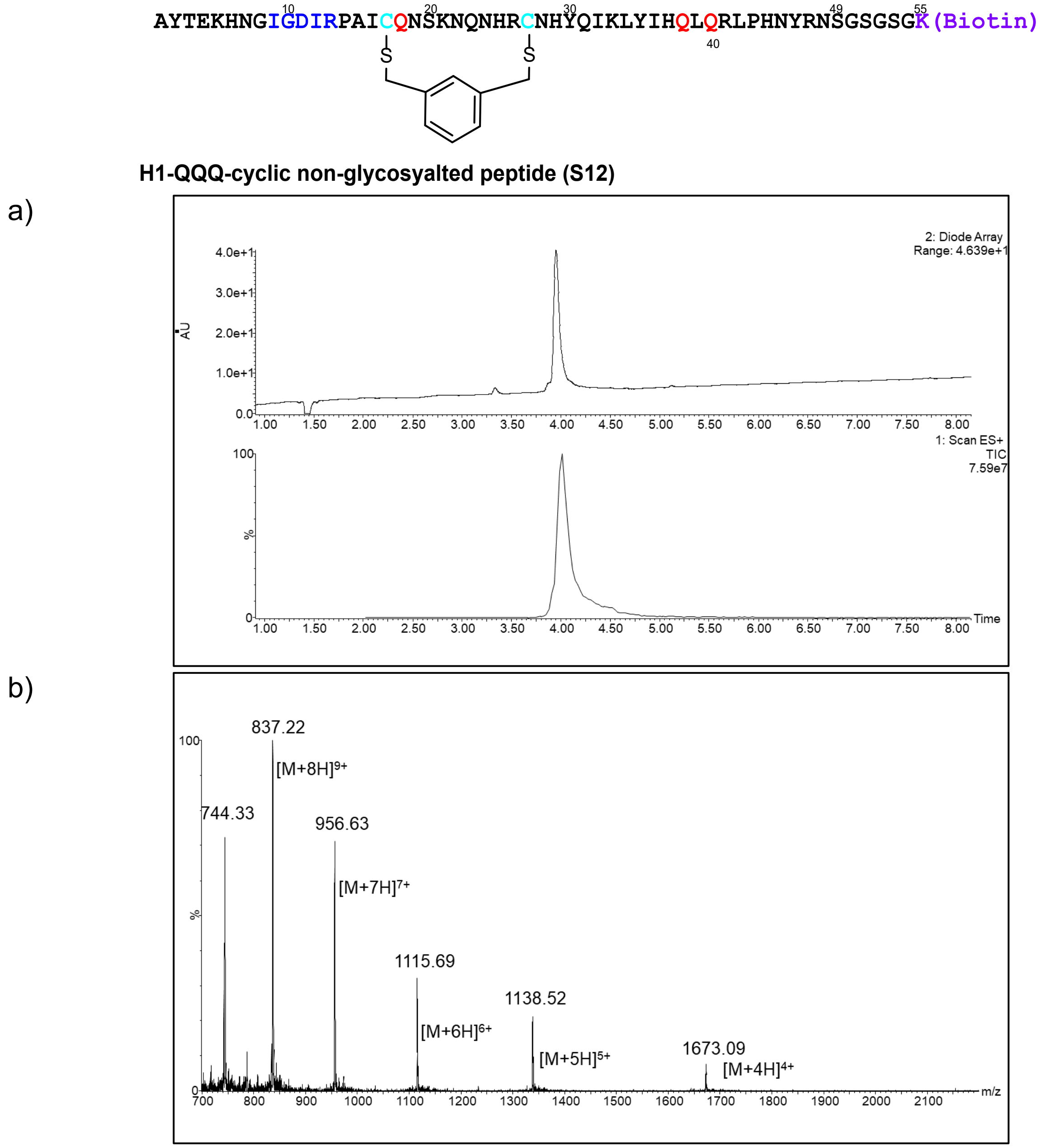

**Figure S25**. LC-MS of **H1-QQQ-cyclic-peptide (S12)**. a) UV and ESI+ TIC chromatogram of H1-QQQ-cyclic peptide, b) ESI+ MS of H1-QQQ-cyclic peptide.

(LC-MS method: 1% A to 10% B over 1 minute then 10%A to 45%B gradient over 7 minutes, solvent A was water/0.07% formic acid and solvent B was acetonitrile/0.07% formic acid, column: Acquity UPLC@Protein BEH C4, 2.1 mm * 150 mm, 1.7 µm; flow rate: 0.25 mL per minute.)

*7. H1-random-glycopeptide (**6**)*

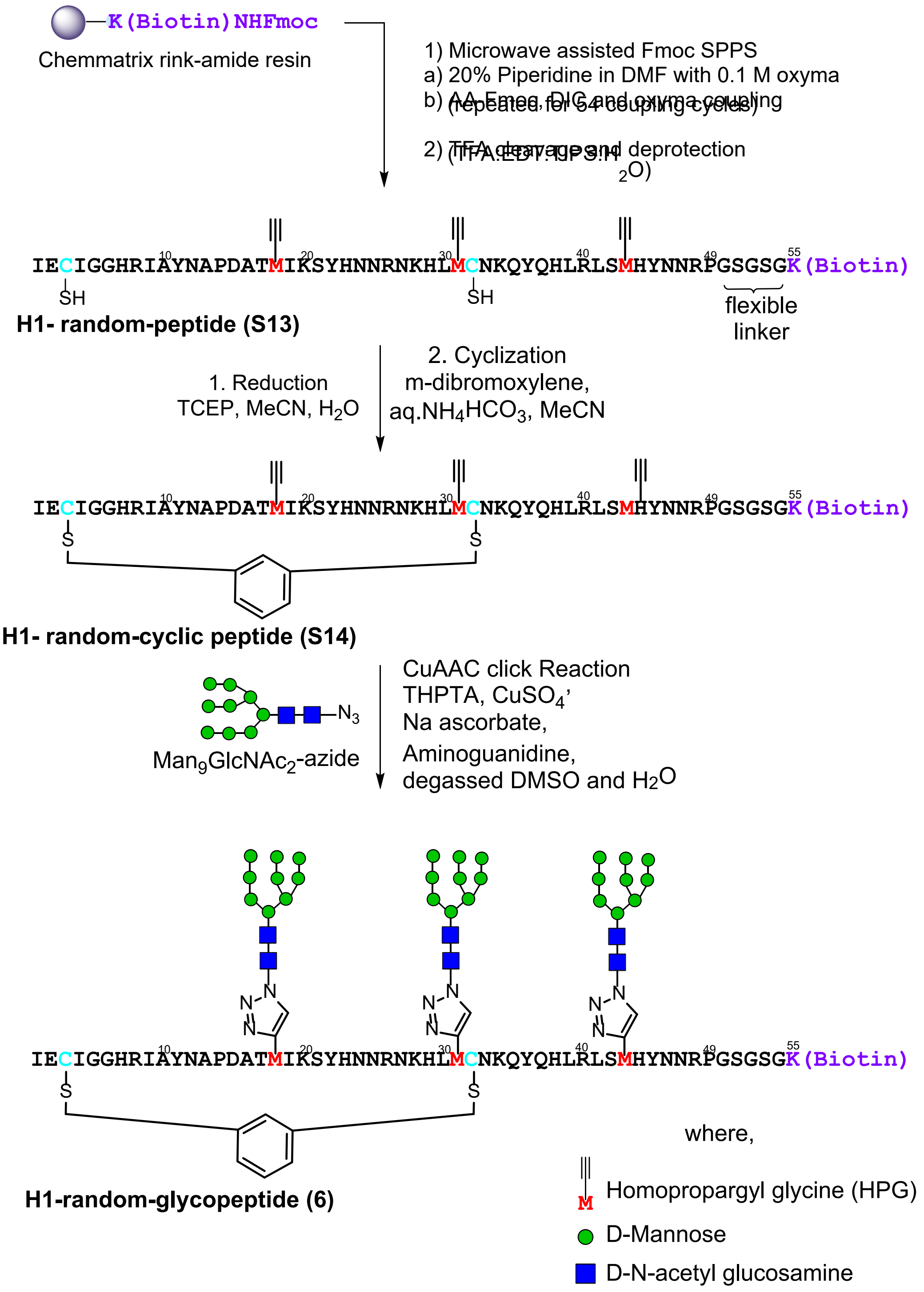

**Scheme S8.** Synthetic scheme to generate **H1-random-glycopeptide (6).**

**H1-random peptide (S13):** H1-random-peptide **(S13)** was prepared as in the representative procedure for solid phase peptide synthesis (Section III.A.4), starting with 180 mg of resin loaded at 0.17 mmol/g (30.6 µmol). Of the crude solid peptide, 40 mg was subjected to purification by RP-HPLC on the 10 x 250 mm C4 column (5-30% B over 45 min). 0.9 mg of pure peptide was obtained (2.2 % of the material that was purified). ESI-LRMS observed *m/z* of multiply charged ions 726.57 $[M+9H]^{9+}$, 817.06 $[M+8H]^{8+}$, 933.63 $[M+7H]^{7+}$, 1089.01 $[M+6H]^{6+}$, 1306.56 $[M+5H]^{5+}$; calculated average m/z values for H1-random-peptide **(S13)** $C_{285}H_{439}N_{93}O_{79}S_3$ : 726.38 $[M+9H]^{9+}$, 817.05 $[M+8H]^{8+}$, 933.62 $[M+7H]^{7+}$, 1089.06 $[M+6H]^{6+}$, 1306.67 $[M+5H]^{5+}$.

**H1-random-cyclic peptide (S14):** The representative procedure for cyclization (Section III.A.7) was followed, using H1-random peptide **(S13)** (0.98 mg, 0.15 µmol). The crude was purified on the 10 x 250 mm C4 column (5-35 % B over 45 minutes), to afford 0.43 mg (0.065 µmol) of H1-cyclic peptide (43 % yield) (quantified by UV-nanodrop). ESI-LRMS observed *m/z* of multiply charged ions 737.82 $[M+9H]^{9+}$, 829.81 $[M+8H]^{8+}$, 948.19 $[M+7H]^{7+}$, 1105.98 $[M+6H]^{6+}$, 1327.04 $[M+5H]^{5+}$, 1658.80 $[M+4H]^{4+}$; calculated average m/z values for H1-random cyclic peptide **(S14)** $C_{293}H_{445}N_{93}O_{79}S_3$ : 737.72 $[M+9H]^{9+}$, 829.81 $[M+8H]^{8+}$, 948.21 $[M+7H]^{7+}$, 1106.08 $[M+6H]^{6+}$, 1327.10 $[M+5H]^{5+}$, 1658.62 $[M+4H]^{4+}$.

**H1-random-glycopeptide (6):** The representative procedure for the click glycosylation of synthetic peptides (Section III.A.8) was conducted with H1-random cyclic peptide (**S14**) (0.43 mg, 0.065 µmol) and $Man_9GlcNAc_2$-azide (7.8 µL from 50 mM stock solution in water, 0.74 mg, 0.39 µmol). After HPLC purification on the 4.6 x 250 mm C4 column (5-30% B over 45 minutes), 43 µg (0.0035 µmol) of pure H1-glycopeptide (5 %) yield was obtained (quantified by UV-nanodrop). ESI-LRMS observed *m/z* of multiply charged ions 1123.99 $[M+11H]^{11+}$, 1236.47 $[M+10H]^{10+}$, 1373.65 $[M+9H]^{9+}$, 1545.36 $[M+8H]^{8+}$, 1766.33 $[M+7H]^{7+}$, 2060.42 $[M+6H]^{6+}$; calculated average m/z values for H1-random-glycopeptide **(6)** $C_{503}H_{796}N_{108}O_{244}S_3$ : 1124.32 $[M+11H]^{11+}$, 1236.66 $[M+10H]^{10+}$, 1373.95 $[M+9H]^{9+}$, 1545.57 $[M+8H]^{8+}$, 1766.22 $[M+7H]^{7+}$, 2060.42 $[M+6H]^{6+}$.

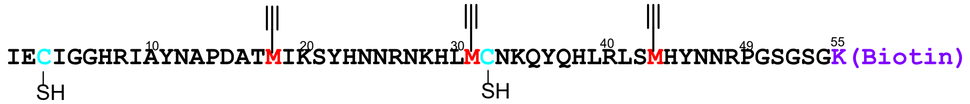

**H1- random-peptide (S13)**

a)

b)

c)

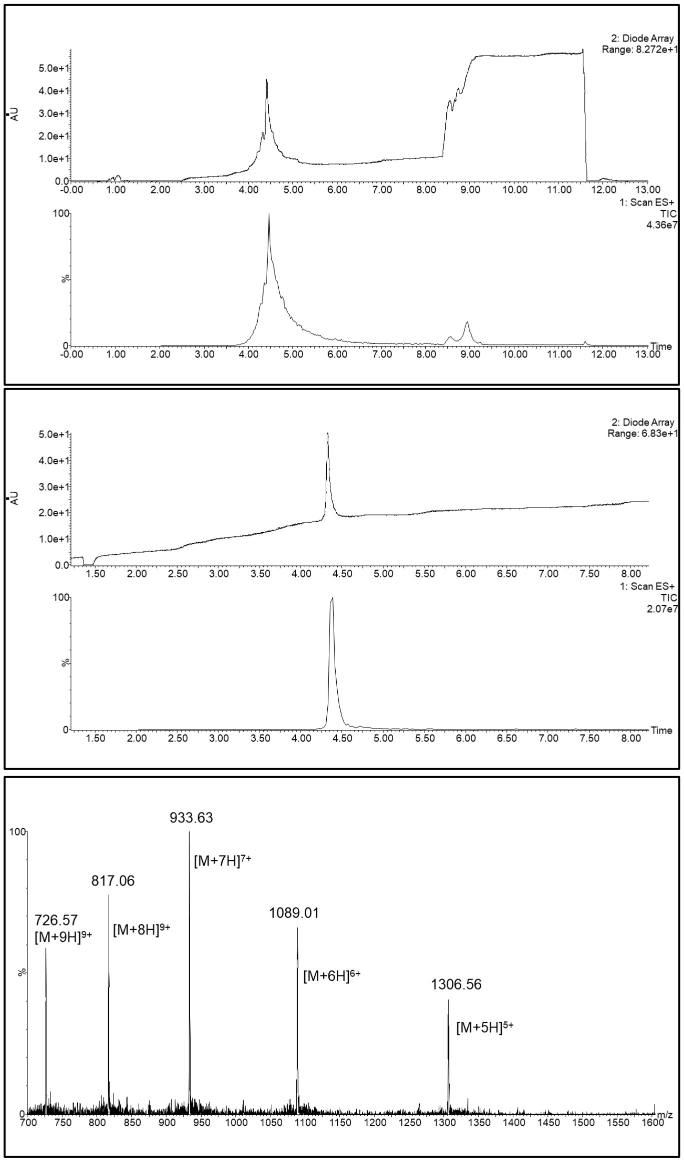

**Figure S26**. LC-MS of **H1-random-peptide (S13)**. a) UV and ESI+ TIC chromatogram of H1-random-peptide (crude), b) UV and ESI+ TIC chromatogram of H1-random-peptide (purified), c) ESI+ MS of H1-random-peptide.

(LC-MS method for crude H1-random peptide: 1% A to 10% B over 1 minute then 10%A to 45%B gradient over 8 minutes LC-MS method for H1-random-peptide (purified): 1% A to 10% B over 1 minute then 10%A to 45%B gradient over 7 minutes, solvent A was water/0.07% formic acid and solvent B was acetonitrile/0.07% formic acid, column: Acquity UPLC@Protein BEH C4, 2.1 mm * 150 mm, 1.7 µm; flow rate: 0.25 mL per minute)

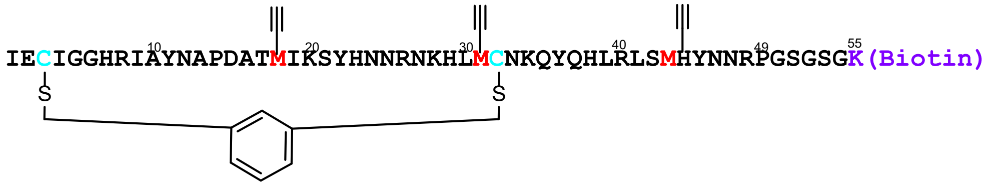

**H1- random-cyclic peptide (S14)**

a)

b)

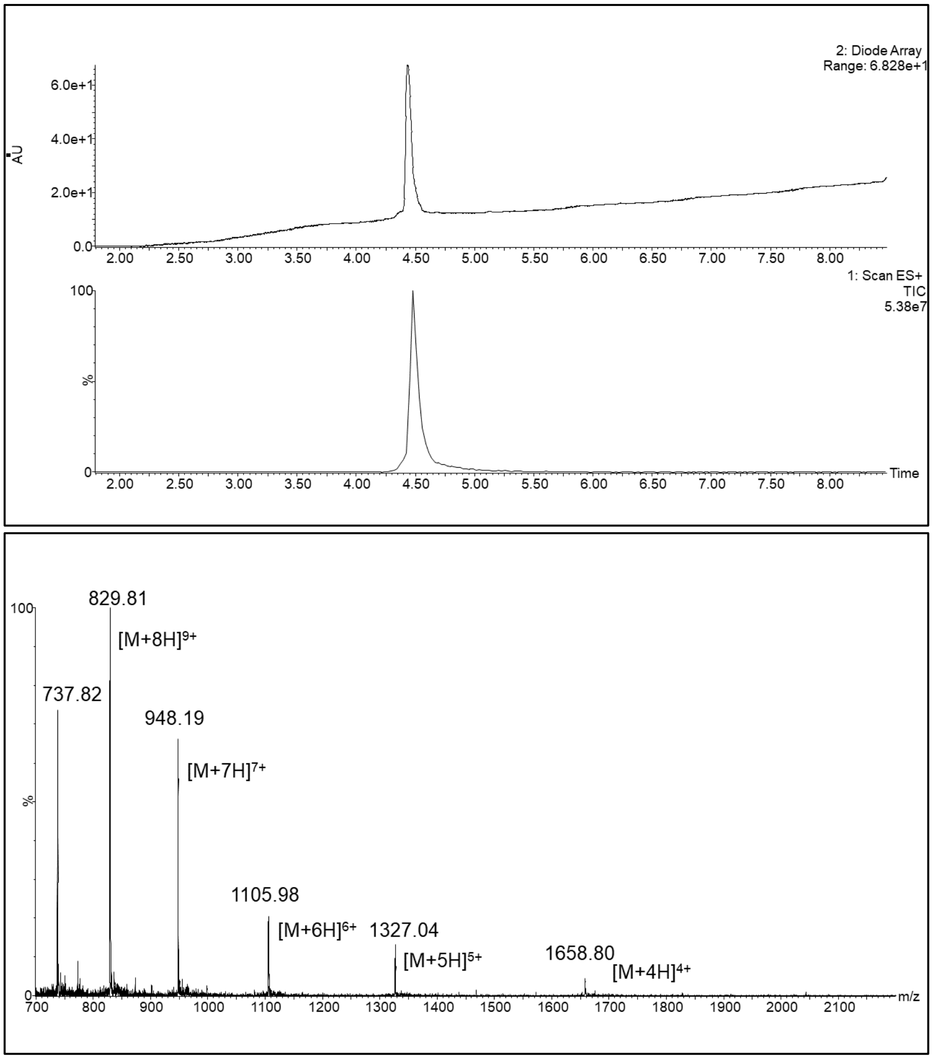

**Figure S27**. LC-MS of **H1-random-cyclic-peptide (S14)**. a) UV and ESI+ TIC chromatogram of H1-random-peptide, b) ESI+ MS of H1-random-cyclic-peptide.

(LC-MS method: 1% A to 10% B over 1 minute then 10%A to 45%B gradient over 7 minutes, solvent A was water/0.07% formic acid and solvent B was acetonitrile/0.07% formic acid, column: Acquity UPLC@Protein BEH C4, 2.1 mm * 150 mm, 1.7 µm; flow rate: 0.25 mL per minute.)

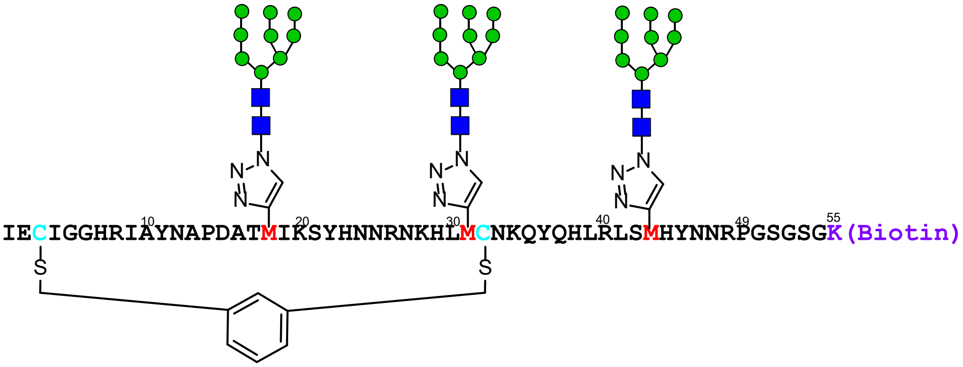

**H1-random-glycopeptide (6)**

a)

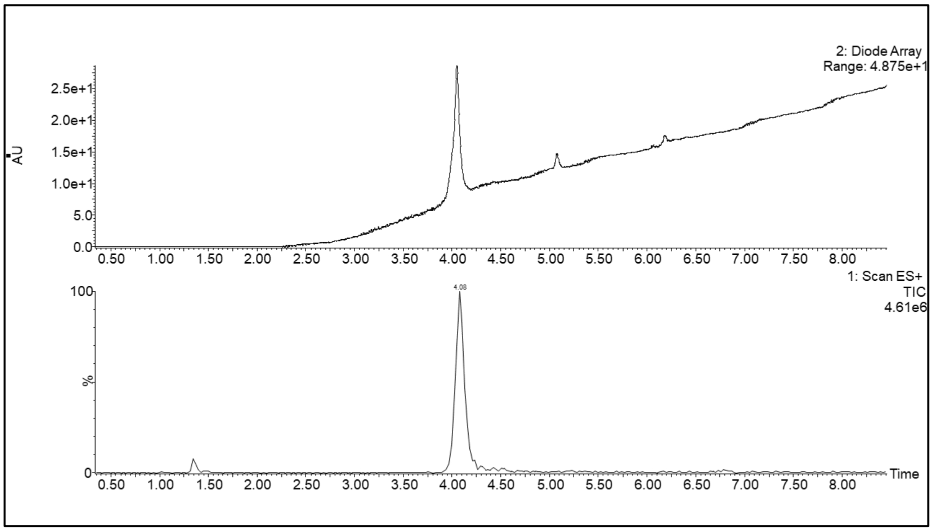

b)

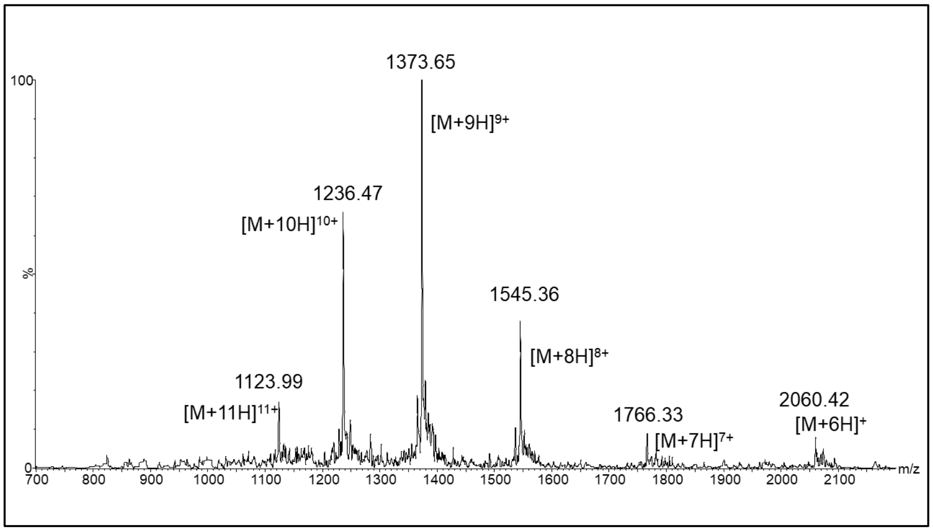

**Figure S28**. LC-MS of **H1-random-glycopeptide (6)**. a) UV and ESI+ TIC chromatogram of H1-random-peptide, b) ESI+ MS of H1-random-glycopeptide.

(LC-MS method: 1% A to 5% B over 1 minute then 5%A to 30%B gradient over 18 minutes, solvent A was water/0.07% formic acid and solvent B was acetonitrile/0.07% formic acid, column: Acquity UPLC@Protein BEH C4, 2.1 mm * 150 mm, 1.7 µm; flow rate: 0.25 mL per minute)

*8. H1-acyclic-glycopeptide (7)*

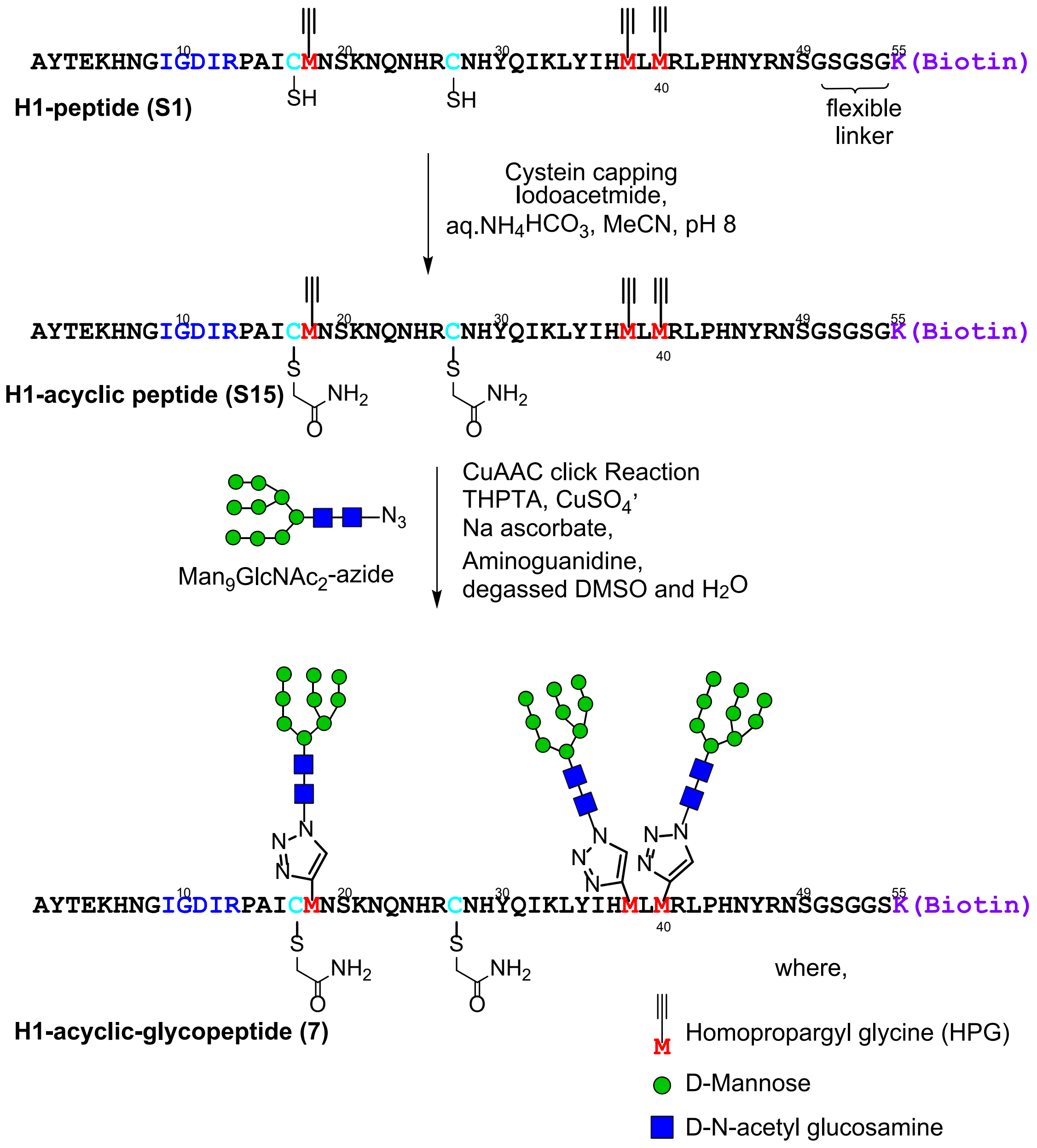

**Scheme S9.** Synthetic scheme to generate **H1-acyclic-glycopeptide (7).**

**H1-acyclic peptide (S15):** To the freshly purified and lyophilized H1-peptide (**S1**) (1.7 mg, 0.25 µmol) was added 500 µL of MQ water, 500 µL of acetonitrile, 16 µL of 40 mM iodoacetamide solution in acetonitrile (2.5 eq., 0.64 µmol) and 12 µL 1M solution of aq. ammonium bicarbonate adjusted to pH 8. The reaction mixture was allowed to react for 1 hour and excess of iodoacetamide was quenched by adding 3 µL of methanol. The crude was purified on the 10 x 250 mm C4 column (5-35 % B over 45 minutes), to afford 0.79 mg (0.12 µmol) of H1-acyclic-cyclic peptide **(S15)** (50 % yield) (quantified by UV-nanodrop). ESI-LRMS observed *m/z* of multiply charged ions 739.52 $[M+9H]^{9+}$, 831.63 $[M+8H]^{8+}$, 950.19 $[M+7H]^{7+}$, 1108.41 $[M+6H]^{6+}$, 1329.48 $[M+5H]^{5+}$, 1661.90 $[M+4H]^{4+}$; calculated average m/z values for H1-acyclic peptide **(S15)** $C_{289}H_{445}N_{95}O_{81}S_3$ : 739.05 $[M+9H]^{9+}$, 831.31 $[M+8H]^{8+}$, 949.92 $[M+7H]^{7+}$, 1108.08 $[M+6H]^{6+}$, 1329.49 $[M+5H]^{5+}$, 1661.61 $[M+4H]^{4+}$.

**H1-acyclic-glycopeptide (7):** The representative procedure for the click glycosylation of synthetic peptides (Section III.A.8) was conducted with H1-acyclic peptide **(S15)** (0.32 mg, 0.05 µmol) and $Man_9GlcNAc_2$-azide (4.5 µL from 50 mM stock solution in water, 0.42 mg, 0.22 µmol). After HPLC purification on the 4.6 x 250 mm C4 column (5-30% B over 45 minutes), 55 µg (0.0044 µmol) of pure H1-acyclic-glycopeptide **(7)** (4 %) yield was obtained (quantified by UV-nanodrop). ESI-LRMS observed *m/z* of multiply charged ions 952.99 $[M+13H]^{13+}$, 1032.36 $[M+12H]^{12+}$, 1125.57 $[M+11H]^{11+}$, 1238.09 $[M+10H]^{10+}$, 1375.38 $[M+9H]^{9+}$, 1547.11 $[M+8H]^{8+}$; calculated average m/z values for H1-acyclic-glycopeptide **(7)** $C_{499}H_{796}N_{110}O_{246}S_3$ : 952.43 $[M+13H]^{13+}$, 1031.71 $[M+12H]^{12+}$, 1125.41 $[M+11H]^{11+}$, 1237.85 $[M+10H]^{10+}$, 1375.28 $[M+9H]^{9+}$, 1547.06 $[M+8H]^{8+}$.

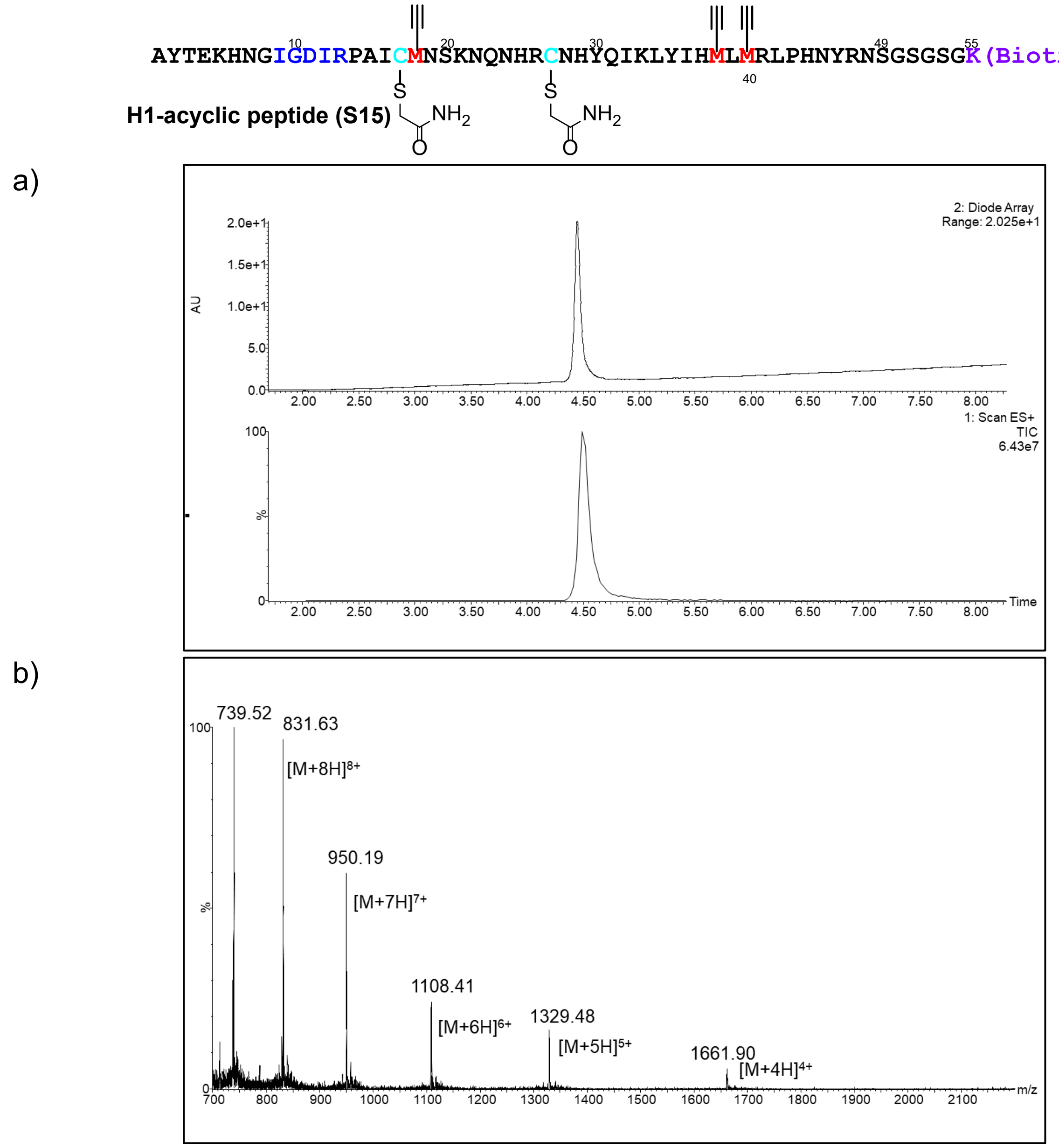

**Figure S29**. LC-MS of **H1-acyclic-peptide (S15)**. a) UV and ESI+ TIC chromatogram of H1-acyclic-peptide, b) ESI+ MS of H1-acyclic-peptide.

(LC-MS method: 1% A to 10% B over 1 minute then 10%A to 45%B gradient over 7 minutes, solvent A was water/0.07% formic acid and solvent B was acetonitrile/0.07% formic acid, column: Acquity UPLC@Protein BEH C4, 2.1 mm * 150 mm, 1.7 µm; flow rate: 0.25 mL per minute.)

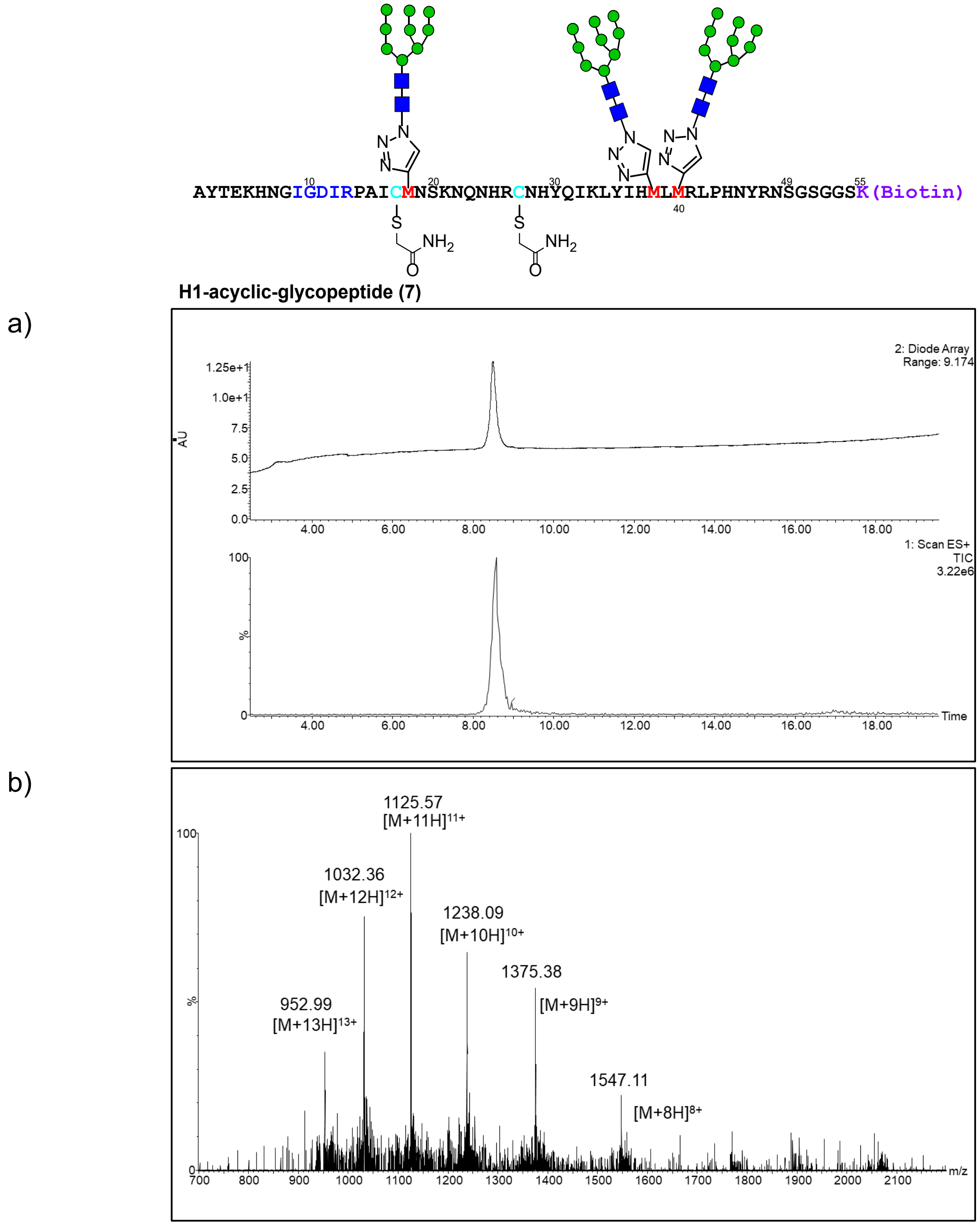

**Figure S30**. LC-MS of **H1-acyclic-glycopeptide (7)**. a) UV and ESI+ TIC chromatogram of H1-acyclic-peptide, b) ESI+ MS of H1-acyclic-glycopeptide.

(LC-MS method: 1% A to 5% B over 1 minute then 5%A to 30%B gradient over 18 minutes, solvent A was water/0.07% formic acid and solvent B was acetonitrile/0.07% formic acid, column: Acquity UPLC@Protein BEH C4, 2.1 mm * 150 mm, 1.7 µm; flow rate: 0.25 mL per minute)

*9. L1-glycopeptide (**8**)*

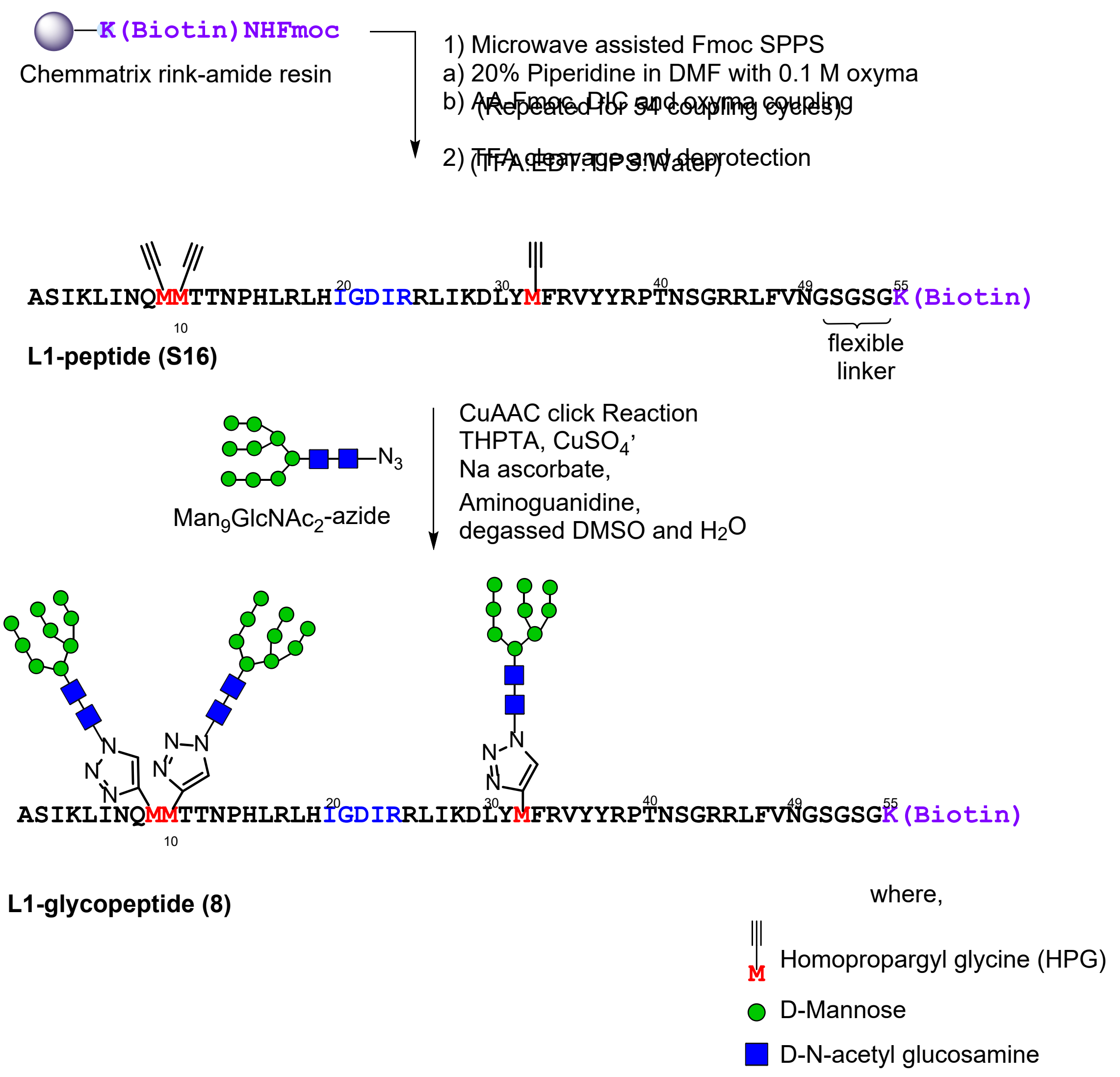

**Scheme S10.** Synthetic scheme to generate **L1-glycopeptide (8).**

**L1-peptide (S16):** L1-peptide (**S16**) was prepared by the general procedure for solid phase peptide synthesis, starting with 417 mg of resin loaded at 0.12 mmol/g (50 µmol). Of the crude solid peptide, 80 mg was subjected to purification by RP-HPLC on the 10 x 250 mm C4 column (5-30% B over 45 min). 1.04 mg of pure peptide (quantified by UV-nanodrop) was obtained (1.25 % of the purified material). ESI-LRMS observed *m/z* of multiply charged ions 731.27 $[M+9H]^{9+}$, 822.46 $[M+8H]^{8+}$, 939.40 $[M+7H]^{7+}$, 1096.13 $[M+6H]^{6+,}$ 1315.12 $[M+5H]^{5+}$; calculated average m/z for L1-peptide (**S16**) $C_{297}H_{473}N_{91}O_{76}S_1$ : 730.63 $[M+9H]^{9+}$, 821.83 $[M+8H]^{8+}$, 939.09 $[M+7H]^{7+}$, 1095.44 $[M+6H]^{6+}$, 1314.32 $[M+5H]^{5+}$

**L1-glycopeptide (8):** The standard procedure for the click reaction was conducted with L1- peptide (**S16**) (0.54 mg, 0.08 µmol) and $Man_9GlcNAc_2$-azide (7.2 µL from 50 mM solution in water, 0.68 mg, 0.36 µmol). After HPLC purification on the 4.6 x 250 mm C4 column (5-30% B over 45 minutes), 120 µg (0.0098 µmol) of pure L1-glycopeptide (**3**) (12 %) yield was obtained (quantified by UV-nanodrop). ESI-LRMS observed *m/z* of multiply charged ions 1367.25 $[M+9H]^{9+}$, 1537.62 $[M+8H]^{8+}$, 1757.01 $[M+7H]^{7+}$; calculated average m/z for L1-glycopeptide (**8**) $C_{507}H_{824}N_{106}O_{241}S_1$ : 1366.85 $[M+9H]^{9+}$, 1537.58 $[M+8H]^{8+}$, 1757.09 $[M+7H]^{7+}$. Fragment ions were observed for higher charged species, corresponding to loss of one $Man_9GlcNAc_2$ from $[M+9H]^{9+}$, calc 1159.56, or two $Man_9GlcNAc_2$ moieties from $[M+8H]^{8+}$, calc 1071.17 and $[M+9H]^{9+}$, calc 952.26.

ASIKLINQMMTTNPHLRLHIGDIRRLIKDLYMFRVYYRPTNSGRRLFVNGSGSGK(Biotin)

**L1-peptide (S16)**

a)

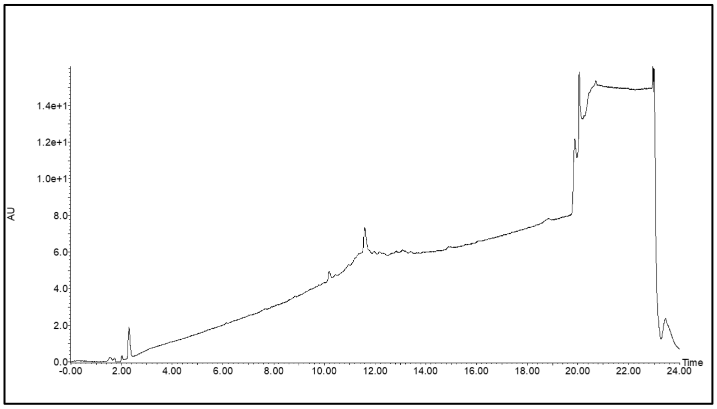

b)

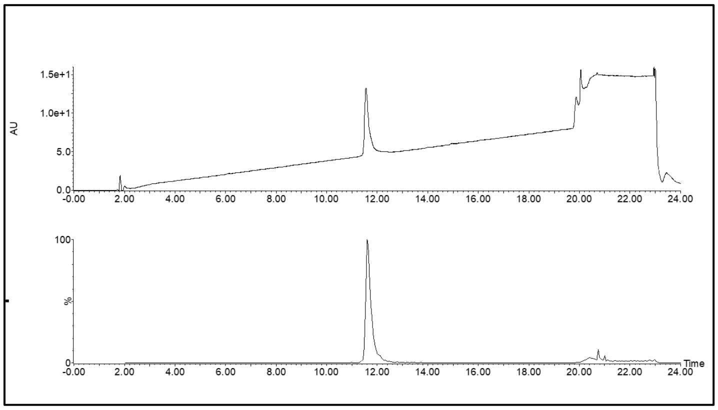

c)

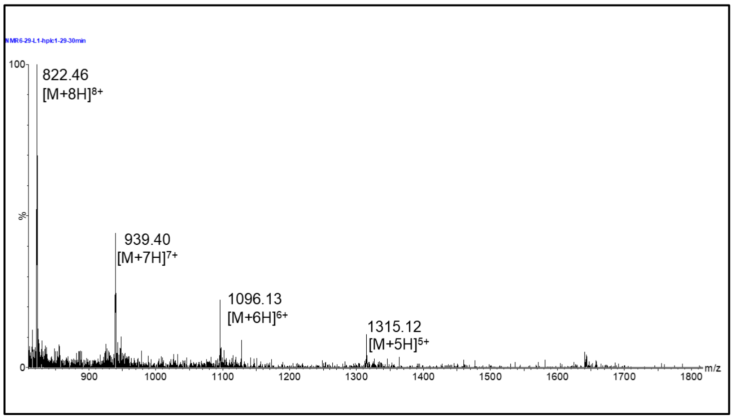

**Figure S31**. LC-MS of **L1-peptide (S16)**. a) UV and ESI+ TIC chromatogram of L1-peptide (crude), b) UV and ESI+ TIC chromatogram of L1-peptide (purified), c) ESI+ MS of L1-peptide.

(LC-MS method for crude L1-peptide: 1% A to 5% B over 1 minute then 5%A to 45%B gradient over 18 minutes LC-MS method for purified L1 peptide: 1% A to 5% B over 1 minute then 5%A to 45%B gradient over 18 minutes, solvent A was water/0.07% formic acid and solvent B was acetonitrile/0.07% formic acid, column: Acquity UPLC@Protein BEH C4, 2.1 mm * 150 mm, 1.7 μm; flow rate: 0.2 mL per minute)

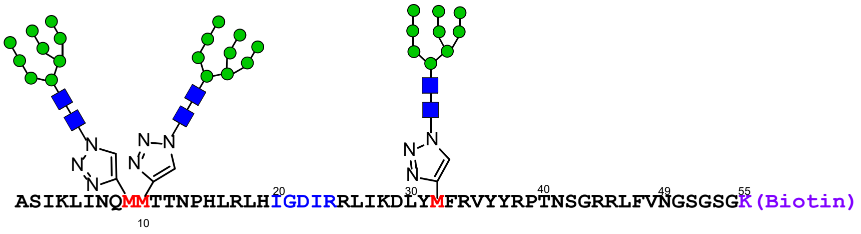

**L1-glycopeptide (8)**

a
)

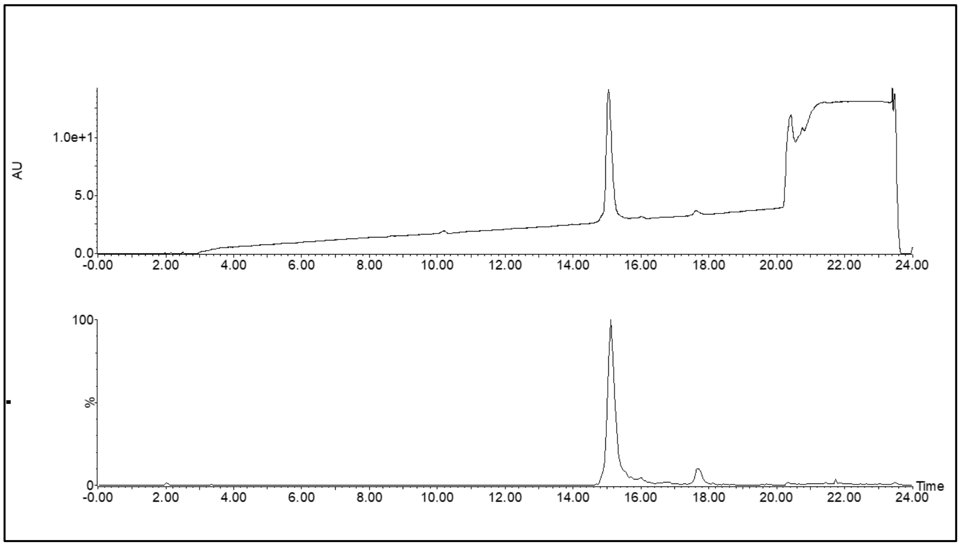

b
)

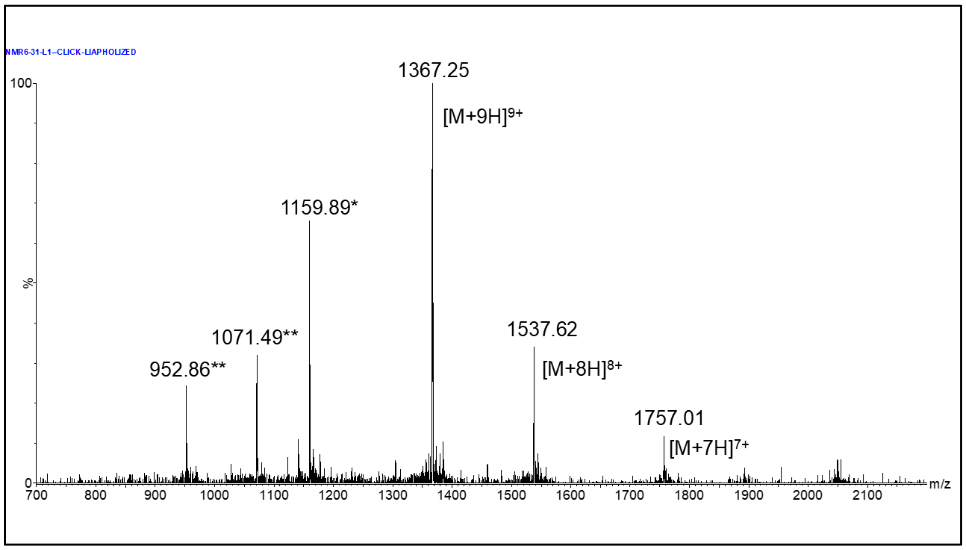

**Figure S32**. LC-MS of **L1-glycopeptide (8)**. a) UV and ESI+ TIC chromatogram of L1-glycopeptide b) ESI+ MS of L1-glycopeptide. (*corresponds to loss of $Man_9GlcNAc_2$ from $[M+9H]^{9+}$, calc 1159.56, ** corresponds to 2 Man9GlcNAc2 moieties from $[M+8H]^{8+}$, calc 1071.17 and $[M+9H]^{9+}$, calc 952.26)
(LC-MS method: 1% A to 5% B over 1 minute then 5%A to 30%B gradient over 18 minutes, solvent A was water/0.07% formic acid, and solvent B was acetonitrile/0.07% formic acid, column: Acquity UPLC@Protein BEH C4, 2.1 mm * 150 mm, 1.7 µm; flow rate: 0.2 mL per minute.)

### C. Binding Analysis of PGT Antibody-Peptide/Glycopeptide Interactions by Biolayer Interferometry

*1. Biolayer Interferometry Protocol*

Biotinylated glycopeptide was loaded onto a streptavidin biosensor in 250 µL buffer 1 (500 to 1000 nM X nM glycopeptide in 20 mM Tris pH 7.5, 150 mM NaCl, 0.2 mg/ml BSA, 0.02% (v/v) Tween-20) for 120s in a 0.5 µL of black Eppendorf tube. The sensor was then washed with 250 µL of blank buffer 1 for 120 seconds in a 0.5 µL black Eppendorf tube. The sensor was washed with 250 µL buffer 3 (10 mM glycine HCl, pH 2.5) for 120 seconds in 0.5 µL of black Eppendorf tube and followed by 250 µL of blank buffer 1 for 120 seconds in a 0.5 µL black Eppendorf tube to immobilize glycopeptides in a range of 0.5 nm to 2 nm response. The sensor was then equilibrated with 250 µL of buffer 2 (20 mM Tris pH 7.5, 150 mM NaCl, 2 mg/ml BSA, 0.1% (v/v) Tween-20) for 120 seconds in a 0.5 µL of black Eppendorf tube PGT antibody (prepared in buffer 2) was associated at several concentrations for 300 seconds in 4 µL drop holder followed by dissociation into blank 250 µL of buffer 2 for 300 seconds in 0.5 µL of black Eppendorf tube. After each dissociation sensor was regenerated to remove the remaining antibody by treatment with 250 µL of buffer 3 for 120 seconds, followed by 120 seconds of wash with 250 µL buffer 1 and re-equilibration with 250 µL buffer 2 in 0.5 µL of black Eppendorf tube. Throughout the experiment, the shaker rate was set at 2200 rpm. Data were fit globally to a 1:1 binding model.

*2. BLI Sensorgram of H1-QQQ-non-glycosylated Peptide with PGT128*

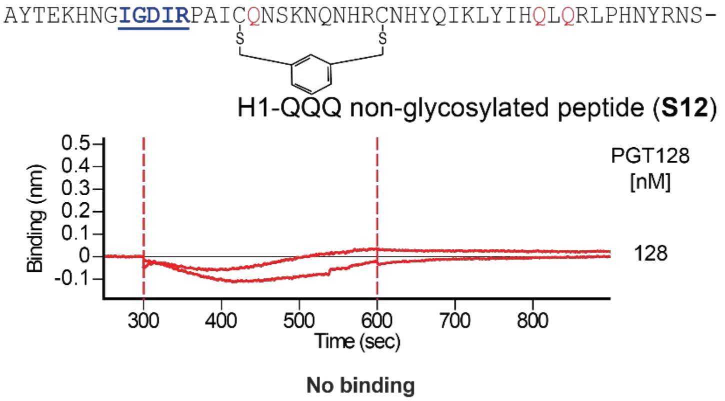

**Figure S33**. BLI Sensorgram of H1-QQQ-non-glycosylated peptide-biotin immobilized on a streptavidin biosensor treated with 128 and 64 nM PGT128 according to the procedure in **Section III.C.1**. Only the experimental curves are shown.

### IV. PGT122/gl-PGT121 Selections

#### A. Library Generation for Round 1 PGT122/gl-PGT121 Selections

*1. Preparation of Library DNA*

Random 193-mer oligos for the Unbiased library (XNS51AS193) were purchased at Keck Biotechnology Resource Laboratory, Yale University (New Haven, CT; 200 nmol) and Biolegio (Nijmegen, Netherlands; 200 nmol), and random 193-mer oligos for the Biased library (X8X33AS193, X21X20AS193, and X34X7AS193) were purchased at IDT (Ultramer® DNA Oligos, 4 nmol; summarized in **Table S10**; theoretical multivalency distribution given in **Figure S34**). The oligos were purified by denaturing urea-PAGE individually. The purified XNS51AS193 oligos from both providers were combined to ~540 pmol, and the purified oligos for the Biased library were combined at an equimolar ratio for a total of ~1 nmol, separately annealed with SSSF1 (1.5 eq.; see **Table S11** for PGT122/gl-PGT121 primer sequences), and extended by DNA Polymerase I, Large (Klenow) Fragment (New England Biolabs) to make each of the library fragments (225 bp). The dsDNAs were diluted 20-fold and used as a template to be further amplified and extended by PCR with Taq polymerase to make a whole library fragment (258 bp; **Table S12**). The PCR was 10 cycles of 94 °C for 30 seconds, 55 °C for 30 seconds, and 74 °C for 30 seconds using a set of primers (SSSF1 and SSSR1) for the Unbiased library and another set of primers (SSSF1 and SSSR2) for the Biased library. The dsDNA libraries were concentrated by isopropanol precipitation and purified by 6% non-denaturing PAGE, yielding ~280 and ~250 pmol of Unbiased and Biased library dsDNA, respectively. The purified library DNAs were diluted 100-fold and further amplified by PCR with Taq polymerase and a set of 1 μM each of primers (Library FP1 and Library RP1.3) for the Unbiased library or another set of primers (Library FP1 and Library RP2.1) for the Biased library via 12 thermal cycles of 94 °C for 30 seconds, 73 °C for 30 seconds, and 74 °C for 30 seconds. The solutions were extracted with Tris-saturated phenol: chloroform: isoamyl alcohol (25:24:1, v/v/v) and then chloroform followed by concentration by butanol. The DNA was recovered by isopropanol precipitation followed by 70% (v/v) ethanol rinse, and the yields of Unbiased and Biased libraries were ~500 pmol and ~430 pmol, respectively. 260 pmol each of the libraries were used for transcription in the next step.

**Table S10**: Individual libraries for selection with PGT antibodies. The Unbiased library comprises a single library with randomized doped *N*NS codons, where *N* is a mixture of 40 % A, 20 % G, 20 % T/U and 20 % C, resulting in an elevated 5% chance of an AUG codon at any single randomized position. The Biased library comprises three individual libraries with the conserved IGDIRXAXCM motif after 8, 21, or 34 randomized codons in the sequence. Codon degeneracy was exploited in the C-terminal constant region to design orthogonal primers for the two library pools. The resulting open reading frames of each individual library are shown, lacking the implicitly-cleaved N-terminal HPG. The antisense DNA sequences of each library, as ordered from Keck, Biolegio, and IDT, are also shown.

| Library | Name | Sequence 5′ to 3′ [a] |
|---|---|---|
| Unbiased | XNS51AS193[b] | AXXXXXXXXXXXXXXXXXXXXXXXXXXXXXXXXXXXXXXXXXXXXXXXXXXX |
| | | TGGCCTAAGCTACCGGAGCCSN5SN5SN5SN5SN5SN5SN5SN5SN5SN5SN5SN<br>5SN5SN5SN5SN5SN5SN5SN5SN5SN5SN5SN5SN5SN5SN5SN5SN5SN5SN5<br>SN5SN5SN5SN5SN5SN5SN5SN5SN5SN5SN5SN5SN5SN5SN5SN5SN5SN5S<br>N5SN5SN5CGCCATTTAGCTGTCCTCCT |
| Biased | X8X33AS193 | AXXXXXXXXIGDIRXAXCMXXXXXXXXXXXXXXXXXXXXXXXXXXXXXXXXX |
| | | TGACCCAGAGAACCGGAGCCSNNSNNSNNSNNSNNSNNSNNSNNSNNSNNSNNSN<br>NSNNSNNSNNSNNSNNSNNSNNSNNSNNSNNSNNSNNSNNSNNSNNSNNSNNSNN<br>SNNSNNSNNCATGCASNNCGCSNNACGGATATCGCCAATSNNSNNSNNSNNSNNS<br>NNSNNSNNCGCCATTTAGCTGTCCTCCT |
| | X21X20AS193 | AXXXXXXXXXXXXXXXXXXXXXIGDIRXAXCMXXXXXXXXXXXXXXXXXXXX |
| | | TGACCCAGAGAACCGGAGCCSNNSNNSNNSNNSNNSNNSNNSNNSNNSNNSNNSN<br>NSNNSNNSNNSNNSNNSNNSNNSNNCATGCASNNCGCSNNACGGATATCGCCAAT<br>SNNSNNSNNSNNSNNSNNSNNSNNSNNSNNSNNSNNSNNSNNSNNSNNSNNSNNS<br>NNSNNSNNCGCCATTTAGCTGTCCTCCT |
| | X34X7AS193 | AXXXXXXXXXXXXXXXXXXXXXXXXXXXXXXXXXXIGDIRXAXCMXXXXXXX |
| | | TGACCCAGAGAACCGGAGCCSNNSNNSNNSNNSNNSNNSNNCATGCASNNCGCSN<br>NACGGATATCGCCAATSNNSNNSNNSNNSNNSNNSNNSNNSNNSNNSNNSNNSNN<br>SNNSNNSNNSNNSNNSNNSNNSNNSNNSNNSNNSNNSNNSNNSNNSNNSNNSNNS<br>NNSNNSNNCGCCATTTAGCTGTCCTCCT |

a Each peptide sequence is followed by −GSGSLGHHHHHHRL.
b “5” is a mixture of 40% T, 20% G, 20% C, 20% A.

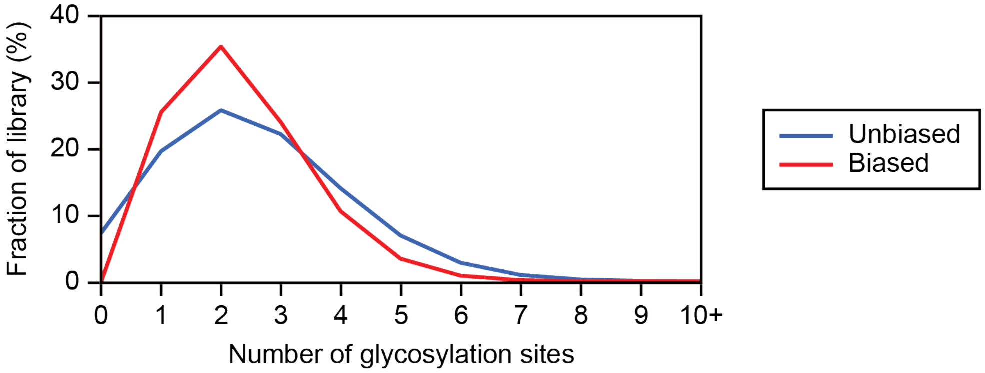

**Figure S34**. Theoretical multivalency distributions for Unbiased and Biased libraries for PGT122 and gl-PGT121 selections.

**Table S11**. List of primers used in PGT122/gl-PGT121 selections.

| Name | Sequence 5′ to 3′ [a] |
|---|---|
| SSSF1 | TAATACGACTCACTATAGGGTTAACTTTAGTAAGGAGGACAGCTAAATGGCG |
| SSSR1 | CTAGCTACCTATAGCCGGTGGTGATGGTGATGGTGGCCTAAGCTACCGGAGCC |
| SSSR2 | CTAGCTACCTATAGCCGGTGGTGATGGTGGTGATGACCCAGAGAACCGGAGCC |
| Library FP1 | TAATACGACTCACTATAGGGTTAACTTTAGTAAGGAGG |
| Library RP1.3 | CTAGCTACCTATAGCCGGTGGTGATGGTGATGGTGGCCTAAGC |
| Library RP2.1 | CTAGCTACCTATAGCCGGTGGTGATGGTGGTGATGACCCAGAG |
| PCB-18-RT-primer1 | /5PCBio//iSp18/TTTTTTTTTTTTTTTGTGATGGTGATGGTGGCCTAAGC |
| PCB-18-RT-primer2 | /5PCBio//iSp18/TTTTTTTTTTTTTTTGTGATGGTGGTGATGACCCAGAG |
| SPL1 | TTTTTTTTTTGTGATGGTGAT |
| SPL1N | TTTTTTTTTTNGTGATGGTGAT |
| SPL2 | TTTTTTTTTTGTGATGGTGGT |
| SPL2N | TTTTTTTTTTNGTGATGGTGGT |

**Table S12**: Extended library DNA sequences. The full sense DNA sequences after single-strand synthesis and PCR with the extending primers are shown, with the added constant regions underlined.

| Library | Name | Sequence 5′ to 3′ |
|---|---|---|
| Unbiased | XNS51AS193 full[a] | TAATACGACTCACTATAGGGTTAACTTTAGTAAGGAGGACAGCTAAATGGCG*N*NS*N*NS*N*NS*N*NS*N*NS*N*NS*N*NS*N*NS*N*NS*N*NS*N*NS*N*NS*N*NS*N*NS*N*NS*N*NS*N*NS*N*NS*N*NS*N*NS*N*NS*N*NS*N*NS*N*NS*N*NS*N*NS*N*NS*N*NS*N*NS*N*NS*N*NS*N*NS*N*NS*N*NS*N*NS*N*NS*N*NS*N*NS*N*NS*N*NS*N*NS*N*NS*N*NS*N*NS*N*NS*N*NS*N*NS*N*NS*N*NS*N*NS*N*NS*N*NSGGCTCCGGTAGCTTAGGCCACCATCACCATCACCACCGGCTATAGGTAGCTAG |
| Biased | X8X33AS193 full | TAATACGACTCACTATAGGGTTAACTTTAGTAAGGAGGACAGCTAAATGGCGNNSNNSNNSNNSNNSNNSNNSNNSATTGGCGATATCCGTNNSGCGNNSTGCATGNNSNNSNNSNNSNNSNNSNNSNNSNNSNNSNNSNNSNNSNNSNNSNNSNNSNNSNNSNNSNNSNNSNNSNNSNNSNNSNNSNNSNNSNNSNNSNNSGGCTCCGGTTCTCTGGGTCATCACCACCATCACCACCGGCTATAGGTAGCTAG |
| | X21X20AS193 full | TAATACGACTCACTATAGGGTTAACTTTAGTAAGGAGGACAGCTAAATGGCGNNSNNSNNSNNSNNSNNSNNSNNSNNSNNSNNSNNSNNSNNSNNSNNSNNSNNSNNSNNSNNSATTGGCGATATCCGTNNSGCGNNSTGCATGNNSNNSNNSNNSNNSNNSNNSNNSNNSNNSNNSNNSNNSNNSNNSNNSNNSNNSNNSNNSGGCTCCGGTTCTCTGGGTCATCACCACCATCACCACCGGCTATAGGTAGCTAG |
| | X34X7AS193 full | TAATACGACTCACTATAGGGTTAACTTTAGTAAGGAGGACAGCTAAATGGCGNNSNNSNNSNNSNNSNNSNNSNNSNNSNNSNNSNNSNNSNNSNNSNNSNNSNNSNNSNNSNNSNNSNNSNNSNNSNNSNNSNNSNNSNNSNNSNNSNNSNNSATTGGCGATATCCGTNNSGCGNNSTGCATGNNSNNSNNSNNSNNSNNSNNSGGCTCCGGTTCTCTGGGTCATCACCACCATCACCACCGGCTATAGGTAGCTAG |

a *N* is a mixture of 40% A, 20% G, 20% C, 20% T.

*2. Preparation of Puromycin-modified RNA*

3.3 mL of T7 transcription reaction contained 80 mM HEPES-KOH (pH 7.6), 40 mM DTT, 2 mM spermidine, 50 mg/mL PEG-8000, 4 mM GTP, 4 mM UTP, 4 mM ATP, 4 mM CTP, 25 mM $MgCl_2$, 2.5 U/mL inorganic pyrophosphatase, 0.05 mg/mL T7 RNA polymerase, and ~80 nM of either of the dsDNA libraries. After incubation at 37 °C overnight, the RNAs were purified by 5% denaturing urea-PAGE with UV shadowing and electroelution followed by isopropanol precipitation and 70% (v/v) ethanol rinse, which yielded 39 nmol and 24 nmol of Unbiased and Biased library RNAs, respectively. All of the RNAs (5 μM) were photo-crosslinked with 1.5 eq. of puromycin-containing oligo XL-PSO in 1X XL buffer by irradiation at 365 nm using a handheld UV lamp for 20 min on ice after an incubation cycle of 70 °C for 3 minutes and slow cooling to 25 °C. The puromycin-modified library RNAs were purified by 5% denaturing urea-PAGE with visualization by staining with RNA Staining Solution (Abnova) and electroelution, yielding 7.8 nmol and 4.8 nmol of puromycin-modified RNAs of Unbiased and Biased libraries, respectively.

*3. Library Translation and mRNA-peptide Fusion Formation*

The radiolabeled mRNA-peptide fusions for round 1 selection were produced as follows: Unbiased and Biased libraries were prepared separately. The components of PURE system with homopropargylglycine and ~3 μM [2,5-$^3$H]-L-histidine (Moravek Biochemicals) were assembled on ice as described in **Section II.A.3** with the inclusion of 100 μM $CoCl_2$, 6.5 μM PDF, and 15 μM MAP. To the PURE system mixture (7.3 mL and 5.0 mL for Unbiased and Biased libraries, respectively), 0.8-1 μM puromycin-modified library RNA for a total of 6.9 nmol and 4.7 nmol of Unbiased and Biased libraries, respectively, were added and incubated at 37 °C for 30 minutes. Next, 0.3 vol. of 2.05 M KCl and 172 mM $Mg(OAc)_2$ were added to facilitate fusion formation, incubated for 15 min at room temperature, and stored at -20 °C overnight.

*4. Library Cyclization and Separation from Free Peptides*

Alternate procedures were used, in which either 1) cyclization was carried out on oligo(dT) magnetic beads before reverse transcription or 2) on Pierce™ Streptavidin UltraLink™ Resin (Thermo Scientific) after reverse transcription. These procedures are described separately in the following two sections. For Round 1 of selection, both strategies for preparing cDNA/mRNA-peptide fusions were employed, but oligo(dT) bead purification was preferred due to higher recovery and more efficient reverse transcription results.

*5. Library Cyclization during Oligo(dT) Purification and Solution Phase Reverse Transcription*

When cyclization was done on oligo(dT) magnetic beads, the procedures were as follows: The amount of oligo(dT) beads used was 5 μg per pmol puromycin-modified RNA. The volume of each buffer used below was 0.8 mL per mg beads (4 x the vol. of the bead slurry from the manufacturer) unless noted otherwise. Oligo $d(T)_{25}$ Magnetic Beads (New England Biolabs) were washed twice with Oligo(dT) Binding Buffer (High EDTA) (20 mM Tris-HCl, pH 8.0, 1 M NaCl, 50 mM EDTA, pH 8.0, 0.2% (v/v) Triton X-100, 5 mM BME). A translation mixture was added together with 1 vol. of 2X Oligo(dT) Binding Buffer (High EDTA), and the mixture was tumbled at room temperature. The beads were separated on a magnetic rack and washed 4 times with Oligo(dT) Binding Buffer TCEP (20 mM Tris-HCl, pH 8.0, 1 M NaCl, 10 mM EDTA, pH 8.0, 0.2% (v/v) Triton X-100, 0.5 mM TCEP), and 3 times with Oligo(dT) Wash Buffer TCEP (20 mM Tris-HCl, pH8, 300 mM NaCl, 10 mM EDTA, pH 8.0, 0.1% (v/v) Tween-20, 0.5 mM TCEP). For cyclization, Cyclization Buffer (20 mM Tris-HCl, pH 8.0, 660 mM NaCl, 0.2% (v/v) Triton X-100, 0.5 mM TCEP, 3.3 mM m-dibromoxylene, 33% (v/v) acetonitrile) were added to the beads and tumbled at room temperature in the dark. Next, the buffer was switched to Cyclization Wash Buffer (20 mM Tris-HCl, pH 8.0, 660 mM NaCl, 0.2% (v/v) Triton X-100, 10 mM BME, 33% (v/v) acetonitrile) and tumbled at room temperature for 10 minutes to quench the reaction. Next, the beads were washed twice with Oligo(dT) Wash Buffer BME, no EDTA (20 mM Tris-HCl, pH8, 300 mM NaCl, pH 8.0, 0.1% (v/v) Tween-20, 10 mM BME), and the cyclic-peptide mRNA fusions were eluted by 0.1% (v/v) Tween-20 (80 μL per mg beads) 5 times at room temperature and 3 times with heating at 70 °C for 2 minutes. The solution was passed through 0.22 μm Ultrafree-MC Centrifugal Filter(s) (MilliporeSigma) with centrifugation at 14,000 ×g.

Reverse transcription was done as follows: The filtered solutions were divided into aliquots (~190 pmol/tube of a mixture of free RNA and RNA fused to peptides), and the RNA and mRNA-peptide fusions were precipitated by isopropanol precipitation with 10 μg/microcentrifuge tube of Linear Acrylamide carrier (Invitrogen) and rinsed with 70% (v/v) ethanol. To each of the pellets containing RNA and mRNA-peptide fusions, 31.25 μL of 16 μM RT primer in 0.16% (v/v) Triton X-100 were added and incubated on ice for ~30 minutes, heated at 65 °C for 5 minutes, and chilled on ice. Next, 10 μL of 5X Reaction Buffer (Thermo Scientific), 5 μL of 10 mM of each dNTP, 1.25 μL of 40 U/µL Ribolock RNase Inhibitor (Thermo Scientific), and 2.5 µL of 200 U/µL RevertAid H minus Reverse Transcriptase (Thermo Scientific) and incubated at 42 °C for 30 minutes.

*6. Library Cyclization and Reverse Transcription on Streptavidin Resin*

A portion (1440 µL) of the Unbiased library was purified, cyclized, and reverse transcribed using Pierce™ Streptavidin UltraLink™ Resin (SAUR; Thermo Scientific) as follows: Unless otherwise stated, all washes were performed with buffer equal to 5 vol. of the 50% slurry resin added to each 1.5 mL tube. For each of 4 aliquots of 360 µL of translation mixture, 90 µL (0.25 vol.) of the resin was washed 3 times with SAUR Washing Buffer 1 (SWB1 Buffer; 50 mM Tris-HCl, pH 8.0, 150 mM KCl, 0.2% (v/v) Triton X-100, 7 mM BME) and then resuspended in 450 µL (5 resin vol.) of a fusion/primer mix comprising 360 µL translation mixture, 50 nM Tris-HCl, pH 8.0, 60 mM EDTA, pH 8.0, 0.2% (v/v) Triton X-100, and 1 µM PCB (photocleavable biotin)-18-RT-primer1. After tumbling for 30 minutes at room temperature, resins were spun down and washed twice with SWB1 Buffer (second wash 2 mL, the maximum capacity of the microcentrifuge tube). To perform the reverse transcription, the filtered resins were heated to 65 °C for 10 minutes (without adding buffer) and then chilled on ice for at least 5 minutes. Resins were resuspended in 135 µL (1.5 resin vol.) of a reverse transcription reaction mix (1X Reaction Buffer (Thermo Scientific), 1 mM of each dNTP, 1 U/uL RiboLock RNase Inhibitor (Thermo Scientific), 10 U/uL RevertAid H minus Reverse Transcriptase (Thermo Scientific)) followed by incubation at 42 °C for 1 hour with occasional mixing. For cyclization, resins were washed 5 times with Cyclization Wash Buffer with TCEP (50 mM Tris-HCl, pH 8.0, 300 mM NaCl, 0.2% (v/v) Triton X-100, 0.5 mM TCEP-HCl), resuspended in SAUR Cyclization Buffer (50 mM Tris-HCl, pH 8.0, 300 mM NaCl, 0.2% (v/v) Triton X-100, 0.5 mM TCEP, 5 mM m-dibromoxylene, 20% (v/v) acetonitrile), and tumbled for 30 minutes at room temperature while covered in aluminum foil. The resins were washed with Cyclization Wash Buffer with BME (50 mM Tris-HCl, pH 8.0, 300 mM NaCl, 0.2% (v/v) Triton X-100, 10 mM BME, 20% (v/v) acetonitrile) and tumbled for 10 minutes at room temperature. To prepare for click reaction, resins were washed twice with 0.2% (v/v) Triton X-100 and resuspended in 180 µL (2 resin vol.) 0.2% (v/v) Triton X-100. To photo-elute mRNA-peptide fusions from the resin, the tubes were covered on the side with aluminum foil and irradiated from the top by a 365 nm handheld UV lamp for 20 min. The supernatant and an equal volume rinse were collected before repeating the irradiation for another 10 min followed by another supernatant collection and rinse. Since SDS-PAGE suggested that libraries were not entirely reverse transcribed, reverse transcription was repeated in the solution phase after isopropanol precipitation as described in **Section II.A.5**. However, the second reverse transcription still failed to complete the generation of cDNA/mRNA-peptide fusion libraries.

*7. Ni-NTA Agarose Purification*

Since the mixture contains cDNA/mRNA duplexes that were not fused to peptide, cDNA/mRNA-peptide fusions were purified by immobilized metal affinity chromatography (IMAC) as follows: The cDNA/mRNA and cDNA/mRNA-peptide fusions were precipitated by ethanol precipitation in the same tubes as the reverse transcription reaction and redissolved in 80 µL of Denaturing Bind Buffer (100 mM $NaH_2PO_4$, 10 mM Tris, 6 M guanidinium hydrochloride, NaOH to pH 8.0, 0.2% (v/v) Triton X-100, 5 mM BME). 20 µL of 50% slurry HisPur™ Ni-NTA Resin were used per tube. The resins were equilibrated and resuspended in 80 µL Denaturing Bind Buffer and then added to each of the tubes individually. After incubation at room temperature with tumbling for 1 hour, ~500 µL of suspensions were combined into a 0.22 µm Ultrafree-MC Centrifugal Filter (MilliporeSigma) and spun down at 2,000 ×g. In further ~500 µL increments, the remaining suspension volume was combined into the same centrifugal filter unit and spun down until all the resin had been transferred to the filter. The resins were washed 2 times with 8 resin vol. of Denaturing Bind Buffer and 3 times with 8 resin vol. of Native Wash Buffer (100 mM $NaH_2PO_4$, 300mM NaCl, NaOH to pH 8.0, 0.2% (v/v) Triton X-100, 5 mM BME) and eluted multiple times with 1 resin vol. of Native Elute Buffer (50 mM $NaH_2PO_4$, 300 mM NaCl, 250 mM imidazole, NaOH to pH 8.0 (stored at 4°C), with 0.2% (v/v) Triton X-100, 5 mM BME). The unbound fractions were re-subjected to IMAC up to 4 times to maximize recovery. The eluate fractions that contained the highest recovery according to LSC were combined and buffer-exchanged using NAP5 columns to Gel Filtration Buffer (10 mM Tris-HCl, pH 7.5, 1 mM EDTA, pH 8.0, 5 mM BME, 0.2% (v/v) Triton X-100) to remove imidazole.

*8. Biotin-primer Purification*

Because reverse transcription did not go to completion using either strategy and because resubjection of the first batch library to reverse transcription did not result in more complete generation of cDNA/mRNA-peptide libraries, an additional purification step was performed with immobilized PCB-18-RT-primers to remove non-reverse transcribed libraries that could be captured by the primers. After gel filtration, fusions were precipitated by isopropanol precipitation with 2.5 µg/microcentrifuge tube of Linear Acrylamide carrier (Invitrogen) and rinsed with 70% (v/v) ethanol. Each pellet was redissolved in 5 µL of a primer solution (0.1% (v/v) Triton X-100, 5 µM PCB-18-RT-primers (PCB-18-RT-primer1 for the Unbiased library; PCB-18-RT-primer2 for the Biased library)). 10 µL of the streptavidin resins were washed 3 times with 50 µL of SAUR Washing Buffer 1 with BME and EDTA (SWB1E; 50 mM Tris-HCl, pH 8.0, 10 mM EDTA, pH 8.0, 150 mM KCl, 0.2% (v/v) Triton X-100, 7 mM BME) and transferred to the fusion/primer solution using 20 µL buffer such that the final 25 µL has composition: 50 mM Tris-HCl, pH 8.0, 10 mM EDTA, pH 8.0, 150 mM KCl, 0.2% (v/v) Triton X-100, 5 mM BME. After 30 minutes of tumbling at room temperature, all suspensions were collected into a single 0.22 µm Ultrafree-MC Centrifugal Filter (MilliporeSigma) and spun down at 5,000 ×g for 1 min. The tubes and resins were washed 7 times with 12.5 µL SWB1E, and all flow-throughs were collected. SDS-PAGE analysis indicated that all bands other than the desired cDNA/mRNA-peptide fusion band were removed.

*9. Click Glycosylation*

As with the PGT128 selection, several batches of fusions were prepared for the Round 1 library. Each batch was subjected to click glycosylation separately, and a summary of all batches of cDNA/mRNA-peptides prior to click reaction (just after PCB-18-RT-primer purification) is given in **Table S13**.

**Table S13**. Summary of cDNA/mRNA-peptides quantities after PCB-18-RT-primer purification as estimated by LSC.

| Batch | Unbiased Library (pmol) | Biased Library (pmol) |
|---|---|---|
| 1 | 7.54 | 10.05 |
| 2 | 21.7 | 7.51 |
| 3 | 29.8 | 19.6* |
| 4 | 123.5* | - |
| Sum: | 182.5 | 37.16 |

* These batches did not require PCB-18-RT-primer purification.

Each batch was subjected to isopropanol precipitation and and the pellets were rinsed with 70% (v/v) ethanol. Pellets were redissolved in 2-4 µL of 250 mM HEPES-KOH, pH 7.6, 0.075% (v/v) Triton X-100 depending on the volume of the click reaction. 1 µL of 50 mM aminoguanidine was added per 10 µL click reaction, and reactions were carried out as described in **Section II.A.8** with scaling of all volumes for reactions carried out at 5, 8, or 10 µL. All batches were subjected to isopropanol precipitation after the first click reaction and the pellets rinsed with 70% (v/v) ethanol before resubjection to another click reaction to improve glycosylation efficiency. A small portion of reaction products were digested with Nuclease P1 (see **Section II.A.9**) and analyzed by SDS-PAGE to monitor click efficiency and library multivalency.

**B. Round 1 Selection**

All batches were buffer exchanged by isopropanol precipitation with 5 µg/microcentrifuge tube of Linear Acrylamide carrier (Invitrogen) followed by 70% (v/v) ethanol rise and redissolved in 1X selection buffer (20 mM Tris-HCl, pH 7.5, 100 mM NaCl, 0.1% (v/v) Triton X-100). After remeasuring specific activity from saved crude translation mixtures, yields were calculated to be 208.7 and 49.8 pmol, equivalent to $1.3 \times 10^{14}$ and $3.0 \times 10^{13}$ sequences for Unbiased and Biased libraries, respectively.

For negative selection against the immobilization carrier, 3.75 and 3 mg Dynabeads™ Protein G (125 and 100 µL of a 30 mg/mL stock; Invitrogen) were equilibrated by prewashing 3 times with 8 bead volumes of 1X selection buffer and resuspended in pooled solutions of Unbiased and Biased library batches to final volumes of 250 and 200 µL, respectively. Libraries were tumbled for 1 hour at room temperature followed by recovery of the supernatant and washing of the beads 4 times with 0.25 vol. 1X selection buffer. To preincubate precleared libraries with target antibodies, the supernatant and washes were combined and mixed with PGT122 and gl-PGT121 such that both antibodies had a concentration of 200 nM in 1200 and 600 µL solutions for Unbiased and Biased libraries, respectively. Libraries were incubated at room temperature for 1.5 hours without tumbling and then 30 minutes with tumbling. Note: This incubation period was reduced in subsequent rounds to 1 hour at room temperature with tumbling. During this incubation, 18 and 9 mg of fresh Dynabeads™ Protein G (600 and 300 µL of a 30 mg/mL stock; Invitrogen) were equilibrated by prewashing with a total of 24 vol. 1X selection buffer over multiple washes. To capture antibodies on the prewashed beads, beads were resuspended in the library/antibody solutions and incubated for 30 minutes at room temperature with tumbling. The beads were magnetically isolated to remove unbound supernatant and washed 3 times with volume equal to the supernatant. Beads were resuspended in 1200 and 600 µL PCR mix A (1X Standard Taq Buffer, 0.1 mM each dNTP, 0.1% (v/v) Triton X-100) for Unbiased and Biased libraries, respectively, and fraction bound values were determined equivalently to the PGT128 selection. The fraction bound was 0.64% for the Unbiased library and 2.79% for the Biased library.

DNA libraries were recovered by on-bead PCR of bound fusion cDNA. For all pilot and large-scale experiments, 1 part resuspended beads in PCR mix A was mixed with 2 parts PCR mix A and 1 part PCR mix B (1X Standard Taq Buffer, 0.1 mM each dNTP, 4 µM forward primer (Library FP), 4 µM reverse primer (Library RP1.3 for the Unbiased library; Library RP2.1 for the Biased library), 0.1 U/µL Taq (New England Biolabs), 0.1% (v/v) Triton X-100). Small-scale pilot experiments as described for the PGT128 selection were used to establish the optimal number of thermal cycles and annealing temperature. Large-scale PCR was performed with an initial incubation at 94 °C for 5 minutes (with brief vortexing before and after incubation to resuspend beads), 18 or 24 cycles (for Unbiased and Biased libraries, respectively) of 94 °C for 30 seconds, 73 °C for 30 seconds, and 74 °C for 30 seconds. DNA was recovered by removing the supernatant from magnetically isolated beads and then washing the beads twice with 3-10 uL 0.1% (v/v) Triton X-100. To purify recovered DNA libraries, solutions were filtered by 0.22 µm Ultrafree-MC Centrifugal Filter (MilliporeSigma) to remove any remaining beads, extracted with phenol/chloroform and then chloroform, and subjected to isopropanol precipitation followed by 70% (v/v) ethanol rinse. Yields were estimated by comparison to samples with known concentration on 8% native PAGE and determined to be approximately 64 and 47 pmol for Unbiased and Biased libraries, respectively.

**C. Subsequent Rounds of Selection**

cDNA/mRNA-glycopeptide fusions were prepared and selected in subsequent rounds similarly as for round 1, but with several changes. Changing condition and selection progress are tracked in **Figures 3 and S35** for PGT122 and gl-PGT121 selections, respectively.

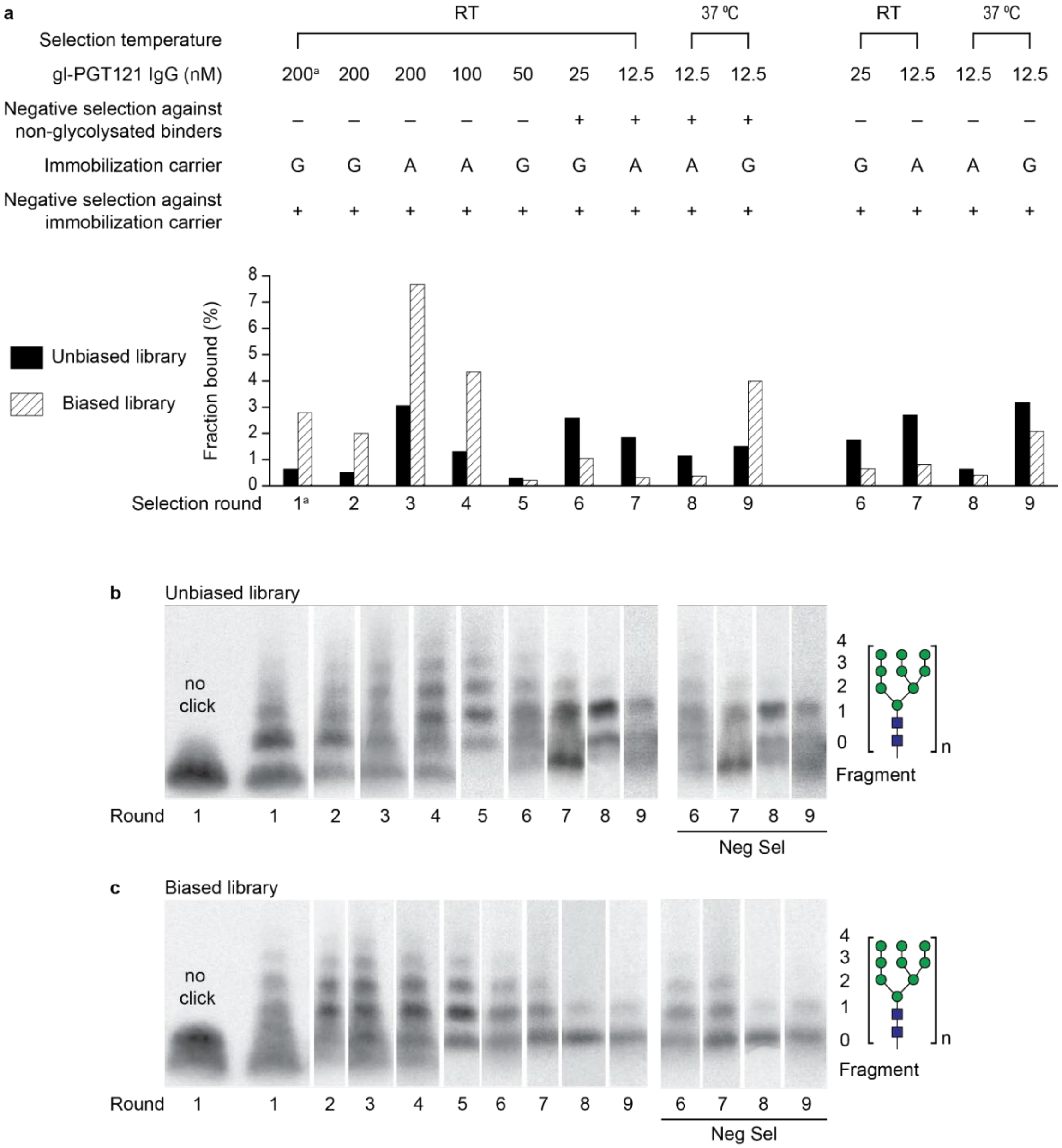

**Figure S35**. Summary of gl-PGT121 selection progress. (a) gl-PGT121 selection conditions and fractions bound for each round. The immobilization carrier used was Protein G (G) or Protein A (A); (b) Glycosylation from round to round for the Unbiased library and (c) Biased library.

[a] Round 1 selection was conducted with both PGT122 and gl-PGT122 in one pot.

*1. Preparation of Puromycin-modified RNA*

For rounds 2-4, RNA produced from T7 transcription was purified by MEGAclear™ Kit (Ambion, AM1908) according to manufacturer instructions instead of by gel purification. For round 5 onward, gel purification was again used because splint ligation was used for puromycin attachment (see below), and this reaction appeared to be more efficient with gel purified RNA template.

For rounds 4-8, a puromycin-containing oligo was attached to the RNA template by splint ligation as previously described.[13] Nucleic acid components were first combined and heated to 94 °C for 1 minute before cooling to 4 °C and keeping on ice for 10 minutes. 10X T4 DNA Ligase Reaction Buffer (New England Biolabs), BSA, RNase Inhibitor (New England Biolabs; M0314), and T4 DNA Ligase (New England Biolabs) were added to the mixture for a final volume of 10 µM template RNA, 10 µM of each splint (SPL1 and SPL1N for Unbiased library; SPL2 and SPL2N for Biased library), 12 µM F30P [$(dA)_{21}$(Spacer 9)$_3$ACC-Puromycin, 5'-phosphorylated using Chemical Phosphorylation Reagent II (Glen Research), purchased from Keck], 1X T4 DNA Ligase Reaction Buffer (New England Biolabs), 0.025 mg/mL BSA, 1 U/µL RNase Inhibitor (New England Biolabs; M0314), and 10 U/µL T4 DNA Ligase (New England Biolabs). After incubation for 1 hour at 37 °C, the reaction was quenched with 4-10 µL 0.5 M EDTA, pH 8.0 per 100 µL of ligation reaction. For purification, solid urea was added to the mixture such that the final solution was ~8 M urea. Ligated RNA was then purified by 5% denaturing urea-PAGE followed by electroelution.

Note: For round 4, both ligated and non-ligated RNA bands seen in 5% denaturing urea-PAGE were recovered by electroelution followed by isopropanol precipitation and 70% (v/v) ethanol rinse. Non-ligated RNA was then resubjected to the same splint ligation and gel purification protocol, at which point it was noted that splint ligation was more efficient with the gel purified template RNA. Ligated RNA from both single and double subjections to splint ligation were used in the subsequent translation.

For round 9 onward, RNA was photo-crosslinked as in rounds 1-3, but purified differently. Purification by 5% denaturing urea-PAGE was performed with staining in ethidium bromide (0.5 µg/mL) and electroelution using Bio-Rad Model 422 Electro-Eluter.

<u>*2. Preparation of cDNA/mRNA-cyclic Peptide Fusions*</u>
For round 3 onward, purification, cyclization, and reverse transcription were conducted only with streptavidin Ultralink resin. Recovered library fusions were able to progress directly from photo-elution from streptavidin resin to click glycosylation. With decreasing library diversity and improved yields, desired yields could be achieved with a single batch each round.

<u>*3. Negative and Positive Selections*</u>
Similar to the PGT128 selection, round 2 libraries were split into selections against only either PGT122 or gl-PGT121 that were conducted in parallel in subsequent rounds. The volume of each round of selection was adjusted to ensure an excess of antibody over library fusions. When selections were performed at 37 °C, both preincubation of libraries with target antibodies and capture onto beads were performed in an incubator at 37 °C with tumbling.

For rounds 6-8 (and round 9 for gl-PGT121), a negative selection was conducted against non-glycosylated binders as follows: After reverse transcription on-resin, photo-elution from streptavidin beads and passage through a centrifugal filter unit, cDNA/mRNA-peptide fusions were concentrated and buffer exchanged by isopropanol precipitation followed by 70% (v/v) ethanol wash and resuspension in 1X selection buffer. The fusions were then subjected to a preclear with the immobilization carrier and selection with 200 nM antibody target as described for round 1 positive selection, except the *unbound* fraction was recovered for subsequent click reaction. Unbound fusions were filtered by 0.22 µm Ultrafree-MC Centrifugal Filter (MilliporeSigma) and prepared for click reaction as described in **Section IV.A.9**. For gl-PGT121 selection, libraries that were never subjected to negative selection against non-glycosylated binders were run in a parallel selection.

For round 9, a negative selection against heat-denatured IgG was performed because high fraction bound values were observed in a pilot selection using a small portion of libraries testing for binding to heat-denatured antibodies. 1600 nM antibody solutions were heat denatured by heating at 70 °C for 20 minutes, chilling on ice for 10 minutes, spinning down at 21,000 ×g for 30 seconds, and resuspending by pipetting. cDNA/mRNA-glycopeptide libraries were incubated with 200 nM respective heat-denatured antibody at room temperature for 1 hour followed by immobilization by incubation at room temperature for 30 minutes with prewashed Dynabeads™ Protein G. The supernatant was recovered from magnetically isolated beads. This negative selection was repeated for a total of 3 times because fraction bound values remained between 5-15%.

For round 9 onward for the PGT122 selection only, positive selection was performed with biotinylated Fabs as follows: For negative selection against the immobilization carrier, 0.25 mg Dynabeads™ Streptavidin M-280 (Invitrogen; 25 µL of 10 mg/mL stock) were equilibrated by prewashing 3 times with 200 µL 1X selection buffer and resuspended in 100 µL Blocking Buffer (20 mM Tris-HCl, pH 7.5, 100 mM NaCl, 0.1% (v/v) Triton X-100, 0.5 mg/mL biotin). After incubation at room temperature for 1 hour with tumbling, the beads were magnetically isolated to remove the supernatant and washed 4 times with 100 µL 1X selection buffer. Libraries were tumbled with these pre-blocked beads at room temperature for 1 hour. To prepare for selection with Fab, another batch of prewashed beads was tumbled with 100 µL of 125 nM biotinylated Fab at room temperature for 1 hour and then washed 4 times with 100 µL 1X selection buffer. Since we determined that ~40% of biotinylated material bound these streptavidin beads, we estimated a final concentration of ~50 nM biotinylated Fab in 100 µL resuspended beads. Beads were then resuspended in 100 µL Blocking Buffer and tumbled for 30 minutes at room temperature followed by 4 washes with 100 µL 1X selection buffer. Fab-bound beads were then resuspended with precleared libraries and tumbled at room temperature for 1 hour. After separating from the unbound supernatant and washing 6 times with 200 µL 1X selection buffer, beads were resuspended in 25 µL 1X selection buffer, heated to 70 °C for 20 minutes, chilled on ice for 5 minutes, and tumbled at room temperature for 10 minutes. At this point, heat eluted selection winners were collected by saving the supernatant and two 25 µL 1X selection buffer washes of the beads.

For round 12 of PGT122 selection, we wanted to improve the proportion of library containing at least one glycosylation site. To achieve this, the click reaction was performed with 5 mM Biotin-PEG$_{11}$-azide (Alfa Aesar) instead of Man$_9$(GlcNAc)$_2$-azide, and the selection step was performed with Dynabeads™ Streptavidin M-280 (Invitrogen). Before selection, fusions were purified by Ni-NTA purification and gel filtration as described in **Section IV.A.7** with minor protocol modifications: Denaturing bind buffer was prepared with 0.5 mM TCEP instead of 5 mM BME, and Ni-NTA suspensions were transferred to a Mini Bio-Spin® Chromatography Column (Bio-Rad) for draining flow-through and washing. The selection step was performed as follows: 0.25 mg Dynabeads™ Streptavidin M-280 (Invitrogen; 25 µL of 10 mg/mL stock) were equilibrated by prewashing 3 times with 200 µL 1X selection buffer with 5 mM BME (20 mM Tris-HCl, pH 7.5, 100 mM NaCl, 0.1% (v/v) Triton X-100, 5 mM BME) and resuspended in 250 µL of libraries. After tumbling at room temperature for 1 hour, the beads were separated from the supernatant by magnetic isolation and washed 6 times with 250 µL 1X selection with 5 mM BME. The beads were resuspended in 100 µL 1X PCR buffer (NEB) + 0.1% (v/v) Triton X-100 for on-bead PCR. On-bead PCR was performed as described above but with 36-fold dilution of beads, 67 °C annealing temperature, and only 3 or 4 PCR cycles for Unbiased and Biased libraries, respectively.

For rounds 14 and 15 of PGT122 selection, a few changes were made to the selection step. After the preclear with pre-blocked beads, the supernatant containing unbound library fusions was passed through a 0.22 µm Ultrafree-MC Centrifugal Filter (MilliporeSigma) to ensure the removal of fusions bound to pre-blocked beads. The collected preclear fusions were mixed with a BSA and salmon sperm DNA solution for a final mixture of fusions in 1X selection buffer with BSA and salmon sperm DNA (20 mM Tris-HCl, pH 7.5, 100 mM NaCl, 0.1% (v/v) Triton X-100, 1 mg/mL UltraPure BSA (Ambion; AM2616), 0.1 mg/mL Salmon Sperm DNA, sheared (Ambion)). After tumbling at 37 °C for 1 hour, beads were separated from the supernatant by magnetic isolation followed by 3 washes with 500 µL pre-warmed 1X selection buffer with BSA and salmon sperm DNA, 3 washes with 500 µL pre-warmed 1X selection buffer, and heat elution as described above.

### D. Sequencing

#### 1. Sanger Sequencing

Recovered fraction bound DNA from round 7 of PGT122 and gl-PGT121 selections was prepared for Sanger sequencing. DNA (~80 ng) was cloned into plasmids and transformed into competent cells following TOPO® TA Cloning® Kit instructions. Individual clones were grown, and their plasmids were purified by miniprep kit (Zymo). Plasmids were quantitated by NanoDrop, assessed for purity by 2% agarose gel, and then sent to GENEWIZ for Sanger sequencing.

#### 2. Preparation of DNA for NGS

For all rounds after round 7 of PGT122 and gl-PGT121 selections, recovered library DNA was prepared for NGS. Either 300 µL of crude on-bead PCR product or 15 µL of DNA purified by phenol/chloroform extraction and isopropanol precipitation were purified using a PCR clean-up kit (New England Biolabs) according to manufacturer instructions, eluting with 16 µL of 10 mM Tris-HCl, pH 8.0. The purified DNA was quantified by NanoDrop and assessed for purity by 8% native PAGE. DNA samples were submitted to GENEWIZ for Amplicon-EZ NGS.

*3. Clustering and Analysis*

As was done for the PGT128 selection, we used NGS data to track multivalency trends over multiple rounds of selection for PGT122 and gl-PGT121 selections (**Figure S36**). Unlike for the PGT128 selection, libraries recovered from round 8 of selection demonstrated predominantly 0 and 1 glycosylation sites, with an average number of 0.7 and 0.3 glycosylation sites for PGT122 Unbiased and Biased libraries, respectively, and ~0.6 glycosylation sites for all gl-PGT121 libraries tested. However, by designing round 12 of PGT122 selection to recover only sequences that successfully underwent click reaction, the multivalency distribution shifted toward more glycosylation sites per sequence after round 12 of PGT122 selection, with the average number of glycosylation sites shifting to 1.4 and 1.2 for the Unbiased and Biased libraries, respectively. While multivalency remained higher in rounds 13 through 15 compared to round 8 values, the Biased library in particular started trending again toward lower multivalency. These trends are again consistent with the glycosylation patterns observed by SDS-PAGE (**Figures 8 and S35B-C**).

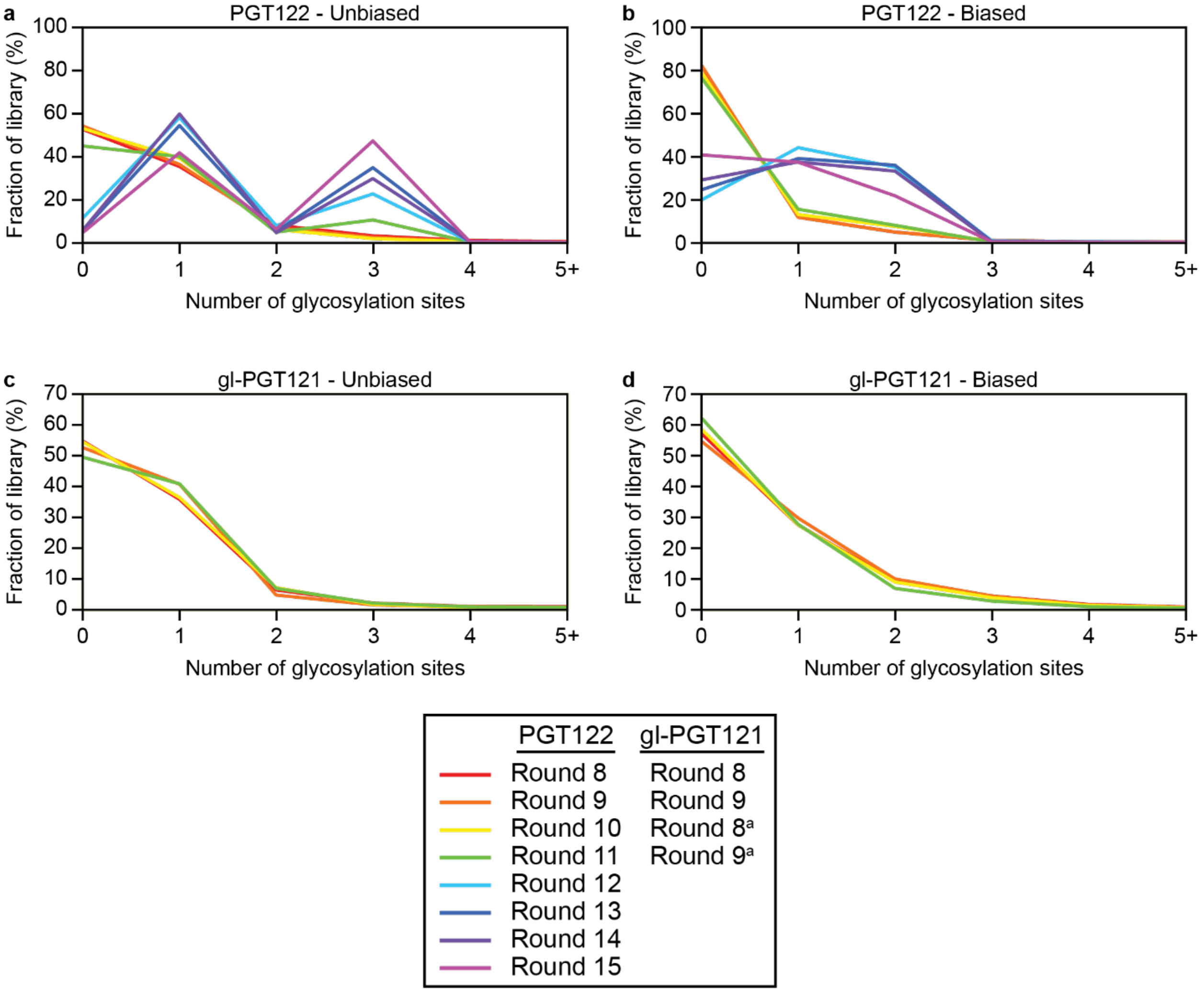

**Figure S36**. NGS analysis of multivalency for (a) PGT122 Unbiased library, (b) PGT122 Biased library, (c) gl-PGT121 Unbiased library, and (d) gl-PGT121 Biased library.

[a] These rounds were performed without negative selection against non-glycosylated binders.

We also noticed that many sequences had frameshift mutations. These mutations are particularly prevalent in the PGT122 Biased library before round 12, where a large majority of sequences had a length other than 153 base pairs in the randomized portion of the ORF (**Figure S37**). Since the majority of the library had 2 and 4 deletions and randomized codons were encoded by NNS, these frameshift mutations had a high likelihood of producing sequences without a glycosylation site. Round 12 of the PGT122 selection strongly favored sequences with the desired length, but the distribution started trending away again in subsequent rounds. While we observed the same pattern in the gl-PGT121 Biased library, the Unbiased library of both selections did not experience a comparable level of frameshift mutations (**Figure S37A**).

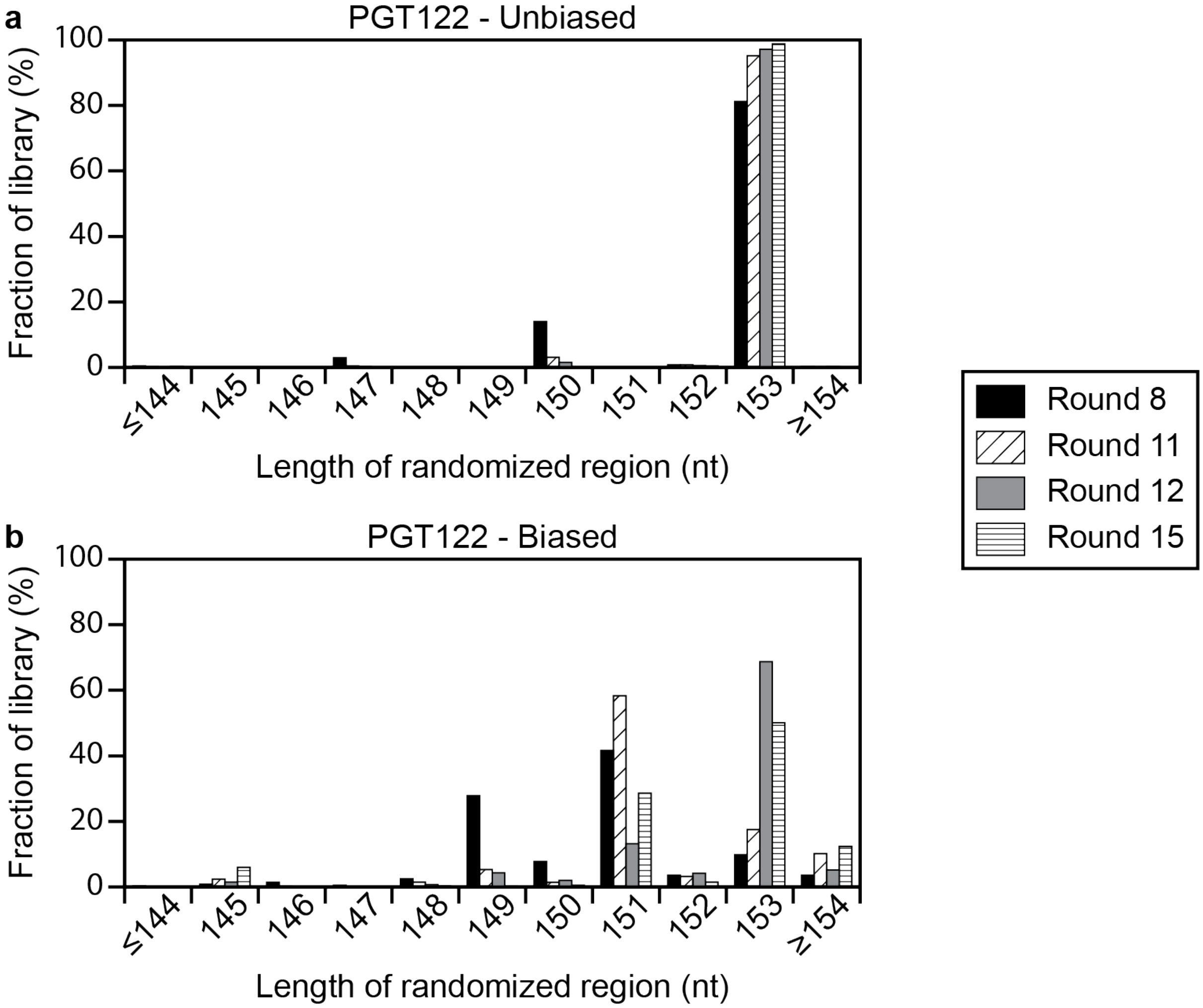

**Figure S37**. Distribution of lengths of randomized region observed in sequences returned by NGS for PGT122 selection (a) Unbiased library and (b) Biased library.

Sequences were clustered in the same way as for PGT128 selection, but with minimum identity threshold of 0.765 (maximum 12 amino acid mismatch allowance) for PGT122 selection and 0.843 (maximum 8 amino acid mismatch allowance) for gl-PGT121 selection. The top 20 clusters are summarized in **Tables S14-S19** with consensus sequence and strength score. Libraries had a greater level of convergence by round 8 than PGT128 selection libraries, with the top 20 clusters representing over 90% of sequences in both libraries by round 15 of PGT122 selection (**Figure S38**).

**Table S14**. Top 20 Most abundant sequence clusters in round 15 of PGT122 selection for the Unbiased library.

| PGT122 – Unbiased, Round 15 (with glycan) | | |
|---|---|---|
| ID | Consensus sequence and score | % of all sequences |
| 0 | KTRTPMNLSGKSNIRYFFTIDNSTMFEDLFMRFPRSKEYHIELEEFEDLIR<br>ededeebeee0eddeeededebdc98eeedaeceedceebeeeeee0eeee | 51.06 |
| 1 | QYIDWEIDINQVPNQMNLSKPTWKSDITINKVKWSFYININQASNENTDNI<br>dee8eeeedcededddcdcdded9ededcbddeeddd0cad9dba29dc5 | 24.86 |
| 4 | KTKTYLIPFHLDILPNDNLRSQKIRLIFGDKATVEGMITFLNDSELEWNFE<br>ddddddd0ebeedeebd7c9ddcdee8dde9bcc0e7cdbb9d7dcdebdd | 6.00 |
| 5 | IQNLNHELTMDSPYWINWSIDVNQVPGTTGDWHVTLQYSDINAISDGVTSI<br>ddddd5dddbcdeeeedeeebede9eeeddceaeeedeeddeac8bdd5ad | 2.48 |
| 10 | KYNNQGSNNVEINPFYNNHTNNTYTFTIKVYNTNNKPSMINYNYLVNLTLI<br>ddcdddd0cdddbe2dbabcdbdedddcaed0ca2acc766caedddeeec | 2.31 |
| 12 | YTKLYLLKYIINRLVNRWRNYARCLICVNSKPLIDPMSSSYHNKIIKLCNK<br>deddeeecdda5decccddbdccbc8c8dabdccdedbccebcbaacc9bb | 1.43 |
| 7 | IIIRRLLRLLISTLVNNFTDLLKRRIRRSAMNKTAMSDALPTNSNMNRLFS<br>bededeeeeedebe0cac1aeebb9c9bdd2ccccbabadedcd7bbdc3b | 1.10 |
| 9 | LYFRIDIVELPYETNKIDNTSPITNFTIPKYLIRFGTETNLNSNRNMFVRN<br>ddc0eeceeeeedcccacbbddbecc0cedeeeedadda96ceb898cd6a | 0.93 |
| 17 | FLLSNTDSFATPTIYDFVVDIAERPGTLDVYECRLSIFRRTKSPSKENATT<br>80ecddecccdd9deeeddeeceeeee0a8edbeecdbbea7a5b9c7ada | 0.93 |
| 2 | YSIHFVCDINETPTNTSKAIFSIKCDITTPESTTRQLMITKHYHFPPTGTR<br>ecdbddeeedededdedddaccdeeccccfebcdbdeddecce9ccb92a7 | 0.86 |
| 36 | INHFSPNYRSSLFLGSIVHMSNDPRKWRTFTLTLNEIPGTFPPAIITNVAI<br>cdc06dcddc92dc80d98d9ceeddddebeedddeeee8eeeddd8acdd | 0.50 |
| 37 | YVISFSIDQPDLDRPNIIRISIGDIRSAKCMKLDTINFAYRDDSSEVCYEK<br>cdc9e0efeefeeefadddeeddeedddbb7adee6ccebbec7cbaccd9 | 0.35 |
| 6 | NTKTQTRSTLKKFHLKGLSNQELPTKYLVFNCNIISATEIICSFEPIASAT<br>eaddcce0cbd965facdebddddbbeda08e6ebecdeeceedceebcda | 0.33 |
| 66 | YTKPTTFIIQLDSVPSDNGLMNPAIWLLTISYPNGTNIASYQIAGSVNRKF<br>eececcdeeeeeeddfd9ddc5bdd8fdeeddcfdeb9deeeddbeacccbc | 0.30 |
| 48 | LKNVKNLIVRIILTIINILTNRIANKLSANESRIAIRKRETNKNMLPNITK<br>e6dddcfdacdfededbbf7cdfda9fbbbadaddcdbcda09bdddbc8a | 0.30 |
| 19 | INTHSSIRTLYNVWYESTDDTNSFNLFIEPTKMVNDDIRPKLWGRVTFIQN<br>bec9c6e3cffcedbe4ddd3dede7fede9d4bcd196bbceeeafd0da | 0.22 |
| 13 | HYKETMEKRTNNSVTNNNRYIIELNIIFVDANQNIKKIPIQLEIVIDLAKH<br>edcddedaeccdbdcbbddb97de6c88dcece88b9bf8eeddcae8e0a | 0.21 |
| 45 | FFVRQLLHRFTANAWNLVYWNTAAIYYLNKVLSSALIRSKQNIETSQNISM<br>a0cbeefce5acccdbfced5e0feceed9debccfbdccdcdfdbeecef | 0.20 |
| 28 | RIFKSLNARGLVKLVNSVISSPNNAKFRYRTRLWTYLKNRPLLFRPLISWT<br>ddd8ce0ccefb8ddddebdc7adc8acfcbdfeaddd8cedfccdfecdf | 0.18 |
| 39 | NNVWSITVTVTQTDPFHPDKLPSKDYNSRPRENVVIFFIANCDDTGTCYID<br>dfbf6cadccedfefcebd0ddd7adcfccbfabceedcfdfeecdacddd | 0.17 |

**Table S15**. Top 20 Most abundant sequence clusters in round 15 of PGT122 selection for the Biased library.

| PGT122 – Biased, Round 15 (with glycan) | | |
|---|---|---|
| ID | Consensus sequence and score | % of all sequences |
| 0 | IIPLYIFNYDNNLADISMVPNIGDIREASCMLTSFVIDQTDSKTTITLFPL<br>4bedddd3ddccd0cdd0acbddec9aa904b67dddeeee0deede9ced | 28.23 |
| 1 | RIYITLADPLGPIELVLDVVDWRYPLRDVHAPVVHLDAAQQQLAHQPGHRG<br>eeeeeede8deedcddeeeeeedddecc5bbdcccdd9bddcbb8aaa983 | 21.17 |
| 6 | YYLNVNYLPTTDSNITHFIVEIGEYPYGELHDLRQPVQHLLQVGGHDLPQV<br>eeeeccebedcddddcaedeedeeeededbddddde6dcddccbdbedcdd | 10.75 |
| 3 | TLNIFSNFSIEPDSFYEIVLPIGDIRIAPCMSSFNRNGSTNNCYHFEINLF<br>d8c9bdcbd9dedbacc8bde9dedddeedcb99cbdddaceeceeeaeb | 8.25 |
| 4 | YVISFSIDQPDLDRPNIIRISIGDIRSAKCMKLDTINFAYRDDSSEVCYEK<br>eeedd9eeeeeeeee9ddededdedcdd990bdddcbdccadd8bdddcdb | 6.07 |
| 12 | NNFICIRNIGDIRSARCMNVSAGRYAPYREFYFDITYDDFNKNVNIVITPV<br>adbbdcabdddcbddddcadcdabdceedeeedeedeee89dcdddddeed | 5.28 |
| 10 | FQFGYDNAPYFTFVIDISNTDHNNSQDTDNPFVLIGDIRSGVHDDVGVHVR<br>dccccdcdeebeeeeedb0bd92cdcccccecdcdeedddc5deda0dcdc | 4.84 |
| 15 | LNLYQIVEFTIDESNDIVQLQIGDIRTADCMSLNKECYNIIFFSWDNHGKN<br>2e5dedeeeeeeeebb7eaeeedebeededded6bedbcac4dddd6b82a | 1.97 |
| 9 | RIYITLADPLGPIELVLDVVDWRYPLRDLHVAEPEQHEVADDLVLLPGDLG<br>eeedeededeeedcddeeededdeeecdcdcdddcdeeeddededededb | 1.78 |
| 8 | YPVYLRLPIGDIRIASCMFGDEPNTWTRDELNDYFYTESYNPQPITITFKC<br>eeeefeeeeeeededeecdcedecccd0cccccccdddac2289aaccc0c | 1.36 |
| 7 | HYPNSINHETSVNHSFVSPLVIDDIRIAGCMNDCYWNNATSYYEIPFELDE<br>dedebddccdcdb9ddcaee8d0ed0ebdfaadee5e0eeeeeedeeeeee | 1.16 |
| 18 | NSYIYVTYQMITPNLAFFTIDIGDIPSGALHAVQPLDELHVARDRAGVHEV<br>ceedcdeedeeeecedeedeedeeeeeeeedcddddeecddcdd4eecdee | 0.93 |
| 17 | FIAYYFDLSQPNLHSPNFTLKIGDIREAQCVSLNSTDELPSSAVDFVIDIQ<br>ddeeeefeefedecdecedeceededeedddefdedeeeedcedeeedede | 0.72 |
| 2 | FPFHSNETIVLYVEDKPDRINIGDIRVANCMFTKKKNSNQLEFFCDLQFMP<br>ddc0dcb1ecee0dda3cbebdcddddedf0eddd98c9dcdecddddc9d | 0.54 |
| 16 | LININFTTNNIRYPTNVIYYTIGDIRHASCMGLDMTKNLINFVIDLYETPN<br>edcc8bc5c9cdddeddbc9ecdecdced720dd6c5bccbdeeddddedec | 0.43 |
| 11 | NNHLNVNTIGDIRSAFCMQQTTPIWFLIDLTQEPKRSISNNNSYILRIQFS<br>dbeded0ec0eebdec29ddccfeeeeedfdefec7cbd76cce9dedcca | 0.37 |
| 171 | IIPLYIFNYDNNLVDISMVPNIGDIREASCMLTSFVIDQTDSKTTITLFPL<br>6adcdbd8dcdca0dddaddddddddd100cdddddddddd4ddfdddddd | 0.32 |
| 41 | YQDPKSKDIEYFDLIIHCYPHIGDIRSATCMVLPMNNTIIPFVLDIRVSSE<br>ebbdadbbcddeedefdfdfcddd8beeedbefc2acdcbfddeedfcdcd | 0.32 |
| 22 | TDTNYITSFVIDWNESTGQWTIGDIRVVKCMDATCPEIEKSLIYDVTVDVW<br>eedeecdeededeceeddeedeeeadadddbec4eceeddddcddeeecc | 0.28 |
| 5 | RIYITLADPLGPIELVLDVVDWRYPLRDVHADEPHELLEVGPRLRLLELPG<br>feedefefdeffeffefffffeeeefece0d00dc00000000000d0000d | 0.24 |

**Table S16**. Top 20 Most abundant sequence clusters in round 9 of gl-PGT121 selection for the Unbiased library (with negative selection against non-glycosylated binders).

| gl-PGT121 – Unbiased, Round 9 (with Negative Selection) | | |
|---|---|---|
| ID | Consensus sequence and score | % of all sequences |
| 1 | NPRIIFFELSIRNNSLKLLRWHDTQGIFNDSMTELSQHAELPRFKFHSNAA<br>eeeeeeeeeeedcddeeeeeeeedeeeeddeddd8ddedddeeeedddbcded | 18.46 |
| 12 | HTKFIDIKNSNSKEYMTLLLLTIRSNQVPGTDTPTIKIGTSIRTLTKSDNI<br>eddddddd9dccdde0deeeeeeeedeeeeededeededcdeeededcddbc | 5.31 |
| 2 | KIAFTFPVIIRAESDTNQWVSKIGSNTNHWFSNWTFRPITPNIYLFTTRVQ<br>dedddeeeeeedddddddeeccdebdbecddeeddedeeedeeeeeeeecded | 3.24 |
| 44 | HDPNNTFNVLVFRISIPFEINPELWVRNPSQVKYSTNKIIPGPRNINVLWT<br>ecdddaddeeeeee8eedeccddeeedddddddecdcddecddccecedea | 2.33 |
| 58 | DINKKYLVQIKLKLKNTKTKKETKNSSHHKSKRKHKQSLKKHKRIWKKKSH<br>edeeeedddeeeeeebdddddddddededdeecddddddedddcededdccd | 2.12 |
| 733 | RTEKKYYERRTLKKLKNNWKEHHKLQKRNSKKKHNFSKKHWKIKIKIKSIK<br>ddeedeedeeeeeeeed5ceddddeddea1eedcddededdeddddeeecddcd | 1.43 |
| 100 | NSMLLTLLLRTNLASPVKVYAFTLRTIDSNEGTSVTSITVHPFLVTQNHHN<br>aebeddeeeeddeededdddedeeeeeeeecceeeeeeeeeee1ededcecc09 | 0.92 |
| 48 | KNLKDRKHKYKLRTNNKYKKKPLKLSSHKLKKKKNKRIRLSKSLKGSKFRI<br>ddededddeddeeeddddddededdeddceedca9aaab99aaa9aabaaa | 0.91 |
| 220 | KLKKKMNSHNKFENKNLKKKKHPKIKGTKSKKKKLRINKIKKKYNWIKLKS<br>eeeec0ccccddbcdcddec0000000000ded0000000cc000000000 | 0.77 |
| 71 | KTYKLDKRWNKVNKKKNTKTKKHLKDNKKSIEKHHVKKMKEKKLKDAANKK<br>eeceeeedecddcddcbdddddddddceddcbcddbdcacbdd9dcdcbdc | 0.77 |
| 7 | KTKNKKLIKKTSHHKLKTRPKPKKPTHKNKSNQTKNHKTNLHKHKKKSKFH<br>eeecddeeedadecddd0deddecdccc9daddbdcdddceddcddcddcc | 0.71 |
| 64 | LKKLKPNNRRVKKKTKPIKTPQKKKYTKRVEKEKNKIKVKVKIKFKFTKKM<br>deeddcdceeedddddddcedededcccdbcbc7cccccbcccddbc0cdc8 | 0.68 |
| 0 | KLWNTKNRAKHPKLTKPKKQNIKKKRLTEKRNKLNKHIRKISRKNLKKSKK<br>edcccddddeddeddeeedccdeddddddabdcddcdcddcddbdbeedcdd | 0.62 |
| 93 | KTYKRIKKRIKFNKNHKNKNSKFILKTLKFSKKNNLKIPNKKIKKIKKYKL<br>feeeeeedeeeeced7e4dddeeded0aeedeeddedededdeeeeeeeee | 0.62 |
| 119 | QQTHLYLIPFYISTNNNNTKLLALTWKNSKEWQLQSMTPSINKIALTMITL<br>deedeeeeeeeeeeecdddddecdeddddcdddadcc3ddecbcdcec0dcd | 0.58 |
| 191 | IIYYPFRITFNQTPNDIENLSGRISFSPSQPLLLIPSVIASNLWRFLMSGS<br>eeeeffeee5eeeeeddeceeeeeeeeeeeddebdddebcadbceeddeeedd | 0.57 |
| 9 | EYKKLKGAKKPGPEKNNKRTQYKKKVKHKRLTKKKWKTSHKRKKQHKKYLI<br>eedeeeddddddddd9cc8dddedcddddddededdeddcddbddddedeec | 0.51 |
| 26 | ITSTPTGLSSHDTYIIKFMVTLNNNNYPRKLKLNFDNSRFSTTPTGFWYTL<br>edddeddeedcdeedcce0deecdcbccccbccabbac9badaccbbcccc | 0.47 |
| 101 | QSHNLKKHSKNKPKKIKLKKKHNVLVKLIAQKNSTEKIKYKKHHIRTKKKK<br>ddedddeeecdeeddeeddcdcdededdbddcbdddccbdddccddedda | 0.46 |
| 78 | KEKKARIKLREKKYINKTKKNHTKKYRKKDKYKNKKKIKYHKKPNDIKKLK<br>eeeeddededcddddaecdeccadddeeedbeeddeddddcdcbccdedcd | 0.45 |

**Table S17**. Top 20 Most abundant sequence clusters in round 9 of gl-PGT121 selection for the Biased library (with negative selection against non-glycosylated binders).

| gl-PGT121 – Biased, Round 9 (with Negative Selection) | | |
|---|---|---|
| ID | Consensus sequence and score | % of all sequences |
| 1 | YPTTHLTSFYVSTNPSTTTIRIGYPSGVLHGHLVPDLVLHLLEQGEPLQLR<br>dededede eededeeddeeeeeeeeedecebececedeeeceeddecedcde | 21.44 |
| 0 | HSLFNFQVRLKQLPLDANGAVIGDIRIAECMNHTYTPLTFYIYSNHPTDIQ<br>ddddeeeeeedeeeeeddeeeeeeeeeeeabccdddeedededdddcdb0bb | 8.46 |
| 32 | HIHNNIDIKDSTRKNAVSFAFIISFNQTPLGDKMIGDIRKAELHDALRDVL<br>ecdcddcccccdcccddeedddeecdedeeedcaeedddddddccdede9ce | 2.35 |
| 2 | HPTWSLLPFLINIRQQQIKYSIGDIHGVLHVVEPLQGEDDQLLVEPLHQVR<br>eeeeeedeeeedeed5deeeeeeecdeeeceddeeeeedddeeddeedced | 1.93 |
| 37 | YPPALDDDWNQFLRFRLNVEDQQVLHIPQNGTWRLAISVSRTDEPEHLHLR<br>edceddddcdddddaecdddeddddddedcddedbceedecdddedddadd | 1.75 |
| 22 | KIKRLKRLEKLQHRKHKKNTSKKRLKNGRHQNSTMIKVSRAQKRVKTERKK<br>eddedeedddecdeccd0ecde8eeddedddbeeccdddeeddedeecdde | 0.96 |
| 15 | TTDFSLPFRLNVDHYGHVSLRRPNYKMTWHDEINIGDIRLARCMFGESTLR<br>edeeeeeedeeeeeeeedeeee0eddc4adedadcdeedeeee67edaaabb | 0.86 |
| 127 | HYFEHLATIGDIRFALHDDQQGVLVVRLRGENHITHPVLLVQLRPVPQVGL<br>ddeecdeeeeeeeeefeedddeeeeeeeeeeeeeddeeeeeeeedeeedeeeee | 0.62 |
| 31 | KLKKKMNSHNKFENKNLKKKKHPKIKGTKSKKKKLRINKIKKKYNWIKLKS<br>ceddb0cdddbeededeeeaba9bbbababdeeaab99aadd9a9aa9a9a | 0.56 |
| 69 | YYLIFVADLNHKGQISRHSEEGRVHLVRLDDHDHWRYPSGGLHDLGPLLEG<br>efeeeedeebebeeeedeedaedacdeeedddbedeeceeedcdeeeeede | 0.52 |
| 23 | FIAYYFDLSQPNLHSPNFTLKIGDIREAQCMLTCDSAKPNPRVFPILALLI<br>defeeddeeeeeedeeddeedceeededdcbcddddeeddded0eeedeee | 0.45 |
| 41 | HSLTNNTHTDVYGSYSCQFYHIGDIRIASCMIHLSNNMWMINHSQDHTLND<br>de9cdcdeedeeeeededeecdedddeeecdcebdbdddedccdeddcce | 0.42 |
| 161 | ITHLNFPLIGDIRWAFLHDRDAPLLELAEAGHVLVPLVVQRDVHGGRLVLA<br>cedddceddecdeeeeeedeeeeefeeeeeedfeeeeeedfeccfedeeef | 0.42 |
| 46 | HSHSPISVLATIKTSNLHNSPTHGLLLIRVSTLDIGDIRADLHVDAHVDEA<br>edcdedceeddddddedccdeddeeeeeddeeddded dddedcdddccdddd | 0.40 |
| 79 | ATSWQLYTLAISVSRVHDVLGEGDPRRDRRLPEVQVRLVLVPLEGVDQLVG<br>eedeeddcbefeeeebadddeddddeeeeedecdbeedeeeeeedede dcdede | 0.34 |
| 35 | TNARRAHIIGDIRSVLHAPHAVRPDDQRQGDPLQHVRGVHFLTIKIKQAPG<br>ededeeddeeedeededdeeedeebddcceeadedefeddeededdafeee | 0.34 |
| 188 | NWPNIYFVKTYVPATDVNNFYIGDIRHLHEVVLPDQVRQDLVPVHLLVLGL<br>eeddeeceecedeeedcddcdeeeeeeeecedeedeededeeeeedeeeedd | 0.31 |
| 255 | TLIFVVHQNTIPGLPHEMRNLRVELLLPDRQVAHWRYPFGGVHDRHPVLDG<br>eedefededdeefe7debdefeeefeeece2eeeeeeddeedddeeeedcd | 0.26 |
| 83 | ISQWSVYRLAISVADLHAAEGDRVRVVVRRVDDPQPDELLDRLYGFNLRIG<br>ddffdafeedeeeeddeeddecfeeeededeebdfdceddedfebddeedd | 0.25 |
| 639 | HNTNDILVLNFVYTQNLGQHKLLVYHSTHEVDDLIGDIRTANDDHPLQHAR<br>eccecdedeeeeededdedeeefeebeddddedOecceccdecd0dddcdbb | 0.24 |

**Table S18**. Top 20 Most abundant sequence clusters in round 9 of gl-PGT121 selection for the Unbiased library (without negative selection against non-glycosylated binders).

| gl-PGT121 – Unbiased, Round 9 (without Negative Selection) | | |
|---|---|---|
| ID | Consensus sequence and score | % of all sequences |
| 0 | NPRIIFFELSIRNNSLKLLRWHDTQGIFNDSMTELSQHAELPRFKFHSNAA<br>eeeeeeeeeedccdeedeeeedeeeddeddd7ddedddeeeeddddccded | 20.83 |
| 7 | HDPNNTFNVLVFRISIPFEINPELWVRNPSQVKYSTNKIIPGPRNINVLWT<br>eddddaddeeeede7eeddcceeeeeddedddcecddcddcdddcddedec | 11.73 |
| 5 | KIAFTFPVIIRAESDTNQWVSKIGSNTNHWFSNWTFRPITPNIYLFTTRVQ<br>eeddddeeeeedddddeecbddbdbecdddeddedeeddeeeeeeedded | 6.92 |
| 36 | HTKFIDIKNSNSKEYMTLLLLTIRSNQVPGTDTPTIKIGTSIRTLTKSDNI<br>eddddddd9dcddce0deeeeeeeceeeeecddeedddcdededeecddcc | 4.93 |
| 25 | NSMLLTLLLRTNLASPVKVYAFTLRTIDSNEGTSVTSITVHPFLVTQNHHN<br>aeaedddeeeeddededddeeeeeeededceedeedeeee0eddececd09 | 1.46 |
| 10 | LNMPIGLTWTPITFYITYKQIPYTNRTSLIANWELIENDPIQMRFPKSRKN<br>ee8eddedeeeedddedeeeeeeeeedddedbaedddccddcd2dcccdbbb | 1.30 |
| 20 | QQTHLYLIPFYISTNNNNTKLLALTWKNSKEWQLQSMTPSINKIALTMITL<br>eeedeeeedeedeeccdccceccdcdbbcccc9bbb0ccccaccbcc0cbc | 1.19 |
| 18 | DNNPKVLWFVTSTIQNDNNFVNKTPKYAINWITFRPDRLTTIETQTFTITF<br>eededeeedecddddccdbecddbebedcdddcddeeccdeddd1ededdd | 0.61 |
| 57 | ITSTPTGLSSHDTYIIKFMVTLNNNNYPRKLKLNFDNSRFSTTPTGFWYTL<br>eeeeedeeeeddeddcdd0deeccdbcccddbcccbaabbbcbcdbcccbc | 0.58 |
| 41 | TIVWELSINQIPWAGTDRTKFRSWYFDPYVPNEPLIYNTSFTTTPNVWSST<br>eeeecfeeeedeedeedddeeeeedeeeecceddedeceededddd0deedd | 0.53 |
| 69 | RTEKKYYERRTLKKLKNNWKEHHKLQKRNSKKKHNFSKKHWKIKIKIKSIK<br>eeeeddedeeeeeeeed8cedecdddd80dedccdeedddeeddedcccd | 0.52 |
| 70 | KNLNQPNDLGKLNKTLPIPLLNFIIRLNQNSDKIRIELRVSDYKNAWDLMI<br>eeee6e0d7edddcedeedcebedeeeeedeededdeedcedcbbdeeec2 | 0.52 |
| 80 | PINNFKTFVVKFILNENFIANTLNPTMLSTLTIIPQTRSLTKSILLNLRRT<br>ddedcdcddecdde9eddcddddcddbcdededeeddccdddddeedecde | 0.44 |
| 129 | DINKKYLVQIKLKLKNTKTKKETKNSSHHKSKRKHKQSLKKHKRIWKKKSH<br>edddeeeddeddcdebcddeeddddedeeeedbddceeedbcbedecddcd | 0.43 |
| 452 | NIKIVLSMHNRVRTEKSLDPSTPFNFTLSVTPPFLTSNVSKVSFMLRLNAD<br>dceeded9d7dbbdcecdeeededdeeedeeeeeeebddddce0eeeeee | 0.39 |
| 86 | KYQLKIQCTPLDERAELIKFTLSISQEPDEQERVEEQHGQPAAQQEEHVKK<br>eeeeedddeeeededeeddeeeeeeedecdddeeeddcedddddeedbddc | 0.37 |
| 181 | LTLYLIIELQDVHEALLLRVDVLDDLGHVLQDPGQLVHDGNRLTITTTLVG<br>eeeeedeeeefddeceeeeeceedeeecacededebcddddeeeeddeeee | 0.35 |
| 328 | IIYYPFRITFNQTPNDIENLSGRISFSPSQPLLLIPSVIASNLWRFLMSGS<br>edeefeeee8efceecddddedeeeddeddedddcecdbcbcddddedebe | 0.33 |
| 326 | KTYKLDKRWNKVNKKKNTKTKKHLKDNKKSIEKHHVKKMKEKKLKDAANKK<br>eebddfdeecdddddcacdcdcdcddddddcccedacd9dcdc8ccdacdb | 0.32 |
| 237 | TKNLHVSDNYRFNLSILQMPHMPNIWLIKLTFTPNKNNEHSKHDKVDSNNY<br>eededecd9deeaeeeee0eecee3eeeeeeecdccccdcdddcedddc9c | 0.30 |

**Table S19**. Top 20 Most abundant sequence clusters in round 9 of gl-PGT121 selection for the Biased library (without negative selection against non-glycosylated binders).

| gl-PGT121 – Biased, Round 9 (without Negative Selection) | | |
|---|---|---|
| ID | Consensus sequence and score | % of all sequences |
| 1 | YPTTHLTSFYVSTNPSTTTIRIGYPSGVLHGHLVPDLVLHLLEQGEPLQLR<br>dededeeeeededeeddeeeeeeeeedecebecdceeeeeceeddecedcde | 26.48 |
| 0 | HSLFNFQVRLKQLPLDANGAVIGDIRIAECMNHTYTPLTFYIYSNHPTDIQ<br>edddeeeededeeeeeddeeeeeeeeeee9bddddeeeededdddcdb0bc | 11.56 |
| 18 | HPTWSLLPFLINIRQQQIKYSIGDIHGVLHVVEPLQGEDDQLLVEPLHQVR<br>eeeeedeeeeedeed6deeeeeeedeeeecedeeeeeeeedddeddeedded | 4.78 |
| 40 | HIHNNIDIKDSTRKNAVSFAFIISFNQTPLGDKMIGDIRKAELHDALRDVL<br>eddcddccddddcdcdddeddceecdedeeedcbedddedddccddde9cd | 2.63 |
| 10 | YPPALDDDWNQFLRFRLNVEDQQVLHIPQNGTWRLAISVSRTDEPEHLHLR<br>edddeddddcddeedaecdddeecdddeddedeecdeedecdddedddaed | 2.28 |
| 47 | TTDFSLPFRLNVDHYGHVSLRRPNYKMTWHDEINIGDIRLARCMFGESTLR<br>edddeeeeeedfeeefeeeee0edeb5ceddadcddeddedd57eebabcb | 0.78 |
| 121 | YYLIFVADLNHKGQISRHSEEGRVHLVRLDDHDHWRYPSGGLHDLGPLLEG<br>feedeeeeecedeeeedeedcec9eeceeedddeeeeeeeeedeeeeeedc | 0.77 |
| 70 | HPNLKYDRIGDIRDACCMYSEPWKHNASVNFSRILKNSTTFYIINTYESTQ<br>eeedeeeeeeceeddee4ddddebdddeebcddde4bbeeddddbbdedee | 0.58 |
| 133 | HSLTNNTHTDVYGSYSCQFYHIGDIRIASCMIHLSNNMWMINHSQDHTLND<br>ee7cecdddfddeeedeedecceeedeeedddcebdddebcdcddeedcde | 0.40 |
| 156 | FIAYYFDLSQPNLHSPNFTLKIGDIREAQCMLTCDSAKPNPRVFPILALLI<br>deedeeeeeeeeddeedddeceeeeeeeddddecddeededded0eddeeee | 0.39 |
| 332 | ATSWQLYTLAISVSRVHDVLGEGDPRRDRRLPEVQVRLVLVPLEGVDQLVG<br>eeceeeeedbddedeebcedceccefeeeedeecd9feedfeeeedcdeddd | 0.37 |
| 96 | HSIMLPPLWFWTLTLRQVPQHINMRTKADTDGYFIGDPSGEVHGEPRLHVR<br>fedceeededeefeeeeeffbbdbeedeeddfecedeeceeddedfeedee | 0.36 |
| 28 | HNTNDILVLNFVYTQNLGQHKLLVYHSTHEVDDLIGDIRTANDDHPLQHAR<br>eaccccceeedeeeefddeceeeeeeebdddcdde0dbcddbdcc0ccddbc9 | 0.36 |
| 5 | VNPQQHLDDDHDHLIDSFQFQIEGLRRTSSSATLVISIRRVHADVALQEAP<br>eeeeedeeeeedddeedeeeddeecffeeeffceedefdefedffeeeeeee | 0.31 |
| 84 | GDYYTFQIIGDIRQRPADGPAHAARVRVLDDRNHHHSIYQIYPFVLSINAG<br>dddccccceefeeeededeeee4cefedfe7cecbbcbddcdddcdcdcccc | 0.30 |
| 178 | ITHLNFPLIGDIRWAFLHDRDAPLLELAEAGHVLVPLVVQRDVHGGRLVLA<br>ceeddaddbfcdfffeededfffffeffeedceedfedfeedcffdefefe | 0.29 |
| 161 | HPELLDHLVVADRREHPAQEHWRYPSRVVHVALRVGQEPGDVAGRHLLRLR<br>eecdecdefddbbddedecddeffeceeee8afeeefeeeeddeedeefed | 0.28 |
| 59 | ISQWSVYRLAISVADLHAAEGDRVRVVVRRVDDPQPDELLDRLYGFNLRIG<br>ddeeebfedeeededceedcfdeeeeeeeedeeddefdedfcdeeddceddd | 0.28 |
| 76 | HSHSPISVLATIKTSNLHNSPTHGLLLIRVSTLDIGDIRADLHVDAHVDEA<br>eededdddeddeedcdeddddddedeeeeedfcdeeededdeeddbddddd | 0.28 |
| 42 | TNARRAHIIGDIRSVLHAPHAVRPDDQRQGDPLQHVRGVHFLTIKIKQAPG<br>ededddcdeeeeeedcdededeeeebeddddef9eedefedeeeefdd9fdee | 0.28 |

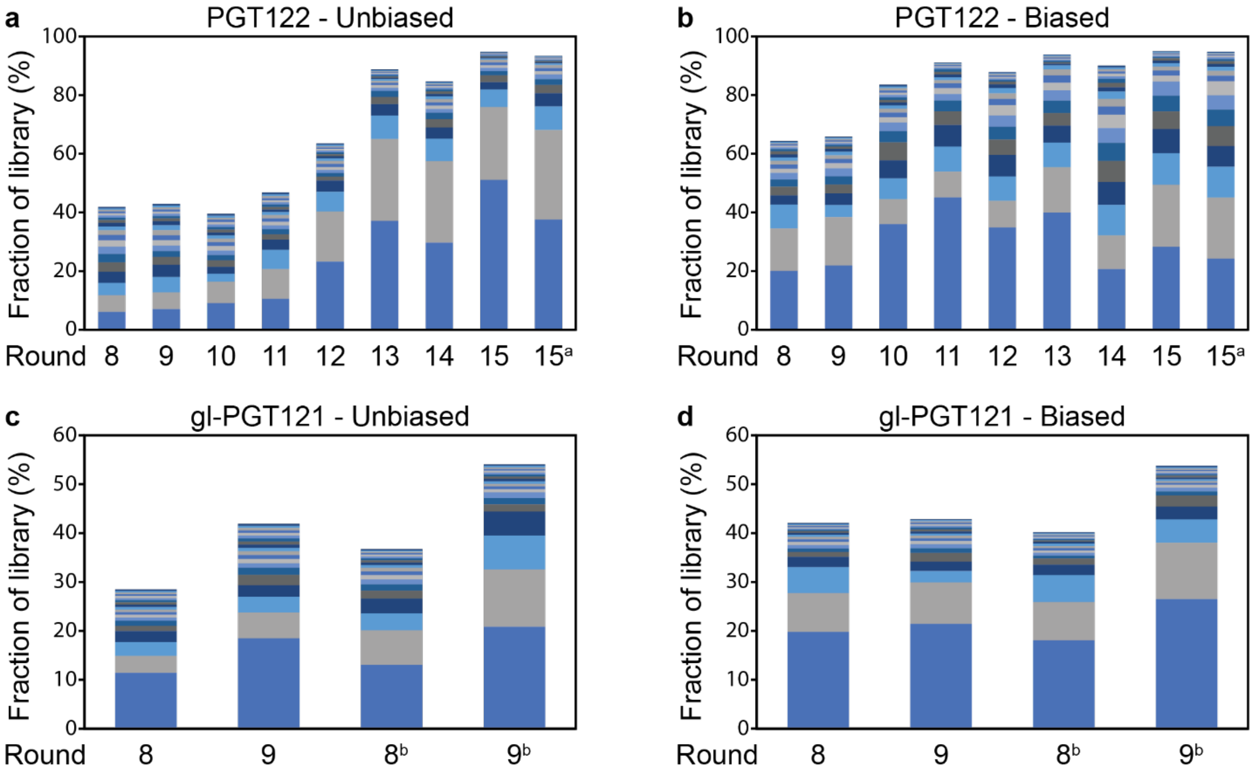

**Figure S38**. Enrichment of the top 20 most abundant sequence clusters for the (a) PGT122 Unbiased library, (b) PGT122 Biased library, (c) gl-PGT121 Unbiased library, and (d) gl-PGT121 Biased library.
a This version of round 15 of PGT122 was performed with non-glycosylated libraries.
b These rounds of gl-PGT121 selection were performed without negative selection against non-glycosylated binders.

**E. Ribosomally-synthesized PGT122/gl-PGT121 Selection Winners**

*1. PCR from Plasmids (gl-PGT121 Selection Clones)*
DNA for preliminary analysis of gl-PGT121 selection winners was PCR amplified from encoding plasmids as a template. Plasmids (10 ng) were added into a 100 µL PCR reaction with final 0.2 mM of each dNTP, 1 µM of each forward primer (Library FP1) and reverse primer (Library RP1.3 for Unbiased library; Library RP2.1 for Biased library), 0.1 ng/µL plasmid template, and 0.025 U/µL Taq DNA Polymerase (New England Biolabs) in 1X Standard Taq Buffer. The following PCR protocol was used: 18 cycles of 94 °C for 30 seconds, 55 °C for 30 seconds, and 74 °C for 30 seconds. The product DNA was purified by extraction with phenol/chloroform and then chloroform followed by ethanol precipitation.

*2. PCR from Library DNA (PGT122 Selection Clones)*
For PGT122 sequences, DNA corresponding to enriched clusters determined from NGS data was prepared by PCR-amplification of libraries using sequence specific primers. As a representative example, 2U7 was chosen for preliminary analysis because it was the second most abundant cluster after round 11 (8.2% of all sequences). A variant of 2U7 with amino acid sequence given in **Table S15** on row ID 0 became the most abundant sequence after round 15. To isolate 2U7 from DNA recovered from round 11 of selection, 40-nucleotide primers were designed with 20-nucleotide complementarity to one end of the randomized region and 20-nucleotide complementarity to the flanking constant region (**Table S20**). The first PCR using these 40-nucleotide primers was set up as a 20 µL PCR reaction with 0.2 mM of each dNTP, 0.5 µM of each forward and reverse primer, 0.05 ng/µL template DNA, and 0.02 U/µL Phusion™ Hot Start II Polymerase (Thermo Scientific) in 1X Phusion HF Buffer. The following PCR protocol was used: 98 °C for 1 minute, followed by 25 cycles of 98 °C for 10 seconds, 72 °C for 10 seconds (annealing and extension steps combined), followed by 72 °C for 5 minutes. DNA products were purified by running in a 2% agarose gel, visualizing with a 365 nm handheld UV lamp, and extracting using QIAquick Gel Extraction Kit (Qiagen) according to manufacturer instructions. The purified DNA from the first PCR was then

extended to full length using the extension primers originally used to extend purchased synthesized libraries. Specifically, DNA was diluted to 1.25 ng/µL in a second PCR reaction (20 µL) with final same final setup as the first PCR reaction but with extension primers (SSSF1 and either SSSR1 or SSSR2 for Unbiased and Biased libraries, respectively) and only 20 PCR cycles. DNA was then prepared for Sanger sequencing, and a plasmid encoding the desired sequence was PCR amplified as described in **Section IV.E.1**. The sequences prepared for preliminary binding assays are summarized in **Table S21**.

**Table S20**. Nucleotide sequence for the most abundant variant of 2U7 at the end of round 11 as well as the 40-nucleotide primers used to isolate the sequence from recovered DNA library. The highlighted nucleotides indicate the regions of complementarity of the primers, with "|" added to visualize the break from constant to randomized regions. Underlined regions were added in the second PCR using extension primers listed in **Table S11**.

| Name | Sequence (5′ to 3′) |
|---|---|
| 2U7 | TAATACGACTCACTATAGGGTTAACTTTAGTAAGGAGGACAGCTAAATGGCG\|AAGACCAGGACCCCGATGAACTTGTCGGGGAAGTCCAACATCCGGTACTTCTTCACGATCGACAACTCGACGATGTTCGAGGACCTCTTCATGCGCTTCCCGCGCAGCAAGGAGTACCACATCGAGCTCGAGGAGTTCAAGGACTTGATCCGC\|GGCTCCGGTAGCTTAGGCCACCATCACCATCACCACCGGCTATAGGTAGCTAG |
| 2U7-1 | AGGAGGACAGCTAAATGGCGAAGACCAGGACCCCGATGAA |
| 2U7-2 | TGGCCTAAGCTACCGGAGCCGCGGATCAAGTCCTTGAACT |

**Table S21**. Clones identified from PGT122/gl-PGT121 selections NGS data that were prepared for preliminary binding analysis.

| Name | Sequence |
|---|---|
| 2U1 | AKTKTYLIRFHLDILPNDNLRSQKIRLIFGDKATVKGMITFLNDSELEWNFE |
| 2U3 | ANPSFSVQFIVDLIGPNVMRITLSNKPFATICPITEGESHISKRFNFQLTIS |
| 2U6 | AQYIDWEIDINQVPNQMNLSKPTWKSDITINKVKWSFYININQASNENTDNI |
| 2U7 | AKTRTPMNLSGKSNIRYFFTIDNSTMFEDLFMRFPRSKEYHIELEEFKDLIR |
| 2U7* | AKTRTPMNLSGKSNIRYFFTIDNSTMFEDLFMRFPRSKEYHIELEEFEDLIR |
| 2U8 | AYSIHFVCDINETPTNTSKAIFSIKCDITTPESTTRQLMITKHYHFPPTGTR |
| 2B2 | AYPVYLRLPIGDIRIASCMFGDEPNTWTRDELNDYFYTESYNPQPITITFKC |
| 2B3 | AFIAYYFDLSQPNLHSPNFTLKIGDIREAQCMLTCDSAKPNPRVVPILALLI |
| 2B4 | ALNLYQIVEFTIDESNDIVQLQIGDIRTADCMSLNKECYNIIFFSWDNHGKN |
| 2B5 | ATLNIFSNFSIEPDSFYEIVLPIGDIRIAPCMSSFNRNGSTNNCYHFEINLF |
| 2B7 | AYVISFSIDQPDLDRPNIIRISIGDIRSAKCMKLDTINFAYRDDSSEVCYEK |
| 2B8 | ANNFICIRNIGDIRSARCMNVSAGRYAPYREFYFDITYDDFNKNVNIVITPV |
| GU1 | ANPRIIFFELSIRNNSLKLLRWHDTQGIFHDSMTELSQHAELPRFKFHSNAA |
| GB1 | AHSLFNFQVRLKQLPLDANGAVIGDIRIAECMNHTYTPLTFYIYSNHPTDIQ |

*Denotes 2U7 K->E variant that became prevalent during last rounds of selection and was prepared synthetically for BLI binding assay.

*3. Non-radioactive Peptides for MALDI-TOF Characterization*

DNA produced from both routes was transcribed, and the crude transcripts were purified by 5% denaturing urea-PAGE. For analysis by MALDI-TOF-MS, translation (20 µL) was carried out in the presence of PDF/MAP but without addition of radioactivity. Depending on whether sequences had two cysteine residues or not, the crude peptides were purified by Ni-NTA agarose either with or without cyclization as previously described for the preparation of 12A in **Section I.B**, though purification with cyclization was slightly modified as described below. A portion of each eluent in 0.2% TFA (1.5 µL) was analyzed by MALDI-TOF-MS to verify sequences were correctly translated and, if applicable, cyclized.

The protocol used for cyclization while captured on resin was as follows: To 20 µL of crude translated peptide was added 80 µL of bind buffer with BME (50 mM Tris-HCl, pH 7.8, 300 mM NaCl, 5 mM BME) and 20 µL of HisPur™ Ni-NTA Resin (ThermoScientific). The mixture was tumbled at room temperature for 1 hour. Next, the mixture was spun down at 17,000 ×g, and the supernatant was carefully removed from the loosely-pelleted agarose. The resin was washed three times with 160 µL of cyclization wash buffer with TCEP (50 mM Tris-HCl, pH 8.0, 300 mM, 0.5 mM TCEP). To the resin was then added 160 µL of free peptide cyclization buffer (50 mM Tris-HCl, pH 8.0, 300 mM NaCl, 0.5 mM TCEP, 3.3 mM m-dibromoxylene, 33% (v/v) MeCN), and the mixture was covered in foil and tumbled at room temperature for 30 minutes. The supernatant was removed and 160 µL of cyclization wash buffer with BME (50 mM Tris-HCl, pH 8.0, 300 mM NaCl, 10 mM BME, 33% (v/v) MeCN) was added. The suspension was tumbled at room temperature for 10 minutes without aluminum foil to quench the reaction. The reaction was spun down, and the supernatant was removed. Next, the resin was resuspended in wash buffer (50 mM Tris-HCl, pH 7.8, 5 mM BME) and transferred to a 0.22 µm Ultrafree-MC Centrifugal Filter (MilliporeSigma). The tube was rinsed with 80 µL of wash buffer, and the rinse was added to the filter. The filter was then spun down at 10,000 ×g for 1 minute. The resin in the filter was washed twice with 160 µL of wash buffer. To elute the peptide, 20 µL 0.2% (v/v) TFA was added to the resin and allowed to sit at room temperature for 2 minutes before spinning down and collecting in a new tube. The elution step was carried out once more, with the eluent collected separately.

*4. Radioactive Peptides for Bead-based Radiometric Binding Assay*

For binding analysis, translation (50 µL) was carried out in the presence of PDF/MAP and $^3$H-histidine for radiolabeling. The crude peptides were purified by Ni-NTA agarose and, if the sequence contained two cysteine residues, was cyclized while captured on the Ni-NTA resin. The peptide eluents were combined and dialyzed as follows: To the combined eluents was added a solution of Triton X-100 in water for a final concentration of 0.1%. Next, the peptides were dialyzed against 0.1% (v/v) Triton X-100 using a Slide-A-Lyzer™ MINI device (0.1 mL capacity, 3.5 kDa MWCO) (Thermo Scientific) at room temperature overnight. The dialyzed peptides were retrieved from the devices. The peptide solutions were concentrated to dryness by speedvac for the click reaction.

The click reaction (5 µL) with $Man_9(GlcNAc)_2$-azide was carried out as previously described. Following the reaction, samples were diluted with 0.1% (v/v) Triton X-100 to 30 µL. Upon observation of insoluble material in the tube, the solution was acidified with TFA to improve solubility. To neutralize the glycosylated peptides before binding assays, peptides were dialyzed against 0.1% (v/v) Triton X-100, as before. The glycosylated peptides were analyzed by 4-20% SDS-PAGE (Bio-Rad) to verify efficient glycosylation. Next, 5X selection buffer was added to the peptide solutions to final 1X.

Sequences were screened for their binding to PGT122 as follows: Each glycosylated peptide (3K CPM/sample) was incubated with 0, 100, 200, or 400 nM PGT122, diluted to 16 µL with 1X selection buffer, and tumbled for 1 hour at room temperature. While tumbling, Dynabeads™ Protein G (Invitrogen) were equilibrated with 1X selection buffer as follows: Steps were carried out using a multi-channel pipettor where possible. For each sequence, 8 µL of Protein G bead suspension was transferred to a 0.2 mL PCR tube in a strip. Next, 64 µL (8 bead vol.) of 1X selection buffer was added to the beads. The mixture was resuspended by gently vortexing, spun down briefly, and placed on a magnet for 1 minute. The supernatant was removed, and this wash was repeated two more times. The final supernatant was removed just prior to addition of the incubated peptide/antibody mixture. The mixture was allowed to tumble at room temperature for 30 minutes to capture antibodies and bound peptides. Next, the suspension was briefly spun down, placed on a magnet, and the supernatant was transferred to an individual 0.2 mL PCR tube. To wash the beads, 16 µL of 1X selection buffer was added, gently vortexed, spun down briefly, placed on a magnet for 1 minute, and the wash was transferred to the same PCR tube as the supernatant. This wash step was repeated twice more for a total of three washes, saved altogether with the supernatant as the "unbound" fraction. The beads were then resuspended in 50 µL of denaturing elute buffer. The suspension was heated at 95 °C for 5 minutes to denature antibodies, spun down briefly, placed on a magnet for 1 minute, and the eluent was transferred to an individual 0.2 mL PCR tube. The beads were washed with 50 μL of denaturing elute buffer, and the wash was placed into the same tube as the eluent. The combined eluent and wash represented the "bound" fraction. The beads were resuspended in 50 μL of denaturing elute buffer, vortexed, and directly placed into a vial with liquid scintillation cocktail for radioactivity measurements, though it was not included in calculations of fraction bound. Each of the tubes with the "unbound" and "bound" fractions were placed into vials for LSC. The tube in which the antibody and peptides were incubated were also measured by LSC, but not included in calculations of fraction bound. Fraction bound values for each antibody concentration after background adjustment were input into KaleidaGraph 4.1.1 for curve fitting to a Michaelis-Menten model, and the resulting dissociation constants ($K_D$) are given in **Table S22**.

*5. Clones Prepared as cDNA/mRNA-glycopeptide Fusions*

Selected clones were also prepared as cDNA/mRNA-glycopeptide fusions for binding analysis. RNA (450-1000 pmol) was photo-crosslinked as described in **Section IV.A.2** but with 10 μM transcribed RNA and 15 μM XL-PSO in 1X XL Buffer for annealing, and dilution to 100 μL for irradiation at 365 nm. XL-RNA was then purified by ethanol precipitation and 5% denaturing urea-PAGE with electroelution. Purified XL-RNA was translated in the presence of PDF/MAP and $^3$H-histidine for radiolabeling. Crude translation reactions were captured on Pierce™ Streptavidin UltraLink™ Resin (Thermo Scientific), subjected to reverse transcription, and cyclized as described for selection. Fusions were then washed and photo-eluted with 1X selection buffer instead of 0.2% (v/v) Triton X-10. For clones that did not require cyclization, fusions were directly washed 5 times with 1X selection buffer after reverse transcription in preparation for photo-elution. Note: 2U3 was also cyclized despite having only one cysteine residue to better mimic its structure during selection. Fusions were then passed through 0.22 μm Ultrafree-MC Centrifugal Filters (MilliporeSigma), buffer exchanged by ethanol precipitation, and subjected to click reaction (5 μL) with $Man_9(GlcNAc)_2$-azide as described in **Section II.A.8**. Binding assays were performed analogously as with free peptides but using 2K CPM per sample, and the results of KaleidaGraph calculations are also given in **Table S22**.

**Table S22**. Binding assay results of ribosomally-synthesized PGT122/gl-PGT121 selection winners prepared both as free peptides and cDNA/mRNA-glycopeptide fusions. “c” after the clone name indicates that they were cyclized.

| | | Peptide | | Fusion | |
|---|---|---|---|---|---|
| Name | Click | $K_D$ | Ratio (No click : Click) | $K_D$ | Ratio (No click : Click) |
| 2U1 | - | ND | N/A | 82.5 | 2.8 : 1.0 |
| | + | ND | | 29.8 | |
| 2U3c[a] | - | 1494.9 | 3.8 : 1.0 | 280.6 | 1.0 : 1.6 |
| | + | 397.6 | | 448.8 | |
| 2U6 | - | 165.6 | 1.0 : 4.1 | 96.7 | 1.0 : 1.8 |
| | + | 683.5 | | 173.4 | |
| 2U7 | - | 1428.8 | N/A | 299.9 | 2.2 : 1.0 |
| | + | ND | | 139.2 | |
| 2U8c | - | ND | N/A | 173.4 | 3.6 : 1.0 |
| | + | 151.6 | | 48.4 | |
| 2B2c | - | 1647.8 | 7.0 : 1.0 | 89.0 | 1.0 : 1.3 |
| | + | 235.1 | | 115.6 | |
| 2B3c | - | 4490.5 | 6.9 : 1.0 | 99.7 | 1.0 : 1.4 |
| | + | 651.2 | | 140.0 | |
| 2B4c | - | 157.7 | 1.0 : 1.6 | 44.4 | 1.0 : 3.5 |
| | + | 247.2 | | 155.2 | |
| 2B5c | - | 212.3 | 1.0 : 1.5 | 314.5 | 3.7 : 1.0 |
| | + | 317.7 | | 85.9 | |
| 2B7c | - | 29.8 | 1.0 : 2.4 | 66.2 | 1.0 : 2.2 |
| | + | 71.9 | | 143.3 | |
| 2B8c | - | ND | N/A | 176.3 | 1.6 : 1.0 |
| | + | 247.4 | | 107.6 | |
| GU1 | - | 156.1 | 1.0 : 14.7 | | |
| | + | 2294.4 | | | |
| GB1 | - | 181.2 | 1.0 : 5.2 | | |
| | + | 948.1 | | | |

a 2U3 was cyclized as a fusion but not as a free peptide.

### V. Synthesis of Individual Peptides and Glycopeptides from PGT122/gl-PGT121 Selections

#### A. Synthetic Scheme, Experimental Procedure, and LC-MS Chromatogram of Individual Peptides and Glycopeptides

*1. 2U7-glycopeptide (**9**)*

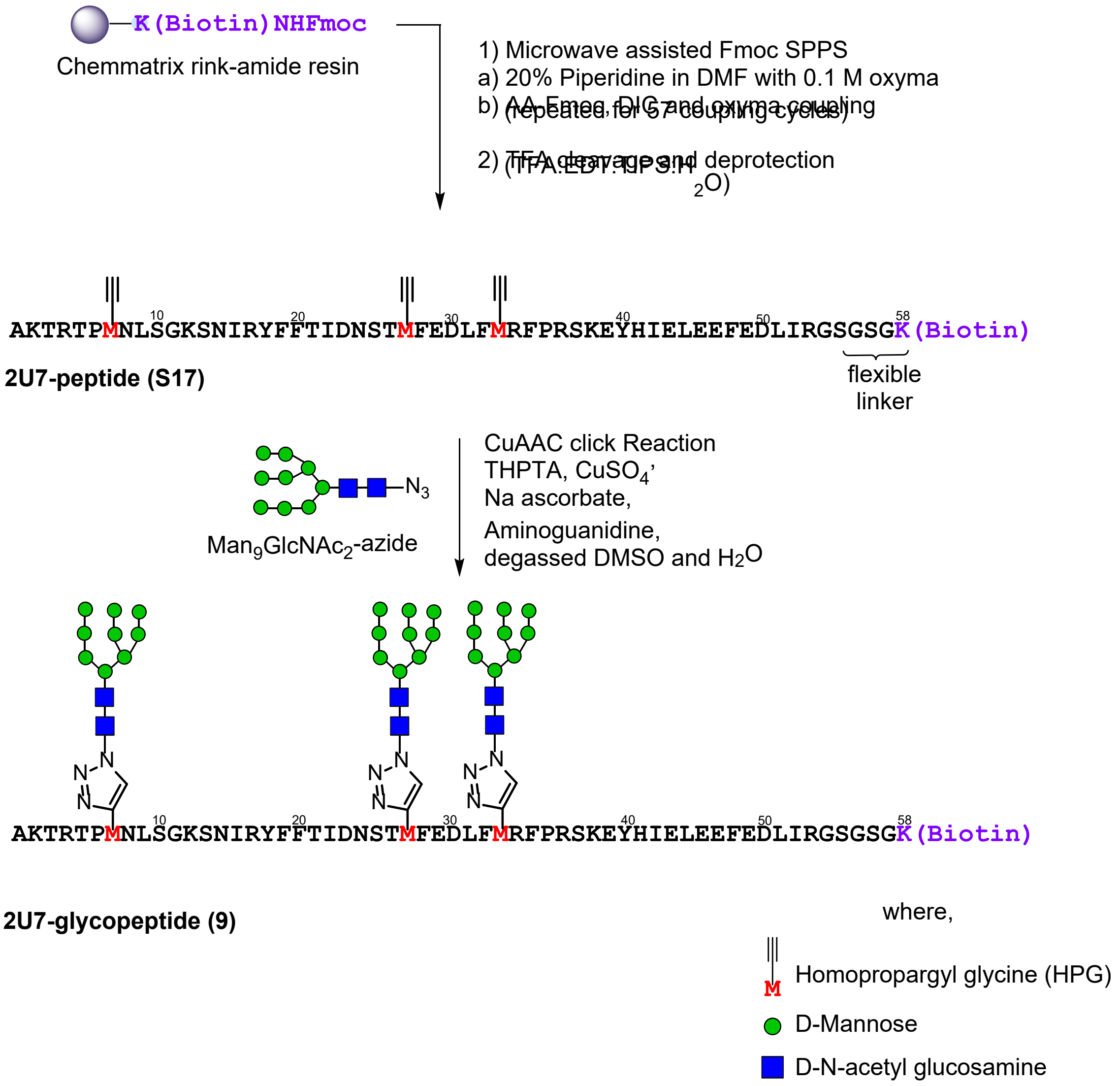

**Scheme S11.** Synthetic scheme to generate **2U7-glycopeptide (9).**

**2U7-peptide (S17):** 2U7-peptide (**S17**) was prepared as in the representative procedure for solid phase peptide synthesis (**Section III.A.4**), starting with 155 mg of resin loaded at 0.17 mmol/g (26 µmol). Of the crude solid peptide, 60 mg was subjected to purification by RP-HPLC on the 10 x 250 mm C4 column (5-30% B over 45 min). 2.1 mg of pure peptide was obtained (3.5 % of the material that was purified). ESI-LR observed *m/z* of multiply charged ions 779.04 $[M+9H]^{9+}$, 876.35 $[M+8H]^{8+}$, 1001.61 $[M+7H]^{7+}$, 1168.02 $[M+6H]^{6+}$; calculated average m/z values for 2U7-peptide $C_{317}H_{477}N_{85}O_{93}S_1$ : 778.65 $[M+9H]^{9+}$, 875.85 $[M+8H]^{8+}$, 1000.83 $[M+7H]^{7+}$, 1167.47 $[M+6H]^{6+}$.

**2U7-glycopeptide (9):** The representative procedure for the click glycosylation of synthetic peptides **(Section III.A.8**) was carried out with 2U7- peptide (**S17**) (0.63 mg, 0.10 µmol) and $Man_9GlcNAc_2$-azide (6.6 µL from 50 mM stock in water, 0.63 mg, 0.33 µmol). After HPLC purification on the 4.6 x 250 mm C4 column (2-30% B over 45 minutes), 67 µg (0.0053 µmol) of pure 2U7-glycopeptide (**9**) (5 %) yield was obtained (quantified by UV-nanodrop). ESI-LR observed *m/z* of multiply charged ions 1414.81 $[M+9H]^{9+}$, 1591.54 $[M+8H]^{8+}$, 1818.67 $[M+7H]^{7+}$; calculated average m/z values for 2U7-glycopeptide $C_{527}H_{828}N_{100}O_{258}S_1$ : 1414.87 $[M+9H]^{9+}$, 1591.61 $[M+8H]^{8+}$, 1818.84 $[M+7H]^{7+}$.

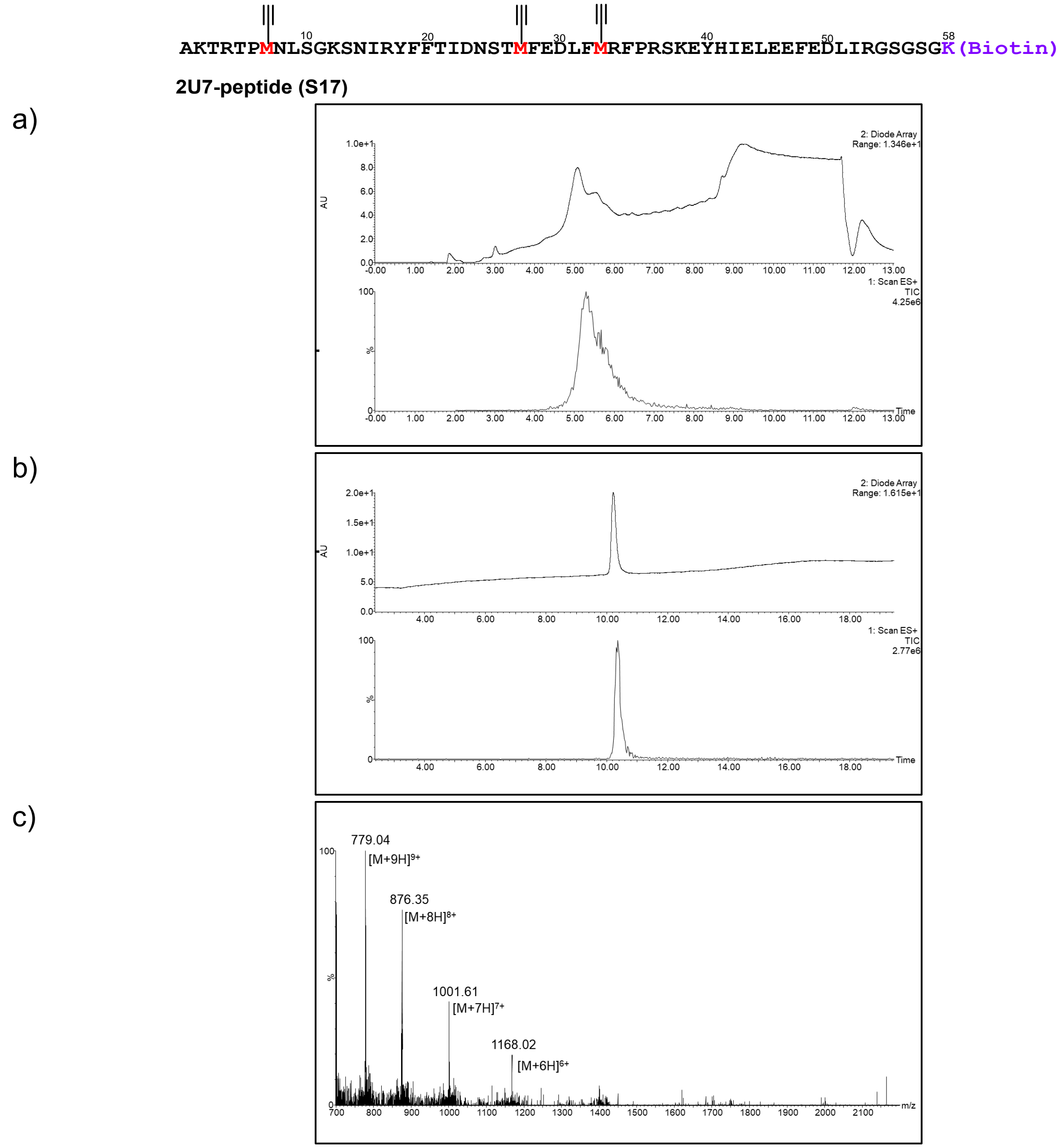

**Figure S39**. LC-MS of **2U7-peptide (S17)**. a) UV and ESI+ TIC chromatogram of 2U7-peptide (crude), b) UV and ESI+ TIC chromatogram of 2U7-peptide, c) ESI+ MS of 2U7-peptide.

(LC-MS method for crude 2U7-peptide: 1% A to 5% B over 1 minute then 5%A to 60%B gradient over 8 minutes; LC-MS method for purified 2U7 peptide: 1% A to 5% B over 1 minute then 5%A to 60%B gradient over 18 minutes, solvent A was water/0.07% formic acid and solvent B was acetonitrile/0.07% formic acid, column: Acquity UPLC@Protein BEH C4, 2.1 mm * 150 mm, 1.7 µm; flow rate: 0.25 mL per minute.)

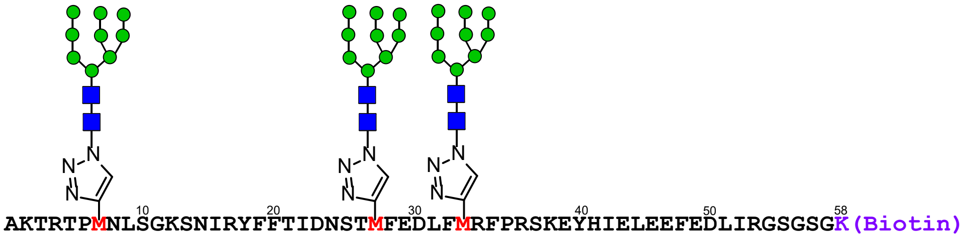

**2U7-glycopeptide (9)**

a)

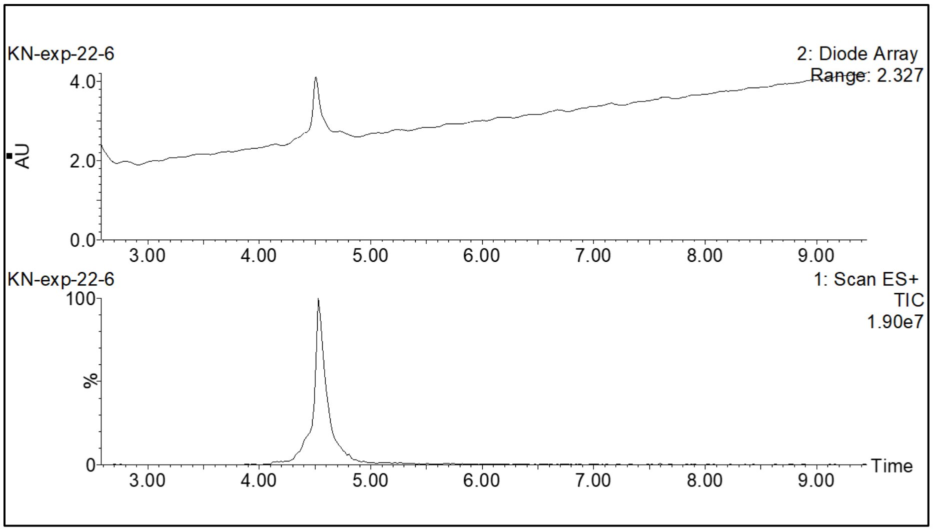

b)

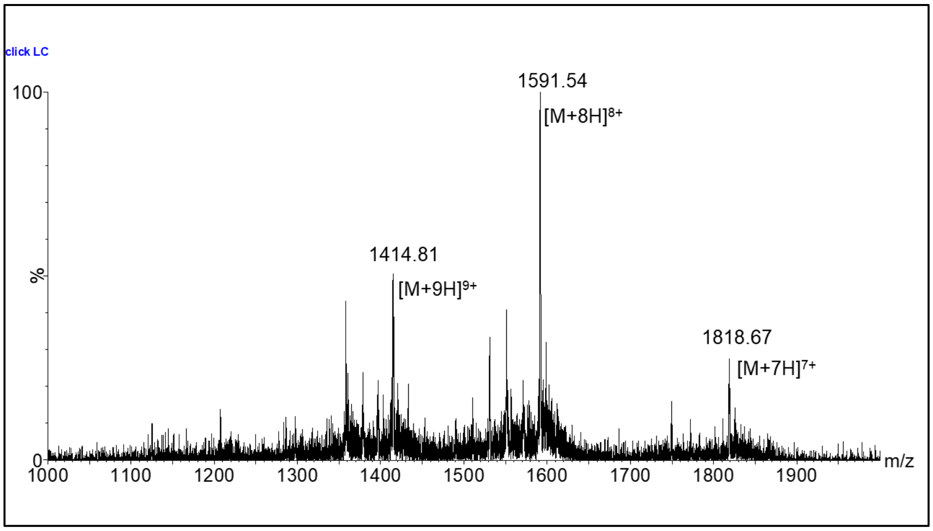

**Figure S40**. LC-MS of **2U7-glycopeptide (9)**. a) UV and ESI+ TIC chromatogram of 2U7-glycopeptide, b) ESI+ MS of 2U7-glycopeptide.

(LC-MS method: 1% A to 10% B over 1 minute then 10%A to 45%B gradient over 10 minutes, solvent A was water/0.07% formic acid, and solvent B was acetonitrile/0.07% formic acid, column: Acquity UPLC@Protein BEH C4, 2.1 mm * 150 mm, 1.7 µm; flow rate: 0.3 mL per minute.)

*2. 2U7-QQQ-non-glycosylated Peptide (**10**)*

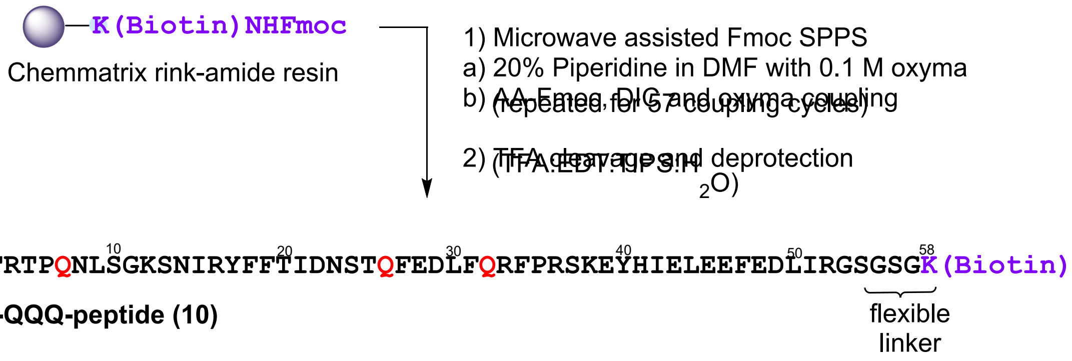

**Scheme S12.** Synthetic scheme to generate **2U7-QQQ glycopeptide (non-glycosylated peptide) (10)**.

**2U7-QQQ peptide (10):** 2U7-QQQ peptide (**10**) was prepared as in the representative procedure for solid phase peptide synthesis (**Section III.A.4**), starting with 155 mg of resin loaded at 0.17 mmol/g (26 µmol). Of the crude solid peptide, 40 mg was subjected to purification by RP-HPLC on the 10 x 250 mm C4 column (5-30% B over 45 min). 1.5 mg of pure peptide was obtained (3.7 % of the material that was purified). ESI+ MS observed *m/z* of multiply charged ions 785.22 $[M+9H]^{9+}$, 883.18 $[M+8H]^{8+}$, 1009.22 $[M+7H]^{7+}$, 1177.18 $[M+6H]^{6+}$, 1412.04 $[M+7H]^{7+}$; calculated average m/z for 2U7-QQQ peptide $C_{314}H_{480}N_{88}O_{96}S_1$ : 784.98 $[M+9H]^{9+}$, 882.98 $[M+8H]^{8+}$, 1008.98 $[M+7H]^{7+}$, 1176.97 $[M+6H]^{6+}$, 1412.16 $[M+7H]^{7+}$.

AKTRTPQNLSGKSNIRYFFTIDNSTQFEDLFQRFPRSKEYHIELEEFEDLIRGSGSGK(Biotin)

**2U7-QQQ-peptide (10)**

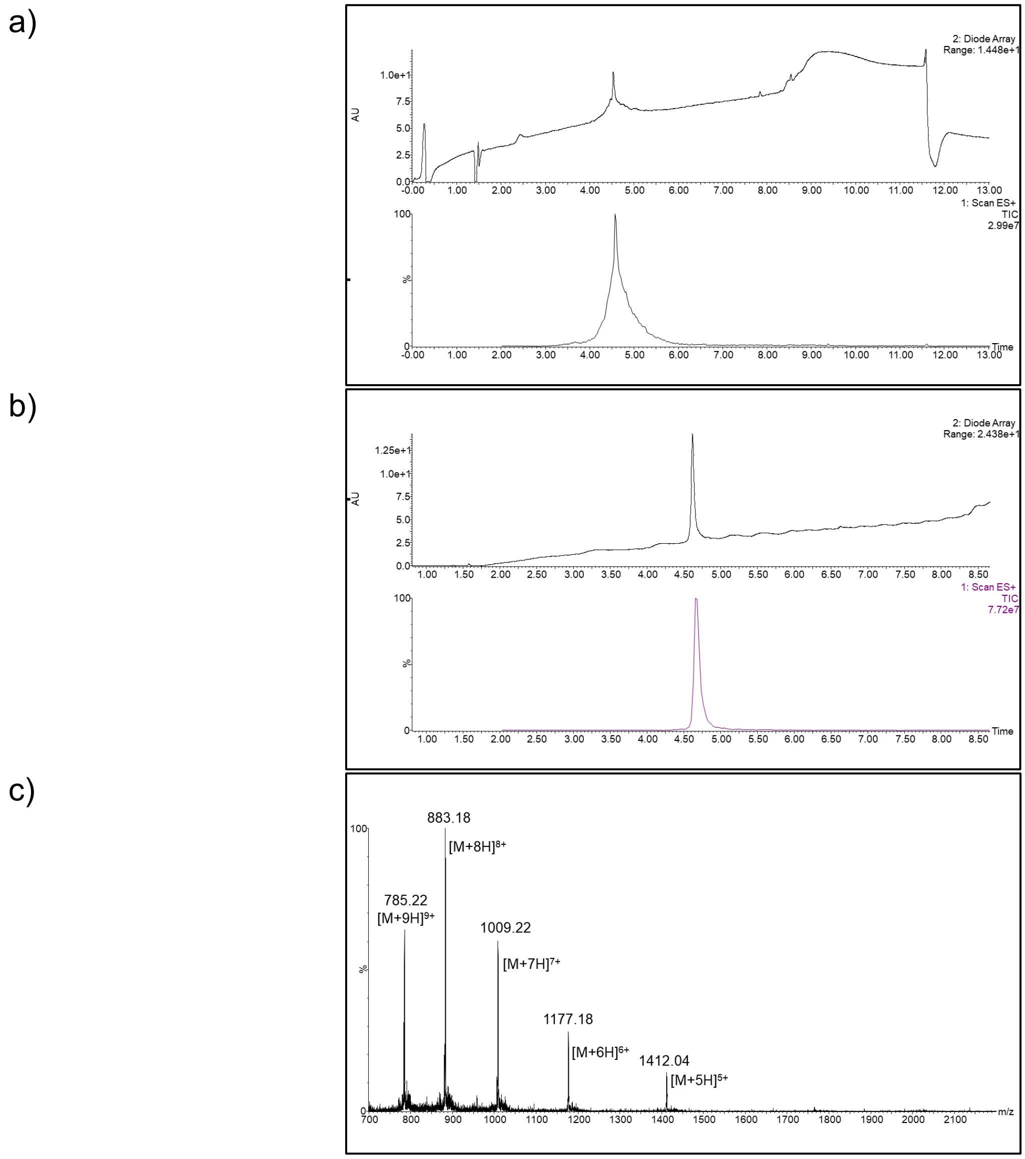

**Figure S41**. LC-MS of **2U7-QQQpeptide (10)**. a) UV and ESI+ TIC chromatogram of 2U7-QQQ peptide (crude), b) UV and ESI+ TIC chromatogram of 2U7-QQQ peptide, c) ESI+ MS of 2U7-QQQ-peptide.

(LC-MS method: 1% A to 5% B over 1 minute then 5%A to 60%B gradient over 8 minutes, solvent A was water/0.07% formic acid and solvent B was acetonitrile/0.07% formic acid, column: Acquity UPLC@Protein BEH C4, 2.1 mm * 150 mm, 1.7 µm; flow rate: 0.25 mL per minute.)

*3. GB1R-glycopeptide (**11**)*

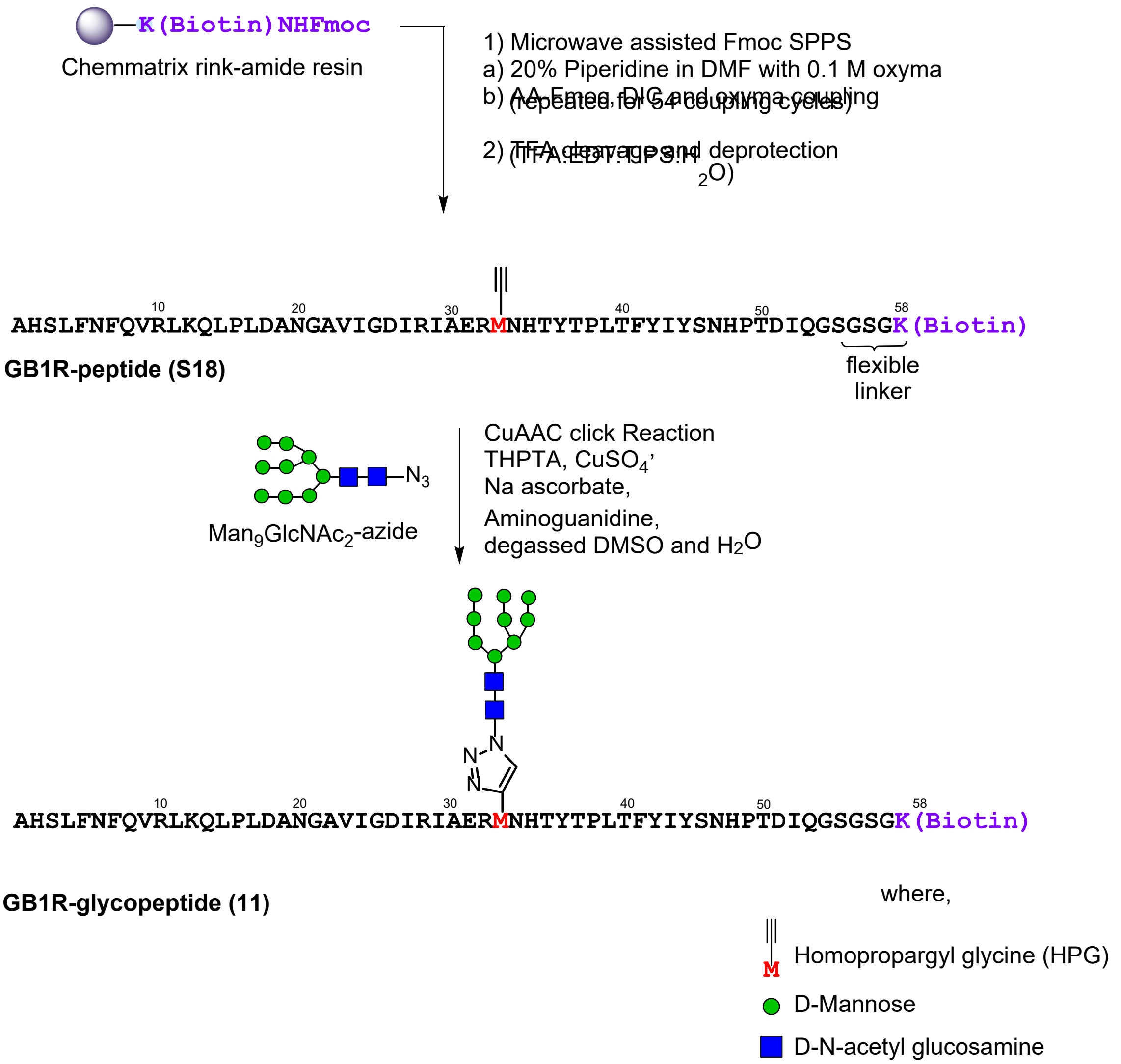

**Scheme S13.** Synthetic scheme to generate **GB1R- glycopeptide (11).**

**GB1R-peptide (S18):** GB1R-peptide (**S18**) was prepared as in the representative procedure for solid phase peptide synthesis (**Section III.A.4**), starting with 172 mg of resin loaded at 0.29 mmol/g (50 µmol). Of the crude solid peptide, 60 mg was subjected to purification by RP-HPLC on the 10 x 250 mm C4 column (5-45% B over 45 min). 1.1 mg of pure peptide was obtained (1.8 % of the material that was purified). ESI-LRMS observed *m/z* of multiply charged ions 834.04 $[M+8H]^{8+}$, 955.79 $[M+7H]^{7+}$, 1114.91 $[M+6H]^{6+}$, 1337.35 $[M+5H]^{5+}$, 1670.94 $[M+4H]^{4+}$; calculated average m/z values for GB1R-peptide $C_{301}H_{461}N_{85}O_{86}S_1$ : 835.82 $[M+8H]^{8+}$, 955.07 $[M+7H]^{7+}$, 1114.09 $[M+6H]^{6+}$, 1336.70 $[M+5H]^{5+}$, 1670.63 $[M+4H]^{4+}$.

**GB1R-glycopeptide (11):** The representative procedure for the click glycosylation of synthetic peptides (**Section III.A.8**) was carried out with GB1R- peptide (**S18**) (1.1 mg, 0.16 µmol) and $Man_9GlcNAc_2$-azide (3.5 µL from 50 mM stock in water 0.34 mg, 0.18 µmol). After HPLC purification on the 4.6 x 250 mm C4 column (2-30% B over 45 minutes), 32 µg (0.0038 µmol) of pure GB1R-glycopeptide (**11**) (3 %) yield was obtained (quantified by UV-nanodrop). ES-LRMS observed *m/z* of multiply charged ions 1074.61 $[M+8H]^{8+}$, 1227.74 $[M+7H]^{7+}$, 1432.81 $[M+6H]^{6+}$; calculated average m/z values for GB1R-glycopeptide $C_{371}H_{578}N_{90}O_{141}S_1$ : 1074.40 $[M+8H]^{8+}$, 1227.74 $[M+7H]^{7+}$, 1432.20 $[M+6H]^{6+}$.

10 20 30 40 50 58
**AHSLFNFQVRLKQLPLDANGAVIGDIRIAERMNHTYTPLTFYIYSNHPTDIQGSGSGK(Biotin)**

**GB1R-peptide (S18)**

a)

b)

c)

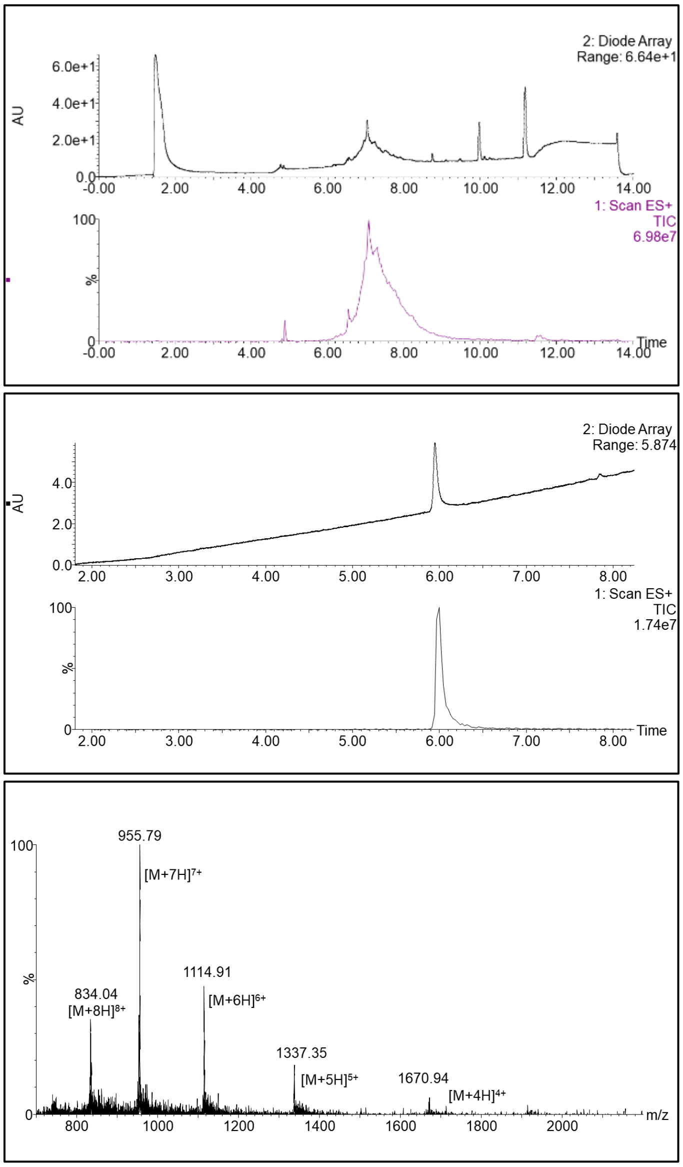

**Figure S42**. LC-MS of **GB1R-peptide (S18)**. a) UV and ESI+ TIC chromatogram of GB1R-peptide (crude), b) UV and ESI+ TIC chromatogram of GB1R-peptide, c) ESI+ MS of GB1R-peptide.

(LC-MS method: 1% A to 5% B over 1 minute then 5%A to 60%B gradient over 10 minutes, solvent A was water/0.07% formic acid and solvent B was acetonitrile/0.07% formic acid, column: Acquity UPLC@Protein BEH C4, 2.1 mm * 150 mm, 1.7 µm; flow rate: 0.25 mL per minute.)

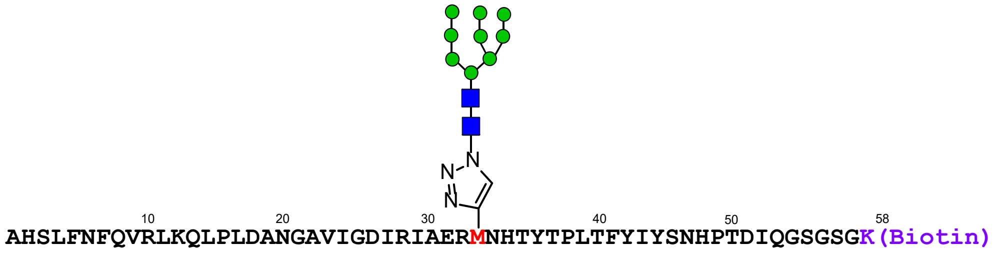

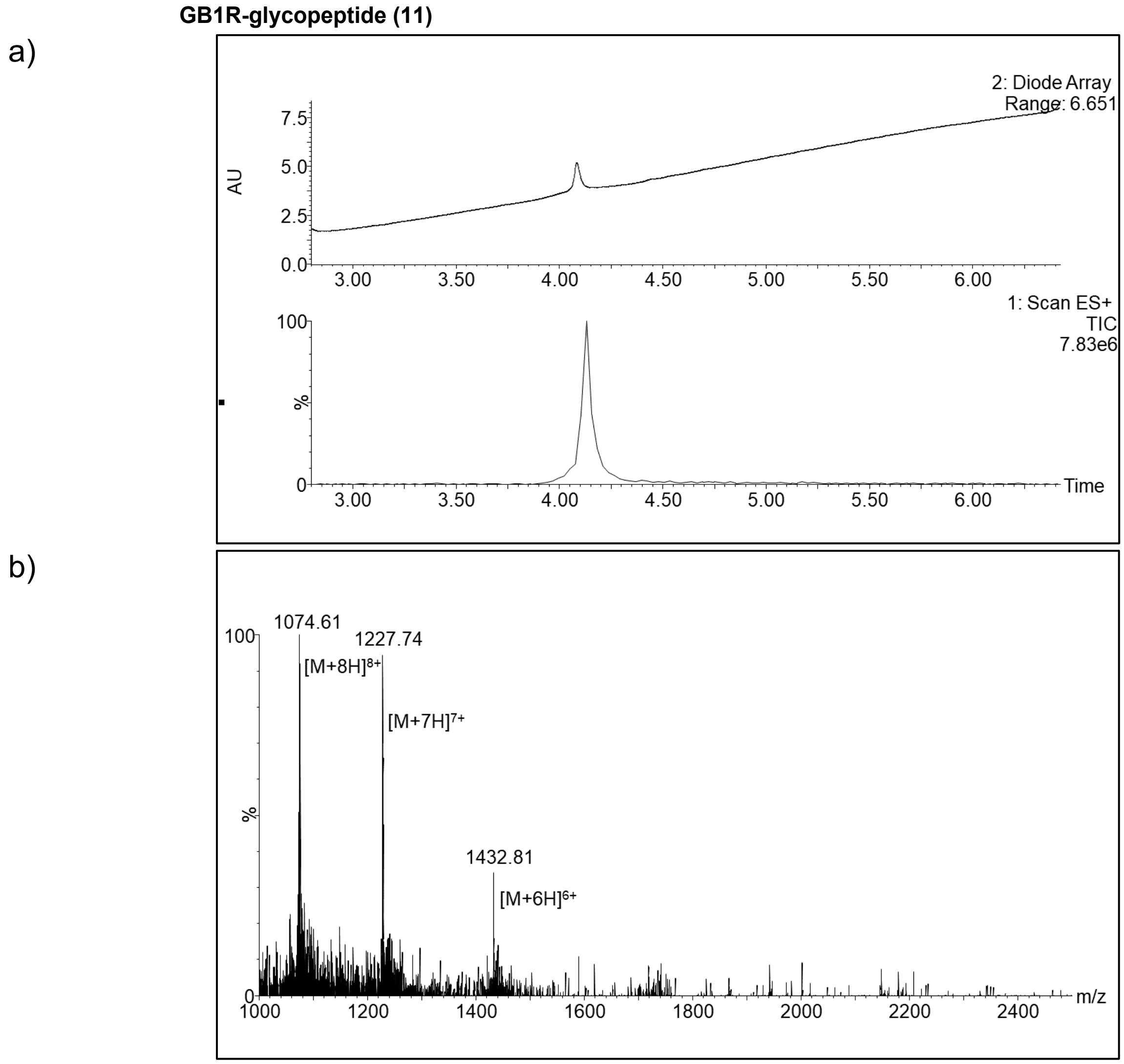

**Figure S43**. LC-MS of **GB1R-glycopeptide (11)**. a) UV and ESI+ TIC chromatogram of GB1R-glycopeptide, b) ESI+ MS of GB1R-glycopeptide.

(LC-MS method: 1% A to 5% B over 1 minute then 5%A to 60%B gradient over 10 minutes, solvent A was water/0.07% formic acid and solvent B was acetonitrile/0.07% formic acid, column: Acquity UPLC@Protein BEH C4, 2.1 mm * 150 mm, 1.7 µm; flow rate: 0.3 mL per minute.)

*4. GB1R-Q-non-glycosylated Peptide (**12**)*

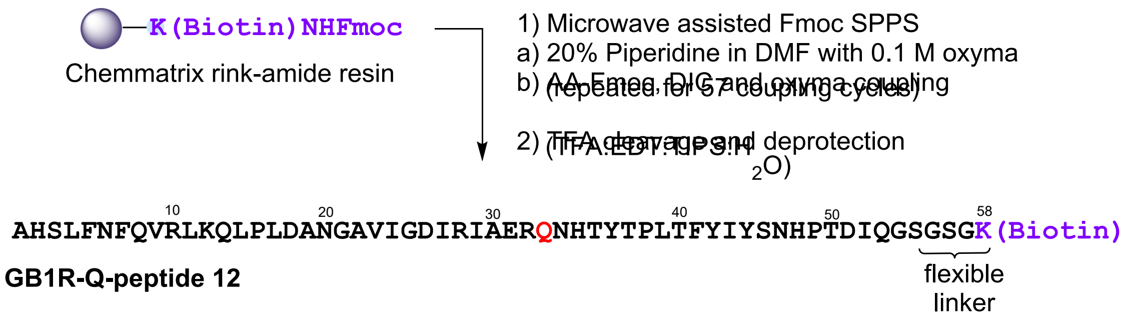

**Scheme S14.** Synthetic scheme to generate **GB1R-Q-non-glycosylated peptide (12).**

**GB1R-Q-peptide (12):** GB1R-Qpeptide (**12**) was prepared as in the representative procedure for solid phase peptide synthesis (**Section III.A.4**), starting with 150 mg of resin loaded at 0.17 mmol/g (25 µmol). Of the crude solid peptide, 40 mg was subjected to purification by RP-HPLC on the 10 x 250 mm C4 column (5-45% B over 45 min). 2 mg of pure peptide was obtained (5 % of the material that was purified). ESI-LRMS observed *m/z* of multiply charged ions 836.25 $[M+8H]^{8+}$, 958.00 $[M+7H]^{7+}$, 1117.38 $[M+6H]^{6+}$, 1340.73 $[M+5H]^{5+}$, 1675.43 $[M+4H]^{4+}$; calculated average m/z values for GB1R-Q peptide $C_{300}H_{462}N_{86}O_{87}S_1$ : 838.19 $[M+8H]^{8+}$, 957.79 $[M+7H]^{7+}$, 1117.25 $[M+6H]^{6+}$, 1340.50 $[M+5H]^{5+}$, 1675.38 $[M+4H]^{4+}$.

AHSLFNFQVRLKQLPLDANGAVIGDIRIAERQNHTYTPLTFYIYSNHPTDIQGSGSGK(Biotin)

**GB1R-Q-peptide 12**

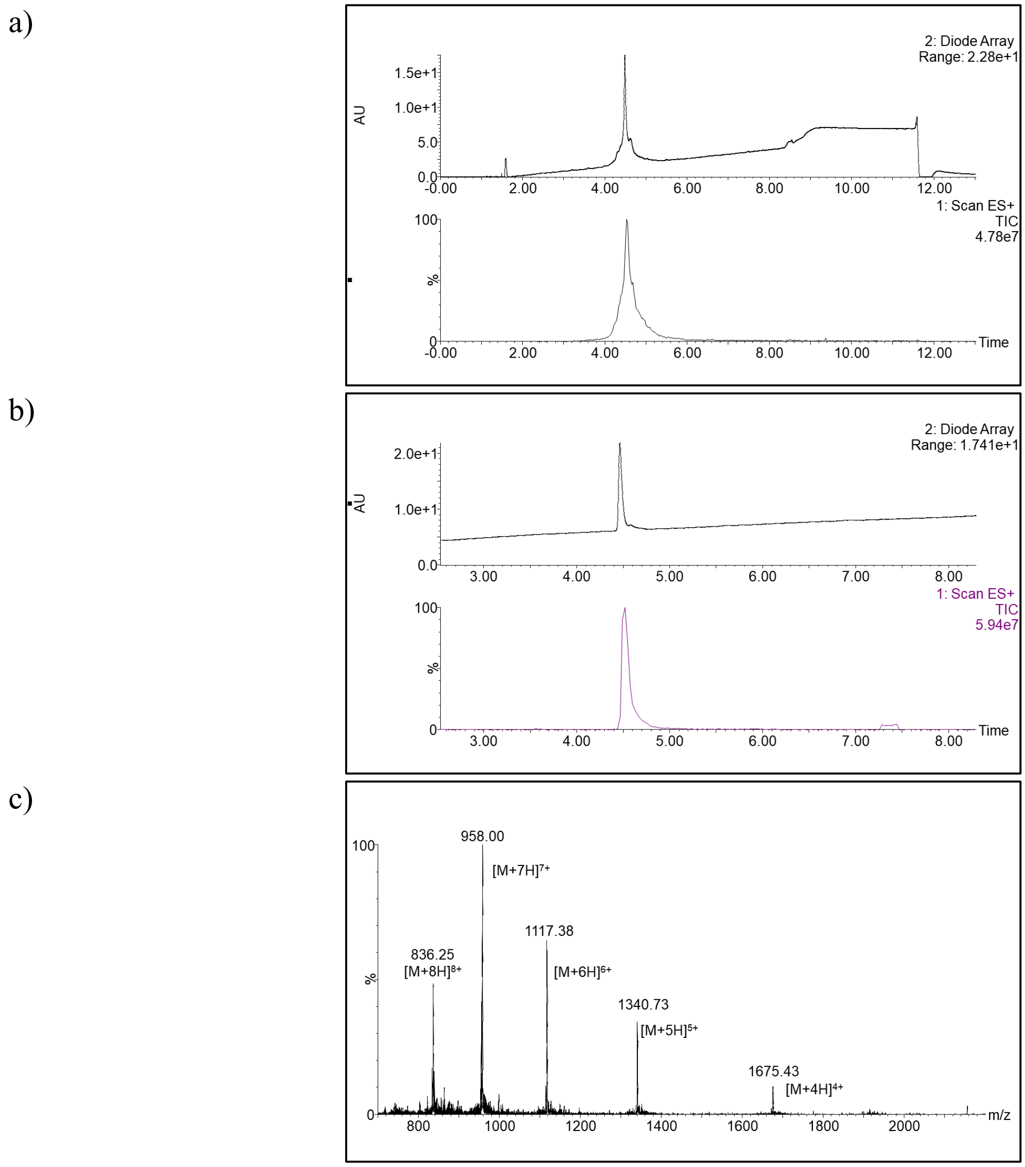

**Figure S44**. LC-MS of **GB1R-Q-non-glycosylated peptide (12)**. a) UV and ESI+ TIC chromatogram of GB1R-Q-non-glycosylated-peptide (crude), b) UV and ESI+ TIC chromatogram of GB1R-Q-non-glycosylated-peptide, c) ESI+ MS of GB1R-Q-non-glycosylated peptide.

(LC-MS method: 1% A to 5% B over 1 minute then 5%A to 60%B gradient over 7 minutes, solvent A was water/0.07% formic acid and solvent B was acetonitrile/0.07% formic acid, column: Acquity UPLC@Protein BEH C4, 2.1 mm * 150 mm, 1.7 µm; flow rate: 0.25 mL per minute.)

**B. BLI Sensorgram of GB1R-glycopeptide with gl-PGT121 versus PGT122**

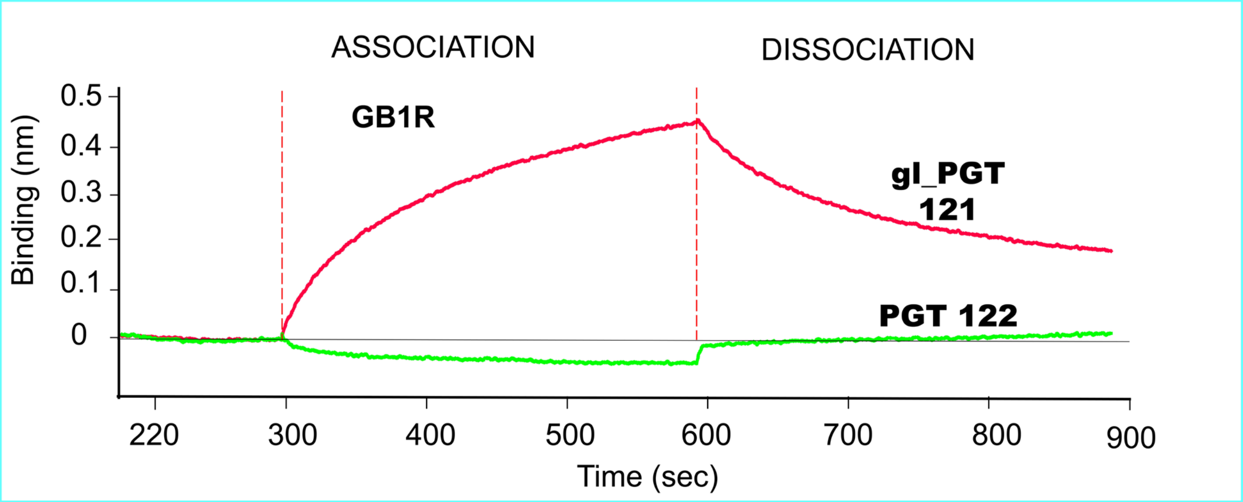

**Figure S45.** BLI Sensorgram of GB1R-glycopeptide-biotin immobilized on a streptavidin biosensor, treated with 2048 nM gl-PGT121 or mature PGT122 according to the procedure in **Section III.C.1**.
